# Heterogeneous Instructive Signals Enable Ensemble Learning in Cerebellar Cortex

**DOI:** 10.64898/2026.05.06.723116

**Authors:** Benjamin S. Ruben, Cengiz Pehlevan

## Abstract

The Marr–Albus framework describes learning in Purkinje cells (PCs) through supervised plasticity associated with complex spikes. However, complex spikes occur with low probability in individual PCs following sensorimotor events, and PCs are organized into “microzones” with shared complex spike tuning and outputs. We unify these observations in a model of associative learning in PC *ensembles* trained under “heterogeneous plasticity,” where each PC randomly receives either depression *or* potentiation during learning. Heterogeneous plasticity critically aids ensemble performance in a pattern recognition task, allowing memorization capacity to scale with ensemble size. An optimal-decoding theory suggests productive roles for recurrent PC–PC dynamics and nonlinear integration in the cerebellar nuclei. We account for additional experimental observations, including bursting and pausing PCs, a bias toward “upbound” or “downbound” learning mechanisms, and correlations in PC physiology and synaptic weights. Together, our results suggest that microzones perform ensemble learning, enabled by heterogeneous plasticity and nonlinear information processing.

## 1 Introduction

The ability to learn associations is an essential function of the nervous system, permitting organisms to adapt their responses to sensory information based on prior experience. The cerebellum is an important locus of associative learning in the brain, essential for classical eyeblink conditioning [1–4], skilled reach adaptation [5], and saccade adaptation [6–11]. Elucidating the computational principles behind associative learning in the cerebellum [12] is therefore essential for understanding how experience reshapes neural circuit function to support adaptive behavior.

In the canonical cerebellar circuit, mossy fibers carry sensorimotor inputs into a massively expanded granule cell (GrC) layer. The GrC axons, known as parallel fibers (PFs), then make excitatory synapses onto Purkinje cells (PCs), the sole output neurons of the cerebellar cortex (fig. 1A). PC dendrites also receive climbing fiber (CF) inputs from the inferior olive (IO), which initiate complex spikes (CSs) and an influx of dendritic calcium associated with the induction of long-term depression (LTD) at the PF–PC synapse when paired with PF stimulation (fig. 1B) [13].

**Figure 1:**
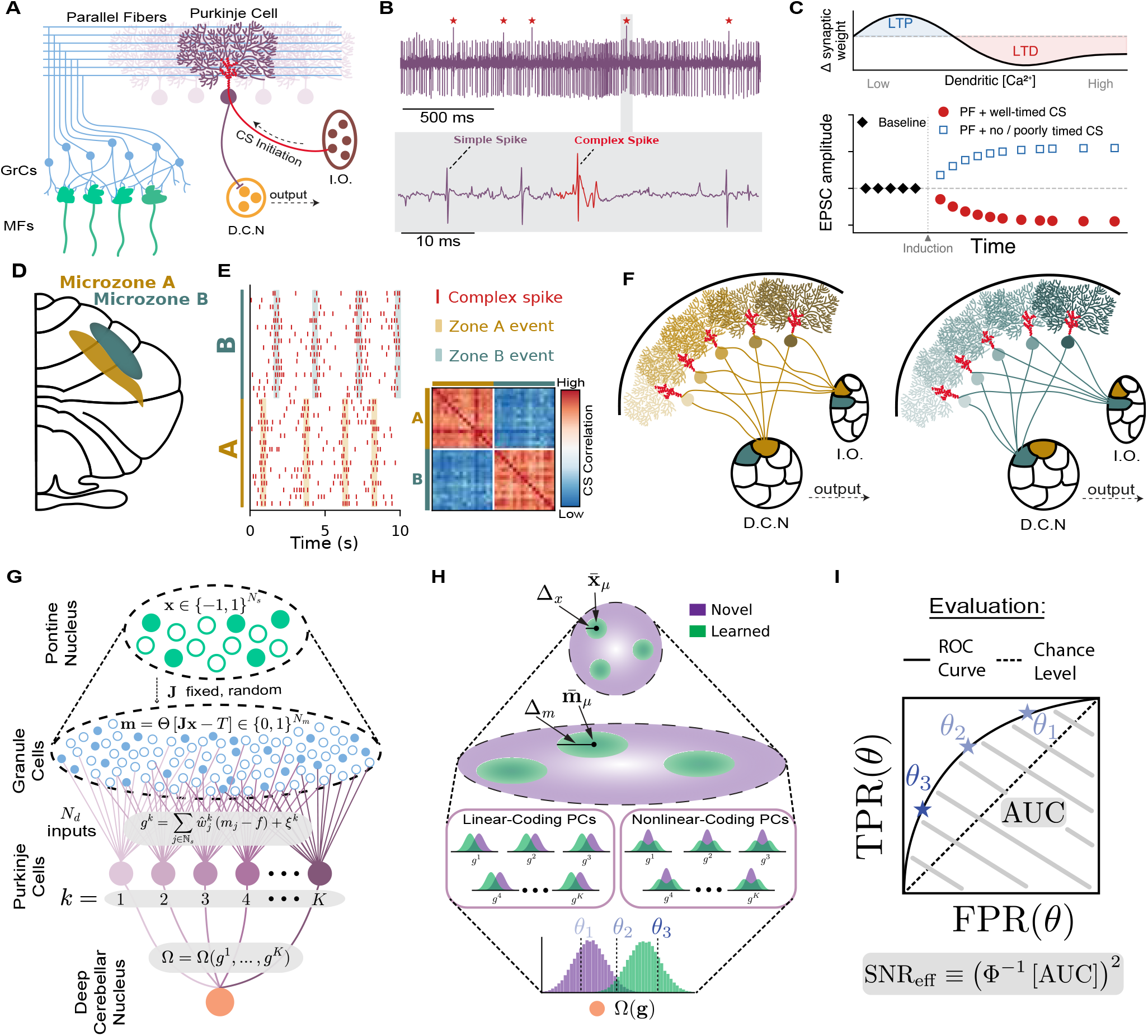
Background and model overview. **(A)** Cartoon depiction of a single operational unit of the cerebellar cortex (MF = mossy fiber, GrC = granule cell). Climbing fibers (CFs) from the Inferior Olive (I.O.) initiate complex spikes in PCs, and PCs project inhibitory axons to the Deep Cerebellar Nuclei (DCN). **(B)** Extracellular PC recording data from the P-sort dataset [49]. Stars mark expert-identified complex spikes. **(C)** Calcium-dependent bidirectional plasticity at the PF–PC synapse: Schematic summary of a calcium-threshold rule for parallel fiber (PF) to Purkinje cell (PC) plasticity. Lower dendritic calcium elevations favor long-term potentiation (LTP), whereas larger and/or more prolonged calcium elevations favor long-term depression (LTD). In this schematic, PF activation paired with a well-timed or otherwise effective complex-spike signal drives calcium into the LTD regime, whereas PF activation with no complex spike, or with a weak or poorly timed complex-spike signal, remains in the lower-calcium regime and favors potentiation [40, 42]. **(D)** Diagram of the cerebellar cortical surface, showing two adjacent microzones A and B (not drawn to scale). **(E)** Synthetic complex spike rasters demonstrating that sensorimotor events are encoded in population-level complex spiking events. CS correlation heatmap demonstrates that zones may be defined by enhanced correlations in complex spike trains across PCs within a zone. **(F)** A cerebellar “micro-complex” consists of a cerebellar microzone, as well as the restricted zones of the inferior olive and DCN which provide inputs to and receive projections from the microzone. **(G)** Overview of model architecture. **(H)** Overview of model associative learning task and possible PC coding schemes. **(I)** Model evaluation metric using the area under the ROC curve (AUC), and definition of SNR_eff_.

The seminal works of Marr, Albus, and Ito proposed that climbing fiber activity provides a supervised teaching signal which instructs plasticity in PCs [14–16] (fig. 1C). In this framework, coincident activation of PF inputs and CF-driven CSs modifies the PF–PC synapses associated with the preceding sensorimotor context, allowing instructive signals to be assigned to the inputs that predicted them. Through these synaptic updates, the cerebellar cortex can transform mossy-fiber representations of sensory and motor state into learned changes in PC output that support adaptive behavior.

While highly influential, these models and subsequent theoretical treatments [17–19] largely formulate cerebellar learning at the level of individual PCs, an assumption that must be reconsidered in light of recent experimental evidence. PCs are organized into sagittal stripes called “microzones,” (fig. 1D) which may be defined functionally through shared complex spike tuning [20, 6, 21] (fig. 1E) or anatomically by the topography of their inferior olive inputs and targets in the deep cerebellar nuclei (DCN) [22–25] (fig. 1F). Histochemical methods also reveal a striped zonal organization [26], which aligns with topographic and functional subdivisions [27–30]. The cerebellar “micro-complex,” consisting of a microzone and its mutually connected subdivisions of the IO and DCN (fig. 1F), has therefore been proposed as a functional computational unit of the cerebellum [31, 32, 9]. Consistent with this view, growing experimental evidence suggests cerebellar computation *must* be understood as a population-level phenomenon:

- *PCs represent task-relevant information in population-level simple spike activity statistics:* During saccades, different PCs within a functional ensemble undergo transient bursts *or* pauses, but a signal tracking eye velocity can be recovered from their population-averaged firing rates [6]. Other experiments suggest population coding through synchronization of PC simple spikes: synchronous firing of only two PCs can elicit time-locked spiking in their shared DCN target neuron [33], and reduced PC synchrony is associated with motor deficits in tottering mice [34].
- *Individual sensorimotor events evoke unreliable plasticity outcomes in individual PCs:* In recordings from single PCs, CSs fire with low probability following individual sensorimotor events including facial touch, movement, changes in sensory load, and visual feedback errors, with typical single-trial CS firing probabilities below 0.3 [35–39, 6–8, 11]. Slice experiments show that PF stimulation without a paired CS induces long-term potentiation (LTP) at PF-PC synapses [40], with more recent results suggesting that CS-paired PF stimulation results in either LTD *or* LTP depending on the phase of IO subthreshold oscillations at the time of CS induction [41] or the relative timing of consecutive CSs [42]. Consistent with these findings, Herzfeld et al. [7] elegantly demonstrate that plasticity outcomes in individual PCs following identical sensorimotor errors vary from trial to trial *in vivo*, in a manner dependent on the presence and timing of complex spikes. Reliable learning is therefore unlikely to be explained by plasticity in isolated PCs alone, and likely requires population-level integration across the attuned microzone, where sensorimotor events are encoded reliably in collective CS activity [20].
- *PCs have recurrent and correlated connectivity:* PCs share extensive recurrent connections with each other both directly and through inhibitory interneurons of the molecular layer [43– 45], providing a substrate for intra-zone communication between PCs that may aid population-level learning and computation. Anatomical reconstructions also suggest correlations in patterns of synaptic connectivity from PFs to PCs which may aid an ensemble learning calculation [46].

Motivated by these findings, we present here *a model of cerebellar learning in a functional PC ensemble converging on shared DCN output*, corresponding to a portion of a microzone receiving shared mossy fiber inputs (fig. 1G). We contrast ensemble performance under a “homogeneous plasticity” learning rule in which PCs receive identical error signals, and a “heterogeneous plasticity” learning rule in which PCs randomly undergo either depression or potentiation during learning. Unreliable plasticity outcomes in individual PCs might naively be considered a nuisance to be overcome by combining the outputs of many PCs. To the contrary, we find that the heterogeneous plasticity learning rule critically *aids* network performance in an associative learning task by reducing cross-talk correlations between the patterns stored in network weights. Following training with the heterogeneous plasticity learning rule, PC activity represents stimulus identity in the distribution of PC population activity, rather than just the mean, therefore behaving as a “nonlinear population code” [47]. When paired with a suitable nonlinear decoding mechanism, heterogeneous plasticity permits the ensemble’s memorization capacity to scale supra-linearly with network width, surpassing the linear scaling of single-PC theories [17, 18], or ensembles trained under homogeneous plasticity. Furthermore, heterogeneous plasticity provides a unified explanation of many experimental phenomena, including bursting and pausing PC dynamics, upbound and downbound learning biases [48], weakly synchronous simple spikes, and weakly correlated PF–PC connectivity. Together, our results provide an updated account of cerebellar learning in which robust associative learning arises from population-level plasticity events and computation in PC ensembles. Furthermore, our analysis identifies heterogeneous plasticity and nonlinear information processing as the critical features enabling this population-level computation.

## 2 Results

### 2.1 A Model of Associative Learning in PC Ensembles

We build on previous models of associative learning in cerebellum-like architectures [17, 18, 50, 19], extending them from a single readout neuron to a population of PCs converging on a common nuclear target (fig. 1G). We consider an associative learning task in which the network must respond selectively to stimuli from *P* clusters in sensory input space [17] (fig. 1H). Binary stimuli 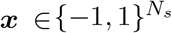 represent the activity of *N*_*s*_ pontine nucleus neurons. Each cluster *µ* = 1, …, *P* has a random, fixed center 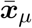, and test patterns from this cluster are generated by flipping each element of 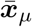 with probability Δ_*x*_*/*2. The network must respond preferentially to learned patterns while remaining quiescent to novel patterns which are drawn uniformly at random from 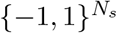.

Following the canonical cerebellar architecture, inputs are first randomly expanded into a mixed-layer representation of *N*_*m*_ ≫ *N*_*s*_ binary granule cells, ***m***(***x***) = Θ[***Jx*** − *T*], where ***J*** is a fixed, random weight matrix and *T* is a sparsifying threshold on GrC activity. In a seminal work, Babadi and Sompolinsky [17] showed that the effect of the sparse expansion is captured by two summary statistics: the cluster sizes Δ_*m*_ following projection to the granule cell layer, and the *effective dimensionality N*_*eff*_ of the PF representations (Methods §3). Here, all granule cells receive fixed, random inputs from a single shared population of pontine nucleus neurons. Microzones, however, often span multiple cerebellar lobules, presumably each representing distinct sensory information. We consider our model to apply to a portion of a microzone which receives inputs from a shared population of mossy fibers.

A layer of *K* PCs then integrates PF activity through learned weights 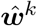, producing an input current

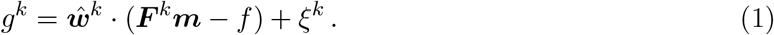

Here, ***F*** ^*k*^ is a binary connectivity mask which selects a random set of *N*_*d*_ PFs “accessible” to PC *k*, which we assume to be sampled randomly (Methods §3), and 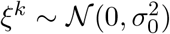 represents intrinsic noise sampled independently across PCs. The centering *f* is the average granule cell activation, which is set by the threshold *T* (Methods §3). Negative weights implicitly capture feedforward inhibition via molecular layer interneurons [43].

Training proceeds online: upon presentation of each cluster center 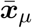, synaptic weights are updated by a two-factor rule

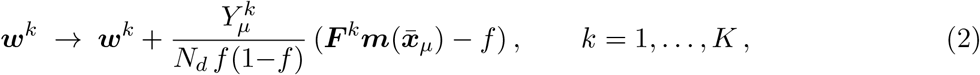

where 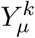 is an *effective plasticity signal* conveyed by the presence/timing of the climbing fiber input to PC *k*. In SI §S11, we justify these centered weight updates as the outcome of an uncentered instructed plasticity update 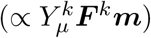 followed by centering homeostatic plasticity.

The final output of the circuit is a DCN neuron that receives convergent input from all *K* PCs. We denote this output Ω(***g***), leaving the transformation from PC inputs to DCN activity in general form – we will later study many possible forms for Ω(***g***), including forms derived from optimization principles. Predicted class *ê*(***x***) is assigned to input ***x*** by comparing the network output Ω(***g***) to a threshold *θ* (fig. 1H), and assigning label *ê* = 1 if Ω(***g***) ≥ *θ*. As the threshold *θ* varies, the false-positive rate FPR(*θ*) and true-positive rate TPR(*θ*) trace out the Receiver Operating Characteristic (ROC) curve (fig. 1I). The standard metric to summarize decoder performance across threshold choice is the area under the ROC curve (AUC). We further define an effective signal-to-noise ratio SNR_eff_ = Φ^−1^ AUC ^2^, where Φ is the standard Gaussian CDF. In the special case of Gaussian-distributed network outputs, SNR_eff_ is equivalent to the standard [51–53, 17, 18] SNR (Methods §3). We therefore consider SNR_eff_ to be a generalization of SNR which allows comparisons across cases with Gaussian and non-Gaussian output statistics, and refer to SNR_eff_ simply as SNR for the remainder of the paper.

### 2.2 Ensembles Yield Limited Benefits under Homogeneous Instruction

Effective ensemble learning requires predictors to learn *different* functions of the input, so that their combined prediction achieves a significant benefit over any individual [54]. One compelling possibility is that PC ensemble diversity is inherited from anatomical constraints. Specifically, PCs receive excitatory synapses from roughly half of the PFs which pass through their dendritic arbor, so that each PC receives a different set of inputs [46]. We may encode this in our model by setting *N*_*d*_ = *N*_*m*_*/*2. To isolate the contribution of sparse PF–PC connectivity, we first train PCs under the “homogeneous LTD” learning rule, setting

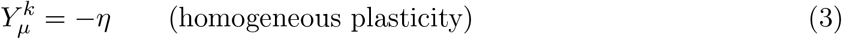

in the two-factor learning rule (eq. 2). Following training, PC response histograms for novel patterns (*P* (***g***|*e* = 0)) and learned patterns (*P* (***g***|*e* = 1)) are then Gaussians distinguished only by a downward mean shift 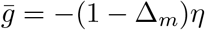 upon presentation of a learned stimulus (fig. 2B). With the plasticity signals constant across *k*, the only sources of ensemble diversity remaining are the random connectivity of PCs to PFs and intrinsically generated fluctuations of variance 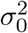. To evaluate their effect on ensemble performance, we must first select the transformation Ω(***g***) used to obtain network output from learned PC input currents. The choice of decoder which maximizes AUC (and therefore SNR) is given by the Neyman-Pearson test, which is simply the likelihood ratio between the classes:

**Figure 2:**
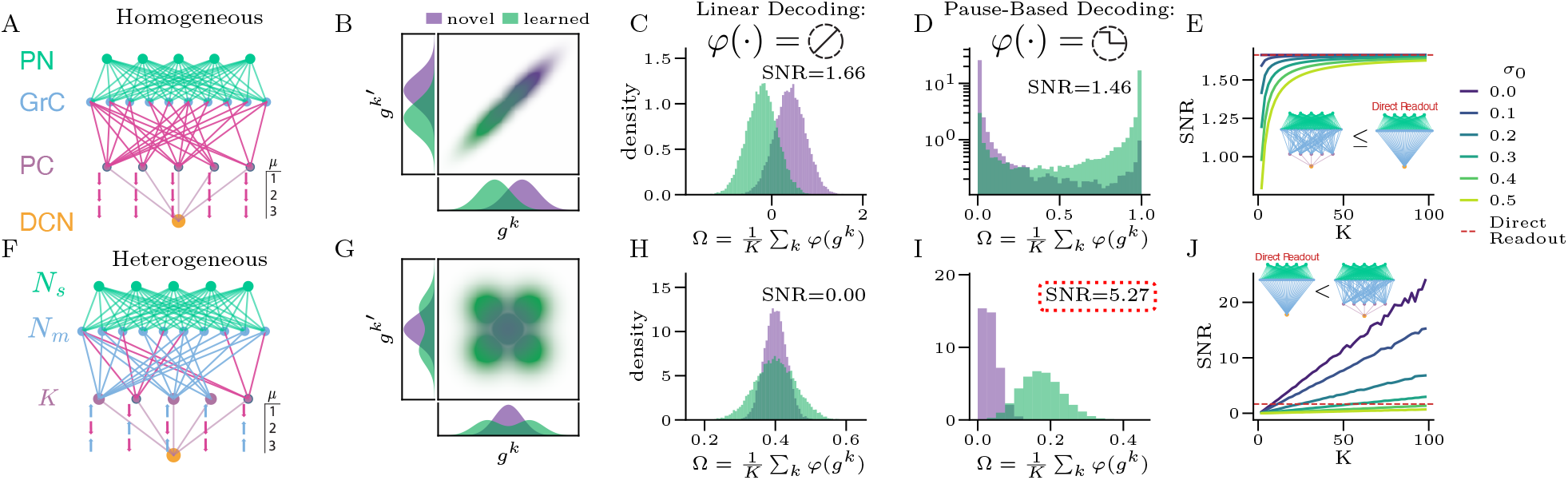
Homogeneous Plasticity Limits Ensemble Performance, while Heterogeneous Plasticity Permits SNR improvements. **Top row:** Homogeneous LTD learning rule. **(A)** Schematic: all PCs undergo LTD during every learning event. **(B)** Joint distribution of PC inputs (*g^k^, g^k^′*) for a representative pair after training. Novel (purple) and learned (green) distributions differ only in their mean, and fluctuations are strongly correlated across PCs. **(C)** Output histograms under linear decoding *φ*(*g*) = *g*. The two classes are separated (SNR = 1.66). **(D)** Output histograms under pause-based decoding *φ*(*g*) = Θ(*κ* − *g*). Nonlinear decoding does not improve separation (SNR = 1.46). **(E)** SNR vs. ensemble size *K* for varying intrinsic noise *σ*_0_. All curves saturate at the direct readout limit (red dashed line; architecture schematized in inset). **Bottom row:** Heterogeneous plasticity learning rule. **(F)** Schematic: each PC independently undergoes LTD (probability *β*) or LTP (probability 1 − *β*) during each learning event. **(G)** Joint distribution of (*g^k^, g^k^′*) after training. PC responses are uncorrelated, and the learned distribution differs from the novel distribution primarily in its variance. **(H)** Linear decoding fails entirely (SNR = 0.00) as mean PC input is identical for the novel and learned classes under balanced plasticity. **(I)** Pause-based decoding recovers a strong signal (SNR = 5.27), vastly exceeding the direct readout limit. **(J)** SNR vs. *K* under pause-based decoding. Performance grows with *K* and surpasses the direct readout limit (red dashed line), in contrast to the homogeneous case (E). Parameters: *K* = 40, *P* = 3,000, *N*_s_ = 1,400, *N*_m_ = 400,000, *N*_d_ = 200,000, *f* = 0.05, Δ_x_ = 0.10, *σ*_0_ = 0.1. Homogeneous: *η*_*−*_ = 1. Heterogeneous: *β* = 0.5, *η*_+_ = *η*_*−*_ = 0.5.

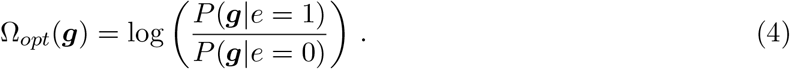

In the case of homogeneous plasticity learning, this reduces to a linear average 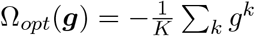 (Methods §3), as has been found in similar settings [55, 53]. With this linear decoder, an analytical form for the SNR is then easily obtained as

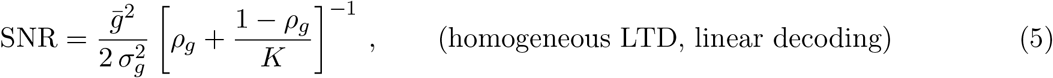

where *ρ*_*g*_ ∈ [0, 1] represents the correlation coefficient and 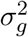 represents the marginal variance of learned PC currents ***g***^*k*^ (full expressions in Methods §3). Because the linear decoder is optimal, eq. 5 is an upper bound on SNR under homogeneous plasticity for all possible decoders Ω(***g***) (fig. 2C,D, fig. S1). Despite integrating different sets of PFs, shared plasticity schedules result in heavily correlated PC input currents across PCs (fig. 2B), which heavily limits ensemble performance. A simple analysis of eq. 5 shows that SNR is monotonically increasing with *N*_*d*_ and decreasing with *σ*_0_, reaching optimal performance when *N*_*d*_ = *N*_*m*_ and *σ*_0_ = 0 (see §S4.2 and fig. S1). At this point, the ensemble reduces effectively to a “direct readout” architecture in which all PFs connect directly to a single output neuron (fig. 2E). When subsampling or intrinsic noise are present *a priori*, increasing PC ensemble size may improve network performance, but only up to a plateau at the SNR of the direct readout architecture (fig. 2E). Under homogeneous plasticity, we therefore find no computational rationale for the existence of the PC layer, as performance never surpasses that of a network in which the PC layer is absent. Furthermore, with realistic parameters (*N*_*s*_ ≈ 1,400, *N*_*d*_ ≈ 200,000, *N*_*m*_ ≈ 400,000, *σ*_0_ ≈ 0 [56], *f* ≈ 0.05), PC fluctuations have strong correlations (*ρ*_*g*_ ≳ 0.8), so that increasing *K* achieves rapidly diminishing SNR improvements (see figs. 2E and S1).

### 2.3 Heterogeneous Plasticity Permits SNR Improvements

Having observed limited benefits from anatomical variations, we next turn to physiology as a source of ensemble diversity. Specifically, CSs are not perfectly synchronized within a functional zone, but rather form heterogeneous patterns [20], and individual PCs may undergo either LTP or LTD depending on the presence and precise timing of CSs following individual sensory events [40, 41, 7, 42]. As a minimal abstraction of these experimental facts, we introduce the *heterogeneous plasticity* learning rule:

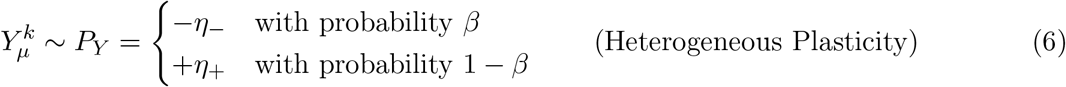

Here, *P*_*Y*_ represents the distribution from which each 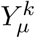 is independently sampled during training (fig. 2F). With *P*_*Y*_ as in eq. 6, *β* corresponds to the probability of a “well-timed” complex spike following presentation of stimulus 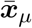 during one-shot learning. The essential results presented here are not sensitive to this exact form for *P*_*Y*_, but rather require that *P*_*Y*_ has nonzero variance. We discuss multi-shot learning and alternative forms of *P*_*Y*_ in the SI. From *P*_*Y*_, we define the mean 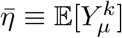, variance 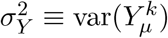, and *plasticity ratio*

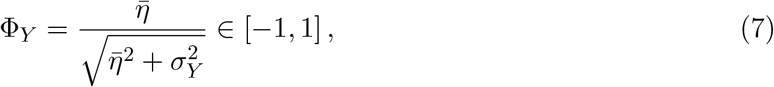

which represents the signed fraction of plasticity power due to its mean direction. Inspired by experimental observations [48], we refer to Φ_*Y*_ *>* 0 as “upbound” and Φ_*Y*_ *<* 0 as “downbound” plasticity, and to Φ_*Y*_ = 0 as “balanced” plasticity.

Heterogeneous plasticity radically changes the statistics of PC population responses. While PCs trained with homogeneous LTD distinguish novel and learned inputs only by a shift in their *mean*, PCs trained with heterogeneous plasticity distinguish learned and novel distributions primarily by increased *variance* across PCs upon presentation of a learned stimulus (fig. 2G). This is precisely the distinction between what have been named *linear* and *nonlinear* population codes [47]: while linear codes represent stimulus identity in mean population activity, nonlinear codes also use variance and higher moments. Furthermore, heterogeneous plasticity results in pairwise correlations that are radically reduced relative to the homogeneous plasticity case (fig. 2G), creating the potential for meaningful benefits from ensembling.

However, extracting the information stored in a nonlinear population code requires a nonlinear decoding mechanism. Under balanced heterogeneous plasticity, linear decoding fails entirely as the mean signal averages to zero (fig. 2H). In fact, under linear decoding, we find that SNR is optimized by setting *β* = 0 or *β* = 1, recovering homogeneous plasticity (Methods § 3). A nonlinear decoder is therefore necessary for heterogeneous plasticity to potentially outperform homogeneous plasticity.

Experimentally, PCs develop “pause responses” to conditioned stimuli in eyeblink conditioning [57, 58], and transient PC pauses precede saccade initiation [59]. This motivates us to consider a decoder of the form Ω(***g***) = ∑_*k*_ Θ[*κ* − *g*^*k*^], in which PCs “pause” if their input current drops below a threshold *κ*, and the DCN is disinhibited in proportion to the number of pausing PCs. Combining heterogeneous plasticity with pause-based decoding yields a dramatic improvement in SNR, far surpassing the direct readout limit (fig. 2I,J). Crucially, *both* heterogeneous plasticity and nonlinear decoding are necessary: using either alone results in worse performance than the direct readout architecture. We also find that the benefits of heterogeneous plasticity are not specific to the pause-based decoder, extending to all tested nonlinearities (fig. S2).

### 2.4 Capacity Scaling under Heterogeneous Plasticity Learning

So far, we have evaluated network performance using the SNR, which measures memorization performance for a fixed number of stimulus clusters *P*. We may also quantify performance using the capacity *P*_max_, which we define here as the largest *P* for which 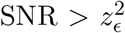 (for some constant tolerance *z*_*ϵ*_, Methods §3). Marr [14]’s seminal work argued that re-coding MF inputs in sparse GrC population activity permitted the memorization capacity of a single PC to scale with the size of the GrC layer rather than the number of MF inputs. For a single PC with *σ*_0_ = 0 and trained with the standard LTD learning rule (or equivalently for the direct readout architecture), we find 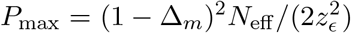, where *N*_eff_ is the effective dimensionality of GrC population activity (Methods §3). Provided GrC population activity is sufficiently sparse (i.e. 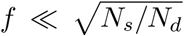), we have *N*_eff_ ≈ *N*_*d*_, so that capacity scales linearly with the width of the GrC layer, in line with Marr [14]’s original insight. While the sparsity of GrC population activity has been debated in the literature [60–63], capacity improvement from sparse recoding of MF inputs remains the prevailing computational rationale for the massive expansion from MF to GrC layers [64].

In fig. 3, we study network capacity under balanced, heterogeneous plasticity learning 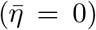, beginning with the pause-based decoding strategy introduced in fig. 2. For simplicity, we assume fully-connected PCs (*N*_*d*_ = *N*_*m*_). Fig. 3A shows AUC as a colormap in the space of ensemble size *K* and cluster number *P* for a pause-based decoder with *β* = 0.1. At each (*K, P*) pair, we separately optimize the pause threshold *κ* (fig. 3B). As PC ensemble size *K* grows, the optimal *β* value scales down so that a smaller fraction of PCs undergo LTD during each plasticity event (fig. 3C). Scaling network size, we therefore converge on a “divide and conquer” strategy, in which network capacity is improved by assigning each input cluster to a sparse subset of PCs, reducing the load on any one PC. The pause-based decoder then distinguishes inputs from learned clusters by the presence of a sparse subset of pausing PCs. When both *β* and *κ* are jointly tuned to their optimal values and *σ*_0_ = 0, we find that network capacity scales as *P*_max_ ∝ *KN*_eff_ */* log(*K*) (fig. 3G,H), which is within a logarithmic factor of the fastest possible scaling for a noiseless two-hidden-layer network [65, 66]. In the case where the size of the Granule Cell layer and the number of PCs are scaled together as *N*_*m*_, *K* ∝ *w*, we obtain *P*_max_ ∝ *w*^2^*/* log(*w*), so that capacity scales near-quadratically with the “width” parameter *w*, beating the linear width-scaling of a single-PC model (fig. 3I).

**Figure 3:**
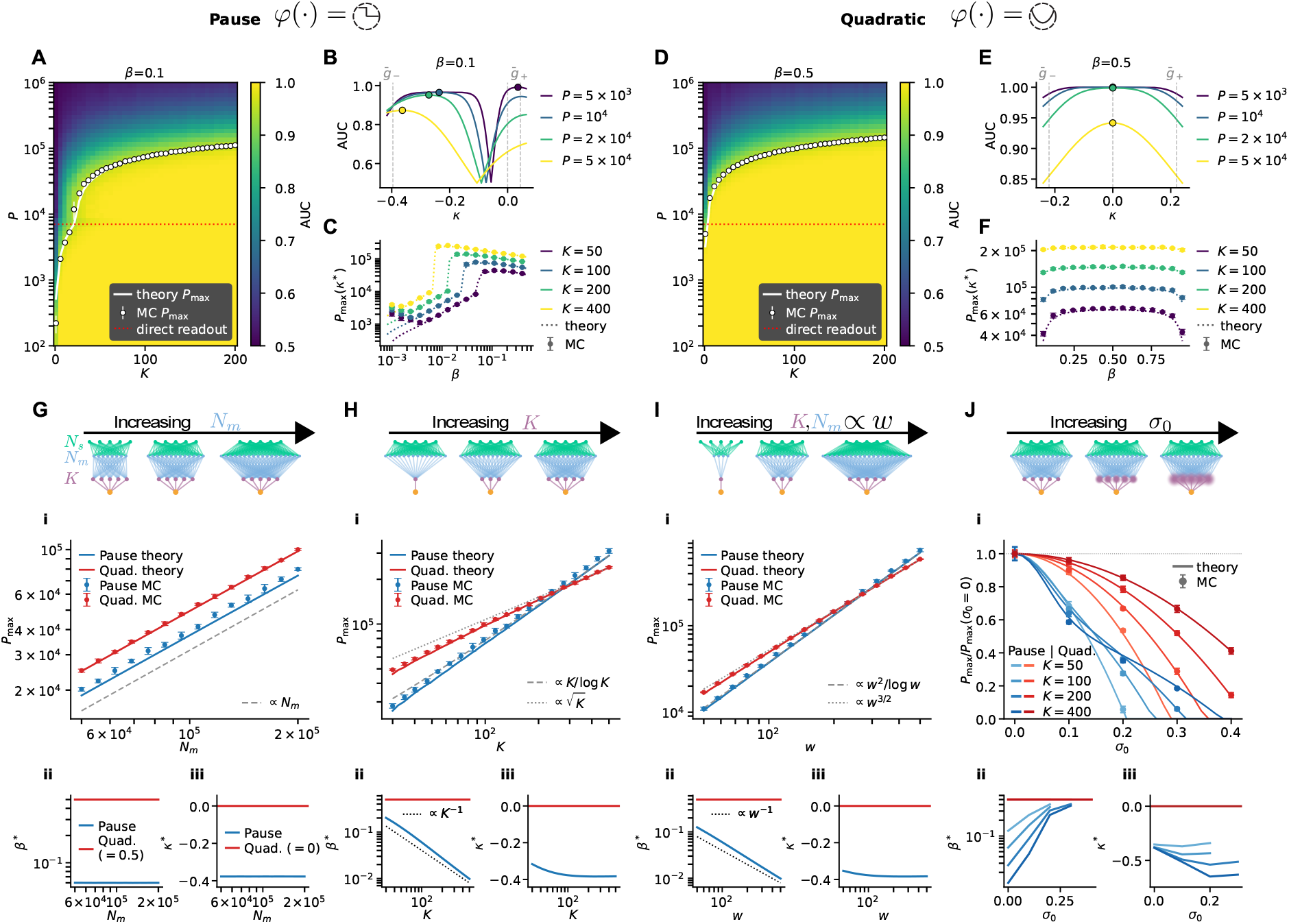
Heterogeneous Plasticity Enables Capacity Scaling. We evaluate the capacity of the model network after training under balanced plasticity (*η*_−_ = 1 − *β, η*+ = *β*) and separable decoder Ω(***g***) = ∑_k_ *φ*(*g^k^*), comparing “pausing” (A–C) and quadratic (D–F) nonlinearities. (A) Heatmaps of AUC in the (*K, P*) plane with *β* = 0.1 and “pausing” nonlinearity *φ*(*g*) = Θ[*κ − g*]. At each grid point, we determine the optimal threshold *κ*^∗^ using the analytical AUC estimate, then calculate AUC at *β* = 0.1, *κ*^∗^ using MC sampling (Methods). Overlaid curves show the analytical capacity estimate and capacity of the direct readout architecture, and markers show capacity determined from MC sampling. (B) Analytical estimate of AUC as a function of pause threshold *κ* with *β* = 0.1 at various *P* values. Vertical dotted lines show 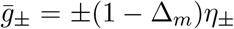 (C) Capacity *P*_max_ at optimal threshold *κ*^∗^ as a function of LTD fraction *β* at various ensemble sizes *K*. (D–F) Same as A–C with *β* = 0.5 and quadratic decoder *φ*(*g*) = (*g − κ*)^2^. At all *K*, we find *β*^∗^ = 0.5 and *κ*^∗^ = 0. (G–J) We measure capacity as a function of various model parameters with both the pausing and quadratic nonlinearities, with optimized parameters *β*^∗^, *κ*^∗^ tuned separately at each parameter value. Scaled parameters are: (G) GrC count *N*_m_; (H) PC ensemble size *K*; (I) both GrC count *N*_m_ and PC ensemble size *K*, in proportion to a network “width” *w*, so that *K* = *w* and *N*_m_ = 10^3^*w*; and (J) intrinsic PC noise scale *σ*_0_. Sub-panels (i) show capacity *P*_max_ at optimized *β*^∗^ and *κ*^∗^. Panels (ii, iii) show the optimal *β*^∗^ and *κ*^∗^ values, respectively.

While some experiments suggest PCs encode information in brief pauses which dis-inhibit their DCN targets, other studies suggest several non-monotonic effects in the pathway from PC inputs to DCN outputs. For example, simulations suggest that transient PC pauses can be induced by strong depolarizing input [67], and strong DCN inhibition by PCs may result in rebound spiking [68]. We therefore examine network capacity with a quadratic decoder Ω(***g***) =∑ _*k*_(*g*^*k*^ − *κ*)^2^, which is sensitive to both increases and decreases in PC activity relative to the threshold *κ*. With the quadratic decoder, we find that, at all scales, capacity is optimized by setting *κ* = 0 and *β* = 0.5 (fig. 3D-F). An analytical estimate then gives that network capacity scales as 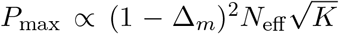 in the relevant regime where 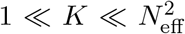 (fig. 3G, H), or as *P*_max_ ∝ *w*^3*/*2^ with network width *w* (fig. 3I). While capacity scales more slowly with PC ensemble size *K* than that attained using the pause-based decoder (and scaling *β* ∝ 1*/K*), it retains many favorable properties. First, capacity scaling as 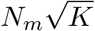 tracks more closely with cerebellar anatomy, as the GrC population dwarfs the PC population. If, instead, capacity scales as *N*_*m*_*K*, the optimal distribution of neurons would be an equal number of GrCs and PCs. Indeed, at realistic scales of *K* ≲ 200, the quadratic decoder with *β* = 0.5 has higher capacity than the pause-based decoder. Second, the quadratic decoder enjoys considerably better robustness to intrinsic noise at the PC soma (fig. 3J). Finally, the capacity scaling attained with the quadratic decoder extends to a multi-shot learning setting (see §S8).

### 2.5 Optimal Decoding of Learned PC Inputs through Cerebellar Dynamics

The linear, pause-based, and quadratic decoders studied above are “separable” decoders of the form Ω(***g***) = ∑_*k*_ *Φ*(*g*^*k*^), representing a narrow slice of all possible decoding strategies. We next study the form of the *optimal* decoder Ω_*opt*_(***g***) under heterogeneous plasticity training, again given by the Neyman-Pearson test (eq. 4). Under heterogeneous plasticity, the learned-class likelihood *P* (***g*** | *e*=1) is a mixture of 2^*K*^ Gaussians—one for each possible combination of LTD/LTP across the *K* PCs. This presents a computational challenge, as naively computing Ω_*opt*_ will require *O*(2^*K*^) computational steps, as well as a challenge to biological interpretability. In the SI, we show that *P* (***g*** | *e* = 1) can be recast as an integral over a single latent variable, evaluable in *O*(*K*) computational steps (Methods §3). Applying a saddle-point approximation to this integral representation then yields an interpretable two-step decoding circuit. In the first step, PC firing rates 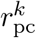 evolve until convergence under learned feedforward drive from PFs, recurrent PC-PC inhibition, and PC-PC dis-inhibition through an inhibitory Purkinje-layer interneuron (PLI) with firing rate *r*_*I*_ (fig. 4A):

**Figure 4:**
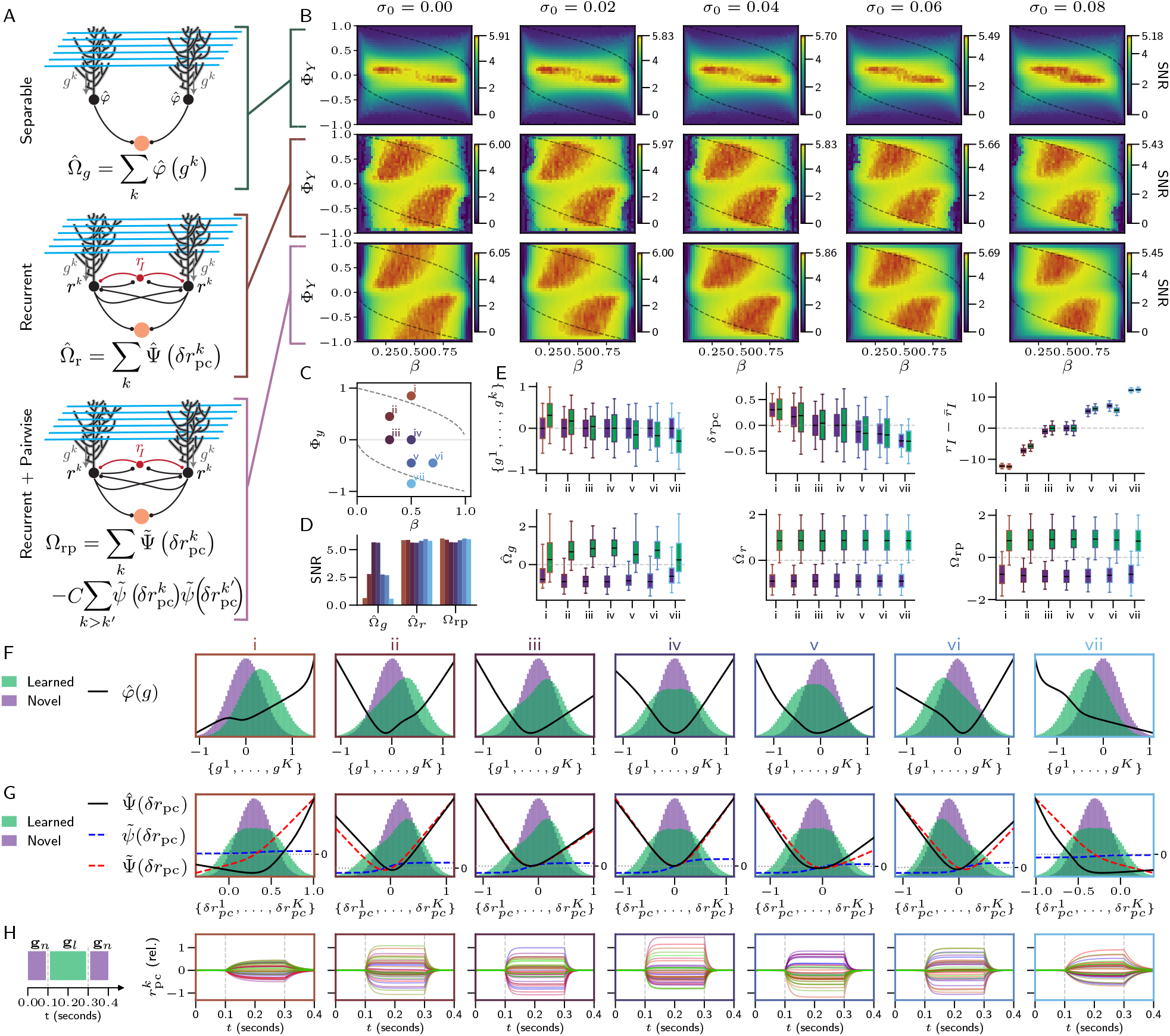
Optimal decoding under heterogeneous plasticity can be implemented through recurrent PC dynamics and nonlinear DCN integration. **(A)** Three decoding strategies of increasing complexity. Top: separable decoder with fitted elementwise nonlinearity. Center: recurrent inhibition between PCs and a PLI followed by a fitted nonlinearity applied to the fixed-point firing rates. Bottom: full saddle-point approximation to the optimal decoder including recurrent dynamics and pairwise interactions. **(B)** SNR heatmaps over plasticity parameters *β* and φ_Y_ (with |*η*_*−*_| + |*η*_+_| = 1) for each decoder and varying intrinsic noise *σ*_0_. Top 5% of each colormap shown in red. Two near-optimal peaks correspond to upbound and downbound regimes. Dashed lines separate the mixed-plasticity regime from pure-LTP 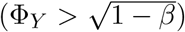 and pure-LTD 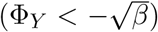. **(C)** Operating points i–vii used in subsequent panels. **(D)** SNR at the indicated operating points for the three decoder types (*σ*_0_ = 0.0). **(E)** Box plots of PC input currents *g^k^*, firing rate deviations 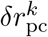, PLI activity 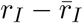, and decoder outputs for novel (purple) and learned (green) inputs at operating points i–vii. **(F)** Marginal distributions of 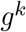 overlaid with fitted nonlinearity 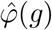. **(G)** Marginal distributions of fixed-point 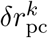 overlaid with nonlinearities from 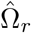 and Ω_rp_. **(H)** PC firing rate dynamics during stimulation with a novel, learned, then (the same) novel stimulus. Parameters: *K* = 40, *P* = 10,000, *N*_s_ = 1,400, *N*_m_ = *N*_d_ = 400,000, *f* = 0.05, Δ_x_ = 0.10, *o* = 1, |*η*_+_| + |*η*_*−*_| = 1. Panels (C–H) fix *σ*_0_ = 0.0.

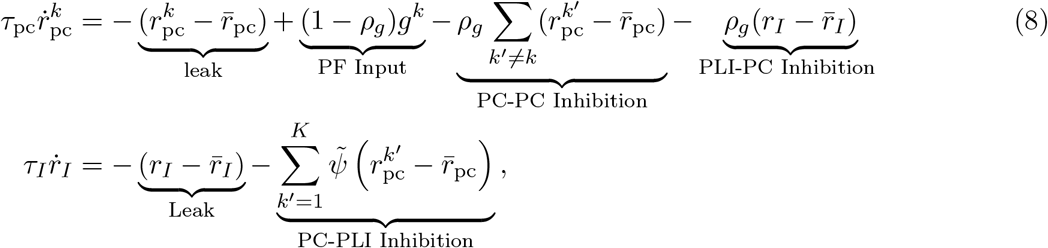

Lugaro cells of the PC layer and molecular layer interneurons are candidate substrates for the PLI in our model, as they are known to provide inhibitory feedback to PCs [69, 43–45]. In the second step, the fixed-point firing rates drive the DCN output through a nonlinear readout:

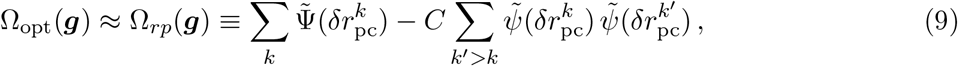

where 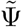 is a nonlinear activation function and we have defined the firing rate deviations 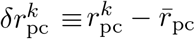 (full expressions in Methods §3). The first term captures the independent influence PCs exert on their DCN targets. The second reflects DCN sensitivity to pairwise correlations in PC activity, possibly acting as a static analogue of DCN sensitivity to synchronous firing in pairs of PC inputs [33, 70]. However, we show next that near-identical classification performance can be achieved without this term.

We examine three decoding strategies of increasing complexity: a separable decoder 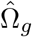 with a fitted elementwise nonlinearity 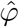 (Fig. 4A, top), an intermediate decoder 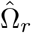 with a fitted separable nonlinearity 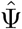 applied *after* running the recurrent dynamics to convergence (Fig. 4A, middle), and the full saddle-point approximation Ω_*rp*_ including both recurrent dynamics and pairwise interactions between PC outputs (Fig. 4A, bottom; eq. 9). The heatmaps in fig. 4B show SNR for each of these decoders in sweeps over *β* and Φ_*Y*_ with |*η*_+_| + |*η*_−_| = 1 held fixed. For the separable decoder, SNR is highest near balanced plasticity Φ_*Y*_ *≈* 0 where PC-PC correlations *ρ*_*g*_ vanish. For the recurrent decoders, SNR shows two distinct near-optimal regions in the upper-left and lower-right quadrants, possibly corresponding to “upbound” and “downbound” plasticity regimes [48]. In addition to increasing the maximum SNR, recurrent interactions permit near-optimal performance over a wider range of plasticity parameters. Ω_*rp*_ performs almost identically to 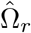, suggesting that pairwise interaction sensitivity is not critical for interpreting PC firing rates when recurrent interactions are present.

We next examine decoder behavior at a handful of selected operating points in the (*β*, Φ_*Y*_)-plane (fig. 4C,D). Across upbound and downbound regimes, recurrent dynamics center PC activity around a set point, leaving stimulus identity encoded in the distribution of firing rates rather than the mean (fig. 4E,F,G). Upbound learning is associated with decreases in PLI activity, and downbound learning with increases (fig. 4E, top right). While individual PC responses to novel and learned stimuli overlap heavily for novel and learned inputs, the decoders reliably distinguish these conditions by sensing nonlinear statistics of PC population activity (fig. 4E, bottom row). The forms of the fitted/derived nonlinearities 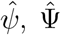, and 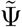 are shown in fig. 4F,G, overlaid with the marginal distributions of the learned PC inputs (F) or fixed-point firing rates (G). At operating points ii-vi, which mix LTP and LTD, fitted nonlinearities are often non-monotonic (fig. 4F,G), possibly consistent with depolarization-induced PC pausing [71] and post-inhibitory rebound spiking in the DCN [68], as previously discussed. Finally, we examine the dynamics of PC firing rates under a stimulation sequence which mimics the transient presentation of a saccade target [6, 7] or a conditioned stimulus [3], resulting in PC dynamics which resemble the transient bursts and pauses observed in experiments (fig. 4H).

### 2.6 Heterogeneous Plasticity Links Pairwise Correlations in PC Anatomy and Physiology

Many studies have measured either electrophysiology or anatomy of multiple PCs simultaneously, revealing weak but statistically significant correlations in both PC firing statistics and their patterns of synaptic input. Simultaneous electrode recordings and calcium imaging show that certain PC pairs fire both simple spikes and complex spikes synchronously more often than expected by chance [72, 34, 73, 8], with the degree of simple spike synchrony varying continuously across pairs and correlating with the similarity of their complex spike trains [72]. To connect with experiment, we express the plasticity signals in terms of a binary LTD-induction matrix ***M*** ∈ {0, 1} ^*P ×K*^, where 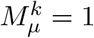 indicates CS-induced LTD for pattern cluster *µ* in PC *k*. We define 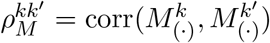 as the Pearson correlation between columns *k* and *k*^*′*^ of ***M***. While 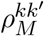 is not identical to the correlation of experimentally recorded complex spike trains, since only a subset of complex spikes effectively induce LTD [7, 42], it provides a natural theoretical analogue for fluctuations in complex spike similarity.

#### Simple spike synchrony

To compare with synchrony measurements (fig. 5A), we model PC simple spikes as inhomogeneous Poisson processes with rate 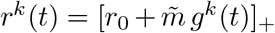, shown in fig. 5. We divide a trial into time windows of 50–100 ms duration, assigning each window PC inputs drawn from the learned cluster response distribution ***g*** ~ *P* (***g*** | *e*=1) with probability *p*_*l*_, or from the novel response distribution ***g*** ~ *P* (***g*** | *e*=0) with probability 1 − *p*_*l*_. Following experimental practice, we quantify synchrony by the fractional excess probability *S*^*kk′*^ of coincident spikes above baseline (see fig. 5A for demonstration). Under these conditions, we find (Methods) the following approximation:

**Figure 5:**
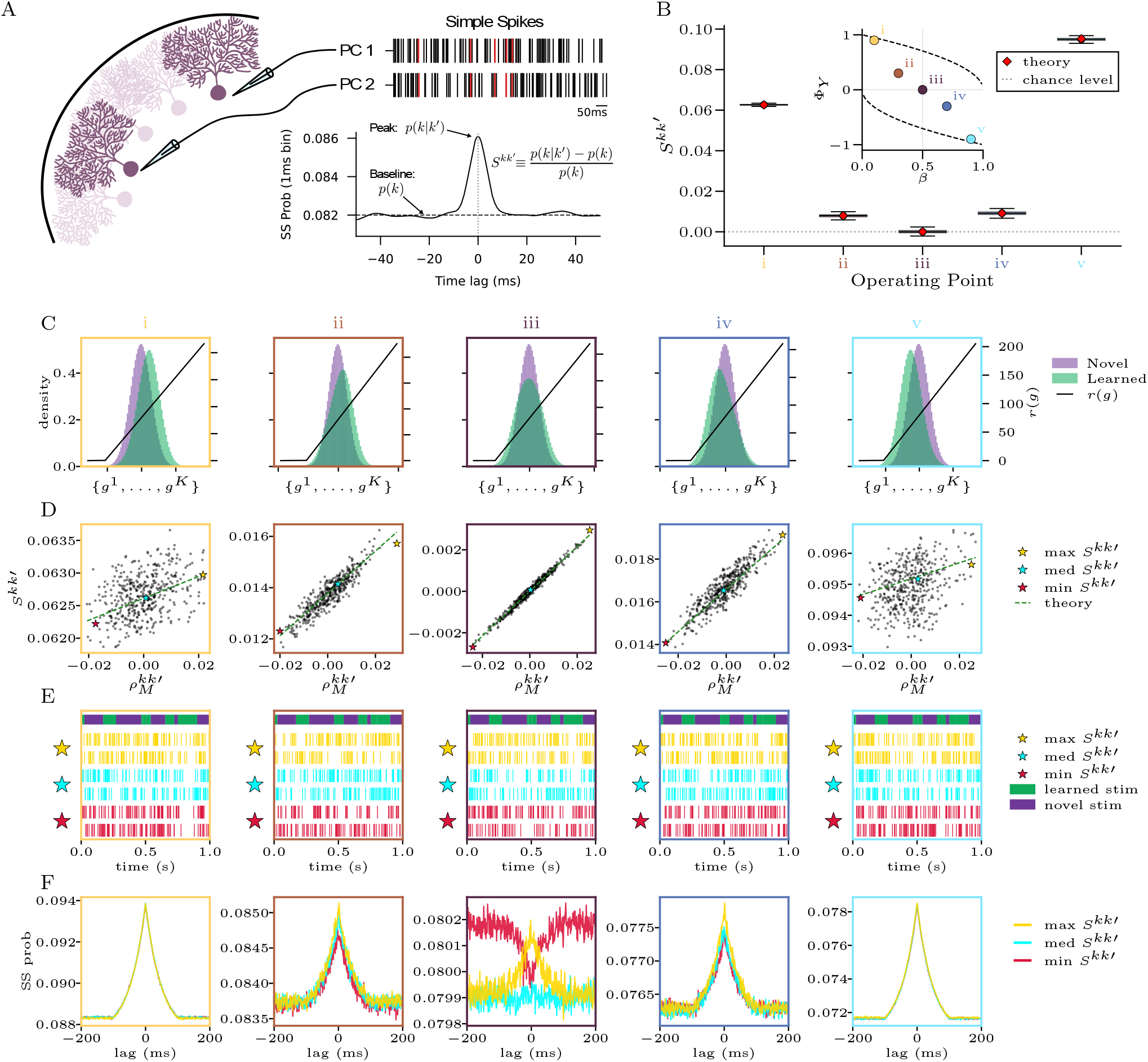
Heterogeneous plasticity produces varying degrees of synchrony across PC pairs. **(A)** Schematic of simultaneous PC recording and synchrony measurement. **(B)** Box plots of synchrony score *S*^kk^*′* across PC pairs at the operating points shown in the inset (*β*, φ_Y_) plane (conditions i–v). **(C)** PC response histograms at each operating point, with the simulated ReLU firing rate function *r*(*g*) overlaid. **(D)** Scatter plots of synchrony score *S*^kk^*′* vs. effective complex spike correlation 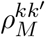 for all PC pairs. Point clouds cluster around the theoretical prediction (dashed lines; eq. 10). Three representative pairs (stars) are selected near the minimum, midpoint, and maximum of the theory line. **(E)** Spike rasters for representative PC pairs during novel and learned stimulus presentation (blue bars). **(F)** Cross-correlograms for the representative pairs. Near balanced plasticity, the zero-lag peak shows greater variability across pairs. Parameters: *K* = 30, *P* = 15,000, *N*_s_ = 1,400, *N*_m_ = *N*_d_ = 400,000, *f* = 0.05, Δ_x_ = 0.10, *σ*_0_ = 0.

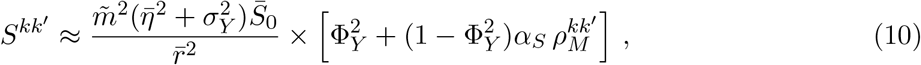

where 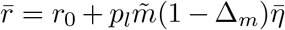 is the mean PC firing rate and 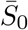 and *α*_*S*_ are constants independent of the plasticity parameters (full expressions in Methods). The scatter plots in fig. 5D confirm that synchrony scores for simulated PC pairs cluster tightly around this prediction. Representative spike rasters for pairs with low, medium, and high 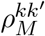 are shown in fig. 5E, and their cross-correlograms in fig. 5F. For balanced and near-balanced plasticity, the relative height of the zero-lag peak exhibits a larger spread across pairs, and fluctuations in 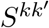 correlate with fluctuations in 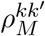, consistent with experimentally observed correlations between SS and CS synchrony scores [72]. Anti-correlated PC pairs are present after training at balanced plasticity, and become less common as plasticity moves into the upbound or downbound regimes.

#### Correlated PF–PC synaptic weights

We next consider how heterogeneous plasticity shapes synaptic weights from PFs to PCs. To measure similarity in the learned weights, we introduce the pairwise weight correlation metric 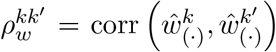, where the correlation is measured over the set of PFs which form synapses with PCs *k* and *k*^*′*^ (fig. 6A, Methods). Under heterogeneous plasticity, we find the following simple approximation (Methods §3):

**Figure 6:**
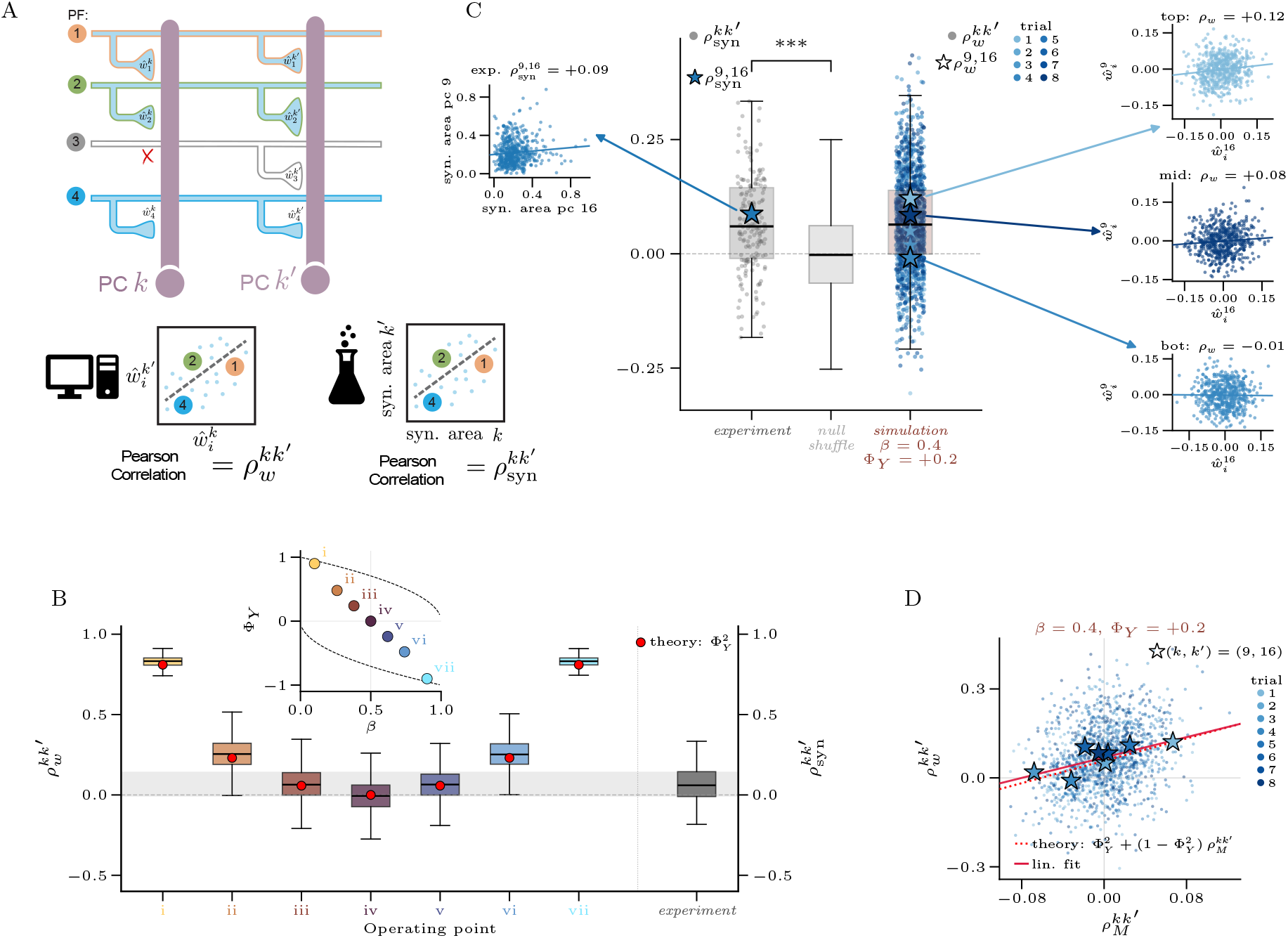
Heterogeneous plasticity predicts weakly correlated PF–PC synaptic weights. **(A)** The weight similarity metric 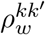 is defined as the Pearson correlation in PF–PC weights, restricted to the set of parallel fibers which connect to PCs *k* and *k*^*′*^. As an experimental analogue which can be readily calculated in a publicly available dataset, we define 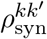 as the Pearson correlation in the total synaptic area of connectivity from PFs to PCs *k* and *k*^*′*^. **(B)** Box plots of 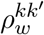 across PC pairs at several plasticity parameter operating points (inset). Red points show theoretical expected overlap 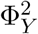. Gray box plot shows the experimental distribution of synapse area vector correlations 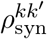, and shaded region shows the inter-quartile range of the experimental distribution. **(C)** Detailed visualizations of box plots for experimental distribution of 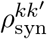 and simulated weight correlations 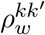 at the best-fitting simulated operating point iii (*β* = 0.4, φ_Y_ = 0.2). Superimposed gray dots show values of individual 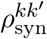, and blue dots show individual 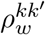 separated by shade across 8 trials with independent draws of the plasticity signals 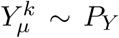. Stars show correlations for PC pair (*k, k*^*′*^) = 9, 16, which had the highest number of *Ñ*^9,16^ = 553 shared PF inputs in the experimental dataset. We also compare to the experimental distribution of 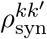 after randomly shuffling synaptic weights among mutually connected PFs independently within each PC. We observe a statistically significant increase in median 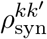 across pairs relative to the null shuffle (*p <* 5 × 10^*−*4^), one-sided Monte Carlo permutation test (*N* = 2000 independent shuffles), see methods and fig S4. **(D)** Scatter plot showing simulated 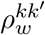 values correlate with pairwise plasticity correlations 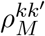. Points are colored by trial, with stars for pair (*k, k*^*′*^) = (9, 16). Solid red line shows linear fit, and dotted red line shows theoretical prediction.

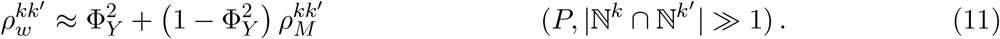

As with synchrony, we expect a baseline similarity depending on the plasticity ratio Φ_*Y*_, with fluctuations in 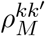 driving correlated fluctuations in 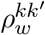. We may also measure an analogous quantity using a publicly available connectome reconstruction of PF–PC connectivity in the mouse cerebellar cortex [46]. Specifically, as an experimental analogue of the weight similarity 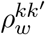, we define 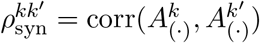, where 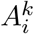 is the total area of synaptic connections from PF *i* into PC *k* (fig. 6A, Methods).

In fig. 6B, we simulate a model network with connectivity matched to the available connectome reconstruction [46], and weights learned using the heterogeneous plasticity learning rule. We find that correlations 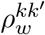 are centered around 0 at balanced plasticity, and increase as plasticity moves into the upbound or downbound regimes, in quantitative agreement with the expected analytical baseline 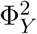 (eq. 11). In fig. 4, we saw that network performance is optimized in regimes with a slight upbound or downbound learning bias. As a result, we expect a small but nonzero 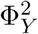, resulting in a slight but significant correlation in learned weights. In line with this prediction, we find that the experimental distribution of 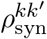 has median and inter-quartile range consistent with a model network trained under heterogeneous plasticity with |Φ_*Y*_| ≈ 0.2 (fig. 6B), with a statistically significant shift upward relative to a null shuffle in the input weights to each PC (figs. 6C, S4C). Finally, in fig. 6D, we show 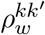 against 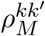 from simulations at the same operating point, confirming that fluctuations in 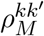 drive correlated fluctuations in 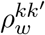, with slope in good agreement with eq. 11.

### 2.7 Wiring of the Olivo-Cortico-Nuclear Loop

Because climbing fibers initiate the complex spikes that instruct synaptic plasticity, PCs innervated by the same inferior olive (IO) neuron will receive synchronous complex spikes and presumably undergo similar plasticity schedules. To investigate the effect of shared inferior olive inputs, we partition the *K* PCs of a functional zone into blocks of size *K*_*b*_, where all PCs within a block share the same plasticity signal 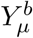 drawn from *P*_*Y*_ (Methods §3). Setting *K*_*b*_ = 1 recovers full heterogeneous plasticity (eq. 6). Following training with this plasticity sharing scheme, the optimal decoding strategy is to first average activity within blocks, 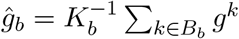, then apply Ω_opt_ to these *N*_*b*_ = *K/K*_*b*_ block averages with appropriately rescaled noise parameters (Methods §3). The ensemble therefore behaves as though it contains only *N*_*b*_ independent learners.

Sweeping over plasticity parameters (*β*, Φ_*Y*_) and intrinsic noise *σ*_0_ for each block size reveals two clear trends (fig. 7B–D). First, increasing *K*_*b*_ has a detrimental effect on optimal SNR, except at *σ*_0_ values so large that ensemble performance is poor at all *K*_*b*_ (fig. 7C). Second, as *K*_*b*_ grows, the optimal operating point migrates toward |Φ_*Y*_| = 1 (fig. 7D), recovering homogeneous plasticity when *K*_*b*_ = *K*. Maximizing the computational benefit of heterogeneous plasticity therefore favors a wiring scheme in which *K*_*b*_ = 1 so that each PC in a functional zone is innervated by a unique IO neuron. This prediction diverges from the “olivocerebellar modules” proposed by Shadmehr [9], in which *K*_*b*_ ≈ 6–8 (fig. 7A). This discrepancy may be attributed to our focus on PCs as the dominant locus of plasticity, while Shadmehr argues, rather, that climbing fiber modularity promotes synchronous complex spikes, necessary to induce plasticity in MF-DCN synapses [73]. A multi-lobule extension of our model may resolve this discrepancy (see discussion).

**Figure 7:**
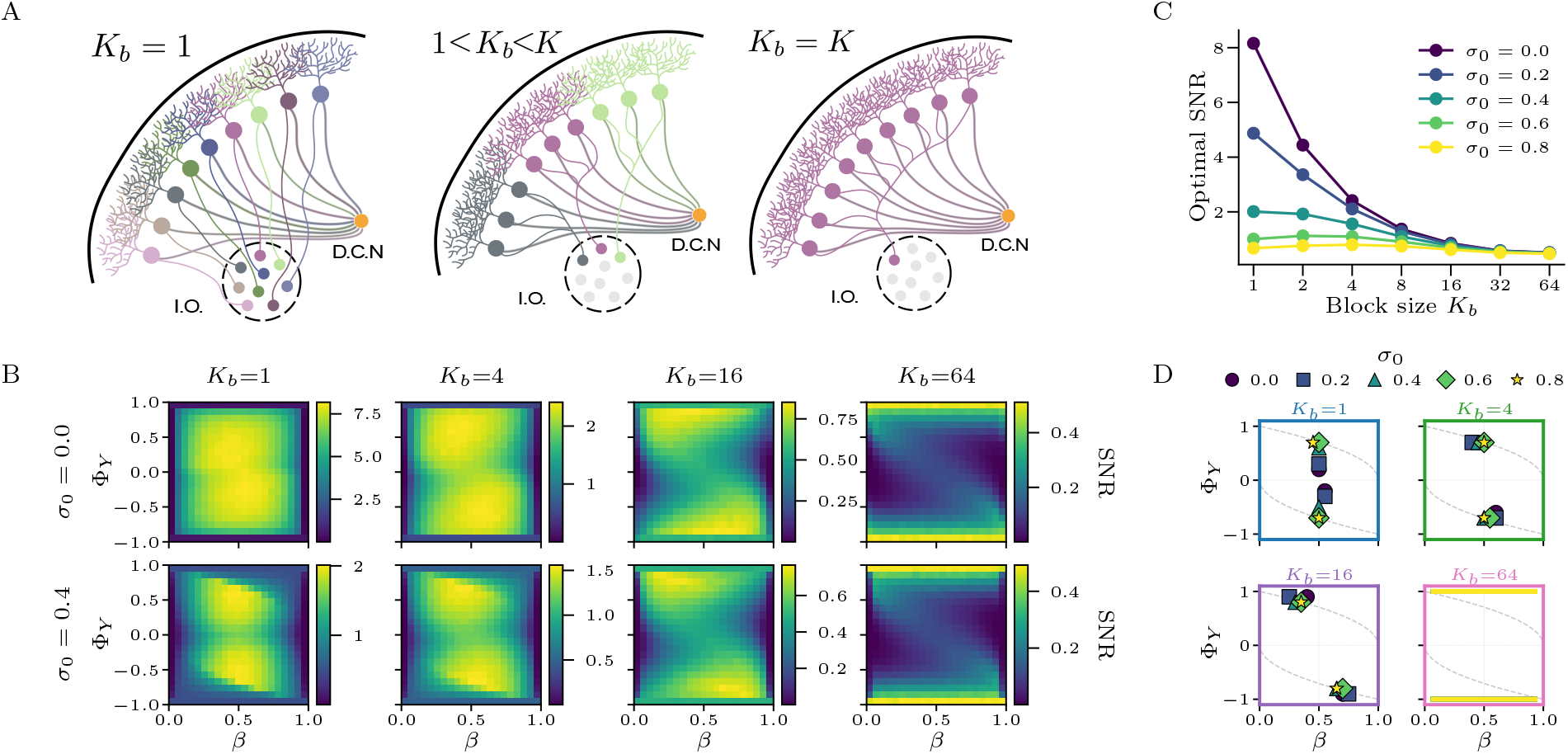
Maximally heterogeneous IO-to-PC wiring optimizes ensemble performance. **(A)** Schematic wiring diagrams of the olivo-cortico-nuclear loop at three block sizes *K*_b_. PCs and their connections are colored by the IO neuron from which they receive climbing fiber input. Left: *K*_b_ = 1 (each PC receives a unique IO input, equivalent to heterogeneous plasticity). Center: 1 *< K*_b_ *< K* (PCs form clusters of size *K*_b_ with shared plasticity). Right: *K*_b_ = *K* (all PCs share one IO input). **(B)** SNR heatmaps in the (*β*, φ_Y_) plane for block sizes *K*_b_ ∈ {1, 4, 16, 64} (columns) and intrinsic noise *σ*_0_ ∈ {0, 0.4} (rows), using the moment-matched decoder applied to block-averaged PC inputs. **(C)** Optimal SNR (maximized over *β* and φ_Y_) as a function of block size *K*_b_ for varying *σ*_0_. Performance decreases monotonically with *K*_b_, except under extreme noise where ensemble performance is always poor. **(D)** Location of the optimal operating point 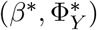 for each *K*_b_ and *σ*_0_. As block size grows, the optimum migrates toward |φ_Y_| = 1, recovering homogeneous plasticity. Parameters: *K* = 64, *P* = 10,000, *N*_s_ = 1,400, *N*_m_ = 400,000, *N*_d_ = 200,000, *f* = 0.05, Δ_x_ = 0.10.

## 3 Discussion

We presented a theory of associative learning in PC populations sharing mossy fiber inputs, DCN targets, and complex spike tuning. Within an ensemble, we assume learning occurs during well-timed population-level plasticity events in which the direction and/or magnitude of plasticity varies randomly between PCs. Our analysis pinpoints two essential ingredients for ensemble learning: *heterogeneous plasticity* and *nonlinear processing of learned PC input currents*. Including either ingredient in isolation offers no performance improvement relative to decoding PF activity with a single large neuron. However, when heterogeneous plasticity is combined with a suitable nonlinear decoding scheme, ensemble memorization capacity can improve well beyond that of a direct read-out. In addition to providing a computational benefit, heterogeneous plasticity learning unifies a variety of experimental observations, including recurrent interactions between PCs, a preference for upbound or downbound plasticity, and weak but significant pairwise correlations in PC anatomy and physiology.

### Beyond the Marr–Albus Theory: multilayer processing in Cerebellar Cortex

Classical theories of cerebellar learning focus on the expansion from the MF to GrC layer as a mechanism for increasing the pattern separation capabilities of an individual PC [14, 15]. Subsequent extensions of the Marr–Albus framework have incorporated noisy sensory information [17], temporal dynamics of GrC activity [74, 75], sparse [18, 76] or structured [77, 78, 46] MF–GrC connectivity, sign constraints in learned PF–PC weights [79, 80], and gradations in task complexity [81, 50], but treat an individual PC as the locus of the learned function. However, when the DCN is considered the true output of the cerebellum, PCs should instead be understood as an additional intermediate layer in the pathway through the cerebellar cortex [9].

Because CSs drive plasticity primarily at the PF–PC synapses, the circuit forms a natural analogue to ensemble machine learning, where multiple predictors are trained independently and their outputs combined. To benefit from ensembling, ensemble members must learn diverse functions [54]. Here we have proposed that the experimentally observed low-probability physiology of CSs following individual sensory events [20] instructs diverse learned mappings across PCs which can be combined to achieve more accurate recall than any single-PC model can attain. Two concurrent works similarly argue for multilayer processing in the cerebellar cortex, but propose alternative mechanisms to diversify the mappings learned by PCs. In one proposal, Hiratani [82] suggests that shunting inhibition by molecular-layer interneurons gates plasticity at PF–PC synapses. In another proposal, Holtrup et al. [83] show that sparse PF–PC connectivity can improve ensemble performance with an alternate learning rule which includes a postsynaptic contribution to PF–PC plasticity. Another related work [84] studies memorization capacity of ensembles of Hebbian-trained classifiers with sparse connectivity to inputs. Our proposal is, to our knowledge, the first to suggest a productive role of CS stochasticity in an ensemble learning computation.

### A computational purpose for PC variability

Achieving reliable behavioral responses despite the variability of individual neurons is a fundamental challenge for the nervous system. A common view is that individual neurons represent noisy estimates of a desired signal, and reliable motor commands are recovered by pooling over many such noisy neurons. Consistent with this view, trial-to-trial fluctuations in firing rates of individual floccular PCs correlate with fluctuations in eye velocity [85], and saccade velocity may be recovered by averaging firing rates of many PCs in the oculomotor vermis [6]. Under this view, the variation between PCs has no computational utility, and is merely averaged away upon convergence in the DCN, as suggested by Walter and Khodakhah [53]. In contrast, we have shown here that heterogeneous plasticity instructs a nonlinear population code [47] in which stimulus identity is represented in the *distribution* of PC input currents *g*^*k*^. Under this view, the degree of PC-PC variance is a critical quantity that conveys stimulus identity to the DCN target. One possible reconciliation of these views is that linear and nonlinear statistics of PC activity convey complementary information to the DCN. Population mean is well-suited to represent graded quantities like movement velocity, while nonlinear population statistics such as transient pausing [59, 86] or synchronization of simple spikes [33, 34] may be better suited to represent binary information, such as whether and when to engage a corrective response or initiate a movement. Determining how heterogeneous plasticity contributes to such mixed coding regimes during continuous motor control remains an important question for future work.

### Reconciling burst-pause dynamics with upbound/downbound learning

The finding that bursting and pausing PC responses coexist within a functional zone [6, 7] is seemingly at odds with the emerging consensus that learning proceeds in individual microzones through either upbound learning driven by LTP or downbound learning driven by LTD at the PF–PC synapses [48]. Our results suggest two possible resolutions to this conflict. First, what experiments describe as “upbound” and “downbound” learning mechanisms may actually consist of a mix of LTP and LTD, with a bias toward one or the other. Indeed, most of the near-optimal operating points fall in this range (fig. 4B). Alternatively, we see that even in the pure-LTP or pure-LTD regimes (operating points i and vii), heterogeneity in the strength of plasticity across PCs is sufficient to create a mix of bursting and pausing responses after centering by the derived recurrent dynamics (fig. 4H). Further experiments may be required to disambiguate these two possibilities.

### A shared mechanistic basis for anatomical and physiological PC–PC correlations

Simultaneous electrode recordings [72, 7] find weak but statistically significant correlations in PC firing rates. Here, we have also demonstrated weak but significant correlations in anatomically defined synaptic weights in pairs of PCs (fig. 6C). We have argued that both of these forms of PC–PC similarity are controlled by the statistics of the complex-spike plasticity signals received by each participating PC during training. Specifically, equations 10 and 11 relate both these forms of PC similarity to the plasticity ratio Φ_*Y*_, with fluctuations driven by spontaneous correlations 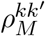 in their plasticity histories. In the cerebellar circuit, shared plasticity arises from similarities in climbing-fiber-driven complex spiking, which is mediated by gap junction coupling between IO neurons [87]. We therefore predict that disrupting gap junction formation in the IO will reduce both simple-spike synchrony and correlations 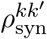 in PF–PC synaptic weights.

### Limitations and extensions

As a first pass at understanding the effects of heterogeneous complex spiking across PCs within a microzone, we modeled learning as a Bernoulli process in which each PC receives a complex spike with the same probability. The resulting heterogeneous plasticity substantially improves ensemble learning performance, and explains a variety of experimental observations. However, CSs exhibit some non-random spatial structure within microzones [88], and several extensions of our model may provide a more complete account of cerebellar learning: First, DCN feedback to the IO allows PCs to influence CS probability in their efferent climbing fibers [89], possibly aiding long-time stability in heterogeneous plasticity learning dynamics. Second, CS probabilities in PCs of the oculomotor vermis vary continuously with the angle of saccade errors relative to each PCs “CS-on” direction [6, 7], suggesting a mechanism by which heterogeneous plasticity learning may generalize to continuous control tasks. Finally, a generalization of our model to the multi-lobule setting is necessary to gain a more complete understanding of the full cerebellar microzone. Such an extension may reveal a productive role for the observed 1:7 divergence from IO neurons to PCs [24] by enabling synchronous complex spikes in PCs which receive distinct sensory information, but converge on common targets in the DCN [90].

### Heterogeneous plasticity beyond the cerebellum

Associative learning occurs in many circuits across the brain. For example, the fly mushroom body shares a similar anatomical structure with the cerebellar cortex, and is responsible for learning appetitive and aversive associations with olfactory stimuli. There, dopamine signaling instructs long-term depression from Kenyon Cells (KCs) to Mushroom Body Output Neurons (MBONs) in a manner analogous to the action of climbing fibers on the PF-PC synapse. A recent experimental study found that LTD is heterogeneously expressed across different KC-MBON synapses [91]. Unlike the heterogeneity *between* PC plasticity directions which we have studied here, Davidson et al. [91] find heterogeneous expression of LTD across the synapses onto a single Mushroom Body Output Neuron. Understanding the benefits of this form of heterogeneous plasticity will require additional theoretical efforts. Another recent model of content-addressable memory formation in the Hippocampus posits that bi-directional plasticity at synapses from area CA3 to CA1 is gated by random, sparse inputs from the Entorhinal Cortex [92]. Random, heterogeneous plasticity may therefore be a widely implemented strategy for high-capacity memorization across many neural circuits.

Heterogeneous plasticity is also closely related to the *bagging* algorithm from ensemble machine learning [93] – famously applied to decision trees in the random forests algorithm [94] – in which an ensemble of predictors is trained on random subsets of data and their predictions combined to yield a stronger overall prediction. In the sparse LTD or sparse LTP limits (*η*_+_ = 0 or *η*_−_ = 0), each training stimulus is assigned to a random subset of PCs and ignored by the rest, directly paralleling the bagging procedure. Because heterogeneous plasticity is a fully local and online learning rule, it may also find applications in on-device learning for neuromorphic hardware, where backpropagation-based training is impractical but local plasticity rules can be implemented efficiently.

## Methods

We will denote the standard Gaussian measure 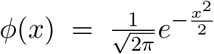, the Gaussian CDF 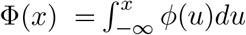 and Gaussian tail function *ℋ* (*x*) ≡ 1 − Φ(*x*). We denote by Θ[·] the Heaviside step function. Bold lower-case letters represent vectors and bold upper-case letters represent matrices. The upper index (·)^*k*^ always iterates over the PCs 1, …, *K* within an ensemble. The lower indices *µ, ν* are used to represent stimulus cluster identity in the range 1, …, *P*. We also use the Kronecker delta function *δ*_*kk*_*′* = 1 when *k* = *k*^*′*^ and 0 otherwise.

### Task and Performance Metrics

Associative learning task is as described in sec. 2.1. Novel inputs are drawn uniformly from 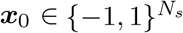. Under the learned-input condition, an input ***x*** is sampled by first selecting a random input cluster *µ* ∼ Unif(1, …, *P*), then randomly flipping each element of the cluster center 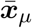 with probability Δ_*x*_*/*2. Performance is quantified using the AUC, or the monotonically related quantity

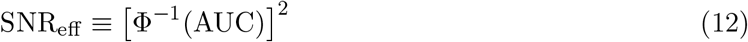

as described in sec.2.1. The AUC is equivalently expressed as a ranking probability [95]:

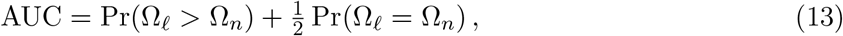

where Ω_*n*_ and Ω_*ℓ*_ are network outputs for a randomly drawn input from the novel and learned input distributions, respectively. In the special case where the decoder outputs are approximately Gaussian under each class, so that 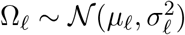 and 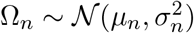, we have

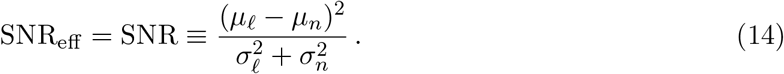

We refer to the SNR_eff_ simply as the SNR in the main text and for the remainder of the methods. We define the *capacity P*_max_(*ϵ*) as the largest number of pattern clusters for which classification performance remains above a threshold:

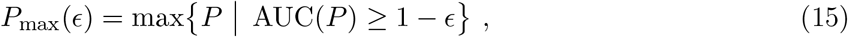

where *ϵ >* 0 is a prescribed error tolerance. In terms of the effective SNR, this is equivalent to the condition 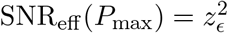 with *z*_*ϵ*_ = Φ^−1^(1 *ϵ*). We expect that at large *K* values the capacity becomes insensitive to the exact choice of *ϵ*.

### Model Architecture and Learning Rule

The model architecture is described in sec. 2.1. In the transformation from input to GrC activity ***m***(***x***) = Θ[***Jx*** − *T*], we sample 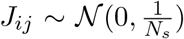. To achieve an average GrC sparsity of *f*, we set *T* to solve *f* = ℋ (*T*). Two statistics of this expansion govern downstream performance [17]. First, the *mixed-layer cluster size*:

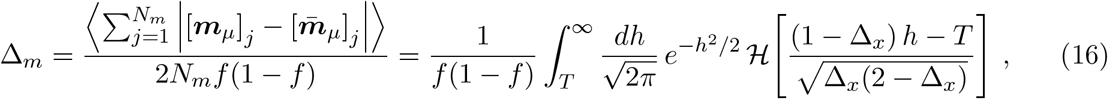

which measures the size of the input clusters after projection to the mixed-layer representation [17]. Second, we have the effective dimensionality of the PF representation:

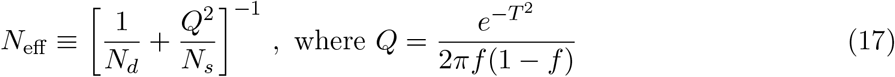

is the sparsity-dependent *excess overlap* statistic [17]. Each PC samples from a subset ℕ^*k*^ ⊆ 1, …, *N*_*m*_ with size *N* ^*k*^ = |ℕ^*k*^|. To reduce parameters, we adopt a *homogeneous overlap assumption*, so that

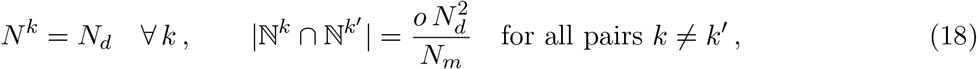

where the parameter *o* controls the degree of overlap in PF sampling. The value *o* = 1 corresponds to random (independent) sampling. In the SI, we show using standard combinatorial arguments that *o* is constrained to lie within the range:

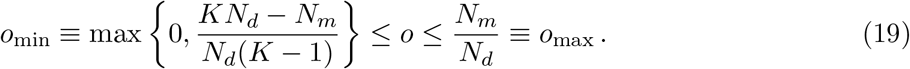

In the main text, we consider a one-shot learning rule in which each cluster center is presented to the network once during training, and PCs undergo weight updates according to eq. 2. After training, the learned weights can be written

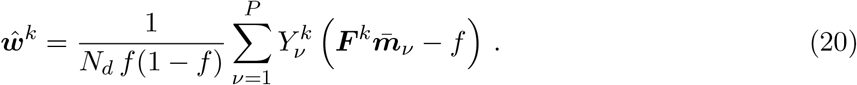

We organize the plasticity signals into a matrix ***Y*** ∈ *ℝ*^*P ×K*^ with entries 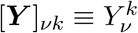.

### Learned PC Response Statistics

With learned weights given by eq. 20, the input currents ***g***_*µ*_ ∈ *ℝ*^*K*^ to the PC layer in response to a test pattern ***x***_*µ*_ drawn from cluster *µ* are Gaussian conditioned on the plasticity signals:

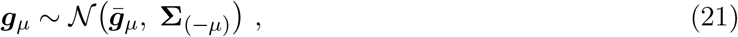

where 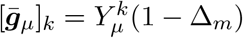 and

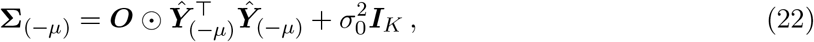

with ***Ŷ*** ∈ R^(*P* −1)*×K*^ denoting the plasticity matrix ***Y*** with row *µ* removed, and ⊙ denoting the Hadamard (elementwise) product. For inputs from the novel pattern distribution (by convention, *µ* = 0), we have 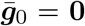 and ***Ŷ***_(−0)_ = ***Y***. Under the *homogeneous overlap* assumption, the overlap matrix ***O*** ∈ ℝ^*K×K*^ takes the form:

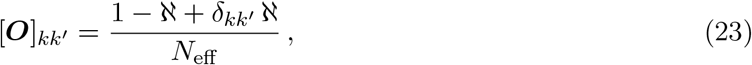

where we define the *effective overlap fraction*:

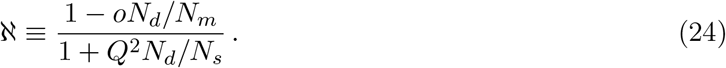

#### Homogeneous Plasticity

Under homogeneous plasticity, we have 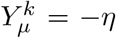 for all *µ* and *k*. The covariance becomes 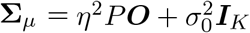 (assuming *P* ≫ 1 so that *P* − 1 ≈ *P*). PC responses can then be decomposed as:

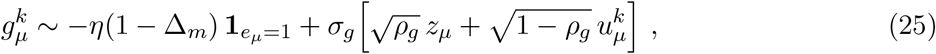

where *z*_*µ*_ ~ *N*(0, 1) is a fluctuation shared across all PCs and 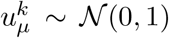 are independently sampled for each PC. The variance and correlation coefficient of cross-talk fluctuations are then given by :

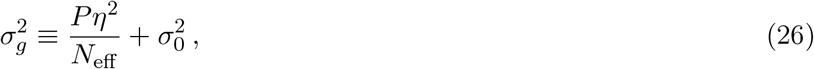

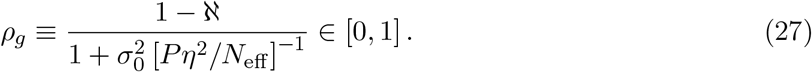

### Monte Carlo Simulations

In Monte Carlo simulations, we determine network SNR through a quasi-analytical procedure. In a single trial, we sample a random ***Y*** matrix according to eq. 6. Then, we generate *N*_*samp*_ Monte Carlo samples of the PC inputs ***g*** from the novel stimulus distribution and the learned stimulus distribution using eq. 21. For each learned stimulus, we sample *µ* ~ Unif(1, …, *P*) then apply eq. 21. We then compute the network outputs Ω(***g***), determine AUC using the empirical ranking definition (eq. 13). AUC may then be converted to SNR_eff_ using eq. 12. To obtain capacity estimates at a particular *K* value, we solve AUC(*P*) = 1 − *ϵ* using a binary search over *P* values, with a single pre-sampled ***Y*** matrix per trial.

### Analytical SNR and Capacity Curves

Here, we state the analytical formulas obtained for the SNR and capacity values under different plasticity conditions and decoders. Full derivations are provided in the SI.

#### SNR under Homogeneous Plasticity

After training under homogeneous plasticity, the statistics of the decoders are described by eq. 25. In this case, the optimal decoder is a simple linear average 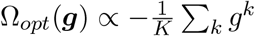, and an analytical form for the SNR follows easily as:

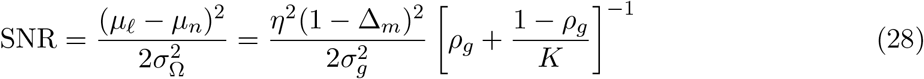

We use this formula to plot SNR under homogeneous plasticity in fig. 2E. It follows from this and eq’s. 27, 26 that SNR is monotonically increasing with PC size *N*_*d*_, reaching a maximum for fully connected networks, when *N*_*d*_ = *N*_*m*_. We may also consider the hypothetical “direct readout” architecture in which the circuit’s output Ω(***g***) is a direct linear readout from all PFs by setting *K* = 1, *N*_*d*_ = *N*_*m*_, and *σ*_0_ = 0. The SNR reduces to:

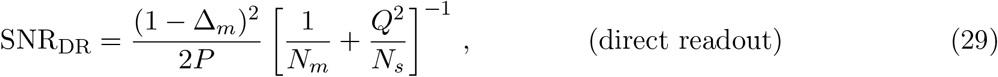

recovering the SNR formula from [17].

### SNR and Capacity under Heterogeneous Plasticity

We show numerically in the SI using Monte Carlo simulations that SNR increases monotonically with *N*_*d*_ when all other parameters are held fixed. We therefore restrict attention here to the fully connected case *N*_*d*_ = *N*_*m*_, which sets and *N*_eff_ =[1/*N*_*m*_ + Q^2^/*N*_*s*_]^−1^. In SI §S5, we derive analytical approximations to SNR for separable decoders of the form Ω(***g***) = ∑_*k*_ *Φ*(*g*^*k*^), with *Φ*(*g*) ∈ *{g*, (*g* − *κ*)^2^, Θ(*κ* − *g*)*}*. Capacity *P*_max_ is then obtained by solving 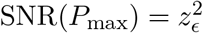.

#### Linear decoding

In the case of the linear decoder *φ*(*g*) = *g*, we obtain

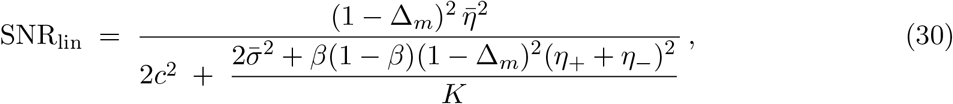

where 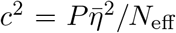 and 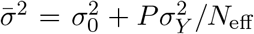. It is easily verified that eq. 30 is maximized over *β* ∈ [0, 1] at the boundary values *β* = 0 (when *η*_+_ *> η*_−_) or *β* = 1 (when *η*_+_ *< η*_−_), recovering homogeneous plasticity. Heterogeneous plasticity therefore offers no benefit when paired with a linear decoder.

#### Quadratic decoding

For the centered quadratic decoder *Φ*(*g*) = (*g* − *κ*)^2^ under balanced plasticity 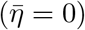, we obtain a closed-form SNR approximation (SI §S5.4):

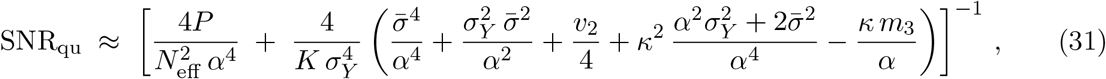

where 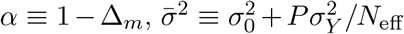 is the per-PC crosstalk variance, and 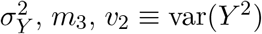 are the second, third, and squared-signal cumulants of the plasticity distribution *P*_*Y*_ (full expressions in SI). SNR_qu_ is optimized by setting 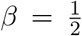 and *κ* = 0. At this optimum, setting 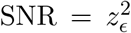 determines the capacity *P*_max_(*K, ϵ*) in closed form as the positive root of a quadratic in *P*_max_ (SI §S5.4), with the scaling regimes

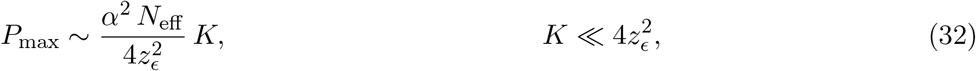

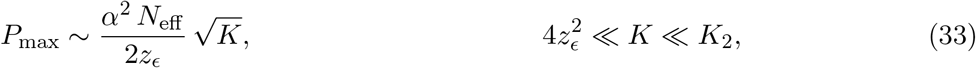

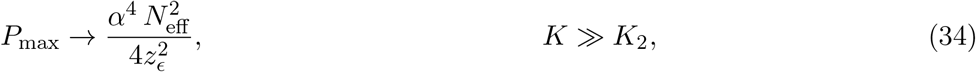

with plateau onset at 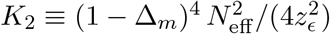.

#### Pause-based decoding

For the pause decoder *Φ*(*g*) = Θ[*κ* − *g*] under balanced plasticity 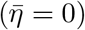, the SNR takes the general separable form (SI §S5.2)

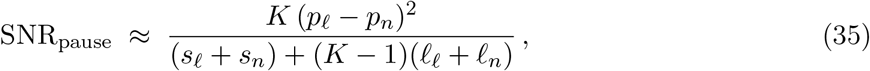

where *p*_*c*_, *s*_*c*_, and *ℓ*_*c*_ (for *c* ∈ {*n, ℓ*}) are the plasticity-averaged per-PC mean, per-PC variance, and PC–PC covariance of the pause indicator. To state these compactly, let *α* ≡ 1 − Δ_*m*_ and write the per-class crosstalk scales 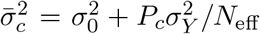 with *P*_*n*_ = *P*, *P*_*ℓ*_ = *P* − 1. The novel and learned inputs standardize the pause threshold as

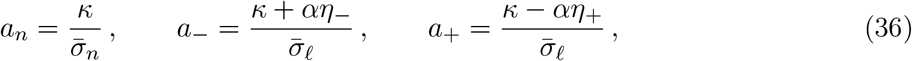

where *a*_∓_ are the LTD/LTP branch thresholds, occurring with probability *β* and 1 − *β*. We denote the corresponding branch average and branch (co)variance by

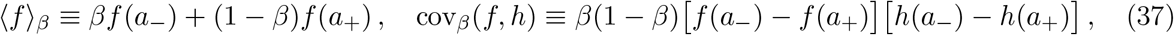

with var_*β*_(*f*) ≡ cov_*β*_(*f, f*). Fluctuations of the crosstalk covariance enter through the diagonal crosstalk 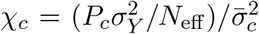 fraction and its strength 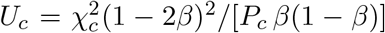, together with the Gaussian-moment functions

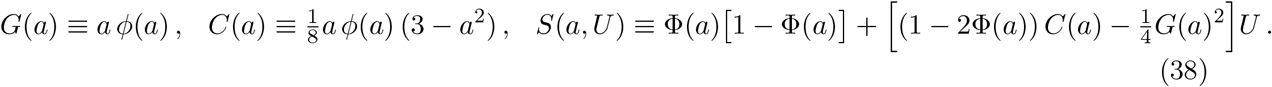

In terms of these, the novel-input moments are

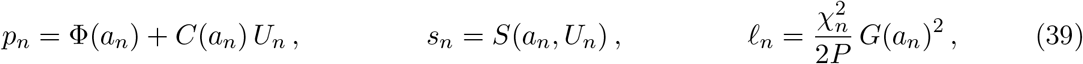

and the learned-input moments are

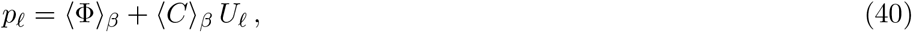

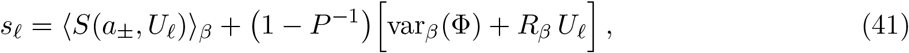

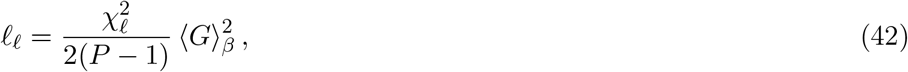

with

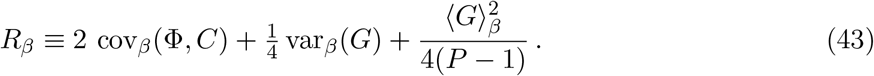

Substituting these into eq. 35 gives the pause-decoder SNR; the capacity *P*_max_ follows by solving 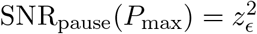. In SI §S5.5.4, we show that when *σ*_0_ = 0 and LTD probability is scaled down as *β* = *λ/K*, the above SNR formula is consistent with a capacity scaling *P*_max_ ∝ *KN*_eff_ */* log(*K*).

### Optimal and Approximately Optimal Decoding of learned PC Inputs

It follows from the Neyman-Pearson lemma [95] that the decoder Ω(***g***) which maximizes the AUC is the log-likelihood ratio of eq. 4. Any monotone transformation of Ω_opt_ preserves the ROC curve, and therefore the AUC and SNR_eff_. The form of Ω_opt_ depends on the plasticity model through the statistics of ***g*** under each class, ranging from a simple linear average (under homogeneous plasticity) to a nonlinear function involving recurrent dynamics (under heterogeneous plasticity). We provide an overview of the results here and defer to the SI for detailed derivations.

#### Optimal Decoder under Homogeneous Plasticity

Under homogeneous plasticity, both *P* (***g***|*e* = 0) and *P* (***g***|*e* = 1) are multivariate Gaussians with the same covariance matrix **Σ** but different means. The log-likelihood ratio reduces to:

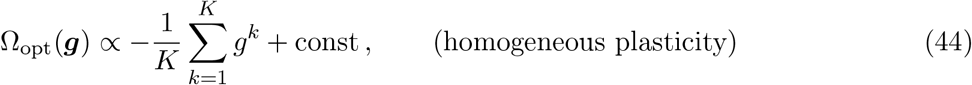

so that the optimal decoder is a simple linear average of the PC inputs.

#### Optimal Decoder under Heterogeneous Plasticity

Under heterogeneous plasticity, the statistics of ***g***_*µ*_ are described by a Gaussian distribution with a random covariance matrix. For the purposes of deriving an optimal decoder, we approximate this distribution by replacing the covariance matrix with its expected value, and considering only the fluctuations in the mean over realizations of ***Y***. Under this approximation, we have

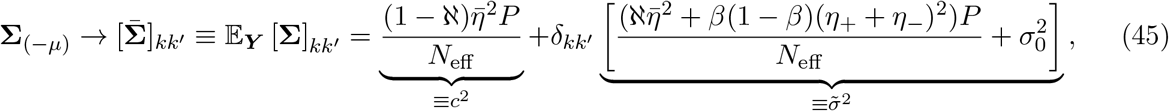

where we have defined 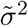 and *c*^2^ as the variance of the independent and shared fluctuations, respectively. The likelihood function for the novel class is then simply 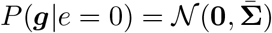. The likelihood function for the learned class *P* (***g*** | *e* = 1) reduces to a mixture of Gaussians 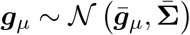 where the mean 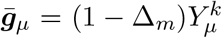 can take one of 2^*K*^ possible values, as each 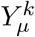 takes the value −*η*_−_ with probability *β* and *η*_+_ with probability (1 − *β*). Evaluating the likelihood directly would require a computation time which scales exponentially with *K*. Using standard manipulations (see SI), we may rewrite the likelihood as the following one-dimensional integral:

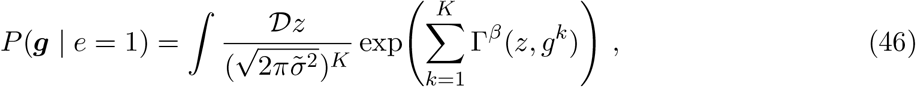

where 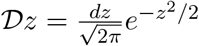, and

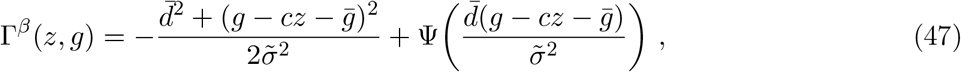

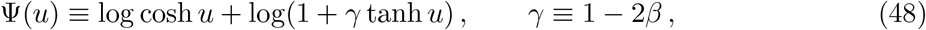

with 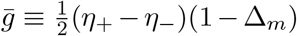 and 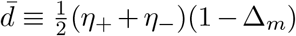, permitting linear time evaluation in *K*.

#### Saddle Approximation to the Optimal Decoder: Define the following nonlinearities

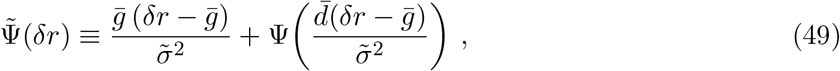

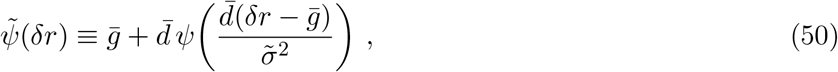

where the scalar nonlinearity Ψ is given in eq. 48, and where

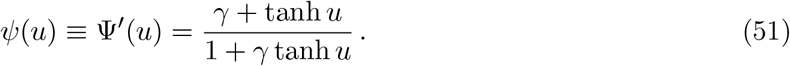

In the SI, we apply a saddle-point (Laplace) approximation to eq. 46, and show the resulting approximation can be equivalently evaluated in a two-step decoding process. The first step is to run the dynamical system of eq. 8 to convergence. Then, expressing the PC and PLI firing rates in terms of their deviations from their baseline firing rates, 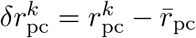 and 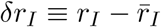, the saddle approximation yields eq. 8, and the final output given by 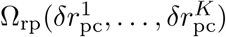 as in main text eq. 9.

#### Moment-Matched Approximation to the Optimal Decoder

As a simpler alternative to the saddle-point approximation, we may approximate Ω_opt_(***g***) by replacing *P* (***g*** | *e*=1) with a Gaussian distribution with matching first and second moments: mean *µ*_1_**1** with 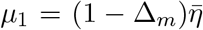, and covariance 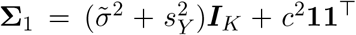, where 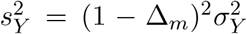 captures the additional variance introduced from fluctuations in the signal term due to randomness in the plasticity signals 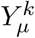. The resulting “moment-matched” approximate optimal decoder may then be reduced to the following form:

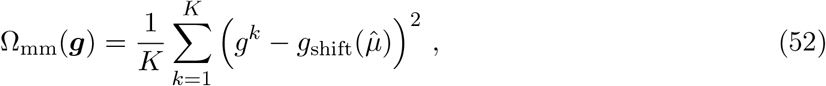

where 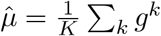 is the ensemble mean and

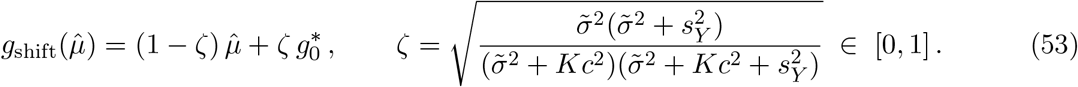

The shift *g*_shift_ interpolates between the ensemble mean 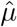 and a fixed reference 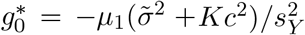, with *ζ* controlling the balance. Under balanced plasticity 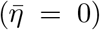, the global noise vanishes (*c* = 0), giving *ζ* = 1 and *g*_shift_ = 0, so that the decoder reduces to the pure secondmoment statistic Ω_mm_(***g***) = ∑_*k*_(*g*^*k*^)^2^. When 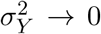 with 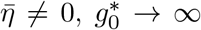, and we recover the linear decoder.

#### Fitting Optimal Separable Nonlinearities

In addition to the analytical approximations above, we numerically fit two decoders with forms:

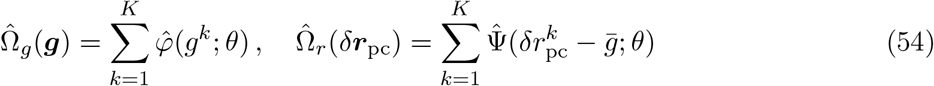

Depending on the decoder class, this scalar signal is taken to be either the raw PC input current *g*^*k*^ or the fixed-point PC firing rates *δ****r***_pc_ under the recurrent dynamics of the optimal decoder. To permit a unified treatment, we may write both in the forms 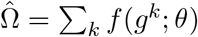, with the understanding that for the recurrent decoder 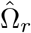, the shifted fixed-point firing rates 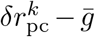 are substituted for *g*^*k*^.

We parameterize the nonlinearity *f* with the following form:

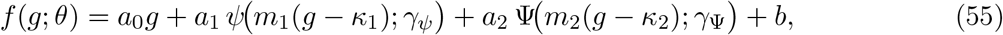

with elementary nonlinearities

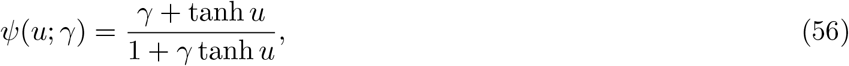

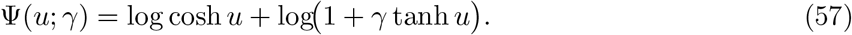

To fit the ten free parameters ***θ*** = {*a*_0_, *a*_1_, *a*_2_, *m*_1_, *m*_2_, *κ*_1_, *κ*_2_, *b, γ*_*ψ*_, *γ*_Ψ_}, we draw *n*_fit_ = 4000 Monte Carlo samples 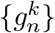 from the class-conditional distributions *P* (***g***|*e*) for *e* = 0 and *e* = 1, and fit *θ* by minimizing the binary cross-entropy

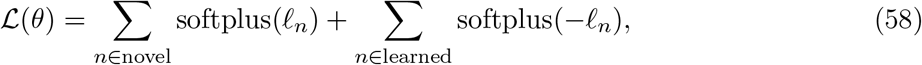

where we have defined the per-pattern logit

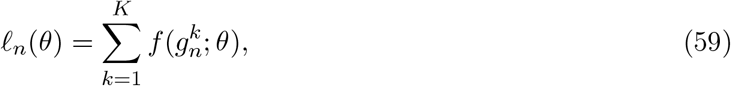

with labels 0/1 for novel/learned patterns. We optimize *ℒ* using `scipy.optimize.minimize` with `method=‘L-BFGS-B’`. We impose lower bounds *m*_1_, *m*_2_ ≥ 10^−3^, leave *η*_ψ_, *η*_Ψ_ unbounded, and use `maxiter=100`, `ftol=1e-8`. To guarantee that 1 + *γ* tanh *u >* 0 for all *u* ∈ ℝ, each shape parameter is reparameterized through an unconstrained latent variable:

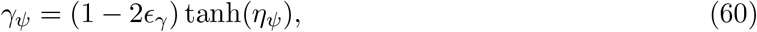

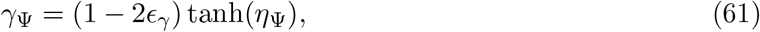

with *ϵ*_γ_ = 10^−4^. The optimizer therefore acts on the unconstrained variables *η*_ψ_, *η*_Ψ_ ∈ ℝ, while always satisfying |*γ*_•_| ≤ 1 − 2*ϵ*_γ_.

### Synchrony and Connectivity Analysis

The central quantity governing pairwise PC statistics is the *pairwise plasticity alignment*, defined as the normalized inner product between columns of the plasticity matrix ***Y*** :

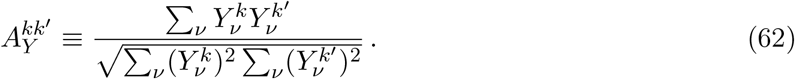

Under heterogeneous plasticity, we represent the plasticity signal matrix as ***Y*** = −*η*_−_***M*** +*η*_+_(**1**_*P ×K*_ − ***M***), where ***M*** ∈ {0, 1}^*P ×K*^ is a binary mask with 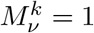 indicating LTD induction in PC *k* during training on pattern *ν*, and 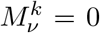 indicating LTP. We define the following per-PC and pairwise statistics of ***M*** :

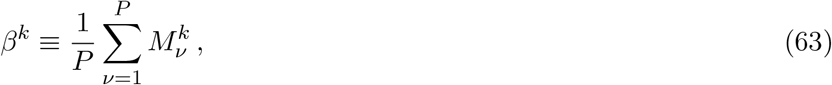

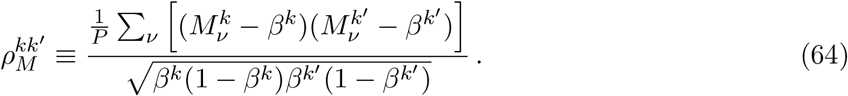

Here *β*^*k*^ is the empirical LTD fraction for PC *k* and 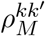 is the Pearson correlation between columns *k* and *k*^*′*^ of ***M***. We note that 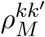 cannot be directly equated with the Pearson correlation of experimentally recorded complex spike trains, since only a subset of complex spikes effectively induce LTD at the PF-PC synapses [7, 42]. We nonetheless expect 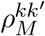 to co-vary with experimentally measured complex spike correlations, and use it as the independent variable in scatter plots relating plasticity history to simple spike synchrony and connectivity overlap. Ignoring fluctuations in *β*^*k*^ (i.e. setting *β*^*k*^ = *β* and treating only fluctuations in 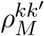), the pairwise plasticity alignment decomposes as

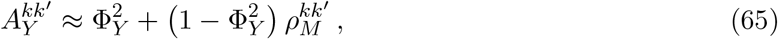

separating into a baseline set by the population-level plasticity ratio Φ_*Y*_ (eq. 7) and pair-to-pair fluctuations driven by empirical correlations between the randomly sampled effective complex spike trains.

#### Temporal Stimulus Protocol and Spike Generation

To study simple spike synchronization we introduce a temporal extension of the model. A simulation interval [0, *T*] is partitioned into consecutive windows whose durations are drawn uniformly from 50 to 100 ms. At the start of each window a new set of PC input currents ***g*** is drawn: with probability *p*_*l*_ from the learned-class distribution *P* (***g***|*e*=1), and with probability *p*_*n*_ = 1 − *p*_*l*_ from the novel-class distribution *P* (***g***|*e*=0). Within each window, PC simple spike trains are generated as independent Poisson processes with a rectified linear rate function:

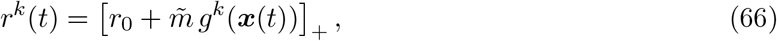

where *r*_0_ is the baseline firing rate, 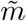 is a gain factor mapping input current to firing rate modulation, and [·]_+_ = max{0, ·} constrains rates to be non-negative.

Cross-correlograms are computed from the resulting spike trains as the empirical probability of a spike in PC *k*^*′*^ at time lag *δt* relative to a spike in PC *k*. The *synchrony score S*^*kk*^*′* quantifies the fractional excess in coincident firing above the rate expected from independent Poisson processes:

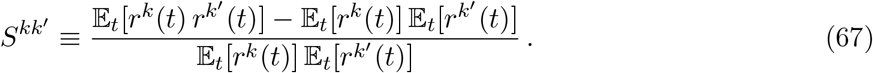

In the SI, we derive the approximate relationship:

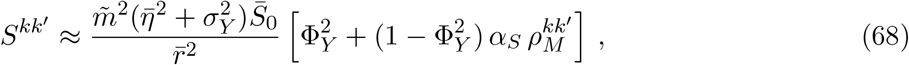

where 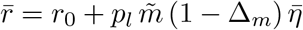 is the mean PC firing rate, and

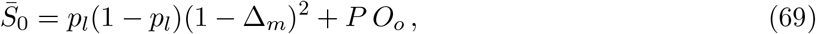

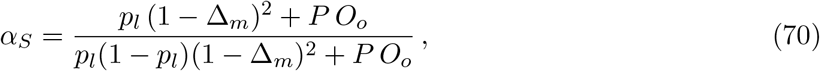

are constants independent of the plasticity distribution. Here *O*_*o*_ = *o/N*_*m*_ + *Q*^2^*/N*_*s*_ is the off-diagonal element of the overlap matrix (eq. 23). The overall prefactor 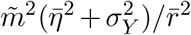 sets the scale of synchrony, while the bracketed factor separates the baseline co-modulation from shared sensory drive (through 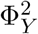) from pair-specific modulations driven by fluctuations in the similarity of the pair’s plasticity histories (through 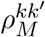). Scatter plots in fig. 5 are generated by sampling rates *r*^*k*^ according to eq. 66 then applying eq. 67. The overlaid analytical comparison is plotted using eqs. 68–70. In simulations, we use *p*_*l*_ = *p*_*n*_ = 0.5.

#### Correlations in Synaptic Weights

We define the learned weight correlation:

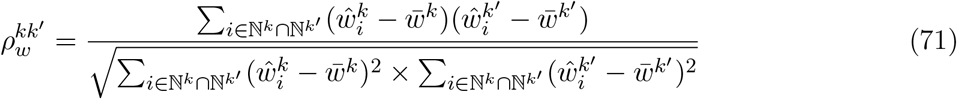

which is the Pearson correlation between the weight vectors learned by PCs *k* and *k*^*′*^, restricted to the subset of parallel fibers ℕ^*k*^ ∩ ℕ^*k*^*′* which form at least one synapse onto both by PCs *k* and *k*^*′*^. It follows directly from the learning rule (eq. 20) that at large *P* the joint distribution of a weight pair 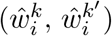 at any PF *i* sampled by PCs *k, k*^*′*^ is bivariate Gaussian with correlation coefficient 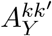. In simulations, 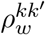 is computed directly from the learned weight matrices (fig. 6B,C,D).

#### Publicly Available Connectome Dataset Analysis

We conduct a novel analysis of the public CB2 connectome analysis dataset published by Nguyen et al. [46], available at `htem/cb2_project_analysis` and online at https://github.com/htem/cb2_project_analysis. The single data file used in our analysis is `analysis/gen_db/pfs/gen_21 0429_setup01_syndb_threshold_10_coalesced.gz`, a PF–PC synapse database. A copy of this dataset, together with our analysis code, is available at https://github.com/benruben87/PSim_HetPlast.git under the directory `pf_pc_effective_weight_portable`.

We filter the database to presynaptic IDs matching the PF families `pf_*` and `ml_pf_*`, and to postsynaptic IDs matching the full-Purkinje regex `pc_\d+`. For each PF–PC pair with at least one synapse, we define 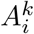 as the sum of the `mesh_area` values across all synapses between PF *i* and PC *k*. For an unordered pair of Purkinje cells (*k, k*^*′*^), let

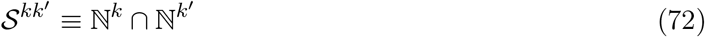

denote the set of PFs shared by the two cells. We then define the experimental Pearson correlation as

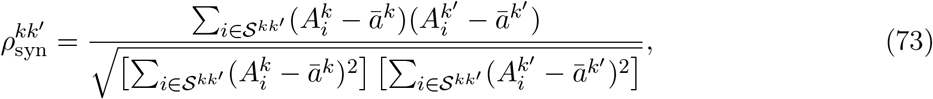

where

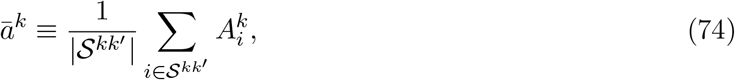

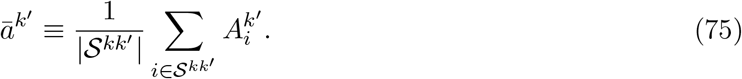

To reduce finite-sampling noise, we restrict the analysis to the 181 PC pairs for which |S^kk^*′* | ≥ 50. We further confirm that pair-to-pair variation in 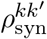 is not explained by variation in the number of observed shared PF inputs (fig. S4F).

To assess significance, we use a one-sided Monte Carlo permutation test on the median pairwise correlation. For each qualifying PC pair (*k, k*^*′*^) and each shuffle trial, we independently randomly permute the entries of the two shared-PF weight vectors

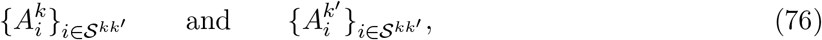

thereby preserving the marginal distribution of synaptic areas for each PC within the pair while destroying the PF-by-PF correspondence between the two cells. We then recompute 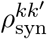 for every qualifying pair and summarize each shuffle by the median across pairs,

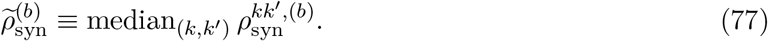

Using *n*_trials_ = 2000 shuffle trials yields a null distribution for the across-pair median correlation. Let

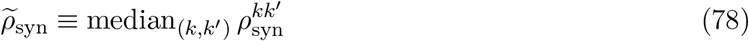

denote the observed median across qualifying PC pairs. The upper-tail Monte Carlo *p*-value is then

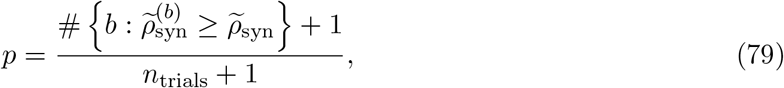

We find in fig. S4C that 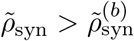 for all 2000 shuffle trials, giving significance *p* ≤ 5 × 10^−4^.

### Extension to Block-Structured Plasticity

To model the scenario where multiple PCs receive climbing fibers originating from the same inferior olive neuron, we introduce a *block-structured plasticity* scheme. The ensemble of *K* PCs is partitioned into *N*_*b*_ disjoint sibling blocks 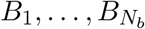, each of equal size *K*_*b*_ = *K/N*_*b*_. Within each block *b*, all PCs share the same plasticity signal:

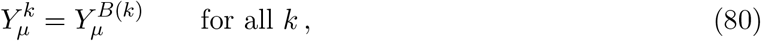

where *B*(*k*) denotes the block index of PC *k*. The block-level signals 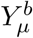 are drawn i.i.d. from the heterogeneous plasticity distribution (eq. 6). We may organize these into a matrix 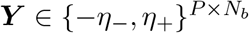. The expected covariance matrix 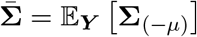 acquires a three-level structure:

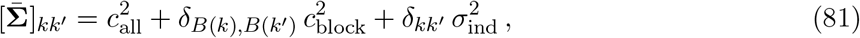

with components

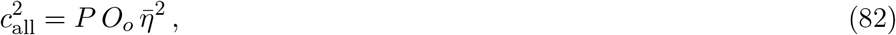

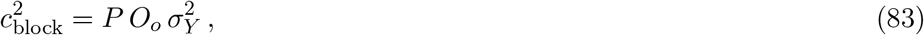

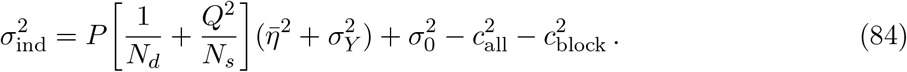

Here *O*_*o*_ = *o/N*_*m*_ + *Q*^2^*/N*_*s*_ is the off-diagonal element of the overlap matrix (eq. 23), and 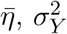 are the mean and variance of the plasticity signal, defined in section 2.3.

Next, we define the block averages 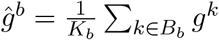. Under the expected-covariance approximation (replacing **Σ**_(−µ)_ with 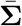 as in section 3), these block averages are distributed as:

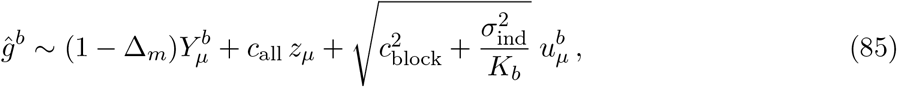

This is the same distributional form as the PC inputs under fully heterogeneous plasticity. In the SI, we show that the optimal decoder Ω_opt_ applied to the full ***g*** vectors under block-structured plasticity reduces exactly to the optimal decoder under heterogeneous plasticity applied to these block-averages 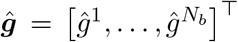, with parameter substitutions 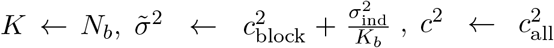. All derived approximate optimal decoders can then be re-purposed for the case of block-structured plasticity when applied to the block averages ***ĝ*** with these parameter substitutions. In all simulations with block-structured plasticity, we apply the moment-matched approximation to the optimal decoder (eq. 52).

## Code Availability

All code used to generate the figures in this study is publicly available on GitHub at https://github.com/benruben87/PSim_HetPlast.git.

## Acknowledgements

C.P. is supported by an NSF CAREER Award (IIS-2239780), DARPA grants DIAL-FP-038 and AIQ-HR00112520041, the Simons Collaboration on the Physics of Learning and Neural Computation, and the William F. Milton Fund from Harvard University. This work has been made possible in part by a gift from the Chan Zuckerberg Initiative Foundation to establish the Kempner Institute for the Study of Natural and Artificial Intelligence. B.S.R. thanks William Qian and Jacob Zavatone-Veth for thoughtful discussions and comments on this manuscript.

---

**Figure S1:**
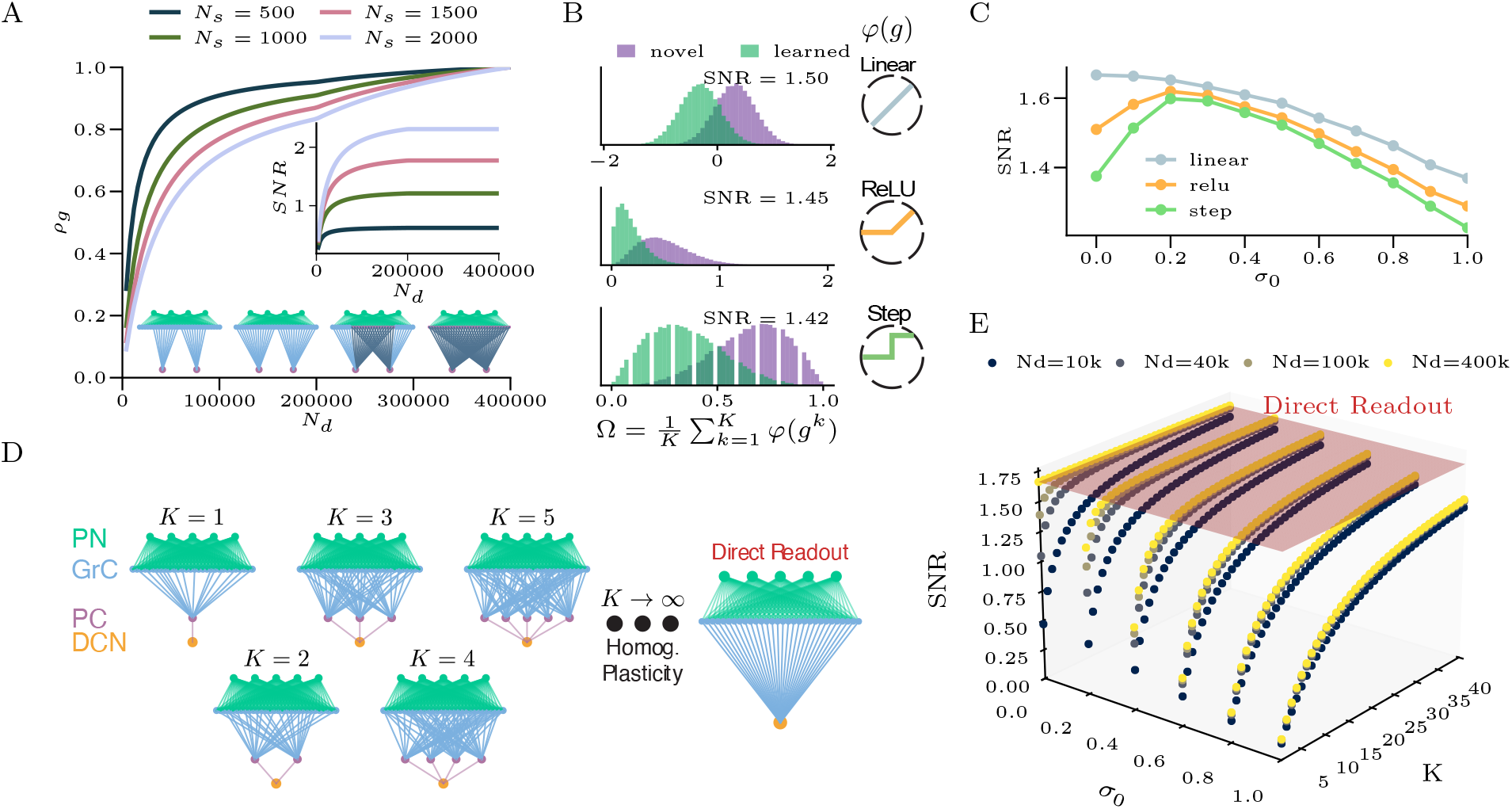
Ensemble utility is limited under homogeneous LTD and subsampled connectivity. **(A)** Correlation coefficient *ρ*_*g*_ between two PCs reading out from *N*_*m*_ = 400,000 PFs as a function of PC dendritic-tree size *N*_*d*_. Line color indicates stimulus dimensionality *N*_*s*_. We set *o* = *o*_min_, *σ*_0_ = 0. Inset: SNR versus *N*_*d*_, confirming monotonic improvement with increasing PC size. **(B)** Histograms of the decoder output Ω(***g***) = *K*^−1^ ∑_k_ *φ*(*g*^*k*^) for separable decoders with three nonlinearities *φ*, applied to novel (purple) and learned (green) pattern distributions. The linear decoder outperforms both nonlinear decoders, in agreement with Eq. S6.11. **(C)** SNR as a function of intrinsic noise *σ*_0_, confirming that linear decoding is optimal across noise levels. **(D)** Schematic: ensemble behavior as *K* → ∞ under homogeneous plasticity converges to a “direct readout” architecture that samples all available PFs. **(E)** SNR for PC ensembles under homogeneous plasticity and linear decoding. The red plane shows the SNR achieved by the direct readout, which upper-bounds ensemble SNR under homogeneous plasticity. Parameters for (B, C): *σ*_0_ = 0.7, *K* = 40, *N*_*m*_ = 400,000, *N*_*d*_ = 200,000, *o* = 1, *f* = 0.05, Δ_*x*_ = 0.10. Parameters for (E): *N*_*m*_ = 400,000, *N*_*s*_ = 1,400, *f* = 0.05, Δ_*x*_ = 0.10.

**Figure S2:**
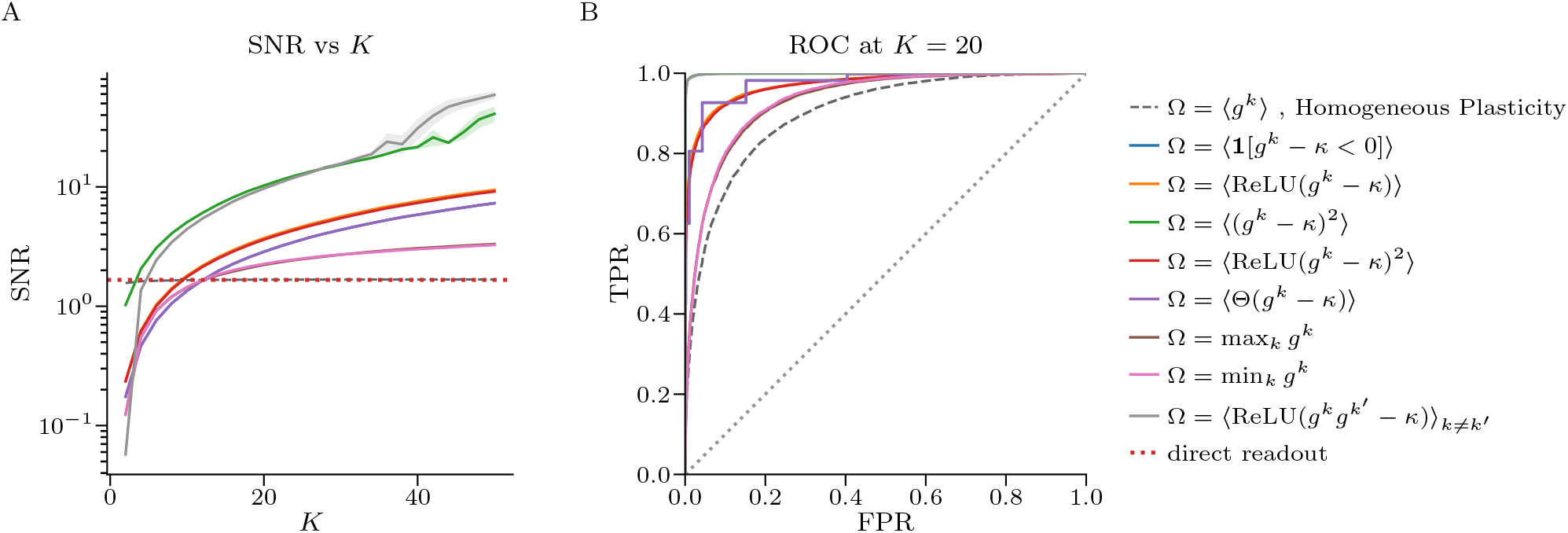
SNR improvement from balanced heterogeneous plasticity is robust to the choice of nonlinear decoder. PC ensembles were trained under heterogeneous plasticity and stimulus class decoded using a variety of nonlinear decoders. **(A)** SNR_eff_ versus the PC ensemble size *K* with a variety of decoding functions Ω(*g*^1^, …, *g*^*K*^). Here, angular brackets ⟨· · · ⟩ denote an average over *k* ∈ {1, …, *K*} unless otherwise indicated. For decoders with a threshold, the threshold *κ* was swept and the curve shows the SNR at the trial-averaged optimum *κ*^∗^(*K*). Shaded bands show 1 ± SEM across 10 trials. For heterogeneous plasticity we use *β* = 0.5, *η*_+_ = *η*_−_ = 0.5. For all networks *σ*_0_ = 0.1, and *N*_*d*_ = 200,000. The gray dotted line shows SNR for the linear decoder trained with homogeneous plasticity (*β* = 1, *η* = 1), and the horizontal red dotted line marks the direct-readout SNR. **(B)** ROC curves for every decoder in a network with ensemble size *K* = 20. For decoders with threshold *κ*, the threshold is fixed to the *κ*^∗^ that maximized trial-averaged SNR at *K* = 20. The diagonal is chance.

**Figure S3:**
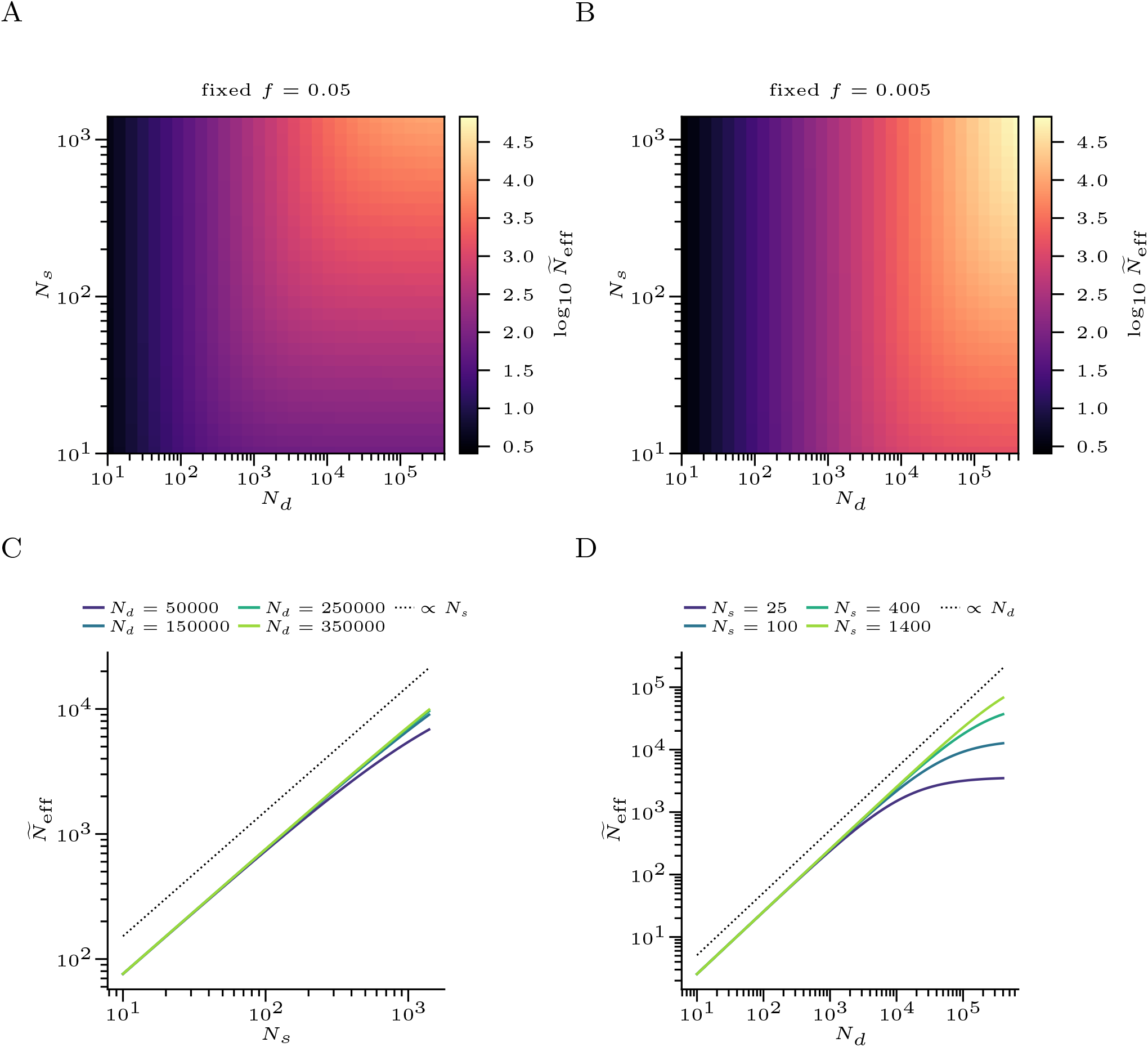
Scaling behavior of effective dimensionality 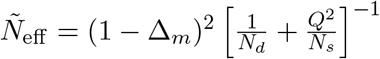. (A) *Ñ*_eff_ as a function of *N*_*d*_ and *N*_*s*_ in the case where GrC coding level is held fixed at *f* = 0.05. (B) Same as (A), except the GrC coding level is set to *f* = 0.005. (C) Cross-sections of the heatmap in A showing that 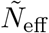 grows linearly with *N*_*s*_ until *N*_*d*_ becomes the bottleneck to dimensionality. (D) Cross-sections of heatmap in B showing that in the sparse-coding regime 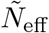 grows linearly with *N*_*d*_ until a plateau determined by *N*_*s*_. All panels use input cluster noise Δ_*x*_ = 0.10

**Figure S4:**
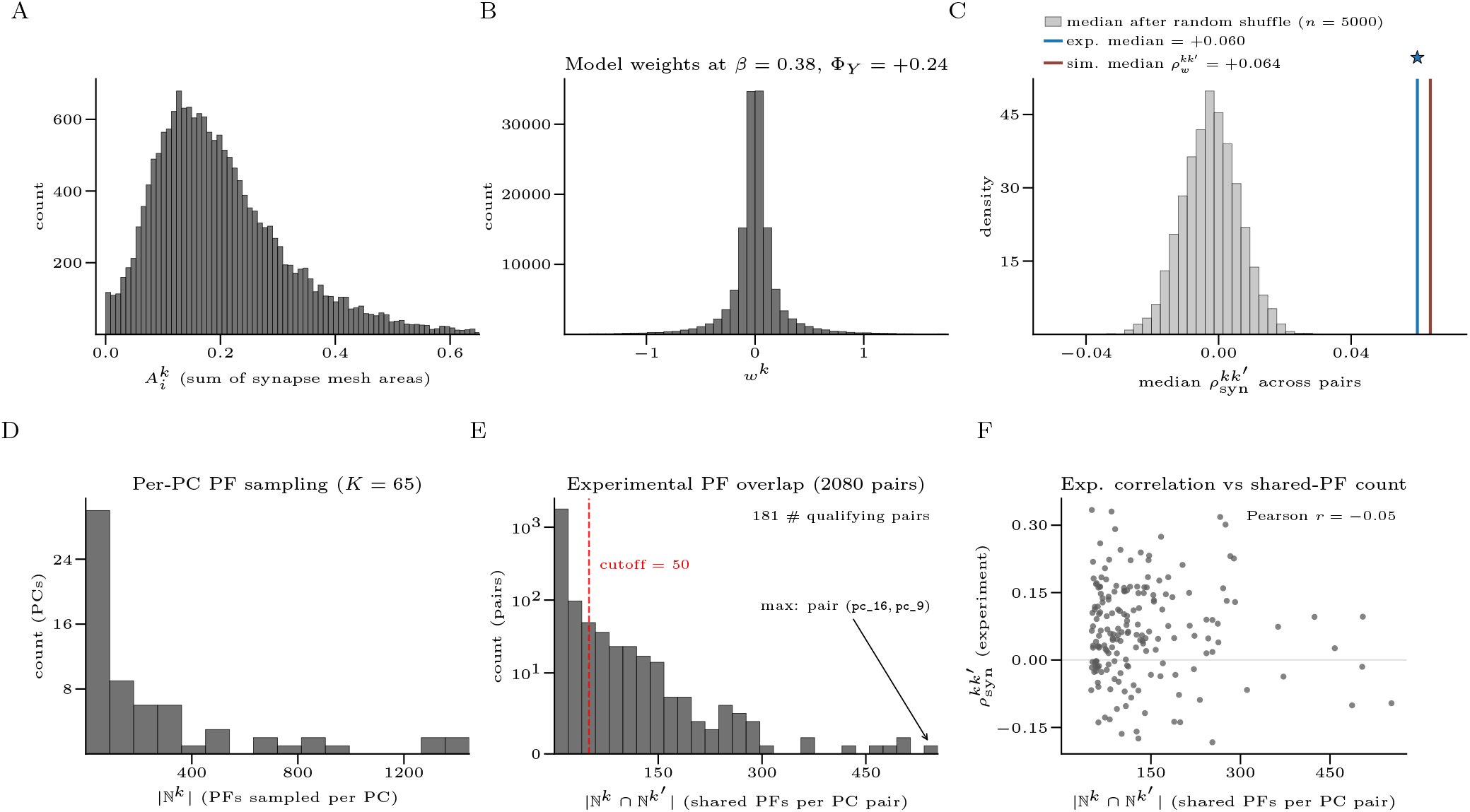
Extended data for experimental weight correlation analysis. (A) Distribution of total synaptic connection areas 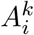 for all measured PF–PC pairs (*i, k*). (B) For comparison, we show the distribution of learned synaptic weights 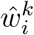 following heterogeneous plasticity training at the operating point indicated in the title. Simulated weight distributions are Gaussian and centered at zero. (C) distribution of median 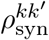 following null shuffle procedure (Methods). (D) Distribution of number of PFs ℕ^*k*^ within the reconstructed EM volume sampled by each PC *k*. (E) Distribution in the number of PFs |ℕ^*k*^ ∩ ℕ^*k*^*′* | jointly sampled by each pair of PCs *k* ≠ *k*^*′*^ visible within the reconstructed EM volume. To reduce finite-sample noise in measured weight correlations, we consider only pairs which share at least 50 PF inputs, yielding 181 qualifying pairs of PCs. (F) Scatter plot showing 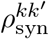 against |ℕ^*k*^ ∩ ℕ^*k*^*′* | for each PC pair included in the pairwise analysis. Similarities in measured synaptic weights are not explained by co-variation with the number of jointly sampled PFs.

## S1 Supplementary Information

This Supplementary Information is organized as follows. In Sec. S2, we establish notation and provide a self-contained summary of the model architecture, task, and learning rules. In Sec. S3, we derive the statistics of learned PC input currents under general, homogeneous, and heterogeneous plasticity. Secs. S4 and S5 present the SNR and capacity analyses under homogeneous and heterogeneous plasticity, respectively. Sec. S6 contains derivations of the approximate optimal decoders, including the saddle-point and moment-matched approximations and the recurrent-circuit interpretation. Sec. S7 derives the Purkinje cell correlation and connectivity results. Sec. S8 extends the model to multi-shot learning. Sec. S9 extends the model to incorporate “block-structured plasticity” in which multiple PCs receive climbing fibers from the same IO neuron, and derives the optimal decoding strategy in that case. Sec. S10 considers the effects of random subsampling of PFs, and analyzes the trade-off between ensemble size and the size of each PC in the case of heterogeneous plasticity learning. Sec. S11 shows that centered weight updates can be decomposed into an uncentered, instructed plasticity signal and a centering homeostatic plasticity. Finally, in Sec. S12, we derive rigorous bounds on network capacity which verify the scaling dependencies on *K* and *N*_eff_ obtained using non-rigorous methods in Sec.S5.

## S2 Preliminaries

Throughout this supplement, we use the following standard notation. The Gaussian tail function is

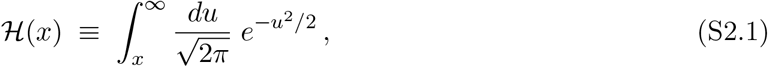

and the standard Gaussian probability density function and cumulative distribution function are denoted

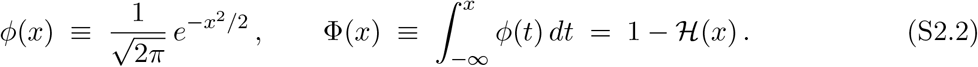

The standard Gaussian measure is written 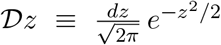, and the symbol ⊙ denotes the Hadamard (elementwise) product of matrices. The indicator function **1**{·} equals 1 when its argument is true and 0 otherwise, and Θ[·] denotes the Heaviside step function. We write **1**_*K*_ for the *K*-dimensional vector of ones and ***I***_*K*_ for the *K* × *K* identity matrix.

### S2.1 Summary of Task and Architecture Description

In this section, we provide a detailed, self-contained description of our modeling procedure, including the model architecture, task, learning rule, and performance evaluation metric.

#### S2.1.1 Model Architecture

The input 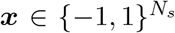 is projected through a random matrix 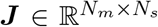 into a mixed layer (granule cells) of *N*_*m*_ binary neurons ***m***(***x***) = Θ[***Jx*** − *T*]. Here Θ[· · ·] is the Heaviside step function and the threshold *T* is selected so that the mixed-layer patterns have, on average, a fraction of *f* active neurons in response to a pattern drawn randomly from the input distribution. A layer of *K* Purkinje cells (PCs) reads out from the mixed layer through learned weights ***ŵ***^*k*^, each sampling *N*_*d*_ PFs, giving *g*^*k*^(***x***) = ***ŵ***^*k*^ · (***F*** ^*k*^***m***(***x***) − *f*). Here 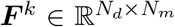 is a projection matrix which selects the subset of *N*_*d*_ GrCs which are sampled by PC *k*. The PC input currents {*g*^1^(***x***), …, *g*^*K*^(***x***)} are decoded by a deep cerebellar nucleus (DCN) output neuron computing a function Ω(***g***), which we leave in general form. In sec. S5, we study network performance with “separable” decoders of the form Ω(***g***) = ∑_k_ *φ*(*g*^*k*^) with *φ*(*g*) ∈ {*g*, (*g* − *κ*)^2^, Θ[*κ* − *g*]}. In §S6 we derive optimal and near-optimal decoding strategies from optimization principles.

#### S2.1.2 Associative Learning Task

The network must learn to respond selectively to *P* stimulus clusters in the space of *N*_*s*_-dimensional binary inputs 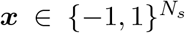. Each cluster *µ* has a randomly chosen center 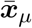, and patterns belonging to cluster *µ* are generated by flipping each element of 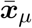 independently with probability Δ_*x*_*/*2. Patterns from learned clusters are assigned the label *e* = 1; novel patterns drawn uniformly at random are assigned *e* = 0.

#### S2.1.3 Two-Factor Learning Rule

We consider the following general form for the learned PC weights:

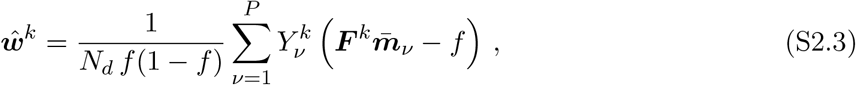

This learning rule can be viewed as the outcome of an online learning process in which one pattern from each cluster (taken to be the cluster center 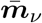 for mathematical convenience) is presented to the network sequentially, and in response each PC *k* = 1, …, *K* receives weight updates of the form:

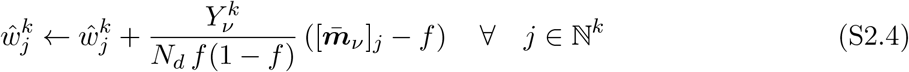

as shown in the main text. For the analysis in S3.1, we leave the plasticity signals 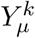 in general form, later specializing to the cases of homogeneous and heterogeneous plasticity.

#### S2.1.4 Performance Evaluation Metric

Let Ω(***g***) ∈ ℝ be any scalar score derived from the PC input currents ***g***, and let Ω_*ℓ*_ and Ω_*n*_ denote independent draws from its distribution under learned and novel inputs, respectively. Given a threshold *θ*, the true-positive rate TPR(*θ*) = Pr(Ω_*ℓ*_ ≥ *θ*) and false-positive rate FPR(*θ*) = Pr(Ω_*n*_ ≥ *θ*) trace out the ROC curve as *θ* varies. The area under the ROC curve,

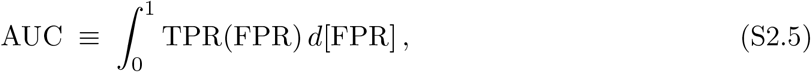

is a threshold-free performance measure ranging from 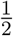 (chance; In cases where AUC *<* 1*/*2, an AUC *>* 1*/*2 can be recovered by a simple sign change Ω(***g***) → − Ω(***g***)) to 1 (perfect) For calculations throughout this work, we make use of the standard result [96] that the AUC equals the ranking probability:

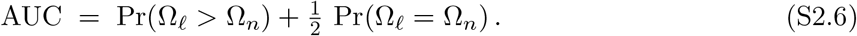

In the special case of Gaussian decoder outputs, 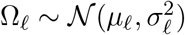 and 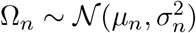, this reduces to 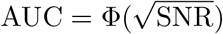, where

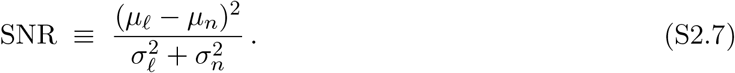

For non-Gaussian decoder outputs, we define an *effective* SNR via the inverse mapping:

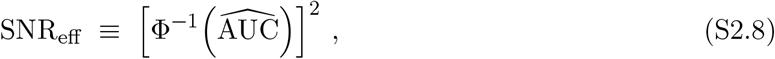

where 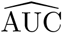 is estimated empirically from Monte Carlo samples using the ranking definition above. By construction, SNR_eff_ reduces to the standard SNR whenever the output distributions are Gaussian, and preserves the monotonic mapping 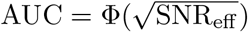 in all cases.

### S2.2 Review of Mixed-Layer Statistics

In the first step of our model architecture, the inputs 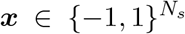 are projected through a random Gaussian weight matrix, and followed by an elementwise application of a step function with threshold *T*, to achieve an average sparsity *f* in the mixed (GrC) layer. The effects of this “sparse expansion” step were studied by Babadi and Sompolinsky [17], and can be summarized using two population-level statistics which govern downstream classification performance. The first is the mixed-layer cluster size:

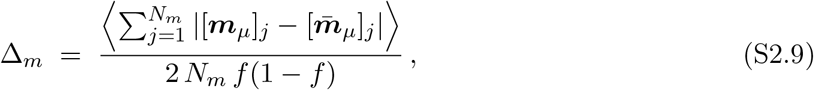

where ⟨·⟩ represents an average over realizations of *m*_*µ*_ in each cluster *µ*, as well as an average over the clusters *µ*. The denominator represents the mean Hamming distance between two independent random sparse patterns with activity *f*, so that Δ_*m*_ varies from 0 (when Δ_*x*_ = 0) to 1. With a random projection matrix and *N*_*s*_, *N*_*m*_ ≫ 1, Babadi and Sompolinsky [17] show that the mixed-layer cluster size is given by:

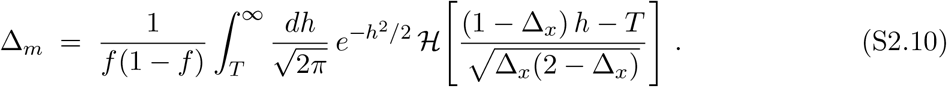

In the regime of small input clusters (Δ_*x*_ ≪ 1) and strong sparsity (*f* ≪ 1), we have the approximation 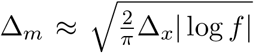. The second important mixed-layer statistic is the “excess overlap” between the representations of inputs from *different* clusters. Specifically, we define the *centered overlap*:

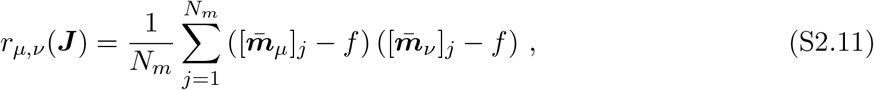

which depends on the input clusters *µ* ≠ *ν*, as well as the particular realization of the random expansion matrix ***J***. It is clear that E_***J***_ [*r*_*µ,ν*_(***J***)] = 0, and Babadi and Sompolinsky [17] calculate the variance as

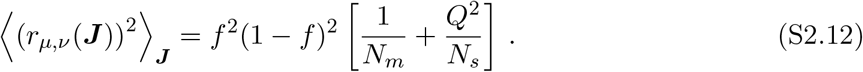

where the excess overlap parameter *Q* = *Q*(*f*) is given by the following:

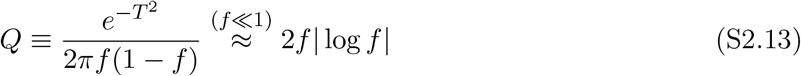

Together, Δ_*m*_ and *Q* fully characterize the mixed-layer expansion for the purposes of downstream classification performance (see §S3).

### S2.3 Minimally Overlapping Homogeneous Subsets

Here, we derive bounds on the overlap between uniformly structured subsets of a finite set. This result will be used later to put bounds on the amount of overlap between subsets of PFs sampled by PCs in the case where *N*_*d*_ *< N*_*m*_. Let *S* be a ground set with |*S*| = *M*, and let *s*_1_, …, *s*_*K*_ be *K* subsets of *S*, each of cardinality *N*. Define the pairwise overlap *O*_*kk*_*′* = |*s*_*k*_ ∩ *s*_*k*_*′* | and the normalized overlap parameter *o* ≡ *O*_*kk*_*′ M/N* ^2^ for *k* ≠ *k*^*′*^, so that *o* = 1 corresponds to the overlap expected from independent uniform random sampling.

#### Incidence function

For each *x* ∈ *S*, define *I*(*x*) ≡ |{*k* : *x* ∈ *s*_*k*_}|. Summing over all *x* ∈ *S*:

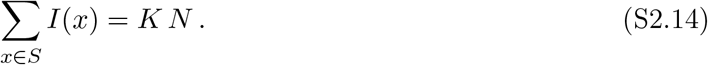

Moreover, 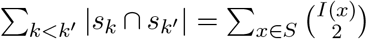, since each *x* contributes to |*s*_*k*_ ∩ *s*_*k*_*′* | exactly when *x* ∈ *s*_*k*_ and *x* ∈ *s*_*k*_*′*

#### Minimizing overlaps

Since 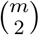 is convex in *m*, the sum 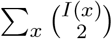 is minimized by distributing the values *I*(*x*) as evenly as possible across *S*. In the limit of large *M*, set *I*(*x*) ≈ *λ* ≡ *KN/M* for all *x* ∈ *S*. Then:

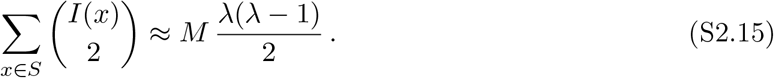

Since there are 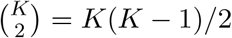 pairs, the average pairwise intersection size is:

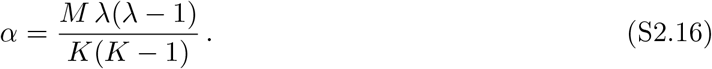

Substituting *λ* = *KN/M* and *o* = *α M/N* ^2^:

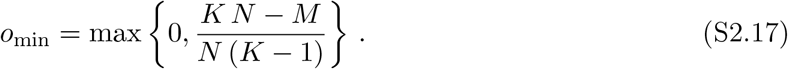

The maximum overlap occurs when all subsets are identical, giving *o*_max_ = *M/N*.

## S3 Derivation of PC Response Statistics

In this section, we derive the statistics of the Purkinje cell input currents ***g*** after training with the two-factor learning rule (eq. S2.3). We will organize the plasticity signals into a matrix ***Y*** ∈ ℝ^P *×*K^ such that 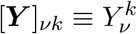. In section S3.1, we treat the case where plasticity signals 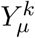 are left in general form, then consider the special cases of homogeneous and heterogeneous plasticity in sections S3.3 and S3.4.

### S3.1 General Case

After training on *P* cluster centers according to the learning rule eq. S2.3, the input current to PC *k* in response to a test pattern ***x***_*µ*_ from cluster *µ* decomposes into signal and noise:

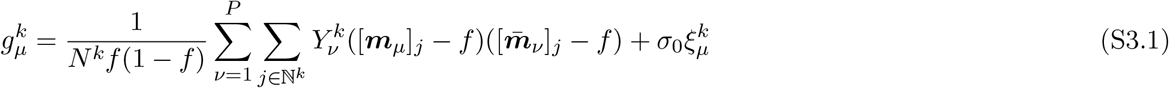

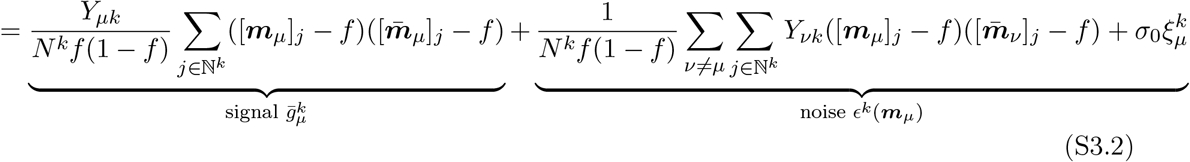

Where 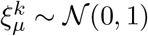 and *σ*_0_ is the variance of an “intrinsic” noise in the Purkinje Cell which results in its baseline spiking activity. Using the identity 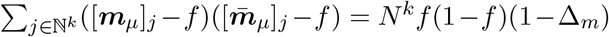, we have the following equation for the “signal” strength in each decoder:

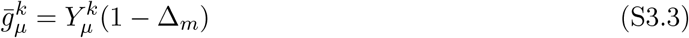

Next, we calculate the variance and covariance of the decoder preactivations.

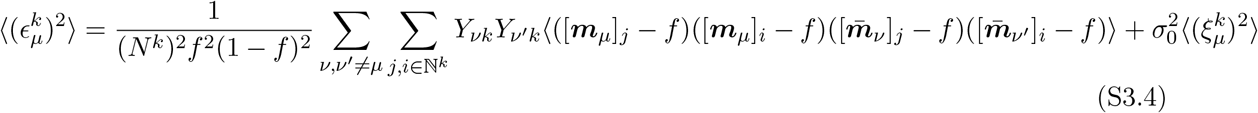

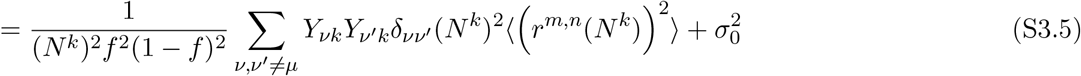

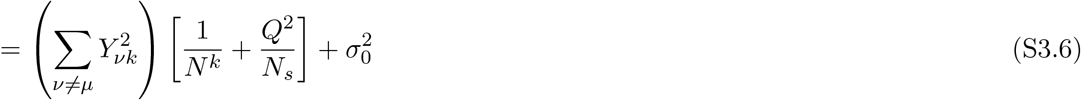

for *k*≠ *k*^*′*^:

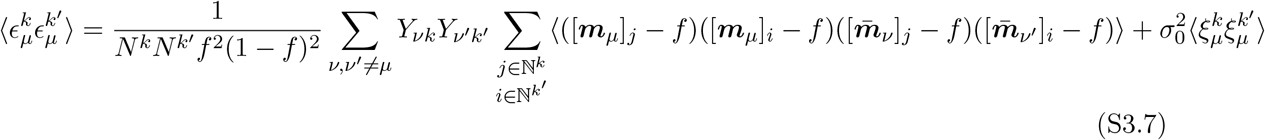

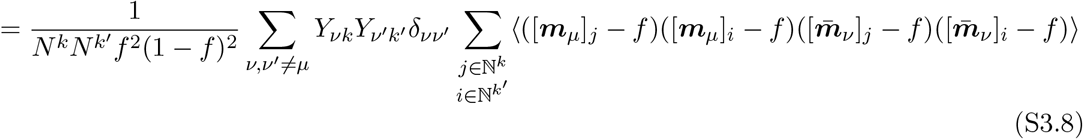

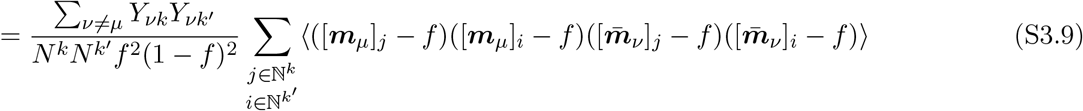

To evaluate the inner sum over ℕ^*k*^ and ℕ^*k*^*′*, we need to handle the overlap between these sets with care. We define the sets 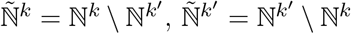, and 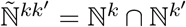. The sum then breaks into four contributions:

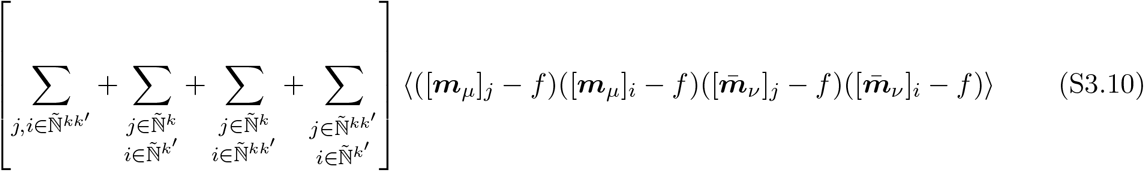

The first term is easily given by

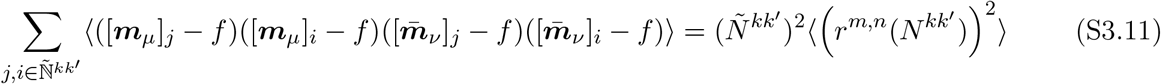

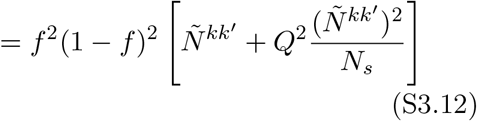

To calculate the remaining terms, we notice that we can rewrite the sums as follows:

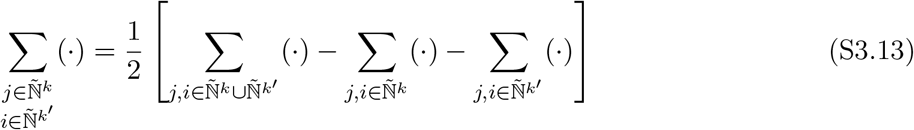

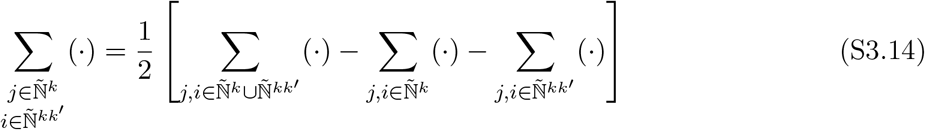

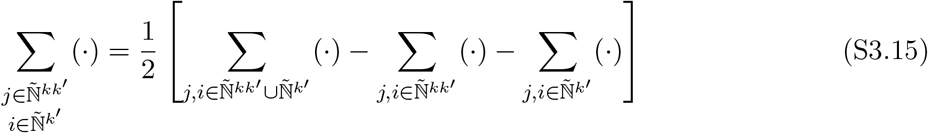

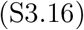

because the sets 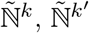, and 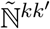 are mutually exclusive. We can then simplify the first inner sum as:

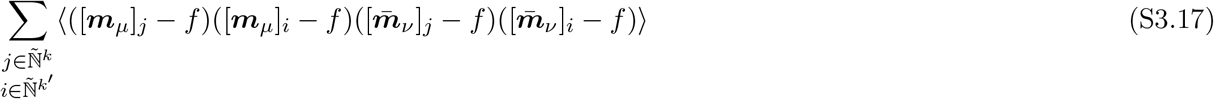

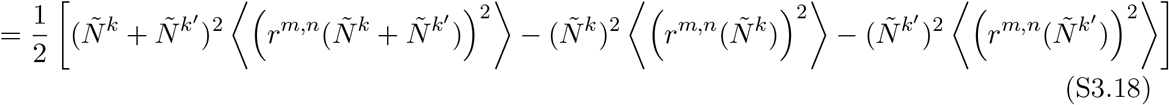

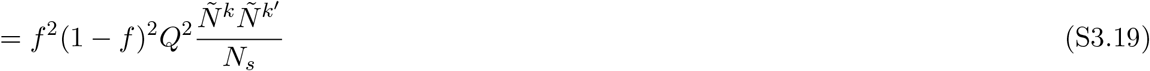

By analogy, we also have

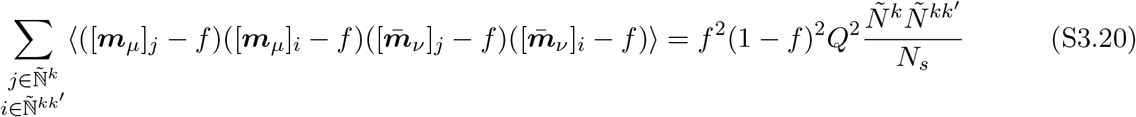

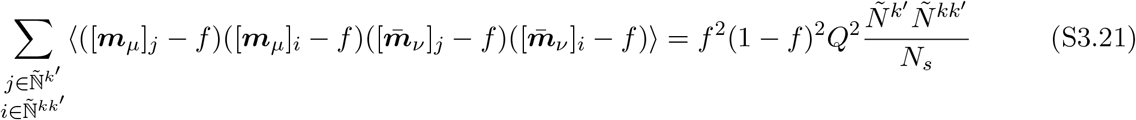

where 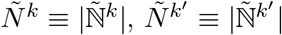, and 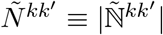. Putting these together, we have

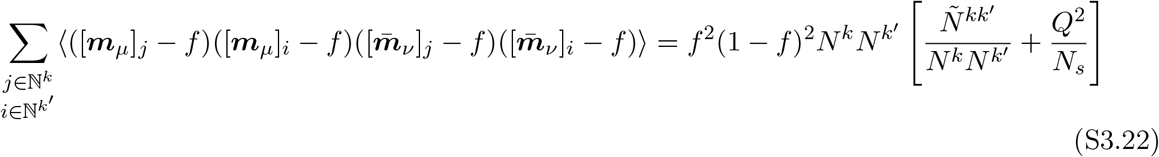

and finally,

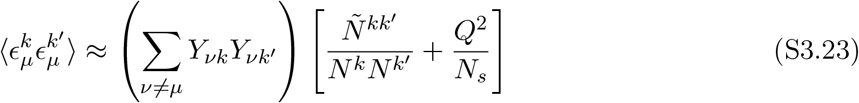

Putting these together, we have that in response to patterns ***x***_*ν*_, generated by flipping elements of 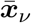 with probability Δ_*x*_*/*2, we have

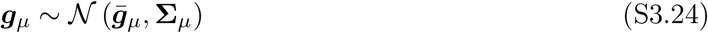

where 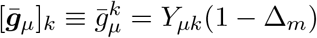. Furthermore, we have

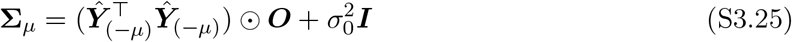

Where ***Ŷ*** ∈ ℝ^(P −1)*×*K^ is the plasticity-signal matrix ***Y*** with row *µ* removed, and we have defined ***O*** ∈ ℝ^K*×*K^ as follows:

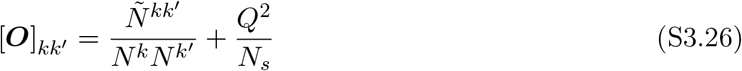

#### Homogeneous Overlap Assumption

From this point forward, we assume homogeneous overlap statistics:

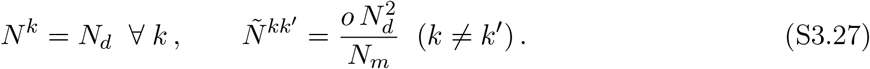

It follows from the standard combinatorial arguments given in section S2.3 that the overlap fraction *o* is constrained to lie in the following range:

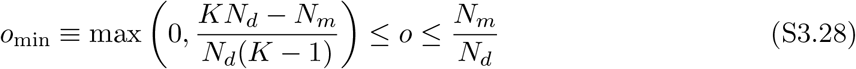

Under the homogeneous overlap assumption, the overlap matrix takes the form:

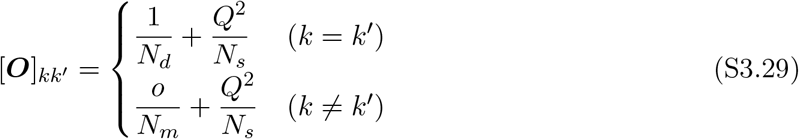

Defining *O*_*o*_ ≡ *o/N*_*m*_ + *Q*^2^*/N*_*s*_ and *κ*_*d*_ ≡ 1*/N*_*d*_ − *o/N*_*m*_, we can write [***O***]_*kk*_*′* = *O*_*o*_ + *δ*_*kk*_*′ κ*_*d*_. In terms of the effective input dimensionality *N*_eff_ and the fraction of independent variance ℵ:

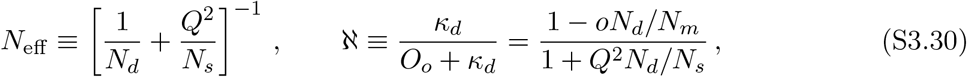

the overlap matrix becomes:

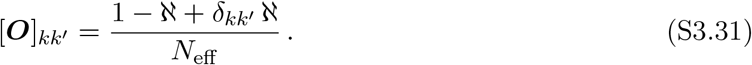

### S3.2 *N*_eff_ as effective dimensionality

Here, we justify the terminology of referring to *N*_eff_ as the “effective dimensionality” of the GrC population code. Specifically, consider the centered covariance matrix of the GrC inputs to a particular PC over the response to a random input ***x***drawn uniformly from 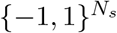.

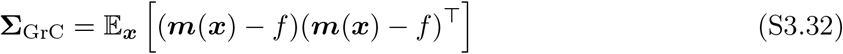

with eigenvalues 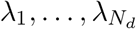. We then define the effective dimensionality via the participation ratio:

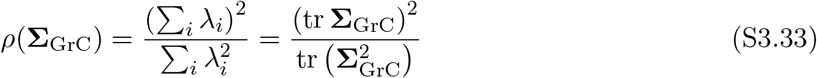

and claim that for *N*_*d*_, *N*_*m*_ ≫ 1, we have the asymptotic equivalence *ρ*(**Σ**_GrC_) ≍ *N*_eff_. Expanding in terms of the components of **Σ**_GrC_, we have

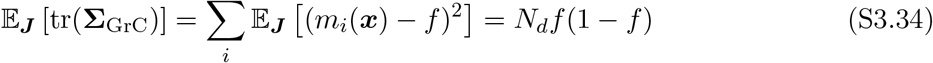

and

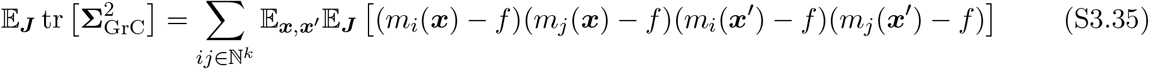

The expectation is equivalent to the expected value in eq. S3.22, with *N* ^*k*^ = *N* ^*k*^*′* = *N* ^*kk*^*′* because ***x***_*µ*_ and 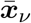 are statistically equivalent to two randomly drawn patterns ***x, x***^*′*^. We therefore have:

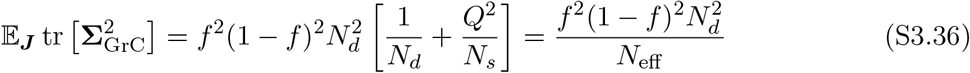

Assuming the numerator and denominator of the participation ratio concentrate separately over ***J***, we arrive at

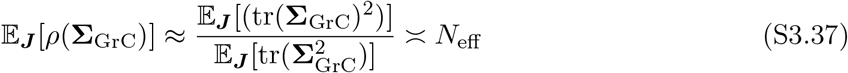

### S3.3 Homogeneous Plasticity Statistics

Homogeneous LTD is recovered by setting 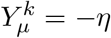 for all *µ, k*, giving ***Y*** ^⊤^***Y*** = *η*^2^*P* **1**_*K×K*_. Under the homogeneous overlap assumption and approximating *P* − 1 ≈ *P* (valid for *P* ≫ 1), the covariance matrix becomes class-independent:

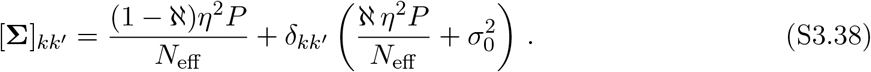

Defining the total variance 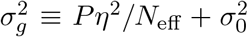 and the correlation coefficient 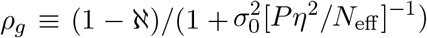, the components of ***g***_*µ*_ may be equivalently represented as:

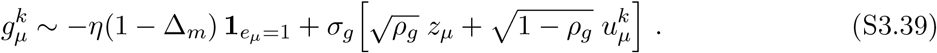

where *z*_*µ*_ ~ *N*(0, 1) is a shared global fluctuation and 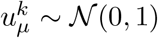 are independently sampled for each PC.

### S3.4 Heterogeneous Plasticity Statistics

In the case of heterogeneous plasticity, we have

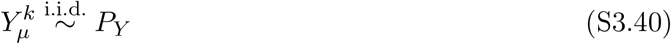

for some plasticity distribution *P*_*Y*_. We define the first two cumulants of the plasticity distribution as

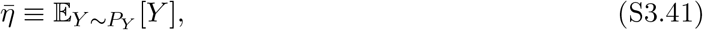

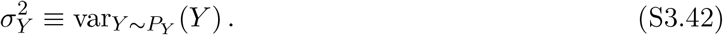

For the one-shot learning rule analyzed in the main text we have,

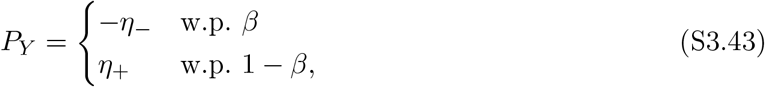

with first two cumulants given by

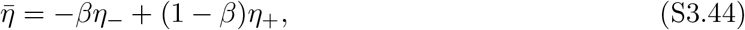

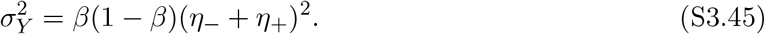

Because ***Y*** is drawn independently each time the network is trained, no exact simplification of the PC response statistics beyond eq. S3.24 is possible. Rather, we expect that for sufficiently large networks, network performance will become consistent, or “concentrate” for typical-case realizations of ***Y***.

## S4 SNR and Capacity under Homogeneous Plasticity

Here, we derive and analyze the analytical expression for the SNR under homogeneous LTD (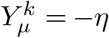 for all), in which case the response statistics of the learned PC inputs are described by eq. S3.39. We consider here the linear decoder 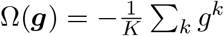. As we show in §S6.1, this corresponds to the optimal decoder in the homogeneous plasticity setting. The SNR and capacity results derived here therefore characterize the best achievable performance under homogeneous plasticity.

### S4.1 SNR Formula

In the homogeneous PC response statistics (eq. S3.39) each PCs input *g*^*k*^ is a linear combination of an input-class-dependent shift, a shared global fluctuation, and an independent fluctuation (represented as 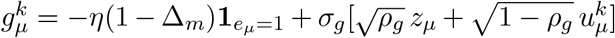. We may then represent the output of the linear decoder as follows:

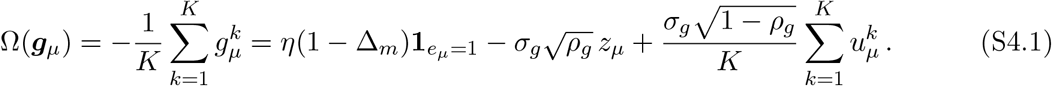

The mean Ω receives the same contribution from each PC due to the shared fluctuation *z*_*µ*_, plus the average of the independent random components *u*^*k*^. Combining the fluctuations using the additive property of the Gaussian, the decoder output distributions are then described as :

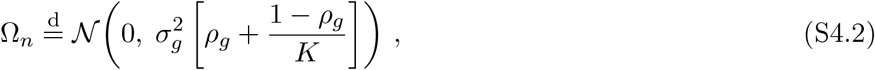

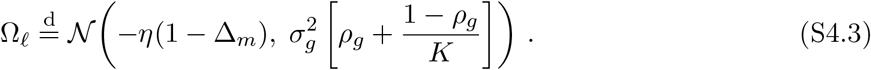

Since both distributions are Gaussian with equal variance, the SNR (see §S2.1.4) is:

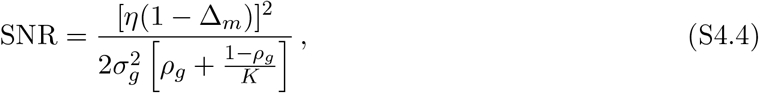

### S4.2 SNR Monotonicity and Direct Readout Upper Bound

For the analysis which follows, it is useful to write the SNR in the following expanded form, obtained by substituting the definitions of *σ*_*g*_ and *ρ*_*g*_, and ℵ:

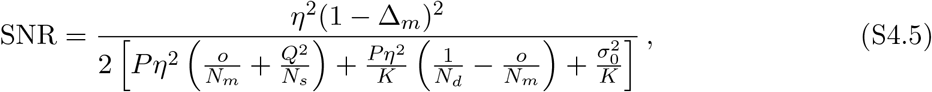

The following facts follow directly from this form.

#### Lemma 1.

SNR *increases monotonically with ensemble size K*.

#### Lemma 2.

SNR *decreases monotonically with the overlap fraction o*

At fixed *K* and *N*_*d*_, SNR is optimized by setting *o* = *o*_min_. In this case, we arrive at the following:

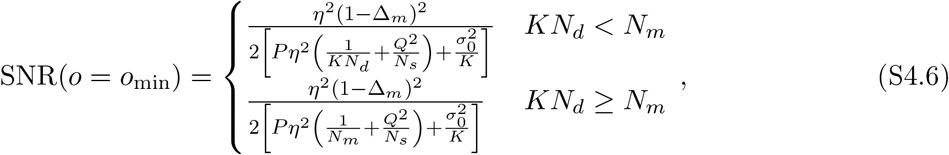

The following lemma is then easily verified:

#### Lemma 3.

SNR *increases monotonically with PC size N*_*d*_, *both when the overlap fraction o is fixed as N*_*d*_ *varies, and when o is set to its optimal (minimum) value as N*_*d*_ *varies*.

For fixed overlap fraction, this can be verified directly from eq. S4.5. For optimal *o*(= *o*_min_), this is verified by noticing that eq. S4.6 is monotonically increasing with *N*_*d*_.

An important point of reference is the “direct readout” architecture in which the network output is read out directly from all available PFs by a single decoding neuron, effectively skipping the PC layer. We can write the SNR achieved by this architecture under homogeneous plasticity by setting *K* = 1, *N*_*d*_ = *N*_*m*_, and *σ*_0_ = 0 (as there are no PCs present to contribute intrinsic noise). We arrive at the following:

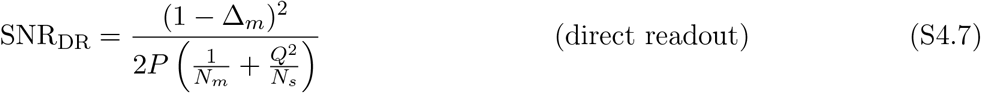

Comparing with eq. S4.6, which upper bounds SNR for ensembles, the following is easily verified:

#### Lemma 4.

*For any ensemble trained under homogeneous plasticity*, SNR ≤ SNR_DR_, *with equivalence when K* → ∞, *or when σ*_0_ = 0 *and KN*_*d*_ ≥ *N*_*m*_ *and o* = *o*_min_.

### S4.3 Capacity Upper Bound under Homogeneous Plasticity

The capacity is defined as the maximum number of patterns that can be stored while maintaining AUC ≥ 1 − *ϵ*, equivalently 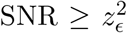 where *z*_*ϵ*_ ≡ Φ^−1^(1 − *ϵ*). Because SNR under homogeneous plasticity is upper bounded by the SNR of a direct readout across all *P* values, it follows that the capacity of any ensemble trained under homogeneous plasticity is upper bounded by the capacity of the direct readout architecture. To obtain this capacity upper bound, we may simply set 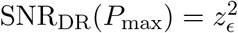 and solve for *P*_max_. In this case, we find:

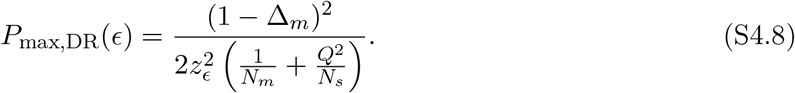

## S5 Analytical SNR and Capacity Estimates for Separable Decoders

### S5.1 Preliminaries: PC Input Statistics for Fully-Connected Networks

In this section, we consider the one-shot heterogeneous plasticity learning rule:

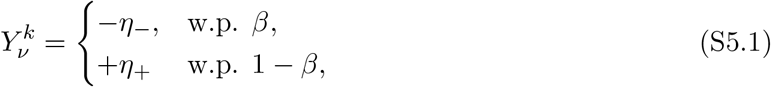

and specialize to fully-connected PCs with *N*_*d*_ = *N*_*m*_. A random novel input then gives ***g***_*n*_ | **Y** ~ *N*(**0, Σ**_0_), while a random learned input from cluster *µ* gives ***g***_*µ*_ | **Y** ~ *N*(***m***_*µ*_, **Σ**_*µ*_) with 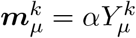. We assume for the learned-input condition that *µ* ~ Unif(1, …, *P*). The covariance matrices are

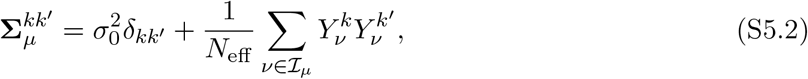

where ℐ_*µ*_ = {1, …, *P*} \ {*µ*} for *µ ≥* 1 and, by convention, ℐ_0_ = {1, …, *P*}. Thus |ℐ_*µ*_| = *P −* 1 for learned inputs and |ℐ_0_| = *P* for novel inputs. For brevity, we will write *P*_*n*_ = *P* and *P*_*ℓ*_ = *P −* 1. To allow for a unified treatment of the novel and learned input cases, we will often represent input class with an index *c* ∈ {*n, ℓ*}. We also define *c*_*µ*_ = *n* for *µ* = 0 and *c*_*µ*_ = *ℓ* for all *µ* ≥ 1.

The entries 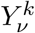 are i.i.d. across patterns *ν* and cells *k*. We denote the single-site moments by 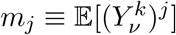, write 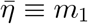, and write 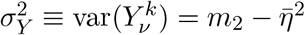.

The matrices **Σ**_*µ*_ depend on the random plasticity schedule **Y**. We decompose them into their **Y**-average and a mean-zero fluctuation matrix,

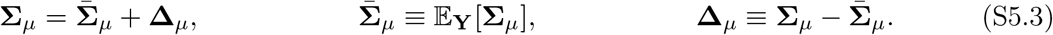

Because the entries of **Y** are i.i.d., 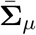 has one diagonal value and one off-diagonal value,

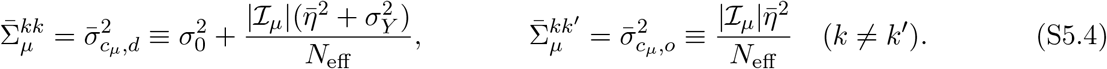

The fluctuations in the components of **Δ**_*µ*_ have the following covariance statistics for individual cells *k* and pairs *k*≠ *k*^*′*^:

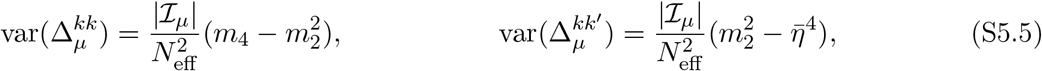

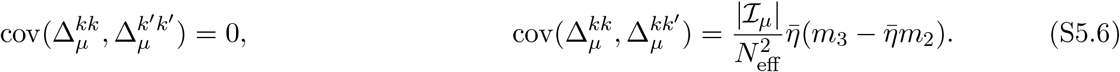

#### S5.1.1 Balanced Plasticity Special Case

For the quadratic and pause-based decoders, we will specialize to the “balanced plasticity” special case, where *η*_−_ = (1 − *β*)Λ and *η*_+_ = *β*Λ so that 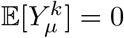 and *η*_−_ + *η*_+_ = Λ. All relevant single-site moments are then written

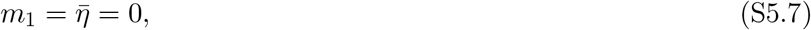

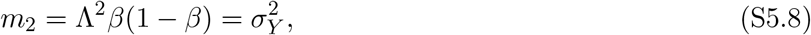

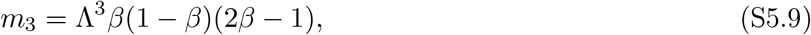

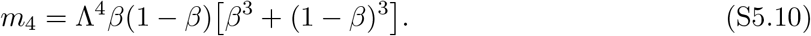

We define the variance of the squared signal as

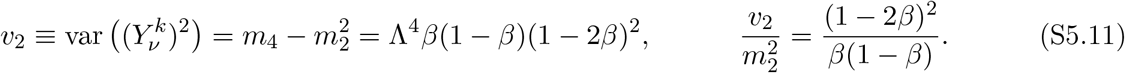

For input conditions *c* ∈ {*n, ℓ*}, the components of 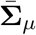 reduce to:

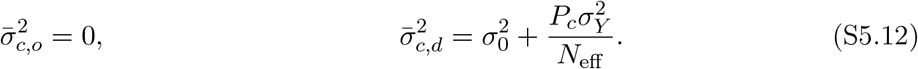

We also define the diagonal crosstalk fractions

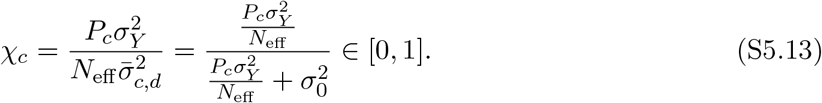

The diagonal and off-diagonal covariance fluctuations decouple, giving

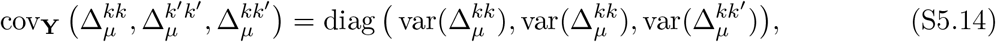

with

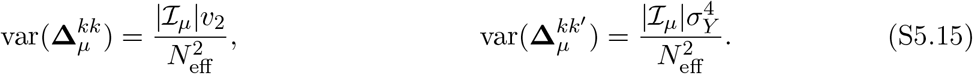

We will also have, for *µ*≠ *ν* and *µ, ν* ≥ 1,

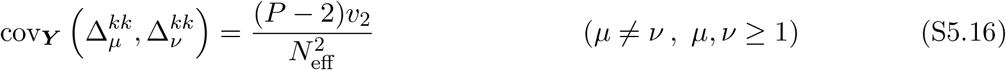

### S5.2 SNR approximation for general separable decoders

From the population response ***g*** ∈ ℝ^*k*^, a scalar readout Ω(***g***) must discriminate a novel input (*c* = *n*) from a learned input (*c* = *ℓ*). For a fixed plasticity schedule **Y**, define

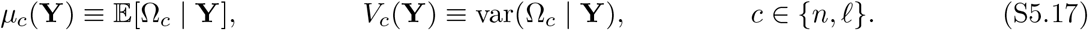

When the conditional readout distributions are close to Gaussian, we may approximate

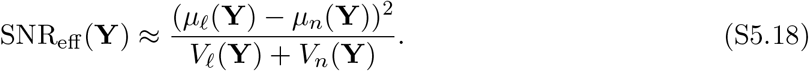

For large networks, we expect the mean and variance statistics to concentrate over the randomly drawn plasticity schedule ***Y***, so that we may approximate *µ*_*c*_(**Y**) and *V*_*c*_(**Y**) by their averages:

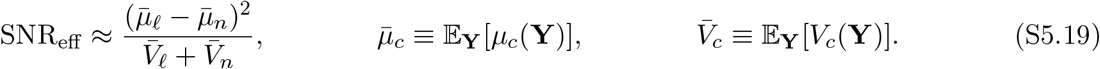

We do not rigorously justify these approximations here, rather appealing to the empirical match between the capacity curves obtained from these approximations and the results of Monte Carlo simulations with explicitly realized ***Y*** matrices (see main text Fig. 3). We also verify the capacity scaling relationships derived here with rigorous (but loose) tail bounds that account for the trial-to-trial fluctuations in ***Y*** in Sec. S12.

Here, we will derive approximate SNR expressions for several decoders of “separable” form,

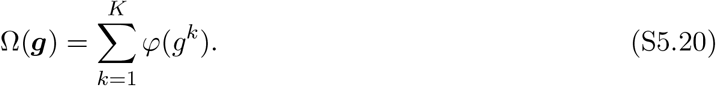

For any scalar nonlinearity *φ*, we define the following mean and variance functionals

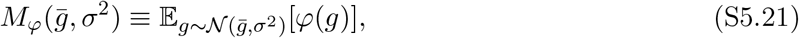

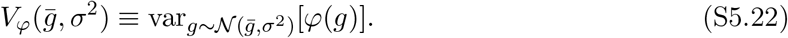

For 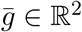 and **Σ** ∈ ℝ^2*×*2^, we define the covariance functional

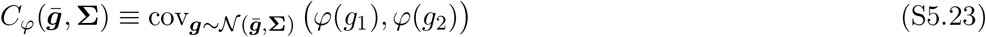

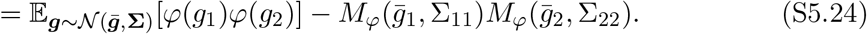

For a pair of cells *k*≠ *k*^*′*^, we write

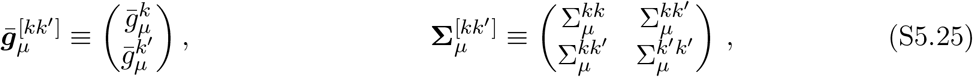

and use the abbreviations

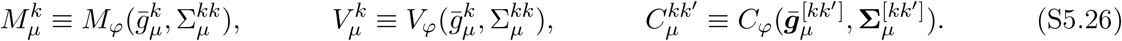

By convention, we write ***m***_0_ = **0** when *µ* = 0, corresponding to the novel input condition. Because 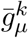 and 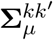 both depend on ***Y***, we may also write the above quantities as functions of ***Y*** where appropriate. Because the distribution of ***Y*** is exchangeable over the PCs *k*, the plasticity-averaged readout moments may be simplified as:

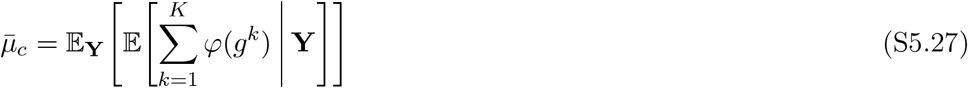

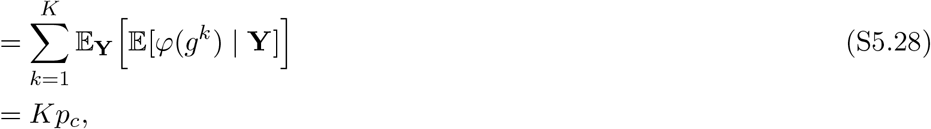

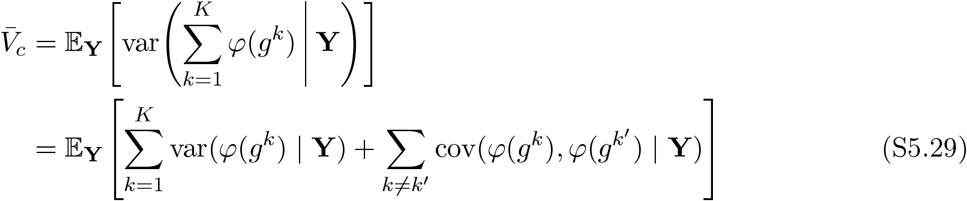

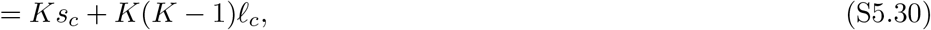

where

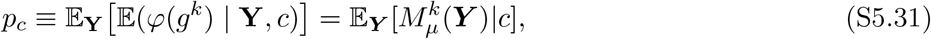

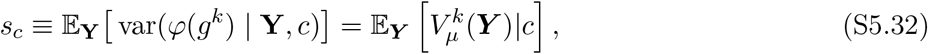

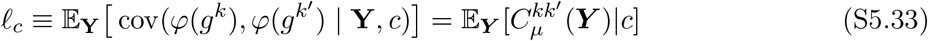

and where it is understood that when *c* = *n* we take *µ* = 0 and when *c* = *ℓ* we may take any *µ* ≥ 1. For the novel condition, there is no learned-pattern index to average over, and

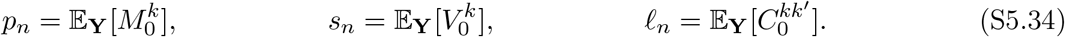

For learned inputs, *µ* ~ Unif(1, …, *P*). The mean is

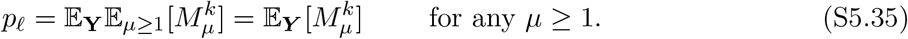

Here, we may remove the expectation over *µ* ≥ 1 because the distribution of ***Y*** is exchangeable over the cluster index *µ* ∈ {1, …, *P*}. Similarly, the law of total variance gives

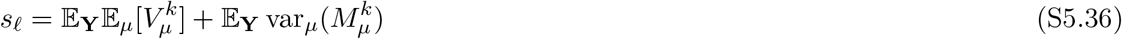

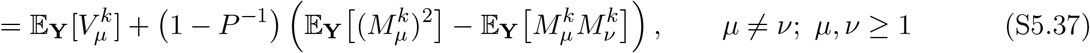

and the law of total covariance gives

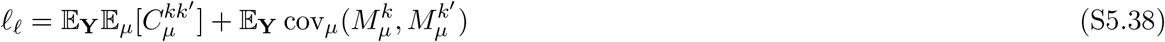

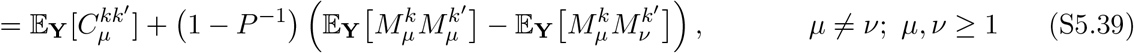

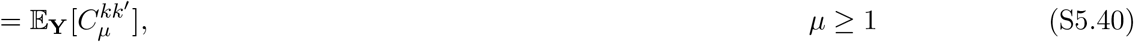

where in the last line, we have used the fact that 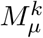 and 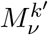 are independent over draws of ***Y*** when *k*≠ *k*^*′*^ for any *µ, ν* (even when *µ* = *ν*), so that 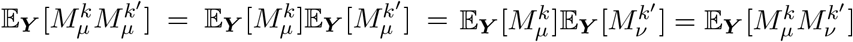. Plugging back into eq. S5.19, we obtain the approximation

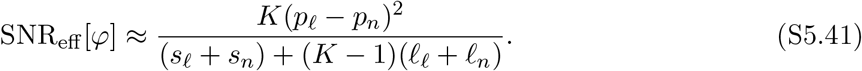

To determine SNR for nonlinearity *φ*, we use the following procedure: first, calculate the expectation and variance functionals *M*_*φ*_, *V*_*φ*_, and *C*_*φ*_. These may then be used to determine *p*_*c*_, *s*_*c*_, and *ℓ*_*c*_ for *c* ∈ {*n, ℓ*}. Finally plug into eq. S5.41 to obtain the approximate SNR. In the sections that follow, we apply this recipe in the special cases of linear (*φ*(*g*) = *g*), quadratic (*φ*(*g*) = (*g* − *κ*)^2^), and pause-based (*φ*(*g*) = Θ(*κ* − *g*)) decoding.

### S5.3 Linear decoder *φ*(*g*) = *g*

For the linear decoder, *φ*(*g*) = *g*, the Gaussian functionals are

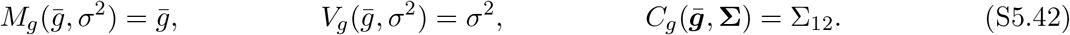

Thus the covariance fluctuations **Δ**_*µ*_ enter only through their means. For the novel condition,

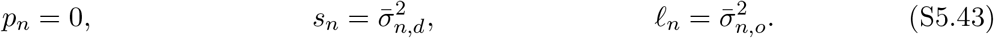

For the learned condition, 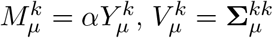, and 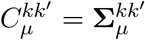. Equations (S5.35)–(S5.40) give

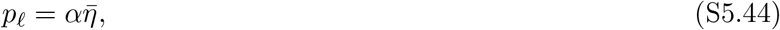

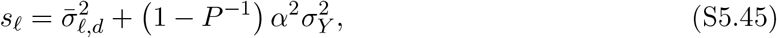

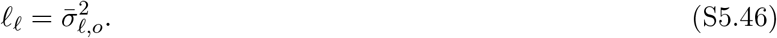

The expected mean gap is

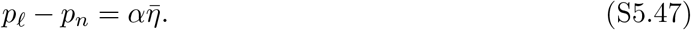

Substituting (S5.43)–(S5.47) into (S5.41) and taking *P* ≫ 1, so that *P* − 1 ≈ *P*, gives

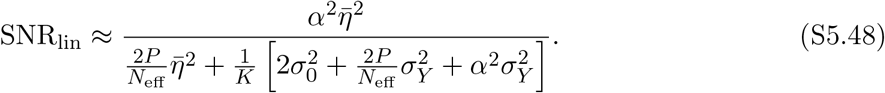

Thus, we see that in the balanced case, 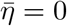 and the linear readout achieves zero discriminability. It also follows easily from eq. S5.48 that under linear decoding, the optimal value of *β* is that which maximizes 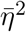 and minimizes 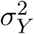, which is achieved by setting *β* = 1 in cases where *η*_−_ *> η*_+_ or *β* = 0 when *η*_−_ *< η*_+_. Defining *η*_⋆_ = max(*η*_−_, *η*_+_), we get the *β*-optimized SNR under linear decoding:

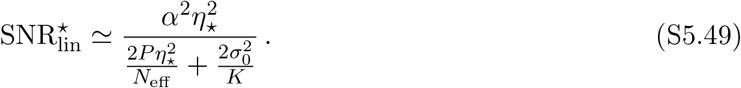

This is equivalent to the SNR achieved by a single PC trained under homogeneous plasticity with weight update strength *η*_⋆_ and intrinsic noise of variance 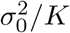. As a result, a linear network can never outperform the direct-readout architecture:

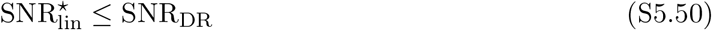

with equivalence when *σ*_0_ = 0 or *K* → ∞ (note that equivalence with *K <* ∞ is possible only because *N*_*d*_ = *N*_*m*_).

### S5.4 Quadratic decoder with balanced plasticity

We take the centered quadratic readout *φ*(*g*) = (*g* − *κ*)^2^, where *κ* is a user-defined shift. Using Isserlis’ theorem for Gaussian moments, one easily obtains

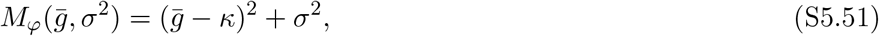

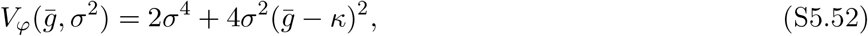

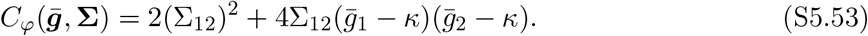

We work in the balanced regime, with 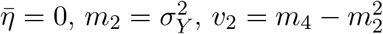, and 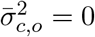. Define

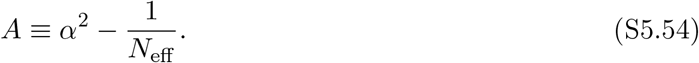

#### Novel moments

For ***m***_0_ = 0, we proceed by substituting S5.51-S5.53 into S5.34, S5.35, S5.37, S5.40 and evaluating the expectations over ***Y*** using the balanced-plasticity covariances of the elements of **Δ**_0_:

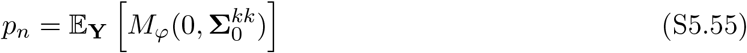

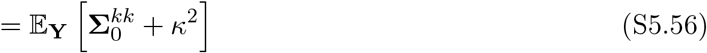

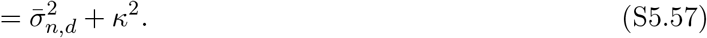

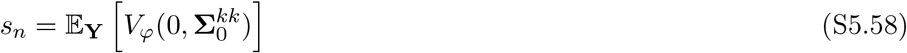

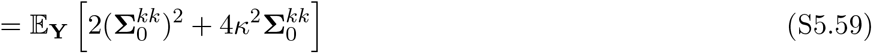

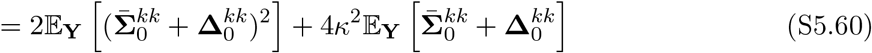

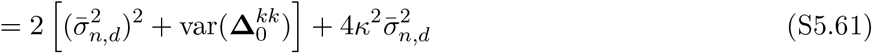

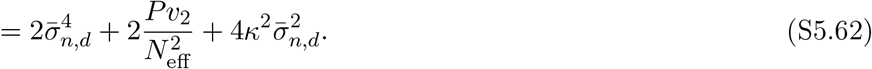

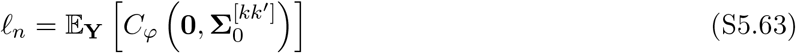

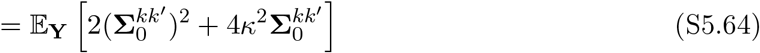

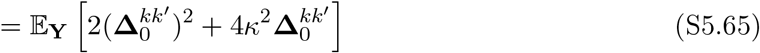

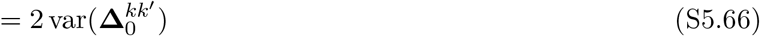

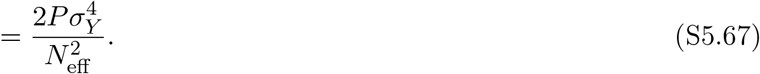

#### Learned moments

For a learned input, *µ* ~ Unif(1, …, *P*). The cued pattern is excluded from the background covariance sum, so **Σ**_*µ*_ is independent of 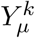 and 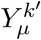. Applying (S5.35) gives

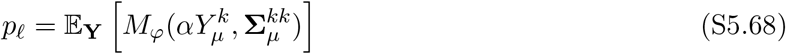

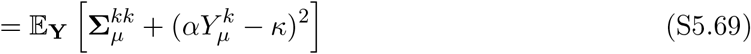

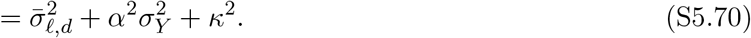

Thus

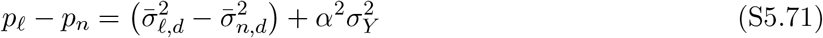

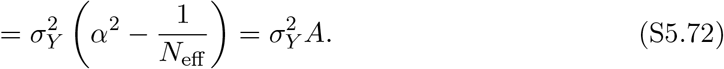

For the single-cell variance, (S5.37), the first term is

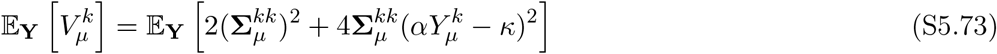

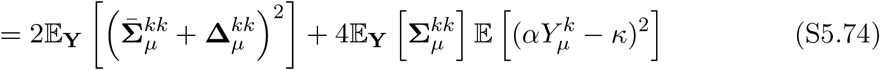

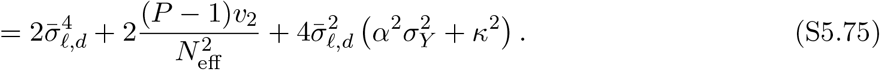

For the second term in (S5.37), write

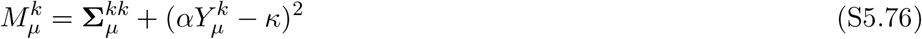

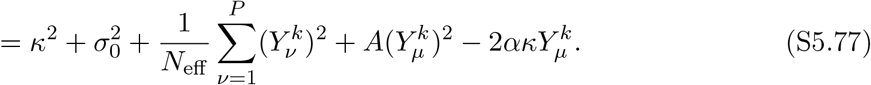

The first three terms are independent of *µ* and drop out of the variance over learned patterns. Therefore

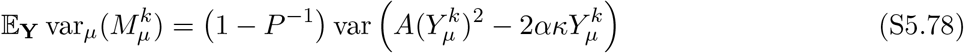

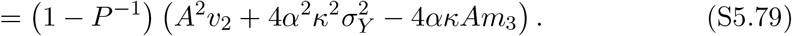

Combining the two terms in (S5.37),

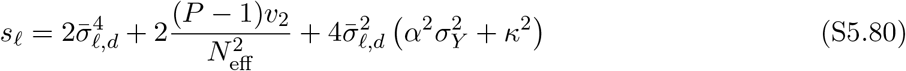

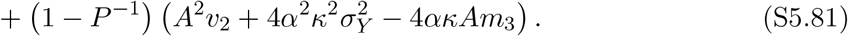

For the learned pair covariance, (S5.40) gives

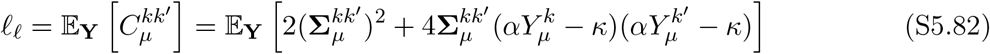

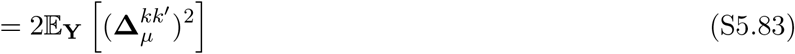

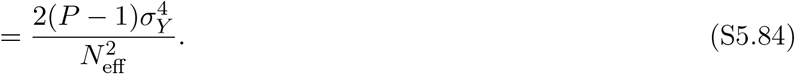

#### Quadratic SNR

We next combine the above terms according to eq. S5.19, taking *N*_eff_ ≫ 1 so that *A* ≃ *α*^2^ and *P* ≫ 1 so that *P* − 1 ≈ *P*, giving

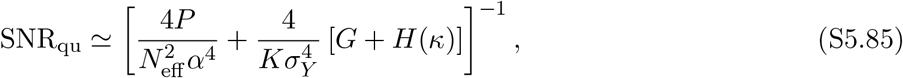

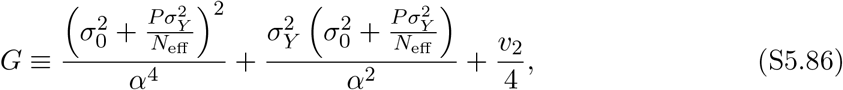

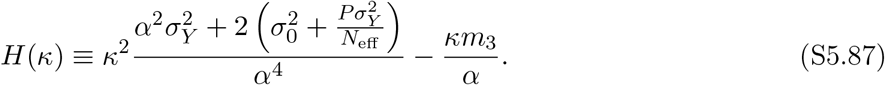

#### Optimization over *β* and *κ*

To maximize SNR_qu_ over *κ* at fixed *β*, we set *H*^*′*^(*κ*) = 0, giving optimal value

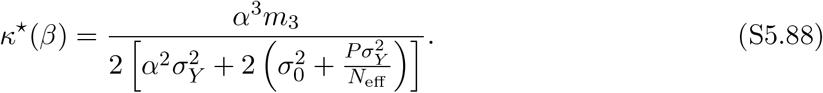

Using 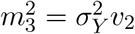 under balanced plasticity, the inverse SNR after optimizing over *κ* becomes

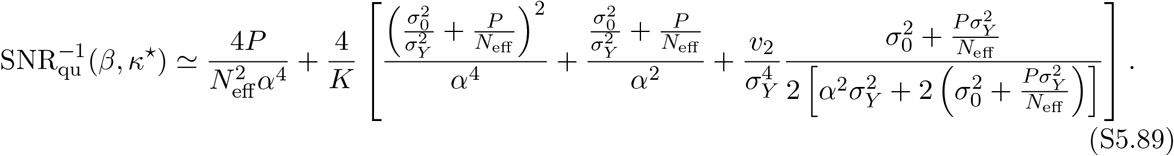

Recall that

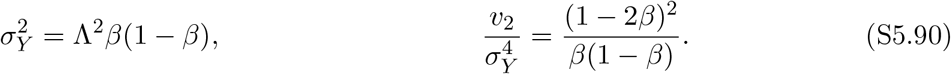

The first two terms inside the brackets in (S5.89) are decreasing functions of 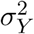, and hence minimized when *β* = 1*/*2. The third term in the brackets in (S5.89) is nonnegative and proportional to *v*_2_, which vanishes at *β* = 1*/*2. Therefore all *β*-dependent terms are minimized at *β*^⋆^ = 1*/*2. At this value, *m*_3_ = 0, and so (S5.88) gives *κ*^⋆^ = 0. Thus the joint optimum is

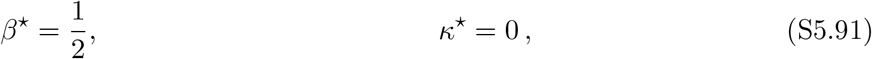

so that *η*_+_ = *η*_−_ = *η*, where we define *η* = Λ*/*2. At 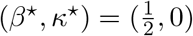, (S5.85) then reduces to

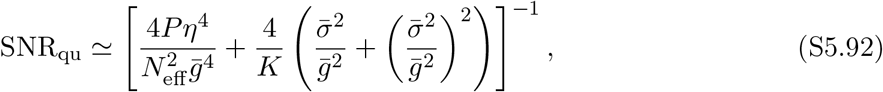

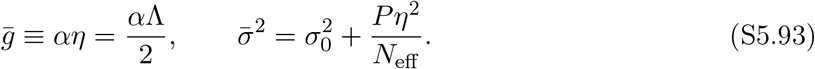

#### Capacity Scaling

To determine capacity with the quadratic decoder, we set 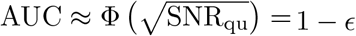 and solve for *P*_max_. Plugging in eq. S5.85, we obtain *P*_max_ in closed form as the positive root of the quadratic equation 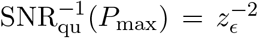, where *z*_*ϵ*_ = Φ^−1^(1 − *ϵ*). At optimal parameters *β* = 1*/*2, *κ* = 0, this may be written explicitly as the positive root of

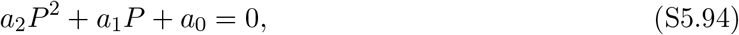

with coefficients

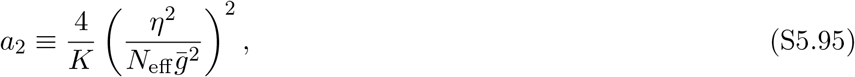

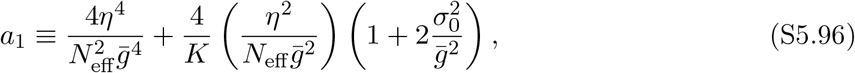

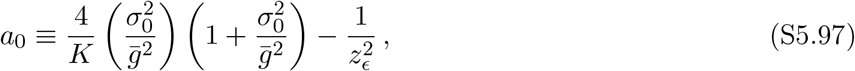

or *P*_max_ = 0 when no positive root exists. We obtain a more transparent capacity approximation at (*β, κ*) = (1*/*2, 0) by writing

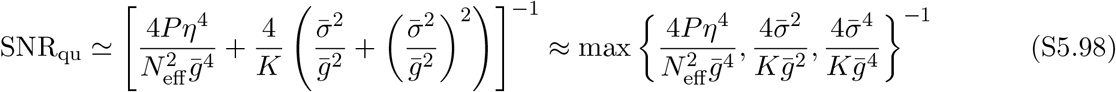

Setting each entry in the last expression equal to 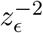 and solving each for *P*, we obtain the approximate capacity:

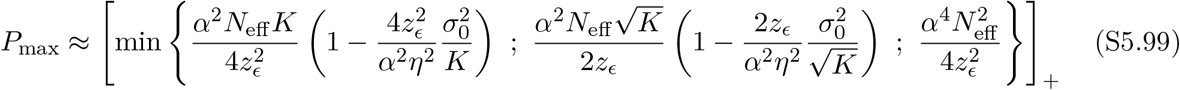

In particular, in the biologically relevant regime where 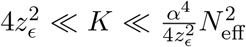 the center branch will dominate, so that 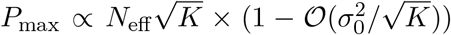. Thus, we see that the network is highly robust to intrinsic noise in this regime, providing a multiplicative effect on capacity which vanishes as 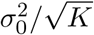. The scaling behavior of *N*_eff_ depends strongly on the sparsity *f* of the GrC population activity. In main text fig. 3, we examine the sparse-coding regime where *N*_*d*_ ≪ *N*_*s*_*/Q*^2^, so that *N*_eff_ ~ *N*_*d*_. Then, provided 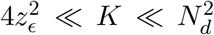, we have 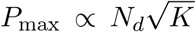. Alternatively, we may consider the scaling regime where *N*_*d*_ ≫ *N*_*s*_*/Q*^2^, so that *N*_eff_ ~ *N*_*s*_*/Q*^2^. In this case, the stimulus dimensionality *N*_*s*_ becomes the bottleneck to capacity:

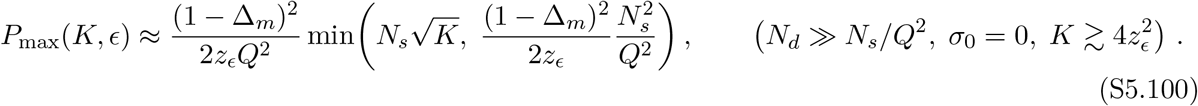

We explore this “dense-coding” capacity scaling regime in fig. S5.

### S5.5 Pause decoder with balanced plasticity

We now specialize to the pause-based decoder *φ*(*g*) = Θ[*κ* − *g*], where Θ[·] represents the Heaviside step function. Here, we interpret *κ* as an input threshold below which a PC will undergo a “pause”. The DCN output Ω(***g***) = _k_ Θ[*κ* − *g*^*k*^] is then dis-inhibited in proportion to the number of pausing PC inputs. We continue to use the balanced two-point plasticity distribution of Section S5.1.1 and write *α* ≡ 1 − Δ_*m*_, so a learned input from cluster *µ* has signal mean 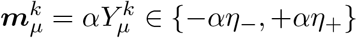.

For *φ*(*g*) = Θ(*κ* − *g*), define

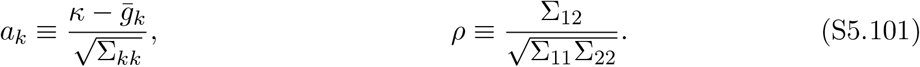

**Figure S5:**
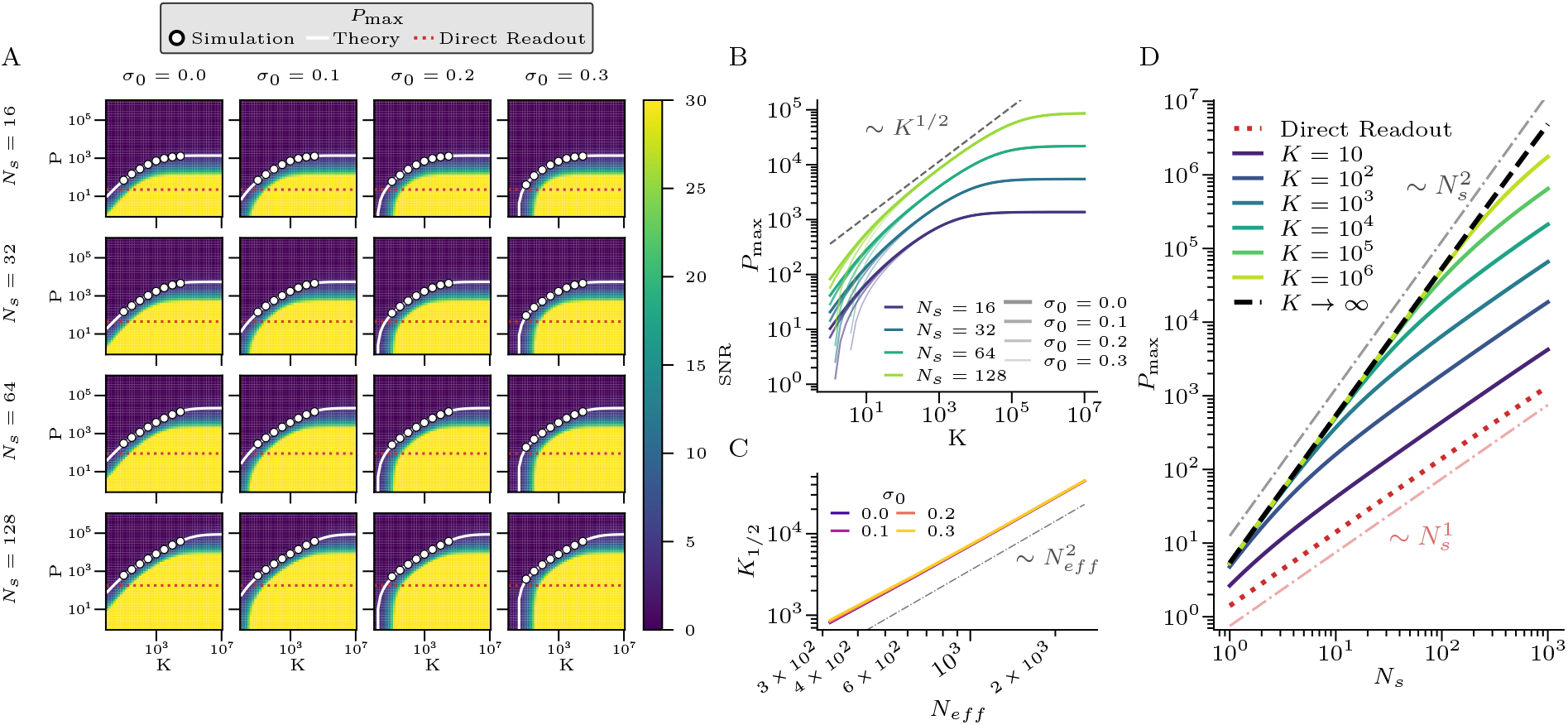
Capacity scaling under heterogeneous plasticity for low-dimensional inputs or dense GrC coding. (A) Heatmaps of SNR in the (*K, P*) plane under balanced heterogeneous plasticity (*β* = 0.5, *η*_+_ = *η*_−_ = 1) with a quadratic decoder *φ*(*g*) = *g*^2^. Grid panels show combinations of stimulus dimensionality *N*_*s*_ ∈ {16, 32, 64, 128} and intrinsic noise *σ*_0_ ∈ {0, 0.1, 0.2, 0.3}. Theoretical capacity curves (white) agree well with numerical experiments (markers). Red dashed lines show the capacity of the direct readout architecture. (B) Capacity vs. ensemble size *K*, aggregated across *N*_*s*_ and *σ*_0_ values. After an initial linear regime, capacity grows as 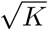 until saturating at an *N*_*s*_-dependent plateau. (C) The ensemble size *K*_1/2_ required to achieve half of the plateau capacity scales as 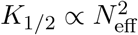. Parameters as in fig. 2 unless otherwise indicated. (D) Capacity vs. effective input dimensionality *N*_eff_ ∝ *N*_*s*_ for fixed ensemble sizes *K*. As *K* increases, the capacity curve transitions from linear to quadratic scaling with *N*_eff_. In the limit *K* → ∞, the capacity scales as 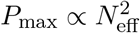, a qualitative improvement over the linear scaling *P*_max_ ∝ *N*_eff_ of the direct readout.

The decoder-specific Gaussian objects are

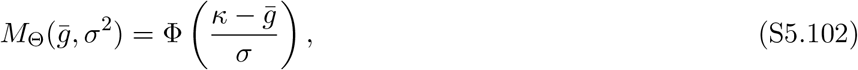

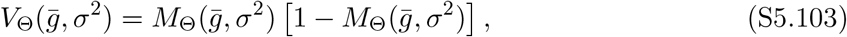

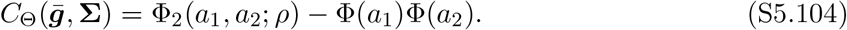

Here Φ_2_(·, ·; *ρ*) is the standard bivariate normal CDF with correlation *ρ*. Taylor expanding the bivariate normal density *ϕ*_2_(*x, y*; *ρ*) about *ρ* = 0 and integrating term by term over *x* ≤ *a*_1_ and *y* ≤ *a*_2_, we obtain:

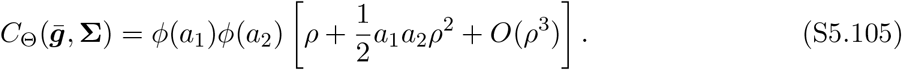

#### S5.5.1 Averages over covariance fluctuations

The dependence of the expressions for *p*_*c*_, *s*_*c*_, *ℓ*_*c*_ for *c* ∈ {*n, ℓ*} on the plasticity schedules ***Y*** can be decomposed into its dependence on the plasticity signals 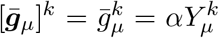 when *µ* ≥ 1 (of 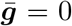 when *µ* = 0 for the novel-input case) and the covariance fluctuations 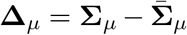. For example, because 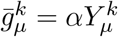 is independent from 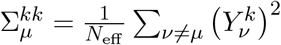 we may write

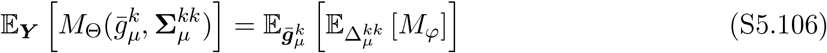

In this section, we evaluate the relevant expectations over the covariance fluctuations **Δ**_*µ*_, leaving the mean signal in general form 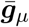. For the novel-input case where *µ* = 0, this encompasses the full expectation over ***Y***. For learned inputs, it will be necessary to evaluate the expectations over the mean signal as well (see sec. S5.5.3). To make these expectations tractable, we will employ the delta method:

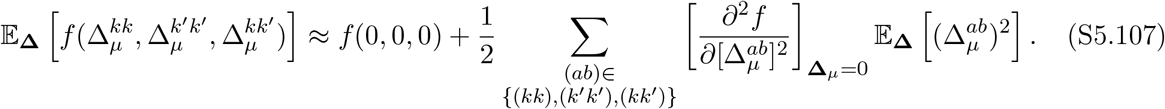

Where we have used that under balanced plasticity, 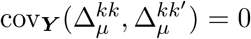. In the balanced regime, 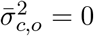, so 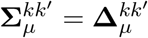 for *k*≠ *k*^*′*^. For the diagonal fluctuations, define

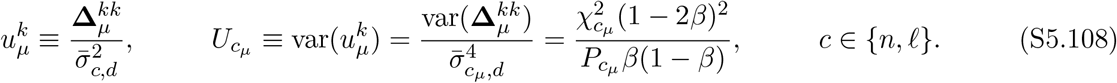

It is also useful to note that

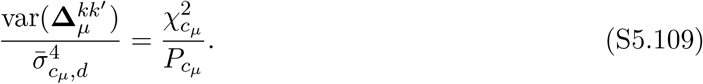

Expanding in 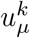, we have

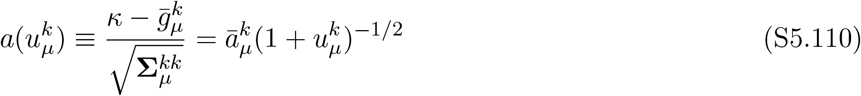

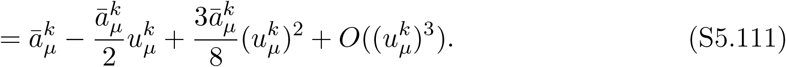

where we have defined

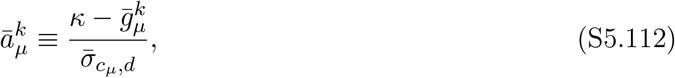

We also define

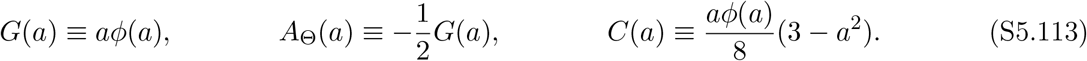

Then

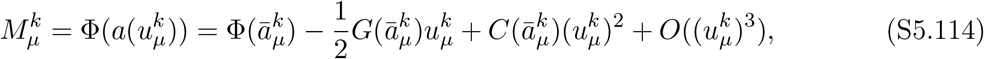

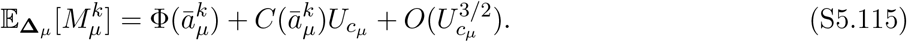

Similarly, for the Bernoulli variance,

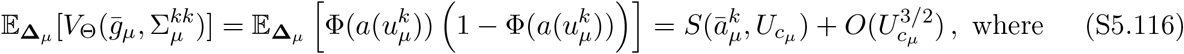

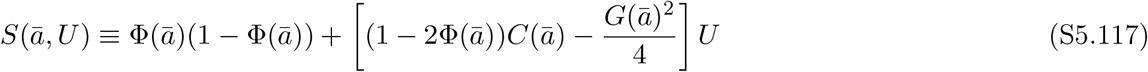

To simplify 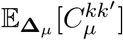 for balanced plasticity, we write

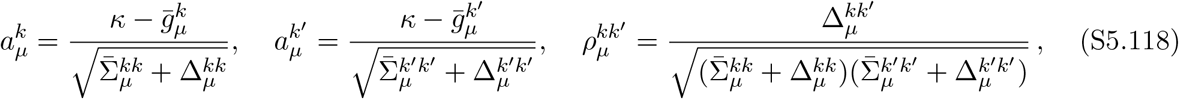

plug into eq. S5.105, and expand to second order in the 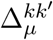 giving

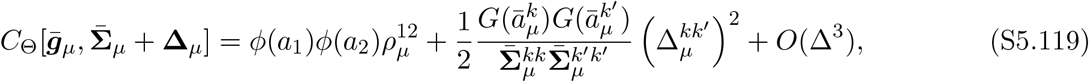

where we have defined 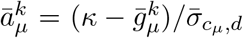. Averaging over the fluctuations the linear term drops out because 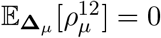. We then obtain

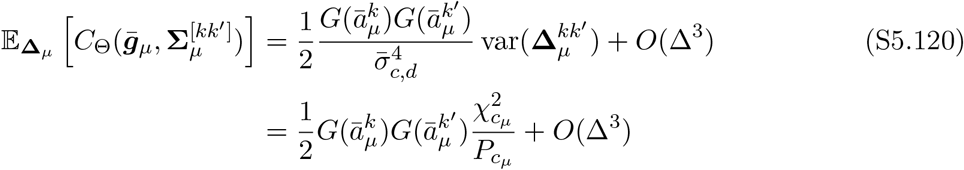

To calculate *s*_*ℓ*_, we will need additional moments of the 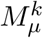:

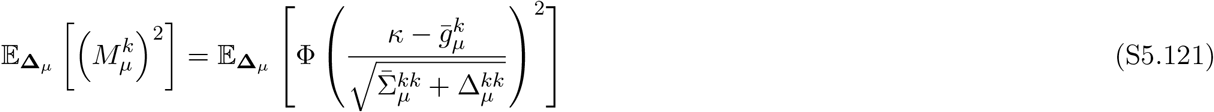

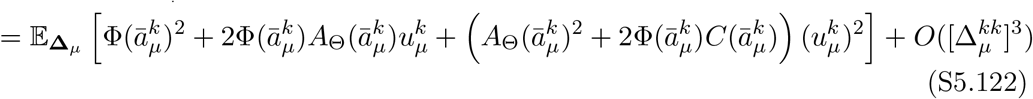

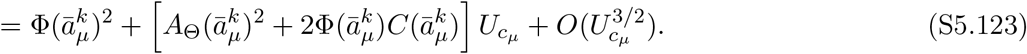

And for *µ*≠ *ν*, define

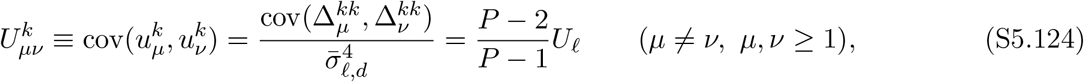

where the last equality uses (S5.16). Then

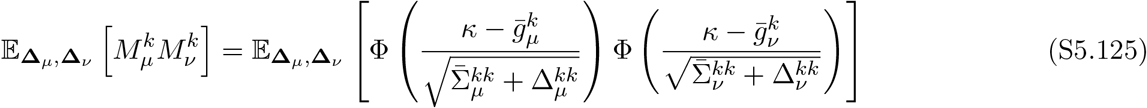

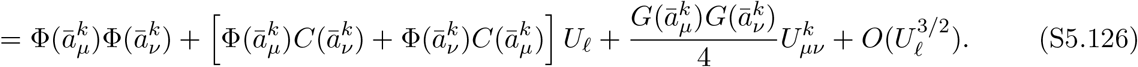

#### S5.5.2 Novel-input moments

For novel inputs, 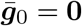, so that 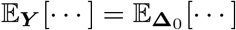. The values of *p*_*n*_, *s*_*n*_, and *ℓ*_*n*_ therefore follow directly from the above expectations. Define

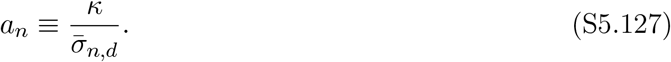

Using (S5.115), we have

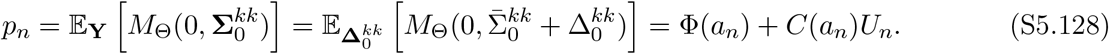

Using (S5.116), the single-cell variance is

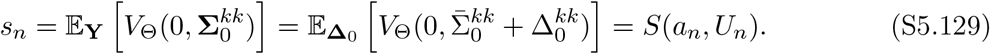

Using (S5.120), the pair covariance is:

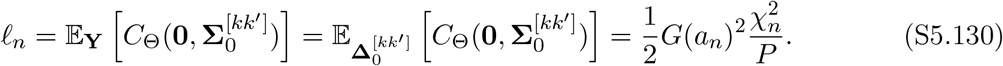

#### S5.5.3 Learned-input moments

For learned inputs we must also account for fluctuations over the mean signals 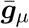. Define the two branch thresholds

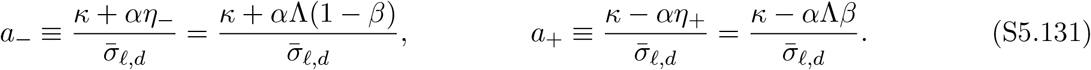

The branch *a*_−_ corresponds to 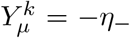, which occurs with probability *β*, while *a*_+_ corresponds to 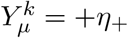, which occurs with probability 1 − *β*. In this section, we use the shorthand

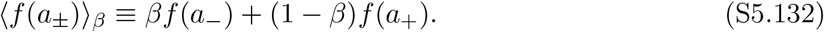

Applying (S5.115),

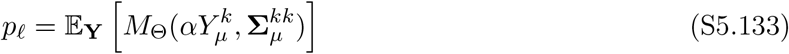

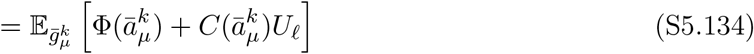

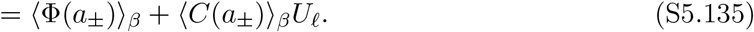

Next, we calculate the terms of *s*_*ℓ*_ as represented in (S5.37). Using (S5.116), the within-pattern contribution is

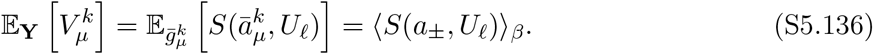

Using (S5.123), the single-pattern second moment is

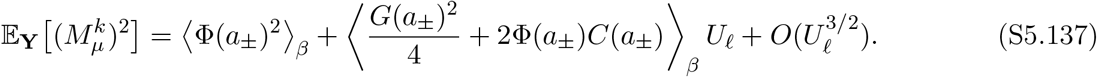

For the cross-pattern moment, use (S5.126). Defining 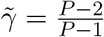, we obtain

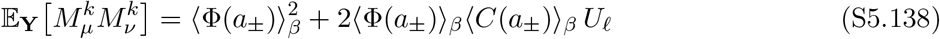

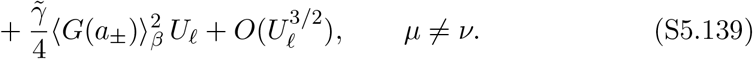

We can then simplify,

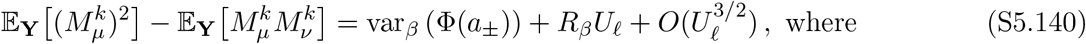

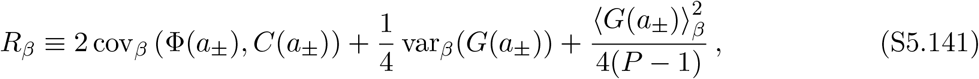

where we are using the shorthands

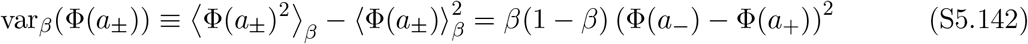

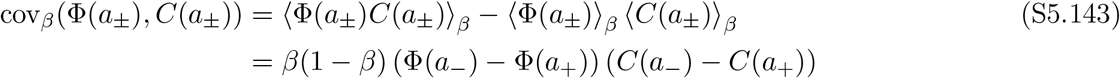

Combining these terms with (S5.37) gives

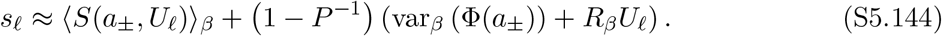

Finally, to evaluate *ℓ*_*ℓ*_ we use (S5.40) and (S5.120), giving

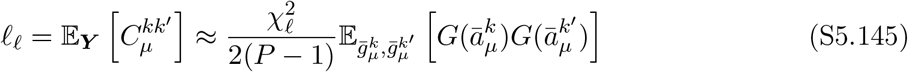

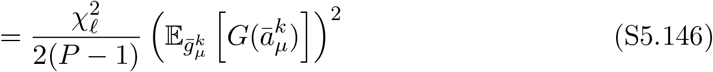

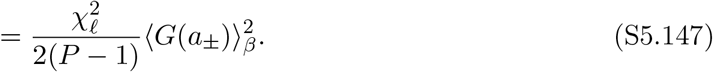

The full SNR is then obtained by substituting the expressions for *p*_*c*_, *s*_*c*_, and *ℓ*_*c*_ for *c* ∈ {*n, ℓ*} into the SNR approximation in eq. S5.19.

#### S5.5.4 Capacity under Pause-Based Decoding with Sparse LTD

Next, we specialize to the noiseless case (*σ*_0_ = 0) and argue that the SNR formula derived above permits capacity to scale as

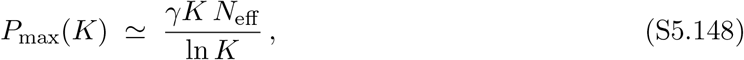

for some constant *γ*, provided *N*_eff_ ≫ log(*K*). To achieve this, we set

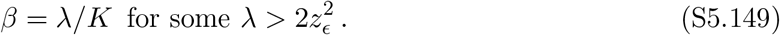

In the noiseless case, we will have *χ*_*c*_ = 1 for *c* ∈ {*n, ℓ*}, so that 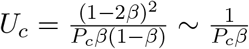. Substituting *P* ~ *KN*_eff_ */* log(*K*), we have *U*_*c*_ ~ log(*K*)*/N*_eff_, which is small by hypothesis. Assuming as well that *P, K, N*_eff_ ≫ 1, we may also simplify the components of the SNR formula as follows. For novel inputs:

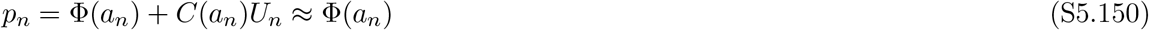

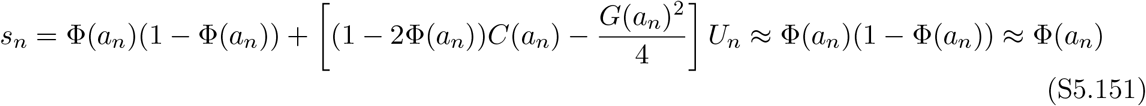

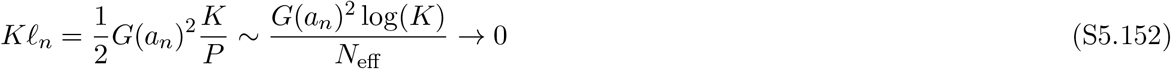

For learned inputs, we will similarly have:

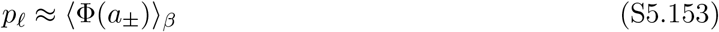

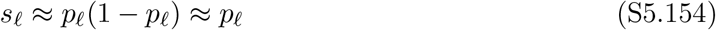

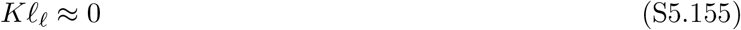

where in the second line we have used that ⟨Φ(1 − Φ)⟩_*β*_ + var_*β*_(Φ) = ⟨Φ⟩_*β*_(1 − ⟨Φ⟩_*β*_). Combining these, we obtain the following approximation of SNR in the regime *P* ~ *KN*_eff_ */* log(*K*):

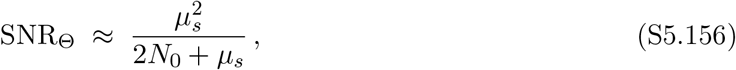

where we have defined the baseline and excess pause counts

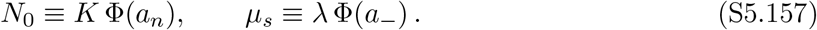

It then suffices to show that for the conjectured capacity scaling, there exists a choice of the threshold *κ* which keeps SNR ~ *O*(1). A sub-optimal but analytically convenient choice is 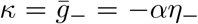, so that *a*_−_ = 0 and *µ*_*s*_ = *λ/*2. We will then have 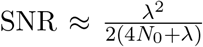. For fixed 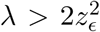, we may achieve 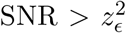 as long as *N*_0_ can be made very small. With 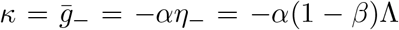 we have *a*_−_ = 0 by construction. For the baseline, recall 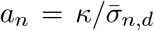 with 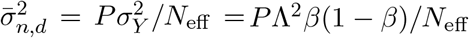, so

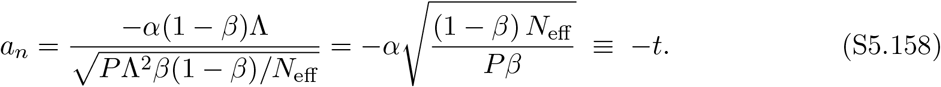

Substituting the conjectured scaling *P* = *γKN*_eff_ */* ln *K* and *β* = *λ/K* gives *Pβ* = *γλN*_eff_ */* ln *K*, hence

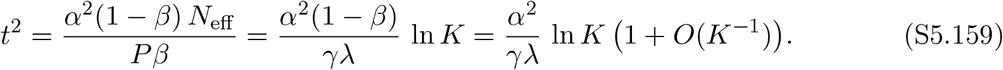

Thus the threshold sits a distance 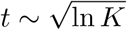 into the lower tail, and *N*_0_ is governed by the Gaussian tail at this depth. Using the following expansion:

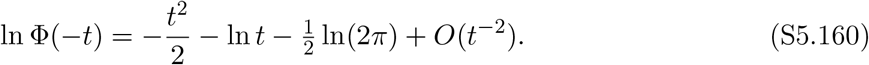

we then have

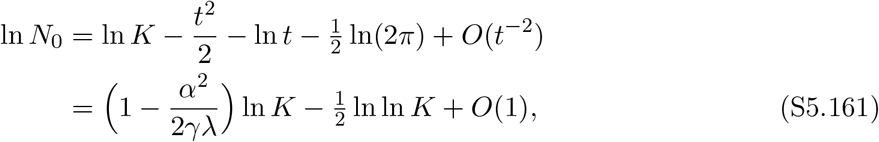

so that,

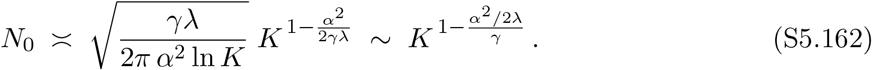

The *K*–exponent is 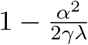, so *N* → 0 polynomially in *K* precisely when it is negative.

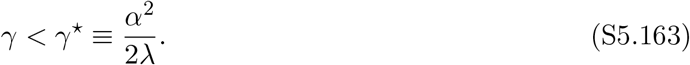

Then, for any *γ < γ*^⋆^, we have *N*_0_ → 0, so

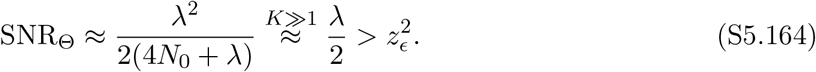

This shows consistency of the *P*_max_ ∝ *KN*_eff_ */* ln *K* scaling. If we write *K* = *c*_1_*w, N*_eff_ = *c*_2_*w* and scale the “width” parameter *w*, we obtain *P*_max_ ∝ *w*^2^*/* log(*w*).

## S6 Derivations of Approximate Optimal Decoders

In this section, we derive the optimal transformation from the learned Purkinje cell (PC) inputs ***g*** = (*g*^1^, …, *g*^*K*^) to the scalar output Ω(***g***) used for classification. We work directly from the class-conditional distributions of ***g*** derived in §S3 and §S3.4. Throughout, we write

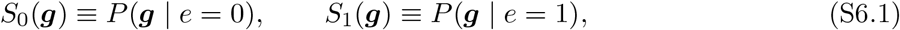

for the novel- and learned-class likelihoods, respectively.

By the Neyman–Pearson lemma [95], the decoder that maximizes the ROC curve is the log-likelihood ratio

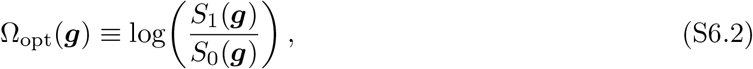

with classification performed by thresholding

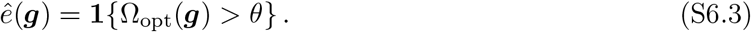

Any monotone transformation of Ω_opt_ yields the same ROC curve, so additive constants and positive rescalings may be absorbed into the threshold *θ*.

### S6.1 Optimal Decoding under Homogeneous Plasticity

Under homogeneous LTD, 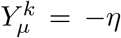 for all *µ, k*, and from §S3.3 the class-conditional PC input distributions are Gaussian with a shared covariance matrix:

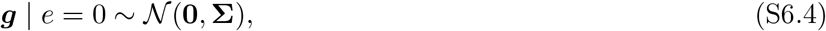

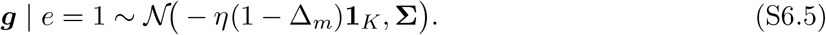

Therefore

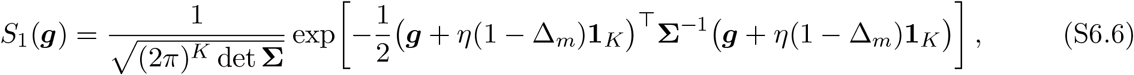

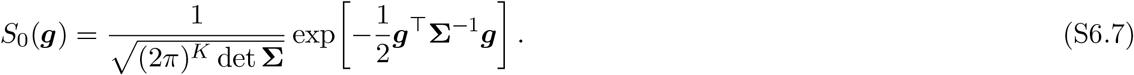

Taking the ratio gives

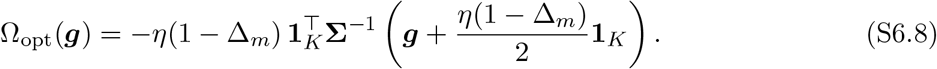

Under the homogeneous-overlap assumption, **Σ** has compound-symmetric form,

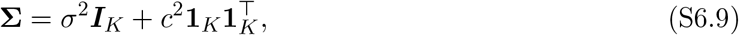

and therefore

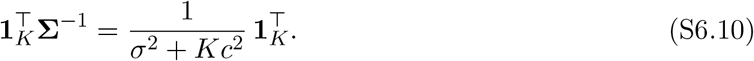

It follows that, up to an additive constant and positive rescaling,

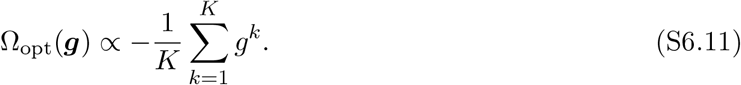

Thus under homogeneous plasticity the optimal decoder is simply the negative population average of the PC inputs.

### S6.2 Optimal Decoding under Heterogeneous Plasticity

Under heterogeneous plasticity, the class-conditional distribution of ***g*** is more complicated because the learned-class mean depends on the random plasticity outcomes 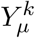. We work under the homogeneous-overlap assumption. For the purposes of deriving an optimal decoder, we will additionally adopt the expected-covariance approximation, replacing the random covariance **Σ**_(−µ)_ by its expectation over ***Y***,

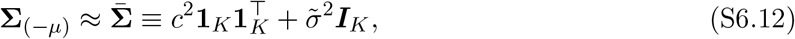

with

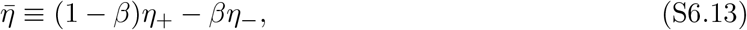

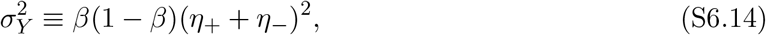

and

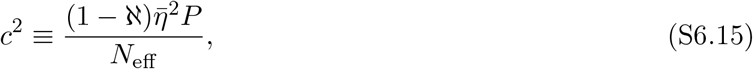

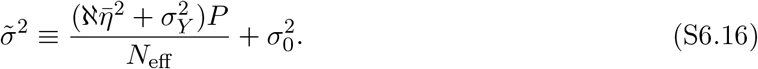

The corresponding pairwise correlation coefficient of the cross-talk fluctuations is

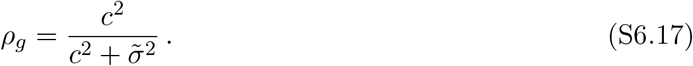

#### S6.2.1 Likelihood for *e* = 0

Under the approximation (S6.12), the novel-class distribution is simply

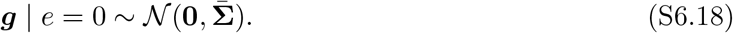

Its likelihood is

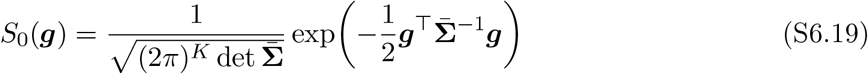

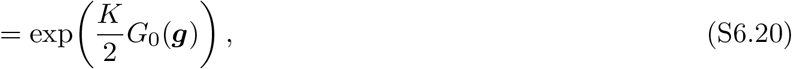

where

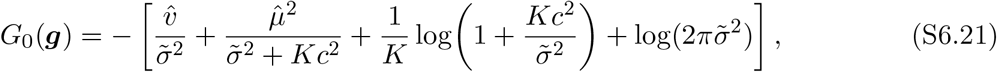

and

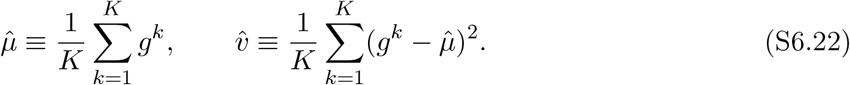

#### S6.2.2 Learned-Class Likelihood as a 2^*K*^-Component Gaussian Mixture

For *e* = 1, we still approximate the covariance by 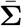, but the mean depends on the plasticity signals *Y* ^*k*^:

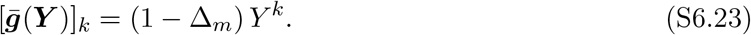

Thus

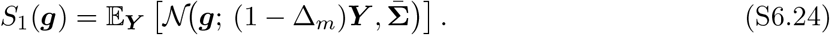

Equivalently, introducing a binary mask ***M*** ∈ {0, 1} ^*K*^ with *M* ^*k*^ = 1 indicating LTD and *M* ^*k*^ = 0 indicating LTP, so that

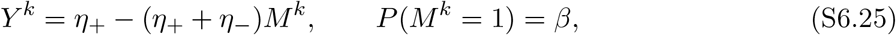

we may write

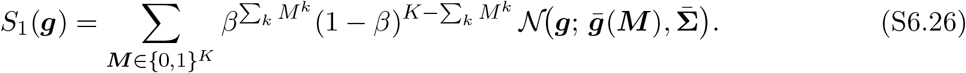

Direct evaluation therefore requires a sum over 2^*K*^ values for ***M***, which may become computationally intractable at large *K* values.

#### S6.2.3 Reduction to a One-Dimensional Integral

The equicorrelated Gaussian 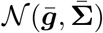 admits the standard one-factor decomposition

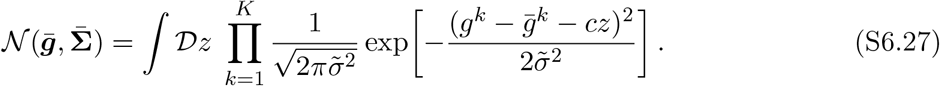

Therefore, conditioned on *z* and ***Y***, the coordinates factorize:

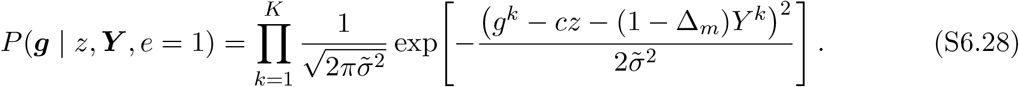

Averaging independently over the *Y* ^*k*^ then gives

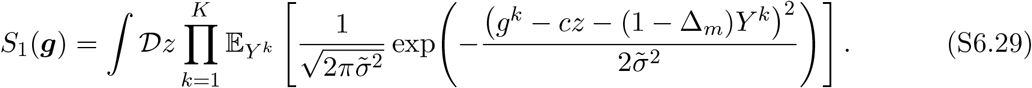

It is convenient to introduce

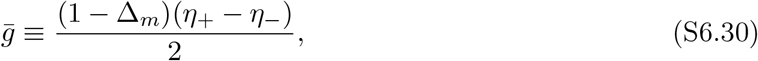

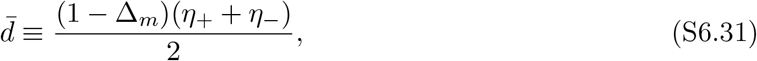

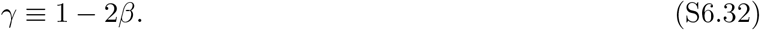

The two possible learned-class means are then 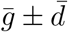. Writing

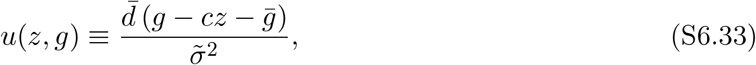

one finds

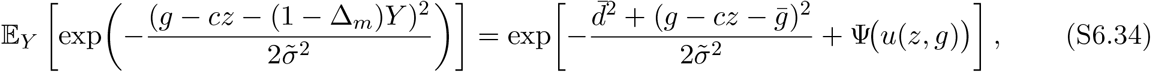

where

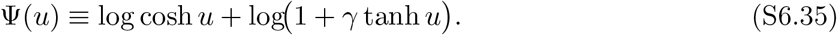

Thus the learned-class likelihood can be written as the one-dimensional integral

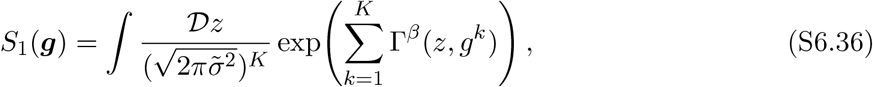

with

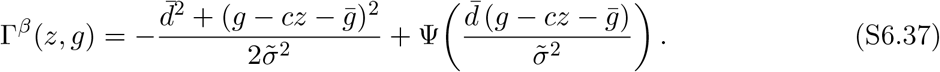

Equation (S6.36) is exact under the expected-covariance approximation and can be evaluated in *O*(*K*) time.

### S6.3 Saddle Approximation of the Optimal Decoder

We now derive a biologically interpretable approximation to Ω_opt_(***g***) by applying Laplace’s method to the one-dimensional integral (S6.36).

#### Novel class

For *e* = 0, the same latent-factor decomposition gives

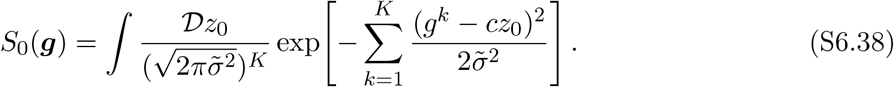

Because the integrand is Gaussian in *z*_0_, the saddle-point calculation is exact. The maximizing value is

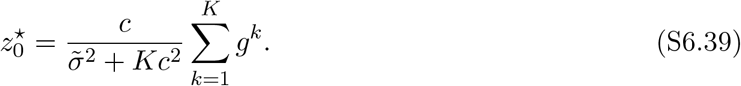

Defining 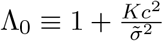, the corresponding log-likelihood may be written as

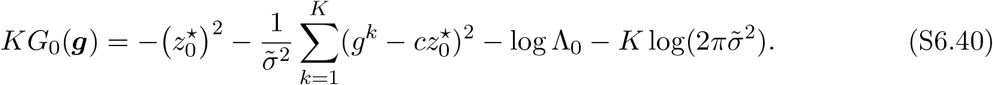

#### Learned class

For *e* = 1, write the log-integrand of (S6.36) as

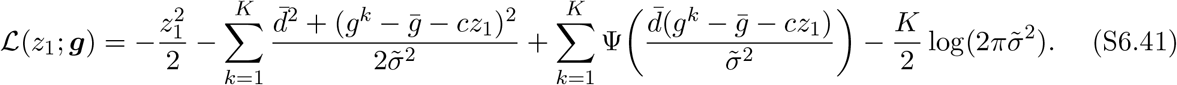

We also define

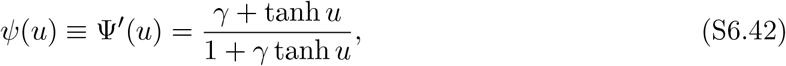

and

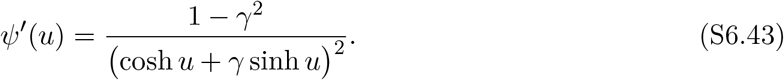

The saddle condition 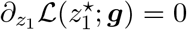 becomes

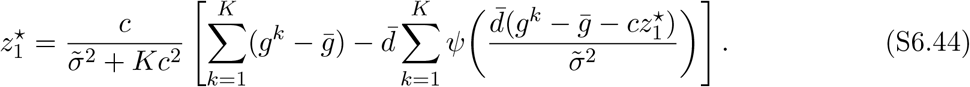

The curvature at the saddle is

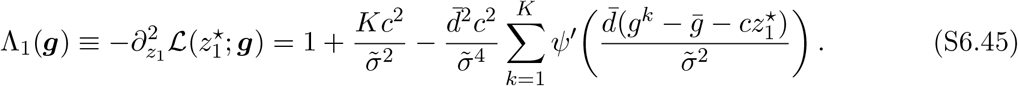

Laplace’s method then gives

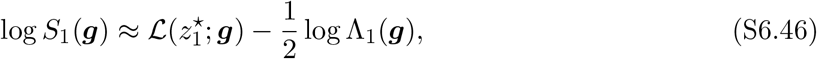

up to an additive constant independent of ***g***.

#### Approximate log-likelihood ratio

To simplify the result, define

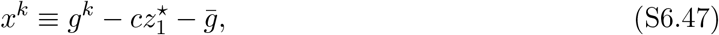

and

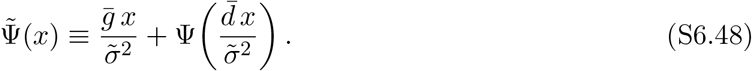

Subtracting the novel-class contribution and simplifying gives

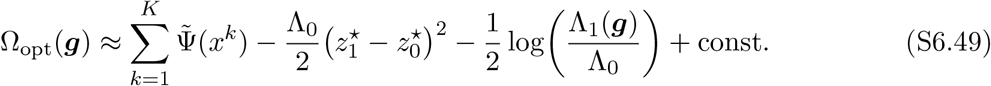

Using (S6.39) and (S6.44), one further finds

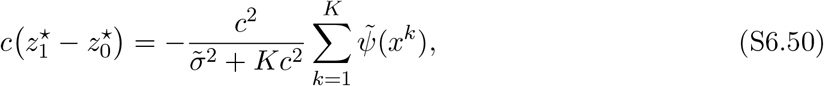

where

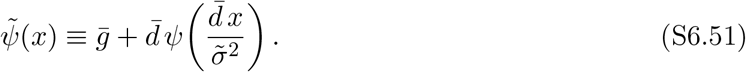

Therefore

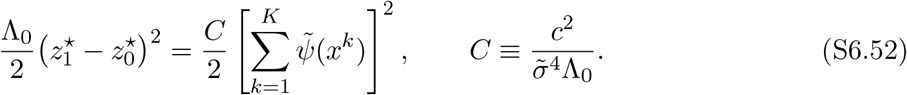

Dropping the curvature correction 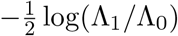, which is numerically small over the parameter ranges studied here, we arrive at the saddle approximation

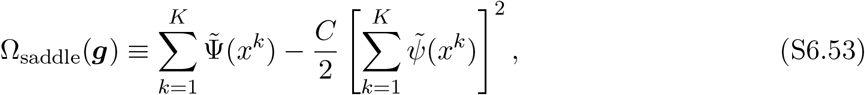

up to an additive constant.

#### Special case: balanced plasticity

When 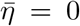, we have *c* = 0, so the global fluctuation vanishes and the pairwise term in (S6.53) disappears. In this case the fluctuations over latent *z* vanish, and the optimal decoder reduces to a separable decoder:

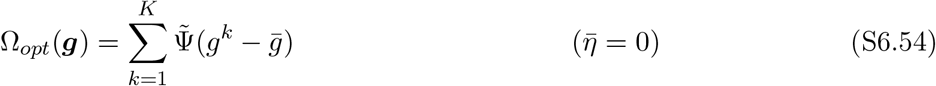

### S6.4 Recurrent-Circuit Interpretation of the Saddle Decoder

The saddle approximation (S6.53) can be rewritten in a form suggestive of recurrent computation in the PC layer. Define the PC firing rate

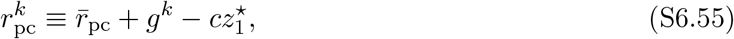

where we interpret 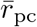 as the PCs baseline firing rate. Here, we are assuming that PCs are in a linear operating regime. For compactness, we will work in terms of the PCs deviation from baseline firing rate

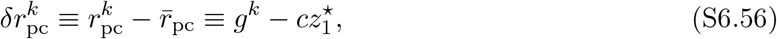

so that 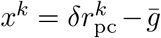. We then absorb this shift into the nonlinearities by a slight abuse of notation:

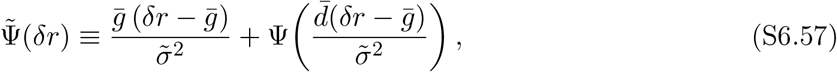

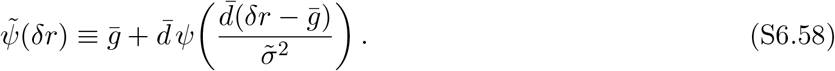

Using 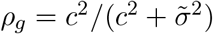, the saddle condition can be rewritten as

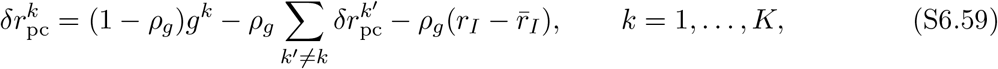

where the Purkinje-layer interneuron (PLI) activity is defined by

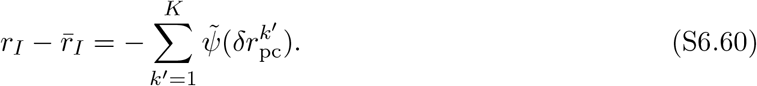

Equations (S6.59)–(S6.60) may then be interpreted as the fixed point of the recurrent dynamics

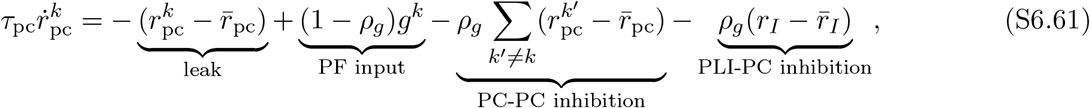

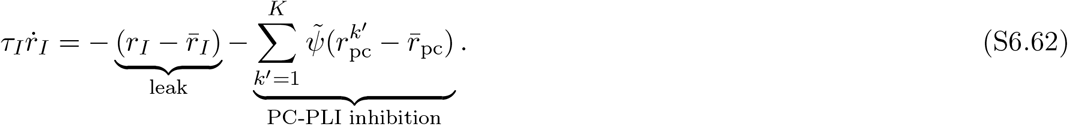

In terms of the fixed-point firing-rate deviations 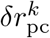, the saddle decoder becomes

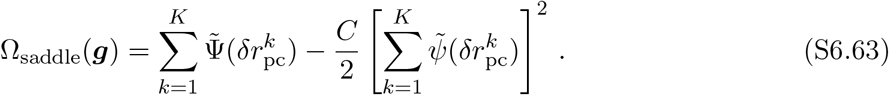

Expanding the square,

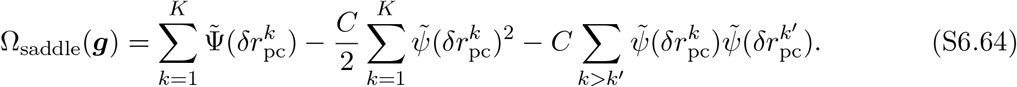

The diagonal term scales as *O*(*K*), whereas the pairwise term scales as *O*(*K*^2^). Ignoring the contribution of the diagonal terms in the double sum (which is sub-leading in ensemble size *K*) yields the simplified decoder used in the main text,

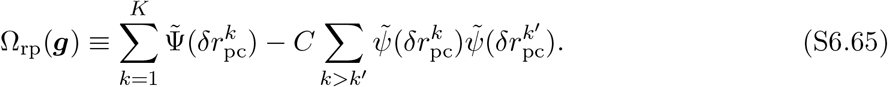

#### Sparse Plasticity Special Case

We consider the special cases where each PC undergoes either LTD or no plasticity during learning (*η*_+_ = 0) and the case where each PC undergoes either LTP or no plasticity during learning (*η*_−_ = 0) in SI figs S6 and S7. In this case, we find that the optimal nonlinearities of the recurrent + pairwise decoder reduce to softplus and sigmoidal forms.

For example, taking *η*_−_ = 0 we get 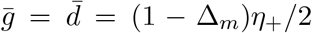 and 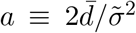 and the relevant nonlinearities reduce to:

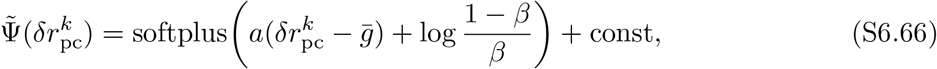

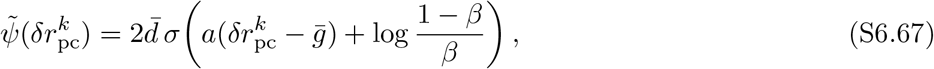

where softplus(*z*) ≡ log(1 + *e*^*z*^) and *σ*(*z*) ≡ (1 + *e*^−*z*^)^−1^.

**Figure S6:**
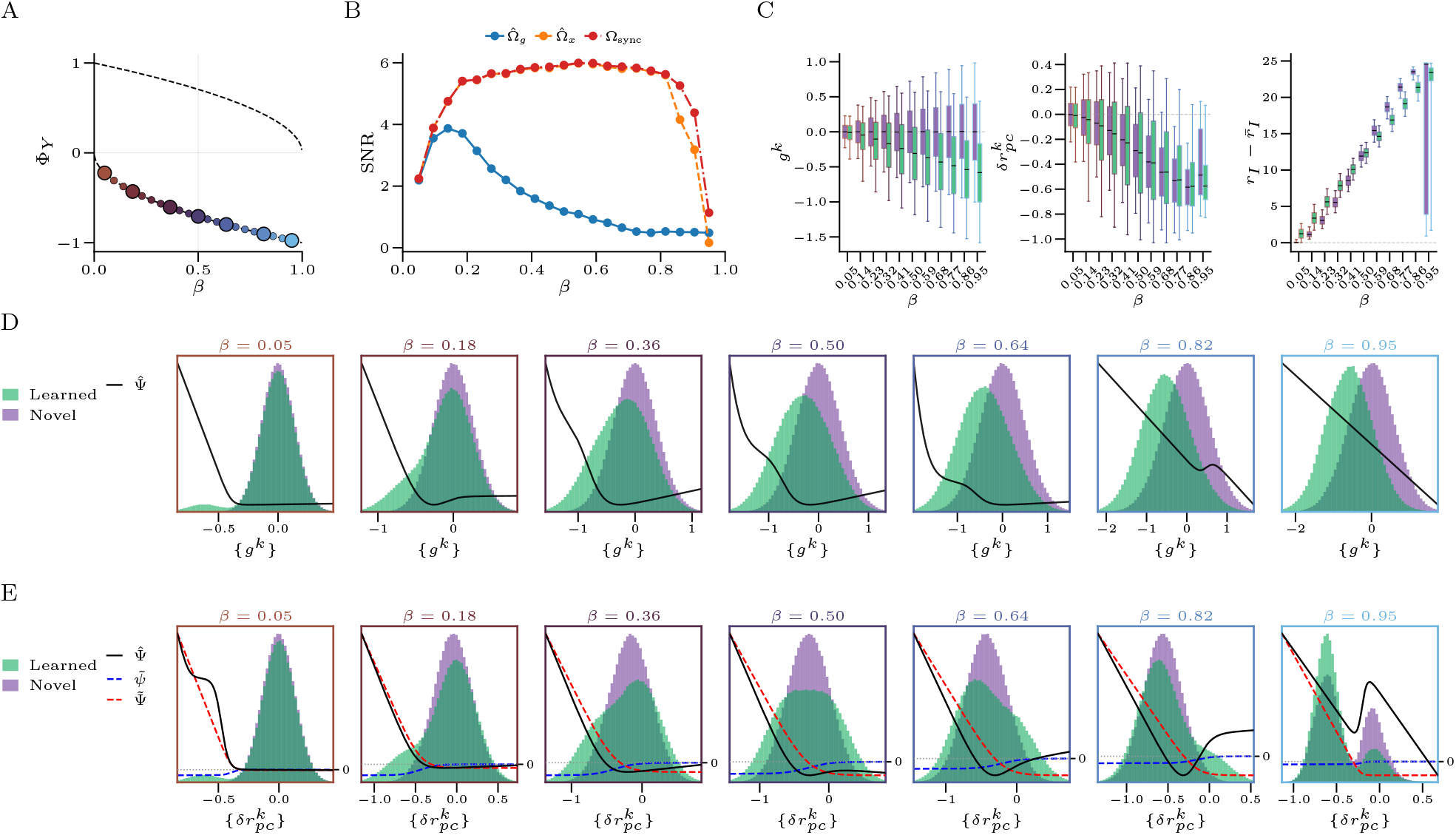
Optimal decoding and PC activity under sparse-LTD learning. **(A)** Operating points used in panels (B–E), plotted in the (*β*, Φ_*Y*_) plane. Colored markers lie on the sparse-LTD line 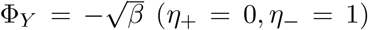. **(B)** SNR versus LTD probability *β* for three decoder classes: the raw separable decoder 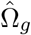, the recurrent separable decoder 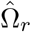, and the full saddle approximation to the optimal decoder Ω_rp_. **(C)** Box plots of raw PC inputs *g*^*k*^, recurrent fixed-point deviations 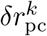, and PLI activity 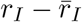 for novel and learned stimuli across the sweep in *β*. (Marginal distributions of raw PC inputs *g*^*k*^ for novel (purple) and learned (green) stimuli at the selected operating points. Black curves show the fitted pointwise nonlinearity 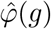 (Methods §3). (**E**)Marginal distributions of recurrent fixed-point deviations 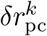 at the same operating points. Black curves show the fitted recurrent nonlinearity 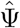 (Methods §3); dashed curves show the saddle-derived nonlinearities 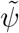 and 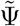 for comparison. Parameters: *η*_+_ = 0, *η*_−_ = 1, *K* = 40, *P* = 10,000, *N*_*s*_ = 1,400, *N*_*m*_ = *N*_*d*_ = 400,000, *f* = 0.05, Δ_*x*_ = 0.10, *σ*_0_ = 0.1, *o* = 1.

**Figure S7:**
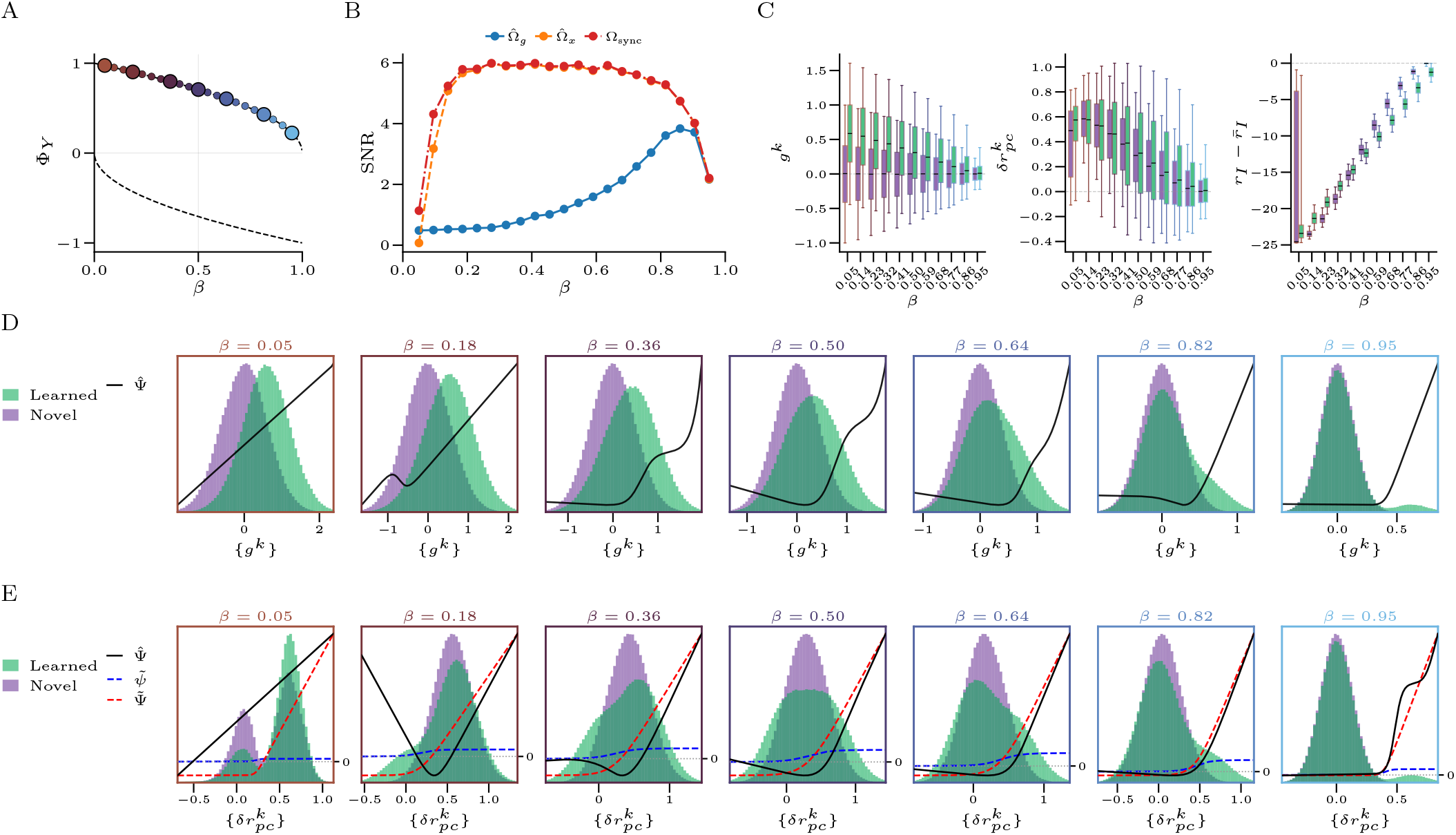
Optimal decoding and PC activity under sparse-LTP learning. **(A)** Operating points used in panels (B–E), plotted in the (*β*, Φ_*Y*_) plane. Colored markers lie on the sparse-LTP line 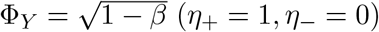. **(B)** SNR versus *β* for three decoder classes: the raw separable decoder 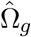, the recurrent separable decoder 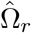, and the full saddle approximation to the optimal decoder Ω_rp_. **(C)** Box plots of raw PC inputs *g*^*k*^, recurrent fixed-point deviations 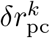, and PLI activity 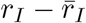 for novel and learned stimuli across the sweep in *β*. **(D)** Marginal distributions of raw PC inputs *g*^*k*^ for novel (purple) and learned (green) stimuli at the selected operating points. Black curves show the fitted pointwise nonlinearity 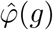 (*g*) (Methods §3). **(E)** Marginal distributions of recurrent fixed-point deviations 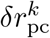 at the same operating points. Black curves show the fitted recurrent nonlinearity 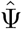 (Methods §3); dashed curves show the saddle-derived nonlinearities 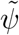 and 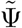 for comparison. Parameters: *η*_+_ = 1, *η*_−_ = 0, *K* = 40, *P* = 10,000, *N*_*s*_ = 1,400, *N*_*m*_ = *N*_*d*_ = 400,000, *f* = 0.05, Δ_*x*_ = 0.10, *σ*_0_ = 0.1, *o* = 1.

### S6.5 Moment-Matched Approximate Optimal Decoder

As a simpler approximation, we may replace the learned-class Gaussian mixture by the Gaussian distribution with matching first and second moments. For *e* = 0, we already have

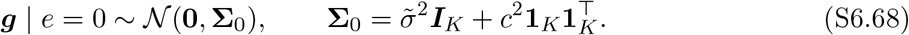

For *e* = 1, the random signal term contributes an additional diagonal variance

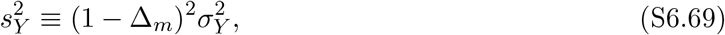

while the mean is

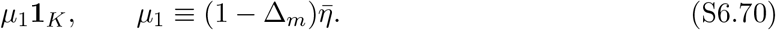

The moment-matched Gaussian approximation is then written

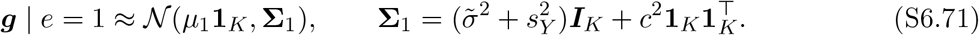

Both **Σ**_0_ and **Σ**_1_ have compound-symmetric form 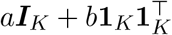. We use the standard identities

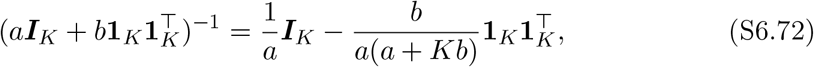

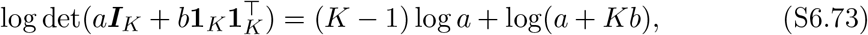

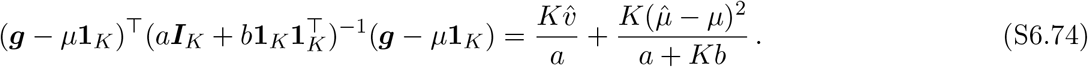

Applying these with 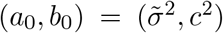 and 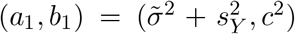, the moment-matched log-likelihood ratio simplifies, up to an additive constant and positive rescaling, to

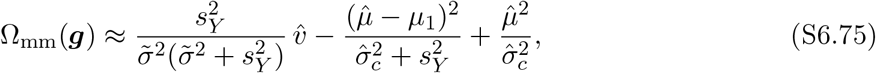

where 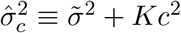.

#### Special case: balanced plasticity

If 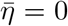, then *c* = 0, *µ*_1_ = 0, and 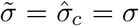. Simplifying, we obtain

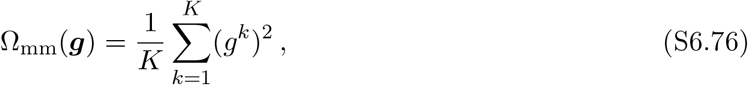

recovering the simple quadratic decoder.

### S6.6 Dynamical Decoding of Recurrent PC Dynamics

We consider here the recurrent decoder

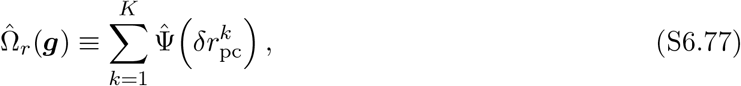

where 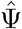 is the fitted separable DCN nonlinearity (Methods §3) and 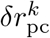 denotes the *k*-th PCs deviation from baseline firing rate at the fixed point of the recurrent dynamics (S6.61)–(S6.62). As written, 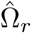 is a function of the *fixed-point* PC firing rates, whereas the biological DCN integrates PC simple spikes continuously. To reconcile this, we may promote the network output Ω to a time-dependent quantity by evaluating the same separable score on the instantaneous firing-rate deviations,

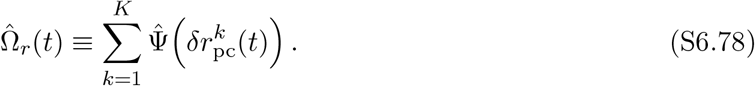

**Figure S8:**
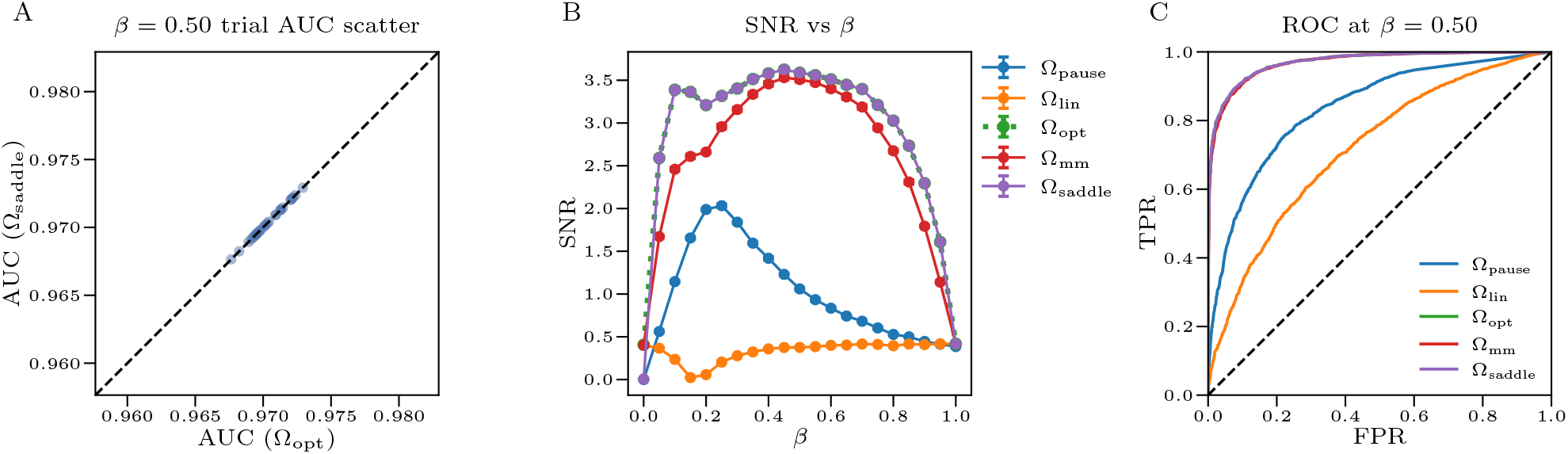
Comparison of decoder architectures under heterogeneous plasticity. **(A)** Trial-level AUC comparison between the numerically evaluated optimal decoder Ω_opt_ (computed via its integral representation) and the saddle-point approximation Ω_saddle_. Each marker represents a single Monte Carlo trial (with fixed ***Y*** matrix, *N* = 5,000 patterns per class); the dashed line indicates AUC(Ω_saddle_) = AUC(Ω_opt_). **(B)** AUC versus LTD probability *β* for five decoders: the optimal decoder Ω_opt_, saddle-point approximation Ω_saddle_, moment-matched approximation Ω_mm_, linear decoder Ω_lin_(***g***) = ∑_*k*_ *g*^*k*^, and pause-based decoder Ω_pause_(***g***) = ∑_*k*_ Θ[*κ* − *g*^*k*^], where the threshold *κ* is optimized separately at each *β*. Error bars indicate s.e.m. across *n* = 15 independent trials. **(C)** Trial-averaged receiver operating characteristic (ROC) curves at *β* = 0.5 for the same set of decoders. Simulations used *K* = 40 Purkinje cells, *N*_*d*_ = 200,000, *N*_*m*_ = 400,000, *N*_*s*_ = 1,400, *P* = 1.2 × 10^4^ learned patterns, granule-cell sparsity *f* = 0.05, sensory-layer noise Δ_*x*_ = 0.10, intrinsic noise *σ*_0_ = 0.1, overlap fraction *o* = 1.0, *η*_+_ = 0.2, *η*_−_ = 1.0. Panels (A) and (C) fix *β* = 0.5.

Differentiating with respect to time and applying the chain rule, we obtain

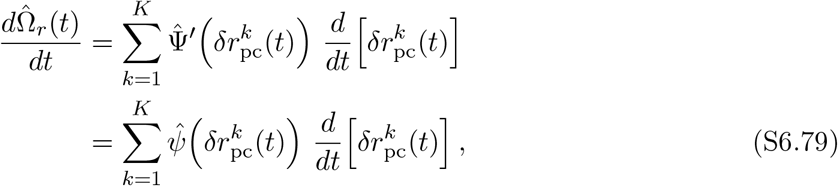

where 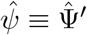. Integrating (S6.79) over the interval [*t*_0_, *t*] gives

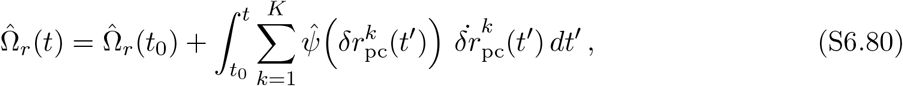

with 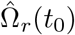 absorbed as an additive constant. Discretizing with step size Δ*t* recovers an incremental update rule:

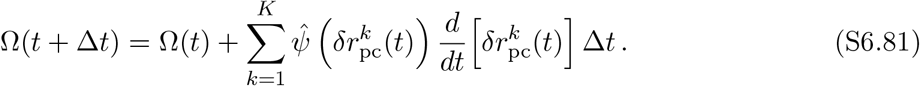

The same derivation applied to the pairwise saddle-point decoder Ω_rp_ (eq. S6.65) yields an analogous update rule with an additional pairwise interaction term:

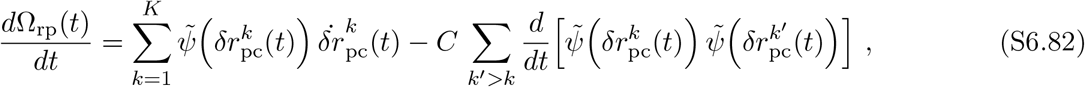

but since near-identical performance can be achieved with further. 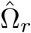, we do not investigate this form

#### Numerical verification over a long stimulus sequence

We investigate the behavior of PC recurrent dynamics, and the incrementally tracked DCN state Ω(*t*) in fig. S9. There, we stimulate PC ensembles trained at the operating points indicated in fig. 4D with an alternating sequence of unique novel and learned input stimuli. As the network evolves under the optimal recurrent dynamics (eq. S6.61), individual PCs increase and decrease their activity seemingly randomly as new stimuli are shown. However, the incrementally tracked Ω(*t*) output reliably increases upon presentation of learned stimuli and decreases upon presentation of novel stimuli. PC ensemble activity may therefore reliably encode the identity of sensory stimuli, even as individual PCs respond erratically to any given stimulus.

**Figure S9:**
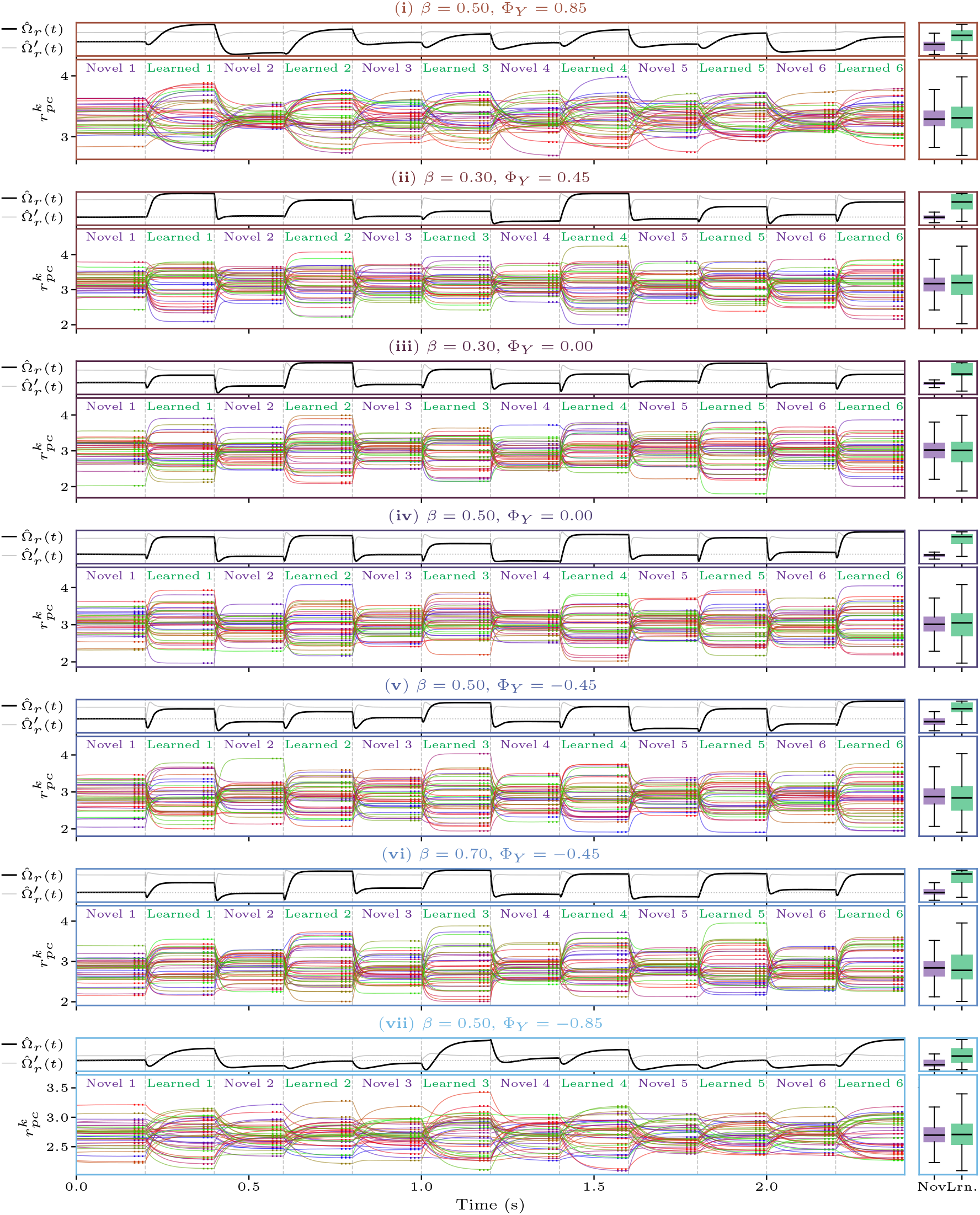
Recurrent PC dynamics and incremental Ω(*t*) across a long alternating stimulus sequence. At each of the seven operating points from fig. 4C (labels i–vii, top to bottom), the recurrent PC–PLI circuit (S6.61)–(S6.62) is driven by a sequence of twelve alternating novel and learned stimuli, each a freshly sampled ***g*** vector presented for *T*_stim_ = 200 ms. **Top subpanel of each block:** running DCN tally Ω(*t*) from eq. (S6.81). **Bottom subpanel of each block:** per-PC firing-rate trajectories 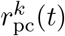, one curve per PC; dotted segments at the right of each interval mark the analytical recurrent fixed points from eqs. (S6.59)–(S6.60). **Side box plots:** distributions of Ω (top) and 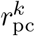 (bottom) sampled on a uniform time grid within the novel (purple) and learned (green) intervals, with *y*-axes aligned to the corresponding main subpanel. Parameters: *K* = 40, *σ*_0_ = 0, *τ*_pc_ = 10 ms, *τ*_*I*_ = 1 ms, Δ*t* = 50 *µ*s.

## S7 Purkinje Cell Correlations

In this section, we derive the pairwise statistics relating PC plasticity histories to (i) the similarity of their learned PF–PC synaptic weights and (ii) the correlations in their firing rates. Throughout, we work in the large-*P* regime and under the homogeneous-overlap assumption.

For any pair of PCs *k, k*^*′*^, define the empirical first and second moments of their plasticity histories:

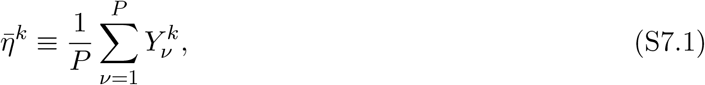

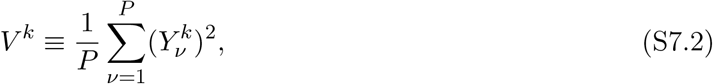

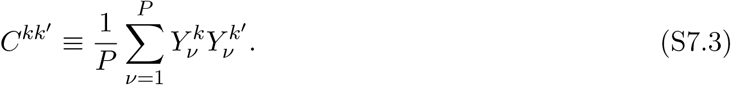

We further define the *pairwise plasticity alignment*

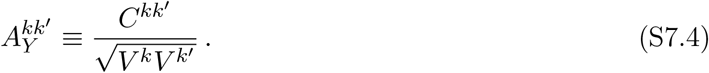

Under heterogeneous plasticity it is convenient to write

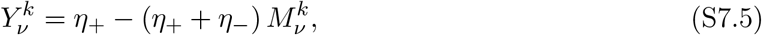

where 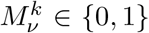, with 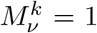 denoting LTD and 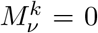 denoting LTP. Define the empirical LTD fraction and pairwise LTD correlation

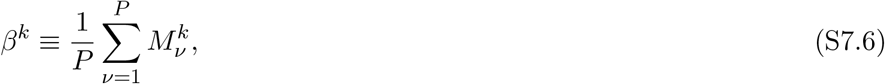

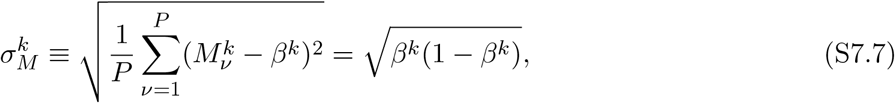

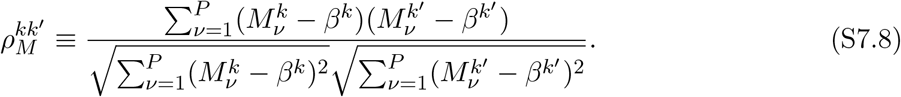

Here, 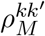 corresponds to the Pearson correlation of the effective plasticity histories in PCs *k* and *k*^*′*^. Though they do not correspond exactly because not all complex spikes necessarily induce plasticity, 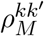 provides a theoretical analogue to the Pearson correlation of complex spike trains as measured in experiments [72].

Throughout this section, we will simplify expressions by replacing the single-PC statistic *β*^*k*^ with its expected value *β*, as we expect the *β*^*k*^ to concentrate at large *P* under heterogeneous plasticity training. Under this approximation, we obtain:

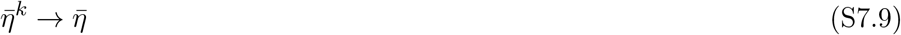

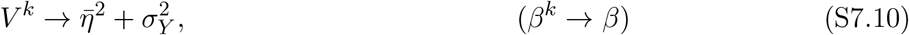

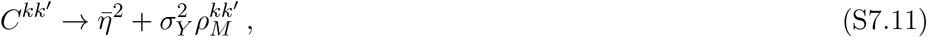

and dividing by 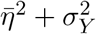 yields

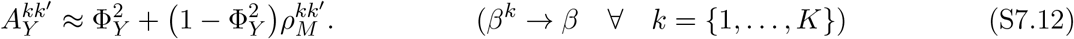

Applying this expression in the following sections, we find that PC–PC similarities have baseline pairwise similarity set by the population plasticity ratio Φ_*Y*_, with pair-to-pair fluctuations controlled by 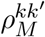.

### S7.1 Correlated PF–PC Synaptic Weights

We now compute the expected weight correlation 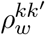 between the learned weights of PCs *k* and *k*^*′*^. Consider a PF *i* that is sampled by both PCs. From the weight formula above,

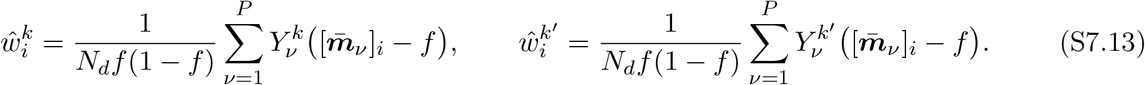

Since 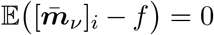 and 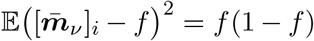, the central limit theorem gives, for large *P*,

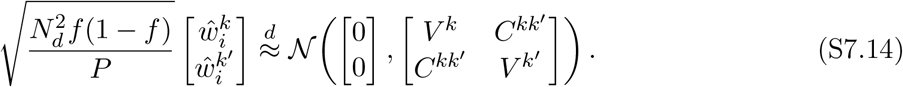

For a particular realization of the learned weights ***ŵ***^*k*^, ***ŵ***^*k*^*′* we define the similarity metric:

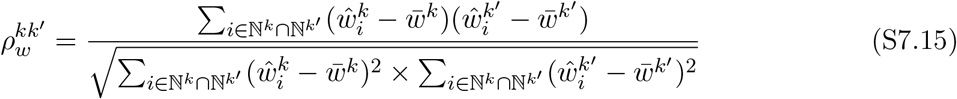

which is the Pearson correlation between the weight vectors learned by PCs *k* and *k*^*′*^, restricted to the subset of parallel fibers ℕ^*k*^ ∩ ℕ^*k*^*′* which form at least one synapse onto both by PCs *k* and *k*^*′*^. It follows from eqs S7.14 and S7.4 that

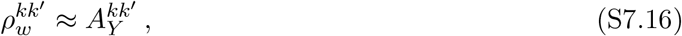

With fluctuations about this expected value vanishing as |ℕ^*k*^ ∩ ℕ|^*k*^*′* grows. Substituting the decomposition (S7.12) yields eq. 11 in the main text. Thus the mean connectivity overlap depends only on the population plasticity ratio Φ_*Y*_, while pair-to-pair fluctuations around this mean are driven by fluctuations in 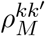. We verify that the expected weight similarity is invariant to *β* for fixed Φ_*Y*_ in fig. S10.

### S7.2 Simple-spike synchrony

We next derive the predicted simple-spike synchrony between PCs *k* and *k*^*′*^. While our model is static in nature, we may introduce a *post hoc* temporal extension by considering PCs to spike as Poisson processes with rate dependent on their integrated inputs *g*^*k*^(*t*):

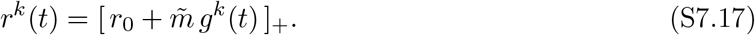

While many past works suggest that coupled PCs dynamically synchronize their spikes on millisecond timescales, recent work Herzfeld et al. [6] find that during smooth pursuit eye movements, floccular PCs fire synchronous spikes at roughly the rate expected from co-modulation of their underlying firing rates. Under the latter view, we may write the instantaneous rate of synchronous simple spikes between PCs *k* and *k*^*′*^ as the product of rates:

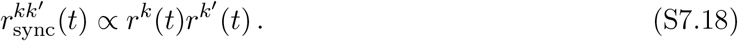

**Figure S10:**
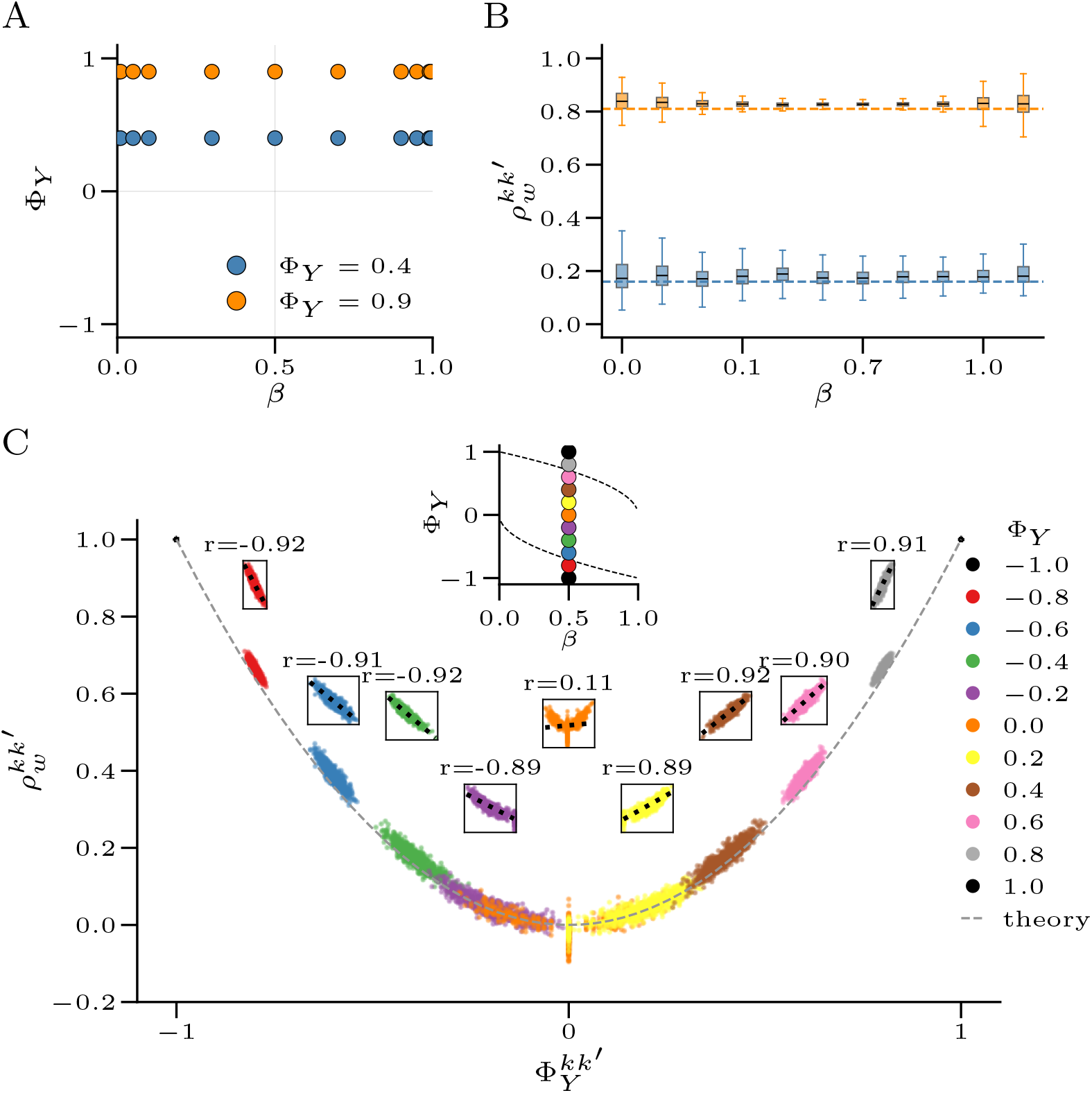
Weight correlation 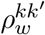 depends on plasticity alignment, not on LTD probability. **(A)** Operating points in the (*β*, Φ_*Y*_) plane used for *β*-sweeps at two fixed plasticity-alignment values, Φ_*Y*_ = 0.4 (blue) and Φ_*Y*_ = 0.9 (orange). **(B)** Box plots of pairwise weight correlation 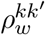 between learned PF–PC weight vectors across *β* for the operating points in (A). Dashed horizontal lines show the theoretical prediction 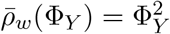, confirming that 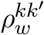 is independent of *β* at fixed Φ_*Y*_. **(C)** Scatter of 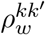 versus the signed root of pairwise plasticity alignment 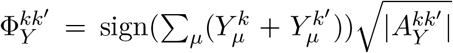 across a sweep of Φ_*Y*_ ∈ {−1.0, −0.8, …, 0.8, 1.0} at fixed *β* = 0.5. Dashed gray curve shows the same theoretical prediction. Insets magnify individual Φ_*Y*_ clusters; dotted lines show linear fits with Pearson correlation *r* annotated above each inset. Simulations used *K* = 40 Purkinje cells, *N*_*s*_ = 1,400, *N*_*m*_ = 40,000, *N*_*d*_ = *N*_*m*_, *P* = 1,000 learned patterns, *f* = 0.05, and Δ_*x*_ = 0.10.

To generate PC inputs ***g***(*t*), we partition a simulation interval [0, *T*] into smaller windows with length drawn uniformly from 50 to 100ms. Within each window, the PC input currents ***g***(*t*) are drawn either from the learned distribution *P* (***g***|*e* = 1) with probability *p*_l_, or from the novel distribution *P* (***g***|*e* = 0) with probability 1 − *p*_l_. We will denote the class presented at time *t* with the indicator variable *e*_*t*_ ∈ {0, 1}.

We choose *r*_0_ to be sufficiently large that the rectification is rarely engaged, so we may write

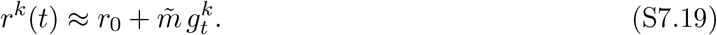

The synchrony score is defined by the fractional excess probability of coincident spikes above baseline:

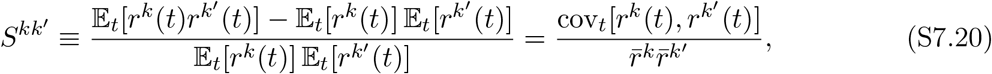

where 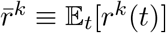.

Using the linear-rate approximation,

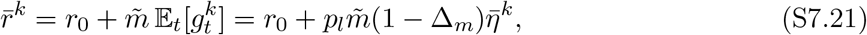

and

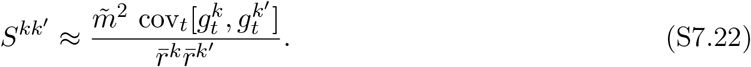

We therefore only need the covariance of the PC inputs across the sequence of time windows. Conditioned on *e*_*t*_ = 0 (novel input), the mean response is zero and

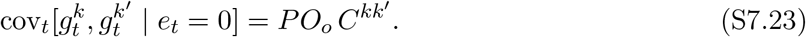

Conditioned on *e*_*t*_ = 1 (learned input), the mean response is 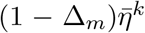, and the law of total covariance gives

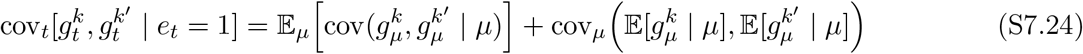

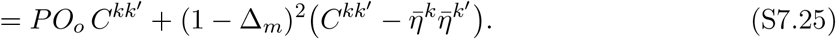

Applying the law of total covariance once more over *e*_*t*_,

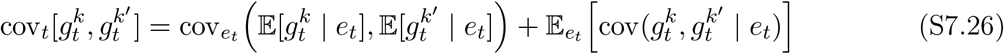

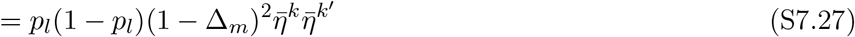

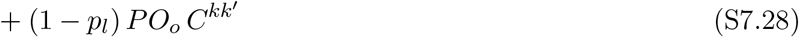

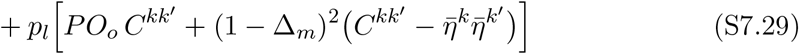

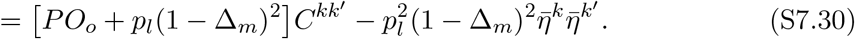

Substituting this into (S7.22) gives the general pairwise synchrony formula

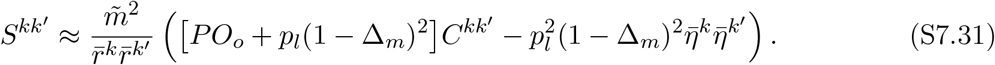

Finally, under heterogeneous plasticity at large *P*, we use the concentration approximations

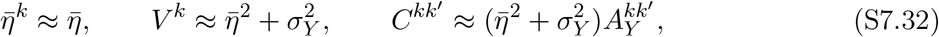

together with (S7.12). Writing

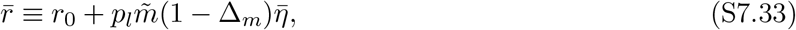

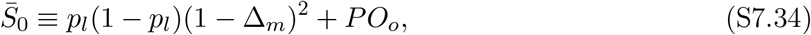

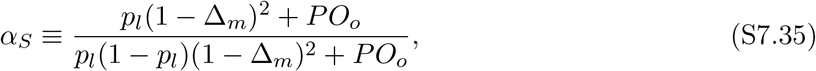

we obtain

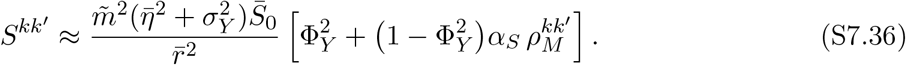

This is the expression used in the main text. We see that *S*^*kk*^*′* clusters around a baseline synchrony level set by the population plasticity bias Φ_*Y*_ as well as the baseline firing rate 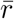, slope 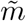, and plasticity variance 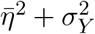. Pair-specific fluctuations around this baseline synchrony level are then driven by the spontaneous correlations in plasticity histories, captured by 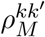.

## S8 Extension to Multi-Shot Learning

In the main text and preceding SI sections, we studied a one-shot learning rule in which the PC ensemble is trained on a single pattern from each cluster (which was taken to be the cluster center 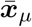 for mathematical convenience). However, in realistic scenarios, an animal may experience repeated pairings between a conditioned and unconditioned stimulus while learning an association. Here, we extend our analysis to a repeated-training, or *multi-shot*, setting in which each cluster is presented *L* times during training. We show that, with an appropriate normalization, the effective plasticity signals converge to a Gaussian distribution as *L* → ∞, and that in the balanced fully connected case the resulting SNR and capacity under quadratic decoding obey the same qualitative scaling laws as in the single-shot setting, including capacity scaling as as 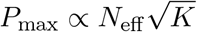 up to a quadratic plateau 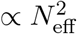.

### S8.1 Effective plasticity signals under repeated training

Suppose that the training set contains *L* samples from each of the *P* learned clusters. Let ***x***_*µ,ℓ*_ denote the *ℓ*-th training example drawn from cluster *µ*, and let ***m***_*µ,ℓ*_ = ***m***(***x***_*µ,ℓ*_) be the corresponding mixedlayer representation. The training patterns are drawn from each cluster *µ* by flipping each element of the cluster center 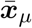 with probability 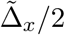, so that 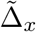 is the training-time cluster size. We then write the multi-shot learning rule:

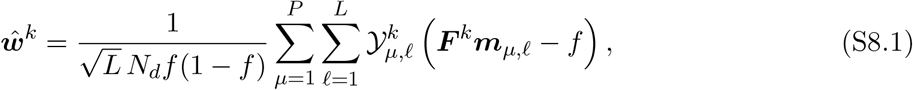

where 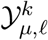 is the plasticity signal received by PC *k* on the *ℓ*-th presentation of cluster *µ*. The prefactor 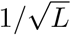 is chosen so that the total variance of the effective plasticity signal remains *O*(1) as *L* grows.

Here, we will again simplify by considering only the limit in which the training patterns are equal to the cluster centers:

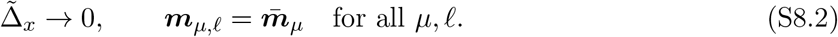

In this case the learning rule simplifies to

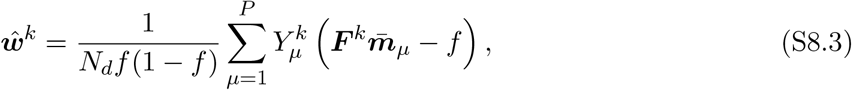

with *effective* plasticity signals

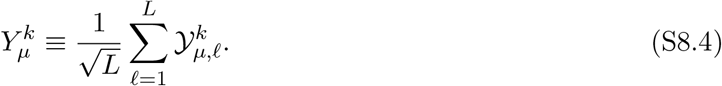

Equation (S8.3) has exactly the same form as the single-shot learning rule analyzed in §S3.1. Therefore all results derived there apply directly once the single-shot variables 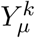 are replaced by the effective multi-shot variables (S8.4).

Now assume that the microscopic plasticity events are i.i.d. draws

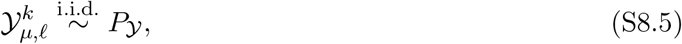

with mean and variance

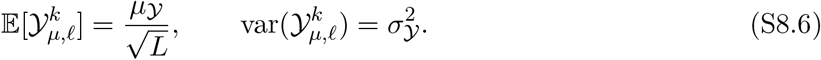

Then the effective signal satisfies

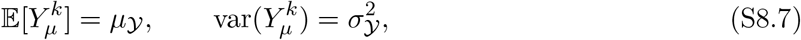

independently of *L*. By the central limit theorem, if *P*_*Y*_ has finite third absolute moment then

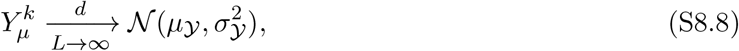

with Berry–Esseen corrections of order *L*^−1/2^.

### S8.2 Gaussian plasticity model

Motivated by the large-*L* limit above, we now study the tractable case in which the effective plasticity signals are themselves exactly Gaussian:

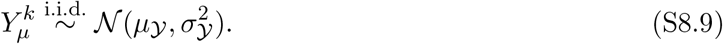

We specialize to the balanced case

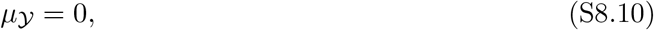

and to the fully connected architecture *N*_*d*_ = *N*_*m*_.

This model is a special case of the fully connected heterogeneous-plasticity analysis from section S3.4, obtained by setting

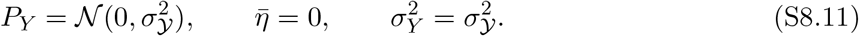

Therefore the fully connected heterogeneous-plasticity statistics apply directly: conditioned on a fixed plasticity matrix ***Y***, the novel and learned PC inputs are Gaussian,

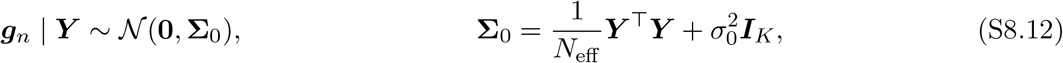

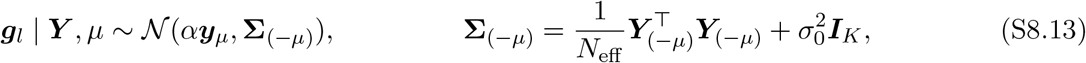

with *α* ≡ 1 −Δ_*m*_, ***y***_*µ*_ the *µ*th row of ***Y***, and ***Y***_(−*µ*)_ the matrix ***Y*** with row *µ* deleted. This is precisely the fully connected heterogeneous-plasticity model analyzed in Section S5, except that the single-site plasticity distribution *P*_*Y*_ is now Gaussian rather than two-point. Averaging the learned signal 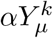 over ***Y*** shows that, marginally, the novel and learned classes differ only in their second moment: learned inputs carry the additional variance

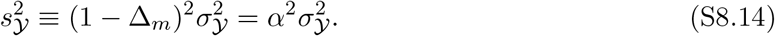

### S8.3 SNR and Capacity under Quadratic Decoding

Because the learned and novel classes differ only through a shift in the conditional variance, a natural choice of decoder is the separable quadratic decoder:

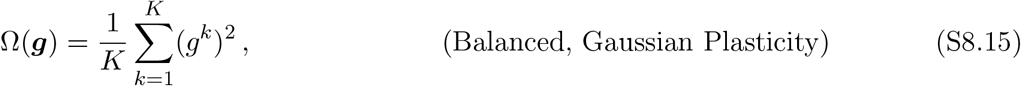

corresponding to the “moment-matched” approximation to the optimal decoder (see Section S6.5). As in sec. S5, we will approximate SNR using the ***Y***-averaged output moments and Gaussian SNR formula (eq. S5.19). A useful identity is the following. For any 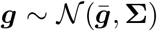, with 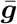 fixed, we will have:

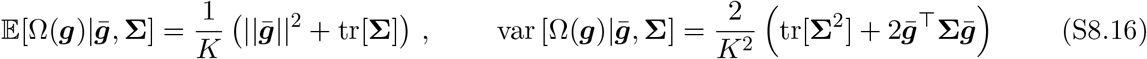

We next take the expectations of these moments over the random plasticity schedules ***Y*** for the novel and learned input conditions.

#### Novel inputs

The novel quadratic score 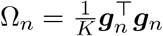 has the fixed-***Y*** moments

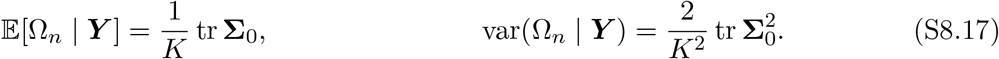

For Gaussian plasticity, 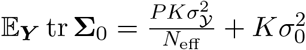 and

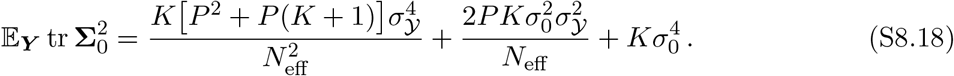

Retaining the leading contributions at large *P* and *K* gives the expected novel mean and variance:

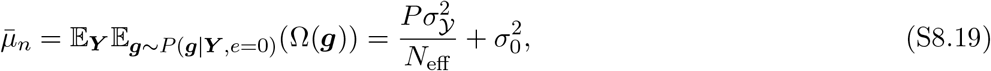

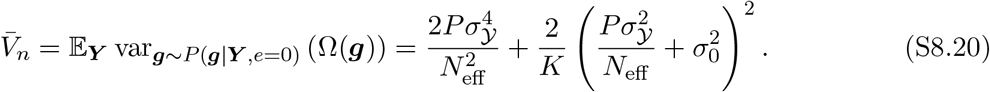

#### Learned inputs

For a learned input from cluster *µ*, ***g***_*l*_ | ***Y***, *µ* ~ *N*(*α****y***_*µ*_, **Σ**_(−*µ*)_), so that for *µ* ≥ 1

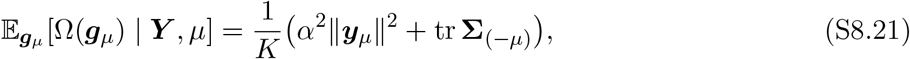

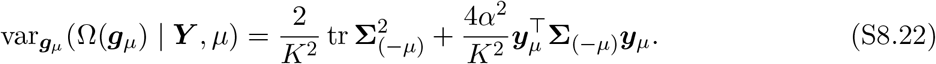

Because **Σ**_(−*µ*)_ and ***y***_*µ*_ are independent, we can convert 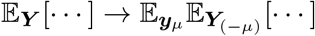. Using standard Gaussian moment results and the law of total variance, we obtain the average moments at leading order for *P, K* ≫ 1, following similar steps to section S5:

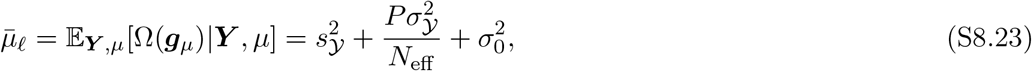

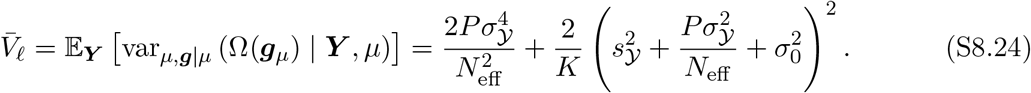

#### SNR Formula

Using the Gaussian SNR approximation (S5.19), we then obtain

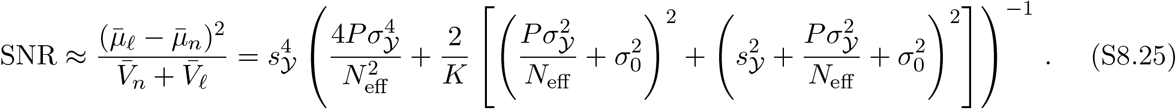

#### Capacity in the noise-free case

To obtain a capacity curve, we set 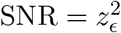 (AUC = 1 − *ϵ*)and solve for *P*_max_. In the special case where *σ*_0_ = 0, we obtain a relatively simple capacity formula:

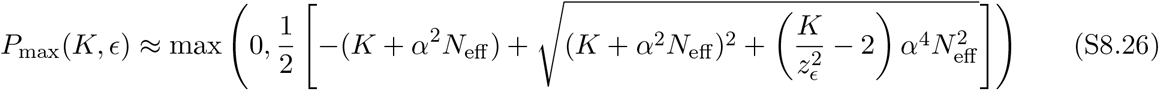

where *α* = 1 − Δ_*m*_.

#### Scaling regimes

Capacity can then be simplified in the following regimes:

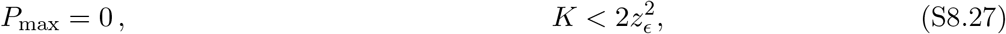

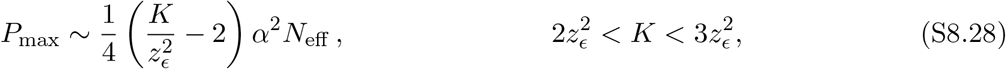

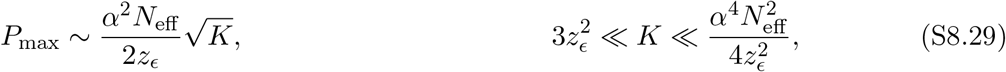

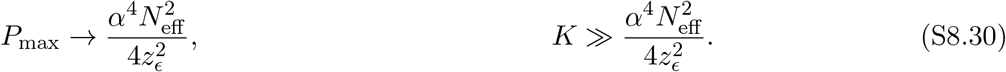

Thus repeated training exhibits a similar capacity scaling profile to networks trained with the one-shot learning rule with quadratic decoder. At small 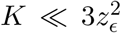, capacity may grow linearly but remains below the direct readout capacity *P*_max,DR_. Following this initial low-capacity regime, capacity grows as 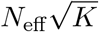, and for sufficiently large ensembles saturates at a plateau proportional to 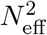. In the limit *N*_*d*_ → ∞, where *N*_eff_ → *N*_*s*_/*Q*^2^, this again yields quadratic scaling with the effective input dimensionality. However, we note that the 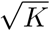 capacity scaling provides rapidly diminishing returns as *K* increases, requiring *K > P*_max_ PCs to achieve the saturation capacity. We therefore do not expect PC ensembles to operate at or near the capacity saturation regime, but rather to benefit from the initial fast growth of the 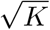 scaling.

## S9 Extension to Block-Structured Plasticity

Axons of neurons from the inferior olive typically branch into ~7 climbing fibers, which innervate separate Purkinje cells. This raises the possibility that multiple PCs in a functional ensemble receive inputs from the same inferior olive neuron, as was suggested in a review by Shadmehr [9]. Here, we consider the case where the *K* PCs in the functional ensemble are partitioned into *N*_*b*_ disjoint blocks 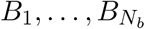 of equal size

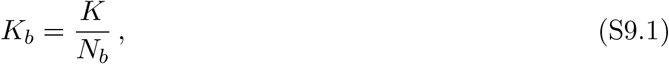

so that all PCs in block *b* share the same plasticity signals from their shared inferior olive neuron:

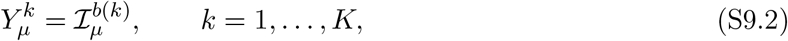

where *b*(*k*) ∈ {1, …, *N*_*b*_} gives the index of the inferior olive neuron which innervates PC *k* and 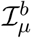 represents the plasticity signal from inferior olive neuron *b* upon presentation of training stimulus *µ*. We then draw the IO plasticity signals independently across *b* and *µ* from the same heterogeneous plasticity distribution as before (i.e. 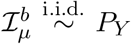, for *P*_*Y*_ defined in eq. S3.43). The limit *K*_*b*_ = 1 reduces to the heterogeneous plasticity learning rule, but when *K*_*b*_ *>* 1, PCs will be grouped into clusters which receive the same plasticity signals during training. In Monte Carlo simulations, we sample the 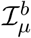, then construct the ***Y*** matrix using eq. S9.2, then sample learned PC inputs by plugging this ***Y*** matrix into the general statistics in eq. S3.24.

To derive the corresponding decoder, we adopt the same expected-covariance approximation used for heterogeneous plasticity: the pattern-dependent covariance matrix generated by a particular realization of the plasticity signals is replaced by its expectation over the block-level plasticity distribution.

### S9.1 Optimal Decoding Under Block-Structured Plasticity

Next, we determine the optimal decoding strategy Ω_opt_(***g***) after training with block-structured plasticity. We will find that the optimal decoder under block-structured plasticity may be implemented by first reducing the full *K*-dimensional activity vector ***g*** to the *N*_*b*_ block averages, and then applying the optimal decoder for the heterogeneous-plasticity case to these averages with suitably renormalized parameters. As in the heterogeneous-plasticity analysis, we work under the expected-covariance approximation, replacing the pattern-dependent covariance matrix by its expectation over the block-level plasticity variables:

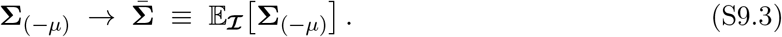

Defining

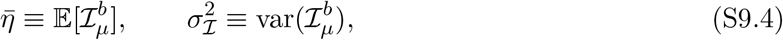

we have

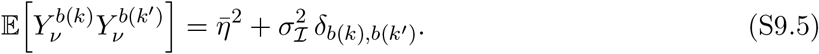

Writing the overlap matrix as

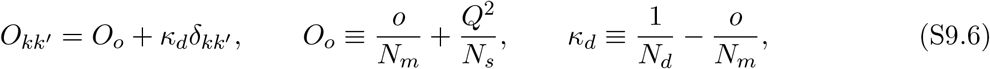

the expected covariance becomes

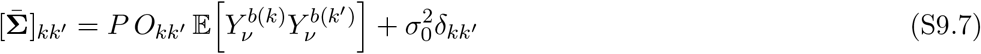

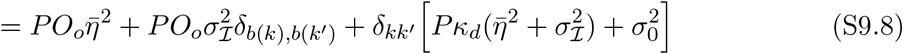

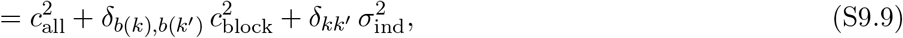

where we have defined the three variance components

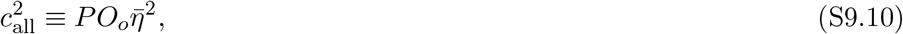

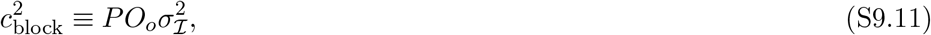

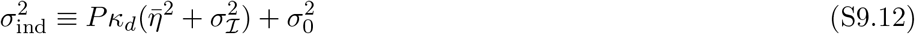

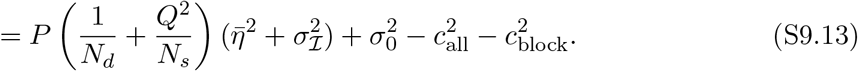

We now compute the optimal likelihood-ratio decoder under this covariance structure. The block covariance in eq. (S9.9) admits the latent-factor representation

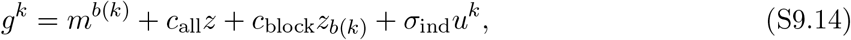

where

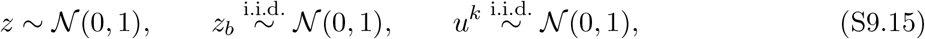

and the class-dependent block means are

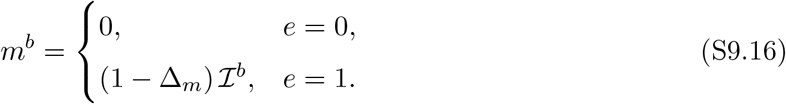

Equivalently, conditioned on *z*, {*z*_*b*_}, and 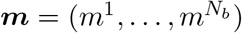, all PCs in block *b* are i.i.d. Gaussians with common mean

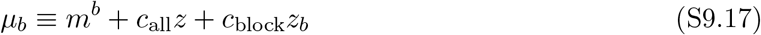

and variance 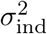. Thus

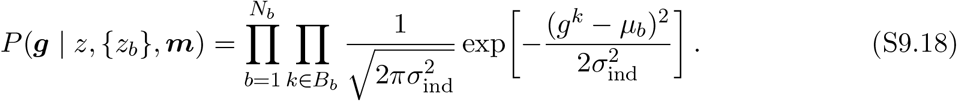

Next, define the block averages

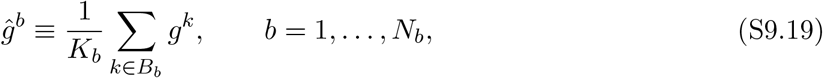

and the within-block residual sums of squares

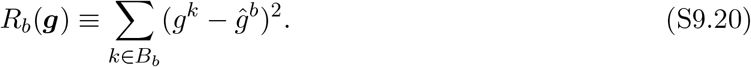

Using the standard decomposition

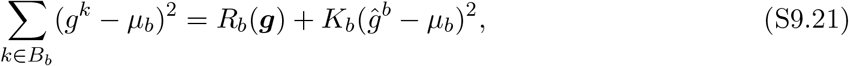

we may rewrite the conditional likelihood as

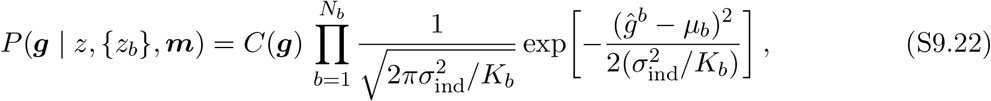

where

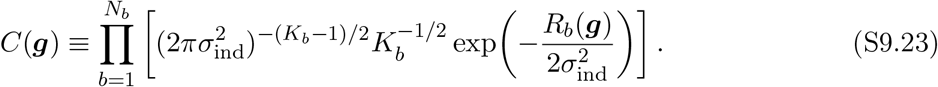

Crucially, *C*(***g***) depends on ***g*** but *not* on the latent variables (*z*, {*z*_*b*_}, ***m***), and therefore appears identically in the likelihoods of both classes. It follows that *C*(***g***) cancels exactly in the likelihood ratio. The optimal decoder therefore depends on the full PC activity vector only through the block averages:

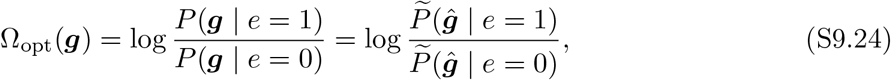

where 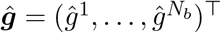, and

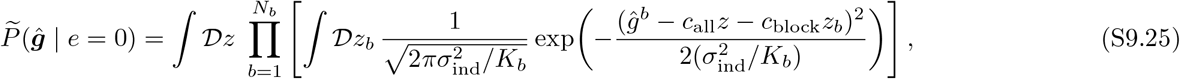

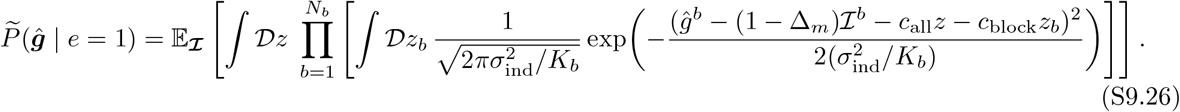

We may now integrate out the block-specific Gaussian latents *z*_*b*_ one block at a time. Defining

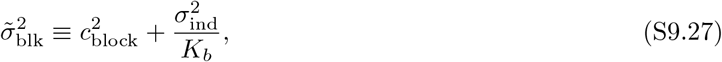

the block averages admit the reduced latent representation

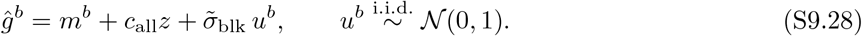

Equivalently,

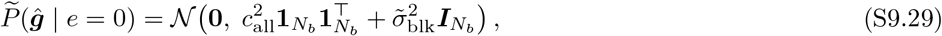

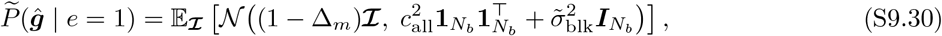

where 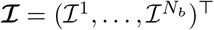.

The statistics of the block-averages therefore have the exact same form as the statistics of individual PCs trained with the heterogeneous plasticity learning rule. Because the optimal decoder depends only on these block-averages, it is equivalent to the optimal decoder for heterogeneous plasticity with the substitutions

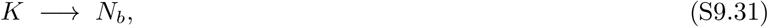

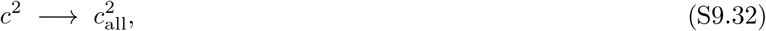

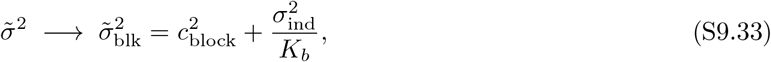

with the scalar activity variable *g* replaced everywhere by the block average *ĝ*. In practice, all approximate decoders derived for heterogeneous plasticity may therefore be reused in the block-structured setting after first replacing the full PC activity vector by the block averages and making the parameter substitutions in eq. (S9.33). In the simulations for block-structured plasticity presented here, we use the moment-matched decoder (section S6.5) applied to ***ĝ***.

## S10 Effects of Random Subsampling of PFs

In the main text, we present most results in the fully connected case (*N*_*d*_ = *N*_*m*_), as partial connectivity is never beneficial to SNR when all other parameters are held fixed. We verified this claim analytically in the case of homogeneous plasticity learning in section S4.2. Here, we provide an analytical argument that randomly subsampled connectivity is never beneficial, provided plasticity schedules 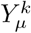 are held fixed across subsampling fractions, and labels are assigned using the optimal decoder Ω_opt_(***g***) (given by the likelihood-ratio/Neyman–Pearson test). We then verify numerically that ensemble SNR increases monotonically with *N*_*d*_ under heterogeneous plasticity when PC activity ***g*** is decoded using the saddle approximation to the optimal decoder. Finally, we study how ensemble SNR varies when *N*_*d*_ and *K* are varied together such that the total synapse count *KN*_*d*_ remains fixed, showing that the trade-off between PC size *N*_*d*_ and ensemble size *K* is dictated by the sparsity *f* of PF activity, with realistic scales *N*_*d*_ ≈ 200, 000 optimizing SNR at optimal PF sparsity *f* ≈ 10^−3^.

### S10.1 Equivalence Between Random Subsampling and Intrinsic Noise under Fixed Plasticity Schedules

We begin by examining the statistics of the learned PC inputs in the case where plasticity signals 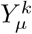 are left in general form, therefore covering both the homogeneous and heterogeneous plasticity settings. For two PCs, we have the following statistics:

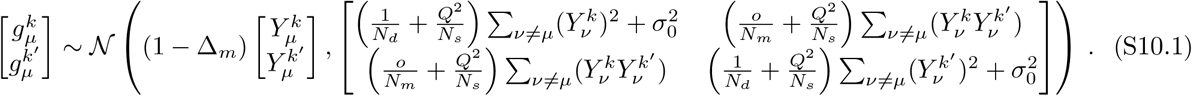

Holding the plasticity signals *Y* and the overlap fraction *o* fixed, we can now examine the effect of subsampling by decreasing *N*_*d*_. We see that this increases the diagonal components of the cross-talk variance, but has no effect on the off-diagonal components of the cross-talk variance. Thus, at fixed *Y* and fixed *o*, decreasing 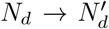 with 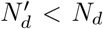 is equivalent in distribution to taking the original PC responses at size *N*_*d*_ and adding an independent Gaussian perturbation to each component of ***g***. We may then make the following statement:

#### Lemma 5.

*For a PC ensemble trained under the general two-factor learning rule (eq. *S2*.3) with a fixed plasticity schedule* 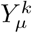 *and fixed overlap fraction o, and when stimuli are detected from PC responses* ***g*** *using the Neyman-Pearson test* Ω_opt_(***g***), *the* AUC *is monotonically increasing with PC size N*_*d*_.

This is a direct application of the “irrelevance theorem” of optimal detection (see page 220 of ref. [97]), which states that the ROC curve achieved by an optimal detector can never improve from the addition of uninformative noise. An important caveat to this argument is that we have assumed *fixed plasticity signals* 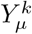 when comparing across different *N*_*d*_ values. Experiments typically describe plasticity in Purkinje cells as driven primarily by the conjunction of parallel-fiber and climbing-fiber activity, rather than by the PCs own firing state prior to complex-spike induction. However, for learning rules which incorporate feedback from PC output into plasticity updates, random subsampling of PFs may implicitly promote heterogeneity in plasticity signals, improving network performance. This effect may explain the benefits to sparse PF–PC connectivity demonstrated in a concurrent work’s simulations of a similar circuit model including PC-specific error feedback signals [98].

Another caveat to this argument is that we have used the PC statistics from eq. S3.24, which were derived in the limit of large *N*_*s*_ and *N*_*d*_, as our starting point. It is possible that sub-leading effects may cause violations of this lemma in small networks.

### S10.2 Varying *N*_*d*_ With Fixed PC Count *K*

Here, we perform Monte Carlo simulations of both homogeneous plasticity and heterogeneous plasticity learning in ensembles of fixed size *K*, varying PC size *N*_*d*_ up to the total PF population size *N*_*m*_ = 400,000. For homogeneous plasticity simulations, we decode using the (optimal in this case) linear decoder, and in the heterogeneous plasticity simulations we decode using the saddle approximation to the optimal decoder, Ω_*rp*_(***g***). The results, shown in fig. S11 (see caption for parameters), verify that SNR increases with *N*_*d*_ across PF sparsity values *f*. In the case of homogeneous plasticity, SNR tends to saturate at relatively small *N*_*d*_ values (around ~ 50,000 or less), which is far smaller than the typical number of ~ 200,000 PF synapses onto a PC. In the case of heterogeneous plasticity, SNR improvement continues even at large *N*_*d*_ values, particularly for very sparse PF activity.

### S10.3 Synapse-Constrained Comparison

Next, we consider the resource-constrained setting in which the total number of PF synapses per ensemble

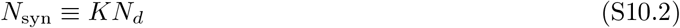

is held constant while *K* and *N*_*d*_ vary. In this regime, increasing *N*_*d*_ induces competing effects, increasing the information available to each PC, but reducing the total number of PCs. Specifically, we fix a total synaptic budget of *N*_syn_ = 8,000,000 and vary *K* and *N*_*d*_ together over the range (*K* = 20, *N*_*d*_ = 400,000) to (*K* = 200, *N*_*d*_ = 40,000). Fig. S12B and C show SNR as a function of *N*_*d*_ in this sweep, after training with both homogeneous (top row) and heterogeneous (bottom row) plasticity, and with varying intrinsic noise scale *σ*_0_ indicated above each subplot. Color bar indicates PF sparsity fraction *f*. In fig. S12B, we again use the optimal linear decoder for homogeneous plasticity. We find that at low noise levels *σ*_0_, SNR is primarily determined by the total number of PFs which are sampled by the PCs. When *o* = 1, we expect approximately *N*_*m*_(1 − *N*_*d*_*/N*_*m*_)^*K*^ PFs will be sampled by none of the PCs. At high sparsity fractions *f* (denser PF activity) this has a negligible effect as PC performance saturates with fewer PFs, but at low sparsity values closer to the optimum this effect encourages a smaller number of larger PCs. At large *σ*_0_, the benefit of noise-reduction gained by averaging over a large number of PCs dominates, and the optimal (*N*_*d*_, *K*) pairing shifts toward a larger number of smaller PCs.

For heterogeneous plasticity simulations (panel C) we use the saddle approximation Ω_*rp*_ in heterogeneous plasticity simulations. Fig. S12C shows that under heterogeneous plasticity, the optimal *N*_*d*_ depends strongly on mixed-layer sparsity *f* and the scale *σ*_0_ of intrinsic noise. With very sparse PF activity (low *f*) and low noise values *σ*_0_ = 0.0 or 0.1, the optimal *N*_*d*_ approaches realistic values on the order 200,000. However, with denser representations (larger *f*) optimal PC size is often far below realistic scales. In figs. S12D,E we aggregate SNR curves at the optimal sparsity *f* for each *N*_*d*_. Under heterogeneous plasticity (fig. S12E) we see the optimal PC size *N*_*d*_ falls at realistic scales of *N*_*d*_ ≈ 200, 000, provided intrinsic noise is limited to relatively small scales. This suggests that the trade-off between PC size and PC ensemble size is not governed by noise reduction, but rather governed by the trade-off between individual PC reliability increasing with *N*_*d*_, and decoder network expressivity increasing with *K*. An important caveat, however, is that this requires extremely sparse representations in PFs, on the order *f* ≈ 10^−3^. Some recent experimental results suggest that PF activity may not be so sparse [62, 61]. Others attribute these estimates to low-dimensional task structure, finding high-dimensional activity in recordings from PFs [63]. Optimal sparsity of PF activity may also be affected by alternative mechanisms unaccounted for in our model, such as the prevalence of silent synapses from PFs to PCs [99]. Alternatively, densely activated PF inputs may drive the PC dendritic tree into a highly nonlinear dendritic spiking regime, so that each PC individually behaves as an ensemble of smaller dendritic compartments [67], increasing the effective ensemble size and decreasing the effective *N*_*d*_ value in a manner which is adaptive to the sparsity of PF inputs. Additional modeling efforts are required to address these possibilities.

**Figure S11:**
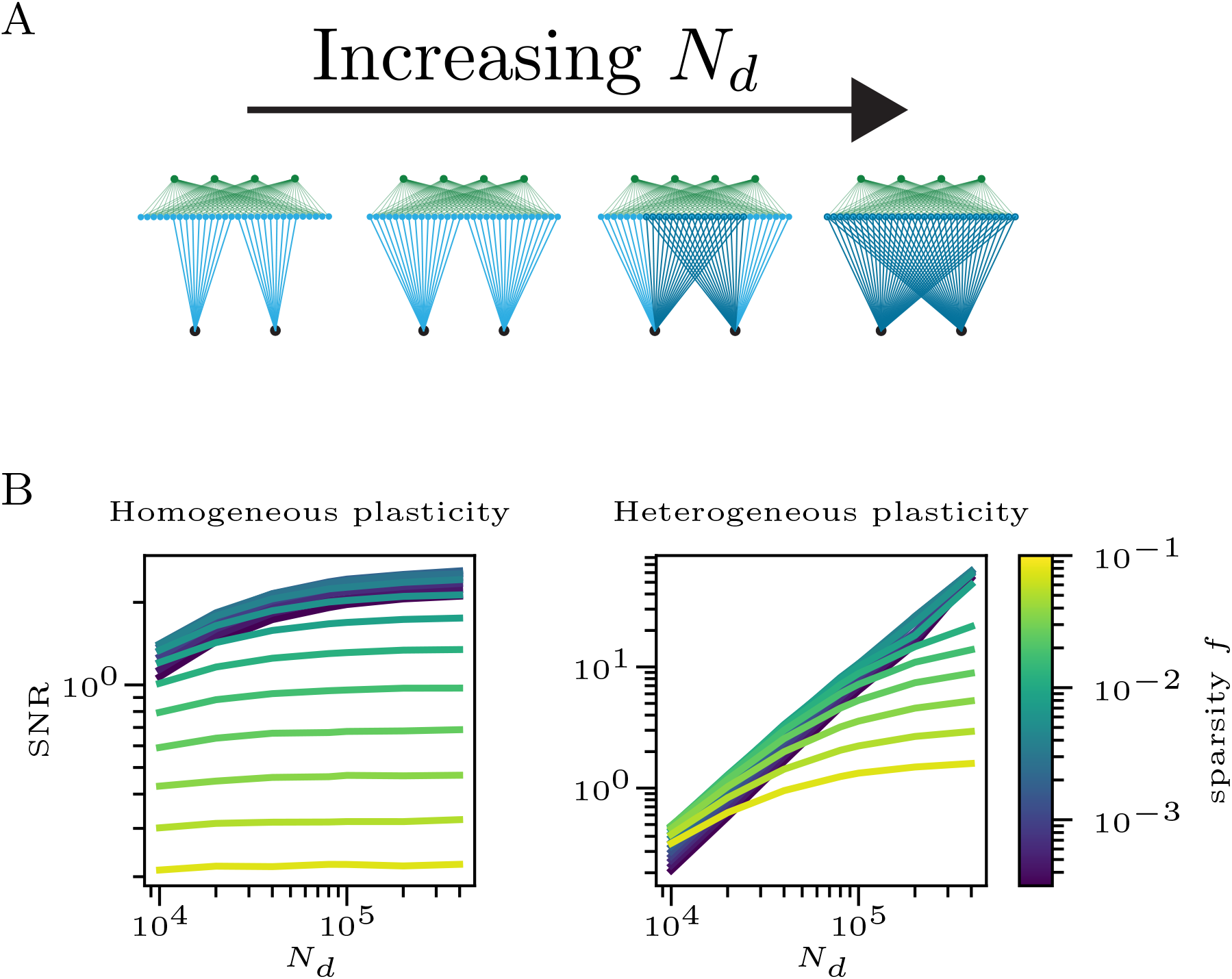
Increasing PC input size at fixed ensemble size. **(A)** Schematic of the fixed-*K* comparison: ensemble size is held constant at *K* = 40 while the number of parallel-fiber inputs per PC (*N*_*d*_) is varied from 10,000 to *N*_*m*_. **(B)** Ensemble SNR versus *N*_*d*_ for homogeneous (left) and heterogeneous (right) plasticity. Curves are colored by granule-cell sparsity *f*. Under homogeneous plasticity, SNR saturates at relatively small *N*_*d*_; under heterogeneous plasticity, SNR continues to improve up to large *N*_*d*_, particularly at low *f*. Simulations used *P* = 15,000, *N*_*s*_ = 1,400, *N*_*m*_ = 400,000, Δ_*x*_ = 0.10, intrinsic noise *σ*_0_ = 0.1, overlap fraction *o* = 1, and plasticity parameters (*β, η*_+_, *η*_−_) = (1, 0, 1) (homogeneous) and (0.78, 1.57, 0.76) (heterogeneous). Shaded regions indicate bootstrap 95% confidence intervals across trials.

## S11 Centered Weight Updates as a Combination of Instructed and Homeostatic Plasticity

We consider learning to occur in two steps upon presentation of a stimulus intended for memorization. Consider 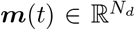 to be PF activity at the inputs to a PC of size *N*_*d*_ at time *t*. Averaging over time, we have 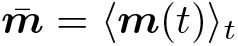. At any given moment in time, the instantaneous spiking rate of the Purkinje cell is determined by its weight vector ***ŵ***:

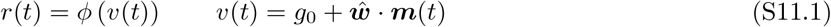

The baseline membrane potential has the form

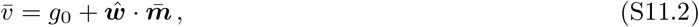

and we seek an update rule for ***ŵ*** which maintains this value of 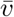. This may be achieved in two steps. Upon presentation of a stimulus ***x***(*t*_*µ*_) with an associated error, the Purkinje cell will receive instantaneous PF inputs ***m***(*t*_*µ*_). We will assume that the patterns ***m***(*t*_*µ*_) can be written as 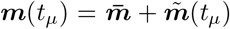, where the components of 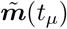 are independent random variables with mean zero. From the central limit theorem, it follows that

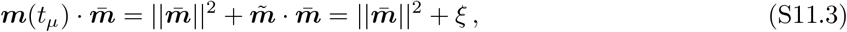

where 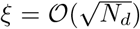 is vanishingly small relative to 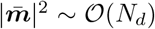 for large *N*_*d*_. In the instructed plasticity step, the plasticity signal *Y*_*µ*_ provides a hebbian weight update *ŵ* → *ŵ′ = ŵ + Y*_*μ*_*m*(*t*_*μ*_). Immediately following this weight update, we will have

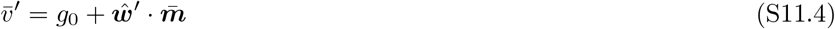

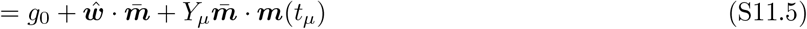

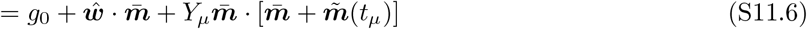

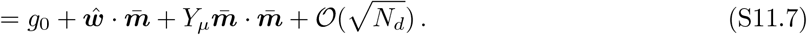

In the second step, suppose a neuron undergoes a slower homeostatic plasticity process which provides a weight update of magnitude 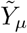 in the direction of the baseline PF activity 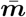, so that 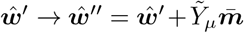. Following the slow homeostatic plasticity step, we will have (at leading order in *N*_*m*_)

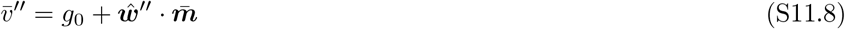

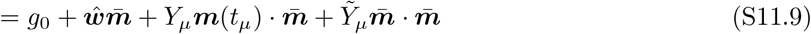

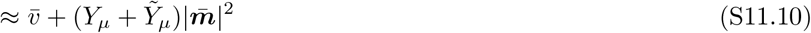

To maintain the baseline membrane potential of 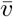 to leading order in *N*_*d*_, we require a homeostatic plasticity signal 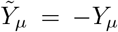. With this choice, the combined effect of the immediate and subsequent homeostatic plasticity will achieve the following net update:

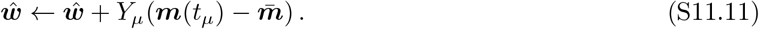

**Figure S12:**
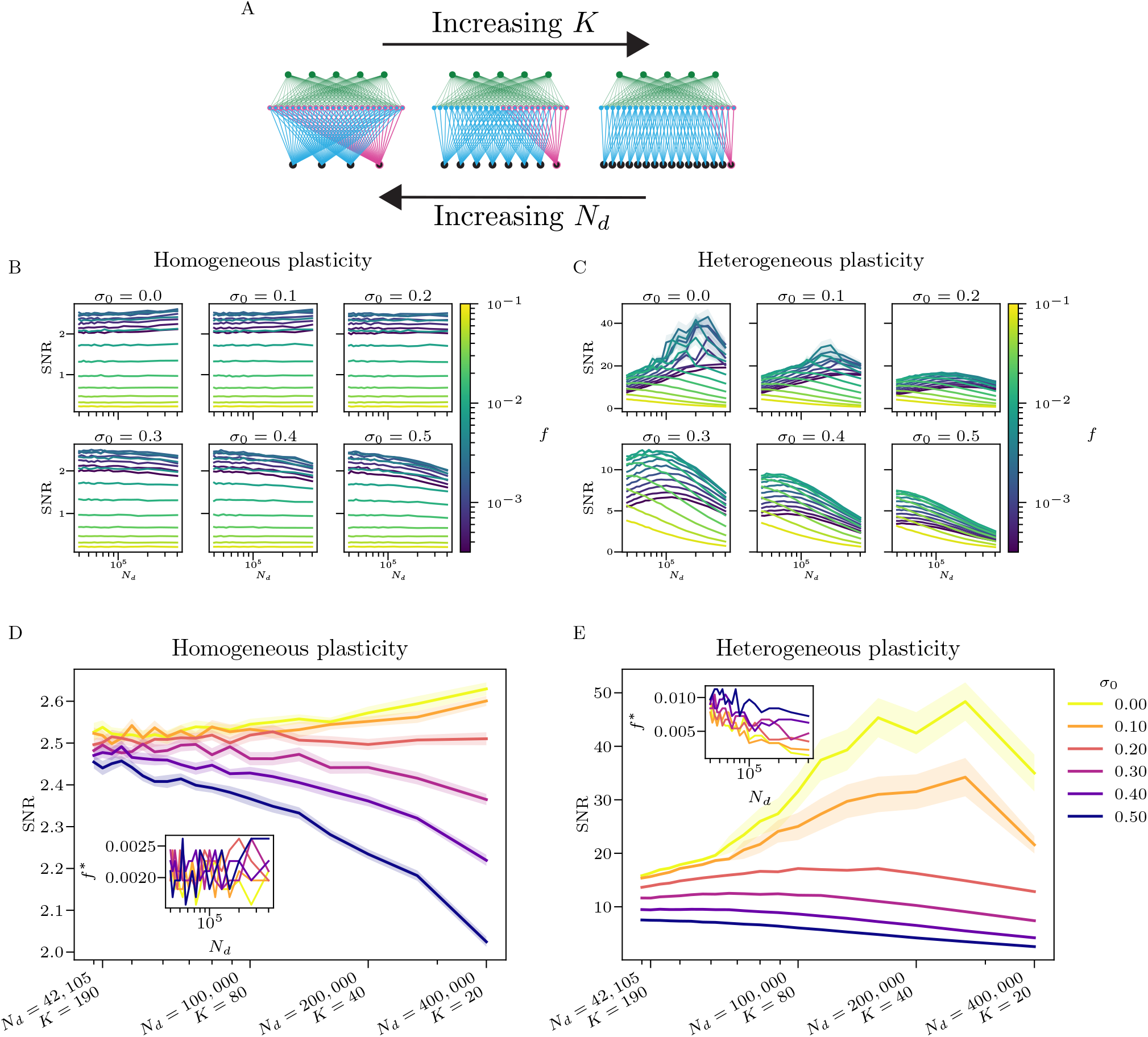
Synapse-constrained trade-off between PC size and ensemble size. **(A)** Schematic of the fixed-budget comparison: total synapse count *N*_syn_ = *KN*_*d*_ is held constant while *K* and *N*_*d*_ vary inversely. **(B, C)** Ensemble SNR versus *N*_*d*_ for homogeneous (B) and heterogeneous (C) plasticity across intrinsic-noise levels *σ*_0_ ∈ {0, 0.1, 0.2, 0.3, 0.4, 0.5}. Curves are colored by granule-cell sparsity *f*. **(D, E)** Optimal-*f* envelopes of SNR versus *N*_*d*_ for homogeneous (D) and heterogeneous (E) plasticity. Insets show the corresponding optimizing sparsity *f*^∗^(*N*_*d*_). Simulations used *P* = 15,000, *N*_*s*_ = 1,400, *N*_*m*_ = 400,000, Δ_*x*_ = 0.10, overlap fraction *o* = 1, fixed budget *N*_syn_ = 8 × 10^6^, and plasticity parameters (*β, η*_+_, *η*_−_) = (1, 0, 1) (homogeneous) and (0.78, 1.57, 0.76) (heterogeneous). Shaded regions indicate bootstrap 95% confidence intervals across trials.

Substituting *Y*_*µ*_ ← *Y*_*µ*_*/*(*N*_*d*_*f* (1 − *f*)) we arrive at the centered update rule used in the main text.

## S12 Rigorous AUC and Capacity Bounds under Heterogeneous Plasticity

In this section we provide a more rigorous treatment of network performance under heterogeneous plasticity learning, verifying the scaling dependence of network capacity *P*_max_ on ensemble size *K* and the effective GrC activity dimensionality *N*_eff_ presented in main text and §S5. This section is somewhat auxiliary to the remainder of the paper, and can be safely skipped by any reader who is satisfied with the level of rigor provided by the physics-style arguments given in §S5. We used the help of AI chatbots Claude^1^ and ChatGPT^2^ for some intermediate steps in the proofs in this section, all of which have been verified by the authors.

We take as our starting point the post-training statistics of PC inputs ***g*** derived in §S3.1. For simplicity, we again specialize to fully connected PCs with *N* ^*k*^ = *N*_*m*_ for all *k* = 1, …, *K*, as in §S5.1; the corresponding covariance matrices are given by eq. S5.2. For clarity, we reproduce the statistics of ***g***_*n*_ and ***g***_*ℓ*_ here:

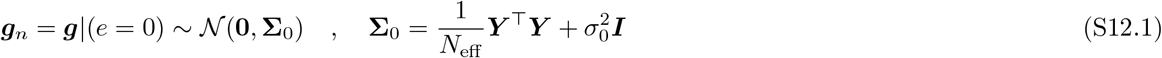

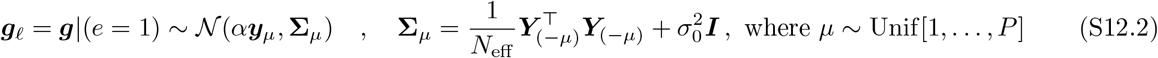

and where we have set *α* = 1 − Δ_*m*_ ∈ (0, 1], ***y***_*µ*_ corresponds to row *µ* of ***Y***, and ***Y***_(−*µ*)_ is the matrix ***Y*** with row *µ* removed.

Because the matrix ***Y*** is randomly sampled as 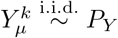 during heterogeneous plasticity learning, network performance fluctuates across realizations of the learning algorithm. These trial-to-trial fluctuations appear to be small (see main-text fig. 3), so the disorder-averaged AUC and capacity estimates derived in §S5 accurately describe the typical performance of a trained network. In this section, we provide a more rigorous treatment of network AUC and capacity which explicitly accounts for the trial-to-trial fluctuations in ***Y***. Specifically, we provide rigorous backing for the following claims (stated informally here):

1. For a network with linear decoding (i.e. Ω(***g***) ∝ ∑_*k*_ *g*^*k*^), homogeneous plasticity learning outperforms heterogeneous plasticity learning, so that the optimal operating point is (*β* = 0, Φ_*Y*_ = 1) or (*β* = 1, Φ_*Y*_ = −1). It follows that no network with a linear decoder can outperform the direct readout architecture. The analogous non-rigorous result is derived in §S5.3 from the approximate SNR in eq. S5.48.
2. For a network with quadratic decoder Ω(***g***) = ∑_*k*_(*g*^*k*^)^2^ trained under balanced, symmetric plasticity (*β* = 1*/*2, *η*_+_ = *η*_−_ = *η*) and *σ*_0_ = 0, network capacity scales as 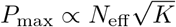 over the biologically relevant range where 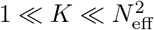. This is the intermediate capacity-scaling regime obtained in the non-rigorous quadratic-decoder analysis in §S5.4 (see eq. S5.99).
3. For a network with pause-based decoder Ω(***g***) = ∑_*k*_ Θ[*κ* − *g*^*k*^] and balanced plasticity (*η*_−_ = (1 − *β*)Λ, *η*_+_ = *β*Λ for some Λ *>* 0), network capacity scales as *P*_max_ ∝ *KN*_eff_ */* log(*K*) when LTD probability scales down as *β* ∝ *c/K*. The analogous non-rigorous scaling argument is given in §S5.5.4, with the target capacity stated in eq. S5.148.

In §S12.1, we prove the first claim directly from the ***g*** statistics for general ***Y*** that AUC for a network with a linear decoder is optimized over all ***Y*** ∈ [−*η*_−_, *η*_+_]^*P ×K*^ by either setting 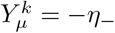 for all *µ, k* or 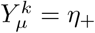 for all *µ, k*.

We treat the second and third statements in §S12.2 and §S12.3, respectively. There, we show that for a fixed error tolerance *ϵ* ∈ (0, 1), for some function 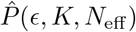 with the prescribed scalings and some small, decreasing function *δ*(*K, P*), a tail bound of the form

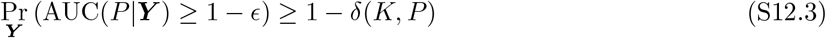

holds for all *P* such that 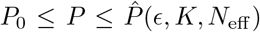. In words, this inequality guarantees that on some high fraction 1 − *δ* of trials, network capacity is no less than 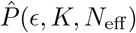. The minimum *P*_0_ is typically some small value below which the concentration arguments of the proof break down. Furthermore, the failure probability *δ*(*K, P*) decreases with system size, so that as the network grows the probability of the network failing to meet the capacity bound on a given trial, due to a rare but pathological realization of ***Y***, vanishes as *δ*(*K, P*). In the case of the quadratic decoder, we will have *δ*(*K, P*) = *e*^−*γ* min{*K,P*}^ and for the pause-decoder we will have *δ*(*K, P*) = 2*K*^−*γ*^ for some *γ, c >* 0.

### Caveat: Scaling with *N*_eff_

Before proceeding to the proofs, we give the caveat that the ***g***_*n*_ and ***g***_*ℓ*_ statistics (eqs. S12.1, S12.2) which we take as our starting point are themselves asymptotics derived under the assumption that *N*_*s*_, *N*_*d*_ ≫ 1 and *N*_*d*_*f* (1 − *f*) ≫ 1. The statements that follow may then be considered rigorous statements about these asymptotically sharp statistics of ***g***_*n*_ and ***g***_*ℓ*_. In the fully-connected case where *N*_*d*_ = *N*_*m*_, the statistics of ***g*** depend on *N*_*s*_ and *N*_*m*_ only through the combined quantity 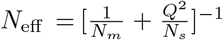. We therefore state the capacity scaling results in terms of *N*_eff_ directly. It is clear that at sufficiently small *N*_*m*_, we will have *N*_eff_ ≈ *N*_*m*_. However, at large *N*_*m*_, we will have *N*_eff_ ≈ *Q*^−2^*N*_*s*_ ∝ *N*_*s*_.

### S12.1 Linear Decoding

Here, taking eqs. S12.1 and S12.2 as our starting point, we prove that under linear decoding, homogeneous plasticity always outperforms heterogeneous plasticity learning. This provides rigorous backing for the claim in §S5.3, which was based on the non-rigorous derivation of SNR for the linear decoder.

#### Theorem 1.

*Suppose PC input statistics are given by eq’s S12*.*1, S12*.*2, and we use linear decoder* 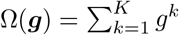. *Suppose also that* ***Y*** ∈ [−*η*_−_, *η*_+_]^*P ×K*^. *Then* AUC *is maximized by setting* 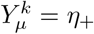 *for all entries µ, k*.

*Proof*. We begin by writing the AUC formula under linear decoding for general ***Y*** matrices.

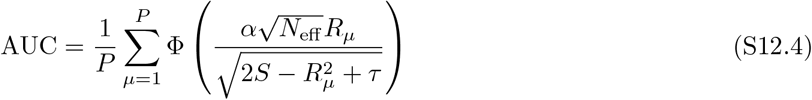

where we have defined the following:

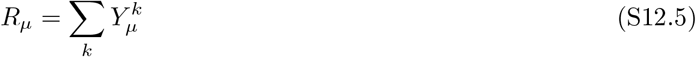

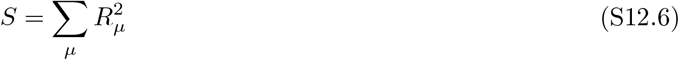

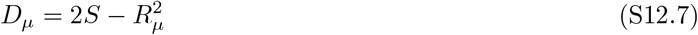

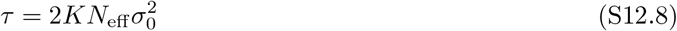

We note that AUC depends on the plasticity matrix ***Y*** only through the row-sums *R*_*µ*_. We may therefore reduce the problem to optimization over the *R*_*µ*_ ∈ [*R*_−_, *R*_+_], where *R*_−_ = −*Kη*_−_, and *R*_+_ = *Kη*_+_, and it suffices to show that AUC is optimized by setting *R*_*µ*_ = *R*_+_ for all *µ*. To show this, we will provide a series of transformations of the *R*_*µ*_ under which AUC is non-decreasing, and which terminates at the desired configuration.

The first transformation is to set all *R*_*ν*_ *<* 0 to 0. Suppose *R*_*ν*_ *<* 0 for all *ν* ∈ *N* ⊆ {1, …, *P*}. Replacing *R*_*ν*_ → 0 in eq. S12.4 for all *ν* ∈ *N* simultaneously, the denominator in the argument of Φ decreases for every *µ*, all positive numerators do not change, and all negative numerators become zero. Therefore every term in the sum is non-decreasing under the transformation.

Now we have *R*_*µ*_ ≥ 0 for all *µ* = 1, …, *P*. We then iterate the following transformation until *M* = {1, …, *P*}: let *r* = min_*µ*_{*R*_*µ*_} and define the set *M* = {*µ* : *R*_*µ*_ = *r*}. Letting *q* = min_*µ*∈*/M*_ {*R*_*µ*_}, put

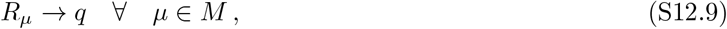

which raises all minimal of *R*_*µ*_ from r to q. To show that this “leveling” transformation (S12.9) cannot decrease AUC, it is enough to consider the continuous path

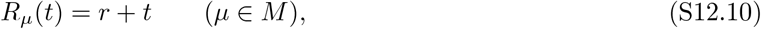

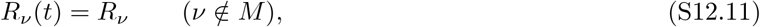

for 0 ≤ *t* ≤ *q* − *r*. Along this path, the entries in *M* remain minimal. Let *ρ* = *r* + *t, m* = |*M* |, and

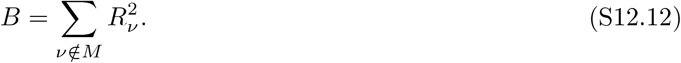

Then

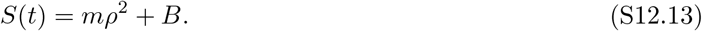

For *µ* ∈ *M*, all terms have the same argument of Φ, which we denote by

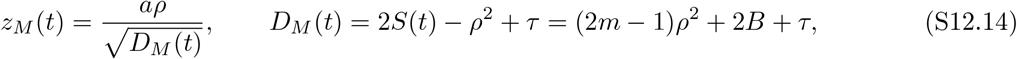

where 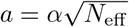. For *ν* ∈*/M*, write

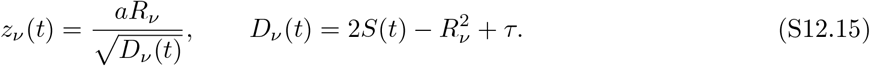

Differentiating gives

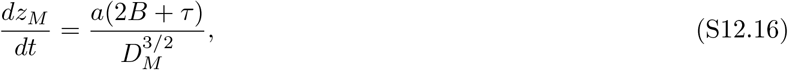

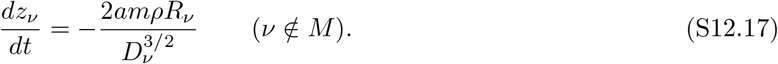

Therefore

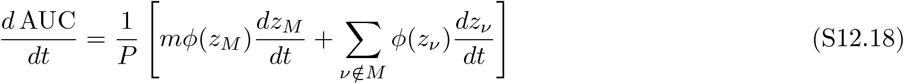

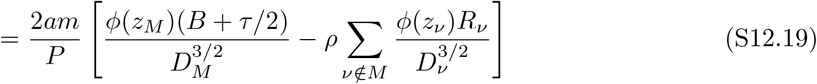

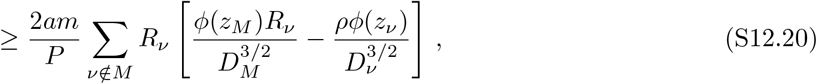

where in the last line we have used 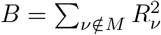 and *τ/*2 ≥ 0. To show that this derivative is nonnegative, therefore suffices to prove

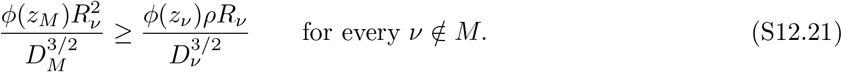

Because the entries in *M* remain minimal along the path, *R*_*ν*_ ≥ *ρ*. Hence

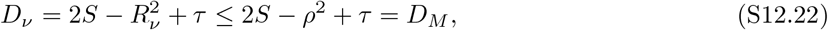

and therefore

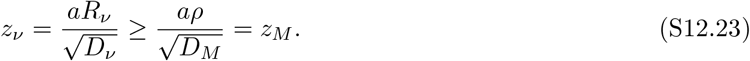

Since *z*_*M*_, *z*_*ν*_ ≥ 0 and *ϕ* is decreasing on [0, ∞), we have

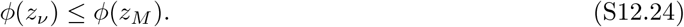

It remains only to check that

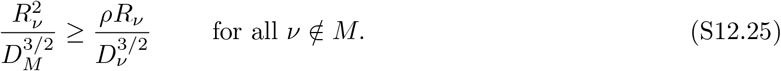

If *ρ* = 0, this is immediate. Otherwise, set *x* = *R*_*ν*_*/ρ* ≥ 1 and write

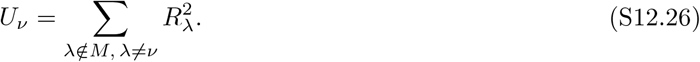

Then

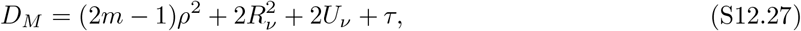

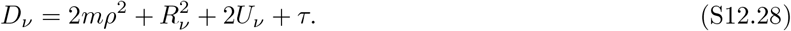

Writing *q*_*ν*_ = (2*U*_*ν*_ + *τ*)*/ρ*^2^ ≥ 0, we have

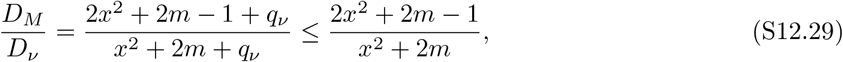

where the inequality follows because the ratio is decreasing in *q*_*ν*_ for *x* ≥ 1 and *q*_*ν*_ ≥ 0. Thus it suffices to show

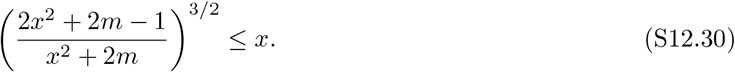

Equivalently, with *y* = *x*^2^ ≥ 1, we need

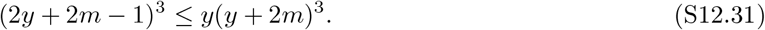

This holds for all *m* ≥ 1 and *y* ≥ 1 due to the identity

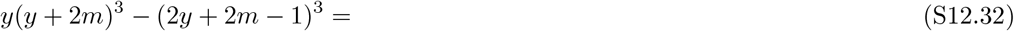

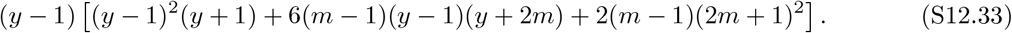

Since *y* ≥ 1 and *m* ≥ 1, every factor on the right-hand side is nonnegative, so that (S12.31) holds. Therefore (S12.21) holds, and hence

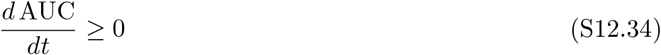

throughout the leveling path. Integrating from *t* = 0 to *t* = *q* − *r* proves that replacing all minimal entries *R*_*µ*_ = *r* by *q* cannot decrease AUC. Iterating the leveling transformation on the set of minimum *R*_*µ*_, we will arrive at *M* = {1, …, *P*} in no more than *P* − 1 steps. With all *R*_*µ*_ = *R* equal, we may rewrite the AUC in simplified form as:

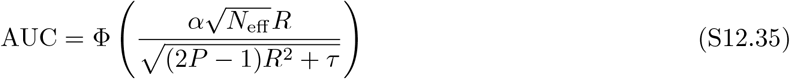

Because *τ* ≥ 0, it follows that AUC is non-decreasing with *R*, so that it is optimized by setting *R* to its maximum value of *R*_+_.

By reflection symmetry, a simple corollary of theorem 1 is that when Ω(***g***) = − *∑*_*k*_ *g*^*k*^ (and assuming −*η*_−_ ≤ 0), the choice 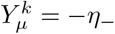 for all *µ, k* optimizes AUC, with AUC given by eq. S12.35 with *R* = *K*|*η*_−_|. It follows that when we are given the flexibility to choose between Ω(***g***) = *± ∑*_*k*_ *g*^*k*^, AUC is optimized by setting 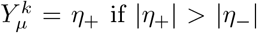 or 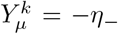 otherwise. It follows that homogeneous plasticity is optimal under linear decoding. Because eq. S12.35 is decreasing with *τ*, and equal to the AUC of the direct readout architecture when *τ* = 0, it also follows that the direct readout AUC is an upper bound on the AUC of any network with linear decoder at fixed *P*. Because this holds for all *P*, it also follows that the capacity of a network with linear decoder is upper bounded by the direct readout capacity.

#### A Useful AUC Lower Bound

Before proceeding to our treatment of the quadratic and pause-based decoders in the following two sections, we note the following lower bound on AUC. From the the ordering definition of AUC, we have that 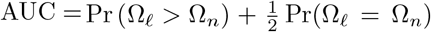 for an Ω_*ℓ*_ and Ω_*n*_ drawn independently from the output distributions for learned and nove linputs, respectively. If we select a threshold *T*, a sufficient condition to achieve Ω_*ℓ*_ *>* Ω_*n*_ is to have Ω_*ℓ*_ *> T* and Ω_*n*_ *< T*. We then have the following inequalities:

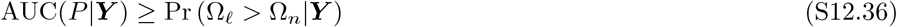

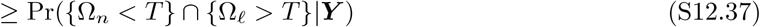

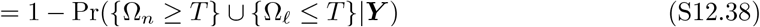

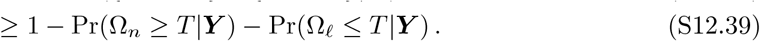

The first line follows from the trivial bound Pr(Ω_*ℓ*_ = Ω_*n*_) ≥ 0, the second from the sufficient condition explained above, the third from De Morgan’s law, and the fourth from the union bound. We therefore see that we may bound AUC by placing upper bounds on the rate of false-poisitives (Ω_*n*_ ≥ *T*) and false-negatives (Ω_*ℓ*_ ≤ *T*) for any threshold *T*. Finally, in the case where *µ* ~ Unif(1, …, *P*) under the learned stimulus condition, we may write

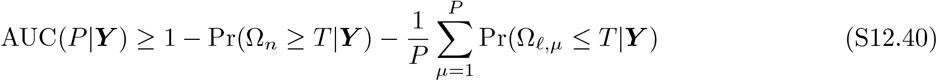

### S12.2 Quadratic Decoding (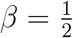, *κ* = 0, *σ*_0_ = 0)

We give a rigorous high-probability lower bound on the AUC, and a corresponding capacity lower bound, for the balanced heterogeneous-plasticity rule paired with quadratic decoding, in the special case 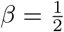, *κ* = 0, and vanishing readout noise *σ*_0_ = 0, so that 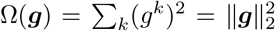. In this section, we use ⪯ for PSD matrices ***A*** and ***B*** to denote Loewner order, so that ***A*** ⪯ ***B*** if and only if ***B*** − ***A*** is PSD.

Let *α* = (1 − Δ_*m*_), *P* ≥ 2, *K* ≥ 1, *η >* 0, and *N*_eff_ *> α*^−2^ *>* 0. Let 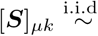 Rademacher for *µ* = 1, …, *P* and *k* = 1, …, *K*, let ***s***_*µ*_ ∈ {− 1, +1}^*K*^ be row *µ* of ***S***, let ***S***_−*µ*_ be ***S*** with row *µ* removed, and set ***Y*** = *η****S***. Writing *α* ≡ 1 − Δ_*m*_ and specializing to *N*_*d*_ = *N*_*m*_, *σ*_0_ = 0, the novel and leave-one-out learned covariances are

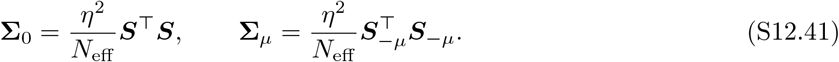

The fixed-***Y*** response distributions are

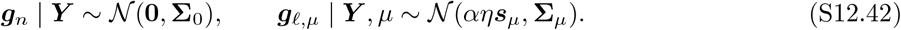

For any nonzero positive semidefinite matrix ***A***, write

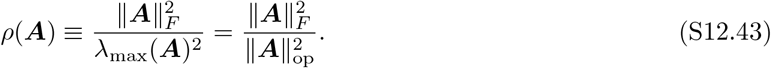

This effective rank is scale-invariant, so *ρ*(*c****A***) = *ρ*(***A***) for every *c >* 0.

Define the mean scores and variance proxies

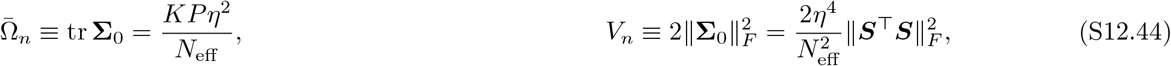

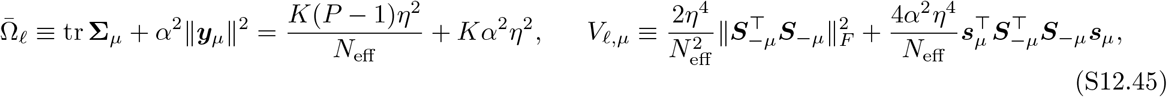

where ***y***_*µ*_ = *η****s***_*µ*_. With the symmetric Rademacher law of ***Y***, both 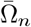 and 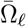 are deterministic and *µ*-independent.

#### Lemma 6

(Upper tail for a weighted central Gaussian norm). *Let* ***ϵ*** ~ *N*(**0, Σ**) *with* **Σ** ⪰ 0, *and let* 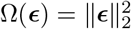, *then for every t* ≥ 0,

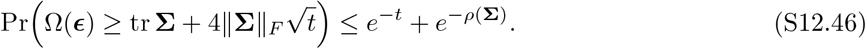

*Proof*. By the upper tail bound in Lemma 1 of Laurent and Massart [100],

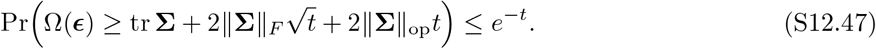

Partition on whether *t* ≤ *ρ*(**Σ**) or *t* ≥ *ρ*(**Σ**). If *t* ≤ *ρ*(**Σ**), then

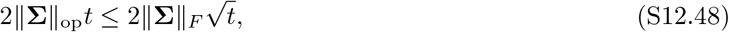

so tr 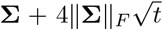 dominates the Laurent–Massart threshold and the tail is at most *e*^−*t*^. If *t* ≥ *ρ*(**Σ**), monotonicity of the tail gives

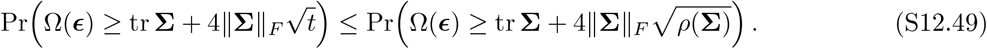

Applying the first case at *t* = *ρ*(**Σ**) bounds this last probability by *e*^−*ρ*(**Σ**)^. Combining the two cases gives the stated bound.

#### Lemma 7

(Lower tail for a weighted noncentral Gaussian norm). *Let* ***ϵ*** ~ *N*(**0, Σ**) *with* **Σ**≠ **0, Σ** ⪰ 0 *and let* ***m*** ∈ ℝ^*K*^ *be fixed. Put* 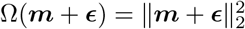,

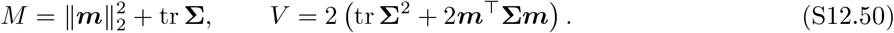

*Then, for every t* ≥ 0,

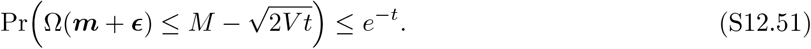

*Proof*. Proof of the special case where ***m*** = 0 is given by the lower tail bound in Lemma 1 of Laurent and Massart [100]. For completeness, we reproduce their argument here with modifications to account for ***m***≠ 0. Diagonalize **Σ** = ***U*** diag(*λ*_*i*_)***U*** ^⊤^ and write 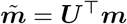. Then

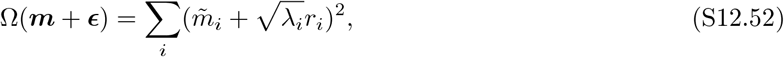

where 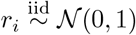, and

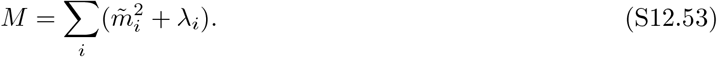

Therefore

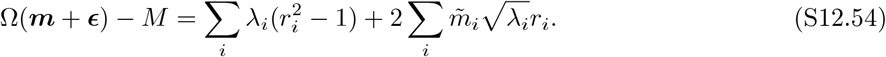

For *u* ≥ 0,

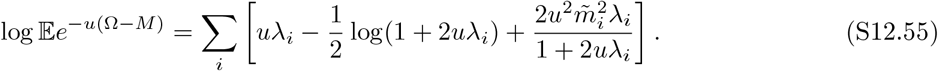

Using 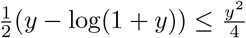 with *y* = 2*uλ*, and using (1 + 2*uλ*_*i*_)^−1^ ≤ 1, gives

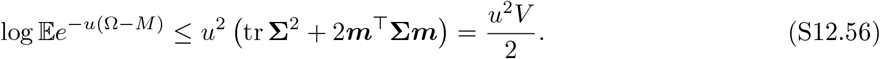

Chernoff’s bound gives, for every *s* ≥ 0,

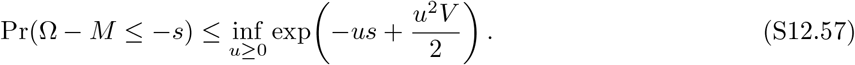

Choosing *u* = *s/V* gives

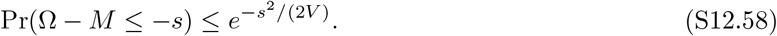

Taking 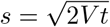 completes the proof.

#### Lemma 8

(Full-Gram spectral control and deterministic noncentral control). *There exists a constant C >* 0 *such that the following holds. Fix a failure level δ* ∈ (0, 1) *and set*

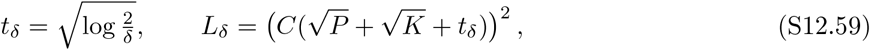

*Then there exists an event G*_*S*_ *with* Pr_***S***_(*G*_*S*_) ≥ 1 − *δ on which*

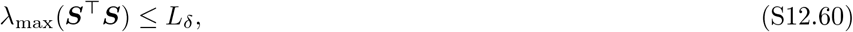

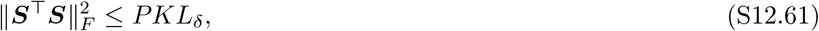

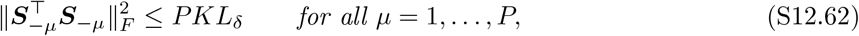

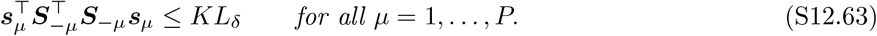

*Moreover, the deterministic lower bounds*

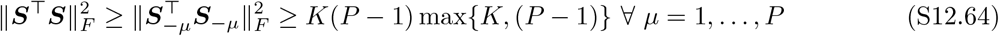

*hold for every realization of* ***S***.

*Proof*. Define

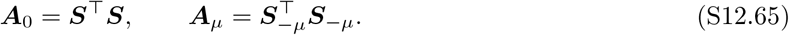

To prove (S12.64), notice first that for all *µ*,

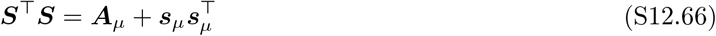

so that 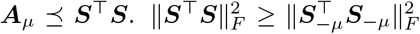 then holds for all *µ* by monotonicity of the Frobenius norm under Loewner order. The *K* diagonal entries of 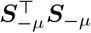 are all equal to *P* − 1, so retaining only the diagonal terms gives

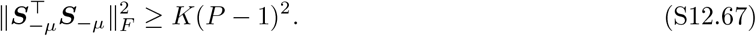

We also have

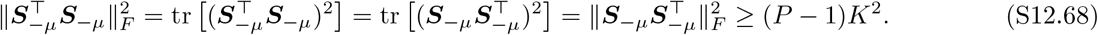

where in the last line we have retained only the diagonal terms. Combining these two deterministic lower bounds gives (S12.64).

Next, ***S*** is a *P* × *K* matrix with independent, mean-zero, sub-Gaussian entries. By Theorem 4.4.3 of **?**], there is an absolute constant *C >* 0 such that, for every *t* ≥ 0,

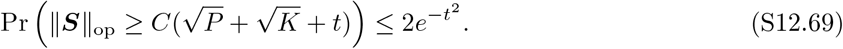

Taking 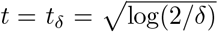 gives failure probability at most *δ*. Therefore, on an event *G*_*S*_ with probability at least 1 − *δ*,

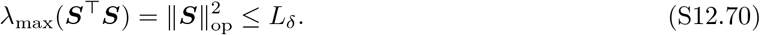

This proves (S12.60).

The Frobenius bound (S12.61) then follows from (S12.60). Writing the singular values of ***S*** as *σ*_1_, …, *σ*_*r*_, with *r* = min{*P, K*},

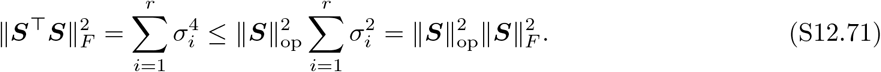

Every entry of ***S*** is *±*1, so 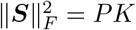. On *G*_*S*_, this gives

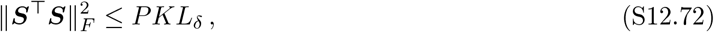

proving (S12.61). Frobenius monotonicity under Loewner order also gives

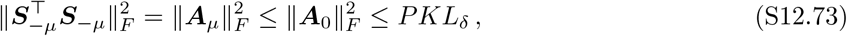

proving (S12.62). Finally, since 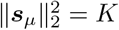 and ***A***_*µ*_ ⪯ ***A***_0_,

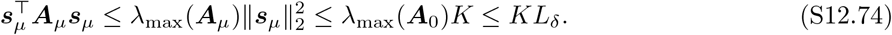

This proves (S12.63) uniformly for all *µ* on the same full-Gram event *G*_*S*_.

#### Theorem 2

(High-probability AUC lower bound). *Fix δ* ∈ (0, 1), *and let t*_*δ*_ *and L*_*δ*_ *be as defined in Lemma 8. Consider the PC statistics under balanced, symmetric plasticity* (S12.41),(S12.42), *assume* 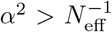, *and define*

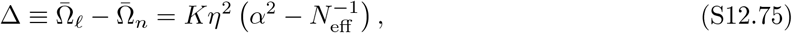

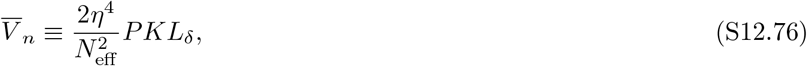

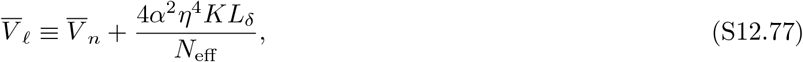

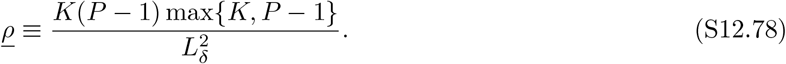

*Then, with probability at least* 1 − *δ over the draw of* ***Y***,

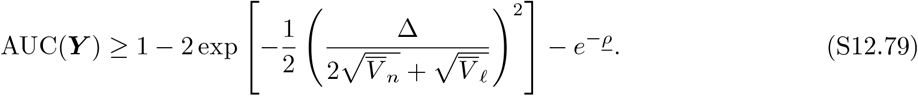

*Proof*. Fix ***Y***, and hence fix ***S***. Following (S12.40), we seek upper bounds on Pr(Ω_*n*_ ≥ *T*) and Pr(Ω_*ℓ,µ*_ ≤ *T*) for a conveneintly chosen threshold *T*. Restrict to 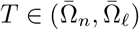 and write

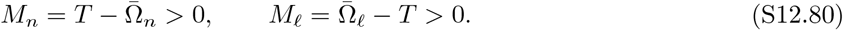

Then *M*_*n*_ + *M*_*ℓ*_ = Δ. Recalling that ***g***_*n*_ ~ *N*(**0, Σ**_0_) and applying Lemma 6 with 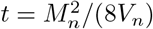 gives

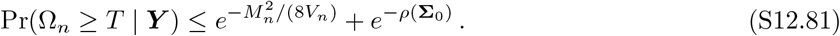

Similarly applying Lemma 7 to ***g***_*ℓ,µ*_ ~ *N*(*α****y***_*µ*_, **Σ**_*µ*_) with 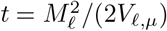 gives

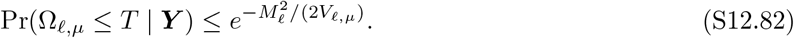

Now restrict to the good event *G*_*S*_ of Lemma 8. On *G*_*S*_, (S12.61) gives

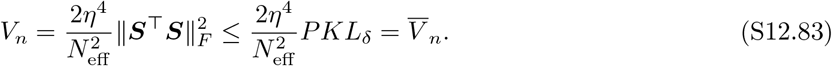

For the learned variance proxy, (S12.62) and (S12.63) give, uniformly for every *µ*,

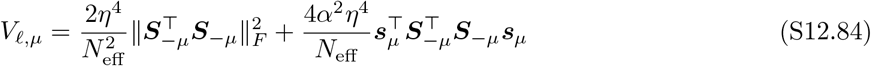

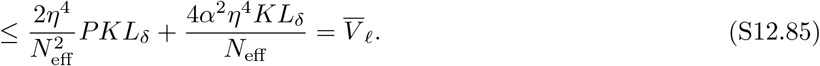

By (S12.64) and (S12.60),

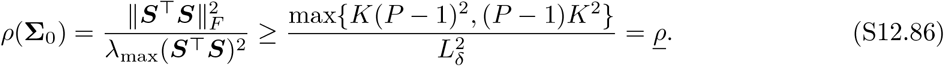

Substituting these deterministic controls into (S12.81) and (S12.82) we obtain, on *G*_*S*_,

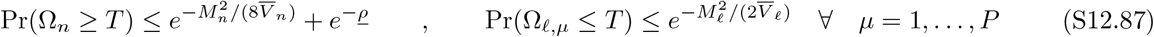

We then select *T* such that

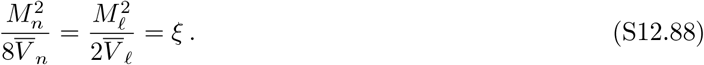

The first equality gives

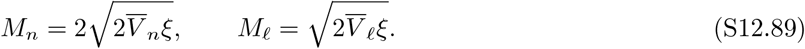

Using *M*_*n*_ + *M*_*ℓ*_ = Δ, we obtain

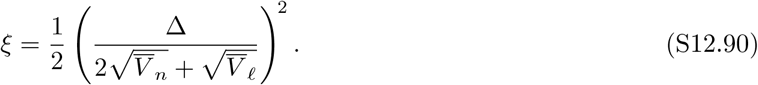

Substituting into (S12.40) and using the fact that (S12.87) is uniform in *µ*, we get

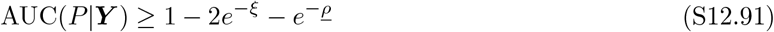

on *G*_*S*_. Finally, Lemma 8 gives Pr_***Y***_ (*G*_*S*_) ≥ 1 − *δ*, completing the proof.

#### Corollary 1

(High-probability capacity lower bound). *Fix a target error ϵ* ∈ (0, 1) *and define* 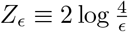. *There exist absolute constants γ, c*_cap_, *c*_0_, *c*_1_ *>* 0 *such that the following holds. Suppose*

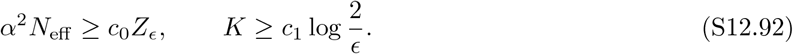

*Then for P satisfying*

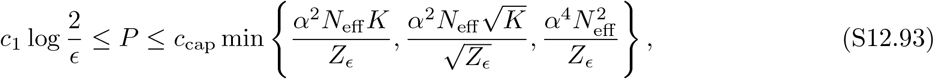

*we have*

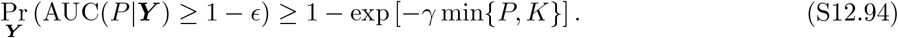

*Proof*. Choose an absolute constant *γ >* 0 and set

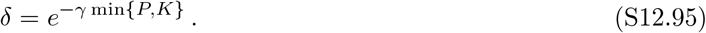

Plugging this choice of *δ* into *t*_*δ*_ from Theorem 2 gives

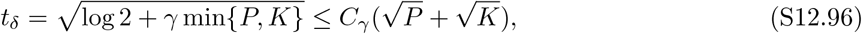

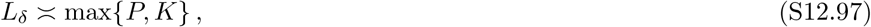

for absolute constant *C*_*γ*_. Also,

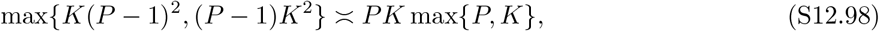

with absolute constants for *P* ≥ 2. Therefore

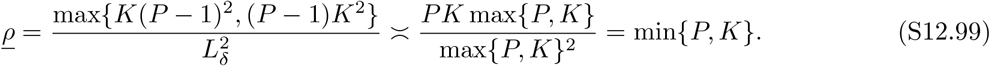

Thus, under the condition min{*P, K*} ≥ *c*_1_ log(2*/ϵ*) with *c*_1_ sufficiently large,

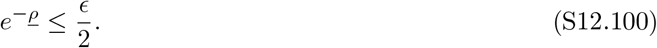

It remains to ensure that the first error term in (S12.79) is at most *ϵ/*2. It is sufficient that

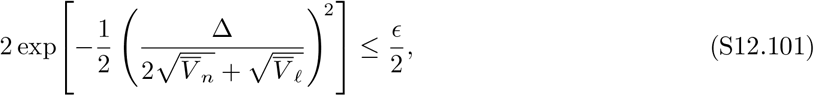

which is equivalent to

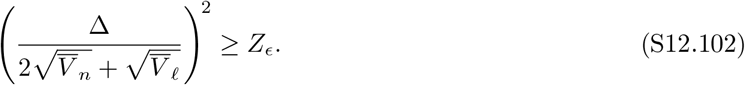

Using Cauchy-Schwarz on the vectors 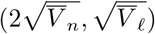 and (1, 1), we get 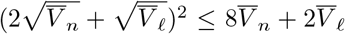. It then suffices to require

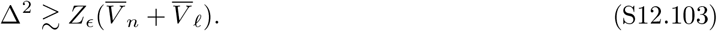

Using *L*_*δ*_ ≍ max{*P, K*}, we also get

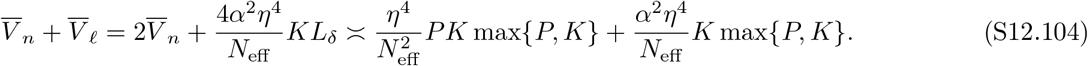

Also, since 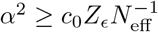, for sufficiently large *c*_0_ we have

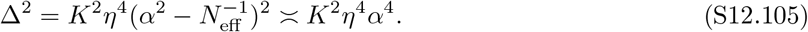

Substituting these scalings into (S12.103) gives the sufficient condition

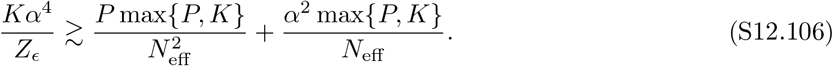

The first term requires

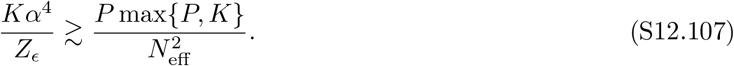

If *P* ≥ *K*, this is implied by

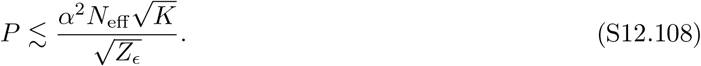

If *K* ≥ *P*, this is implied by

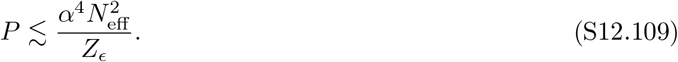

Thus the first term is controlled whenever

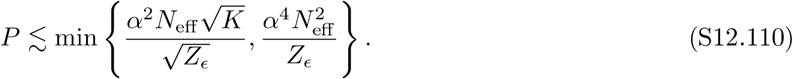

The second term on the right hand side of (S12.106) requires

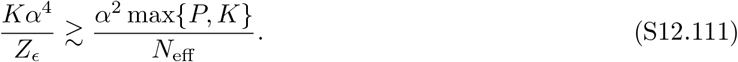

If *P* ≥ *K*, this is implied by

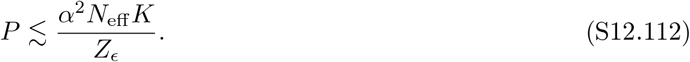

If *K* ≥ *P*, this is implied by the target-separation condition *α*^2^*N*_eff_ ≳ *Z*_*ϵ*_, which holds by hypothesis (see eq. S12.92). Combining these conditions, we obtain the unified sufficient condition

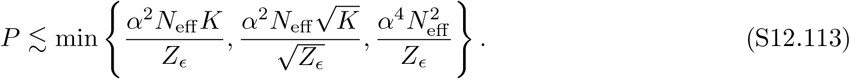

Taking *c*_cap_ *>* 0 sufficiently small to absorb the absolute constants implicit above, every *P* satisfying (S12.93) obeys AUC(*P* |***Y***) ≥ 1 − *ϵ* on the good event of Theorem 2. Since this event has probability at least 1 − *δ* = 1 − exp [−*γ* min{*P, K*}], this proves (S12.94).

### S12.3 Pause-Based Decoding

Next we analyze network capacity with the pause-based decoder Ω(***g***) = ∑_*k*_ Θ[*κ* − *g*^*k*^]. We allow *β* ∈ (0, 1), but restrict to the balanced-plasticity regime where 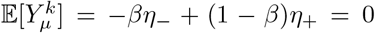 and L1 constraint *η*_−_ + *η*_+_ = Λ. We may then encode LTD events in a matrix ***M*** ∈ {0, 1}^*P ×K*^ with 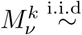 Bernoulli(*β*) and write

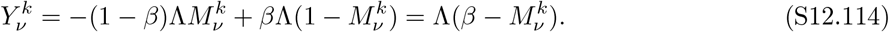

Setting *α* ≡ 1 − Δ_*m*_ *>* 0 under the Gaussian statistics in (S12.1)–(S12.2), the marginal response of PC *k* to a novel input has variance

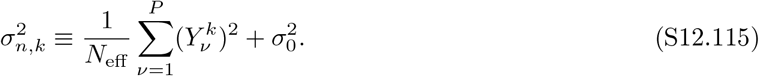

For a learned input from cluster *µ*, the corresponding marginal mean and variance are

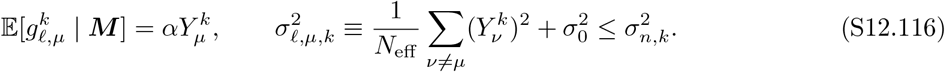

In particular, if 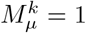, then 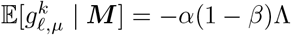. In this section, we will set 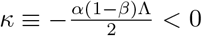, so that it lies at midpoint of baseline and expected depressed PC input current.

#### Theorem 3

(Pause-count AUC guarantee). *Consider the setup above. Fix ϵ* ∈ (0, 1) *and γ >* 0, *and define*

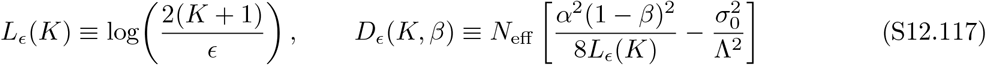

*Suppose the following conditions hold:*

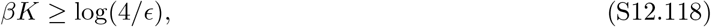

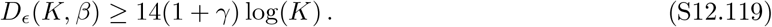

*Then for all P which satisfy*

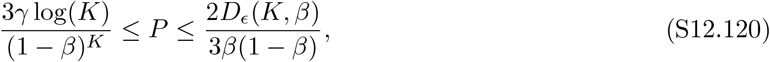

*we have*

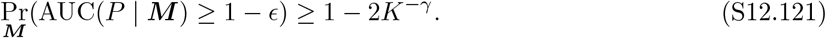

*Proof*. The matrix ***M*** is sampled once at training time and then held fixed. We first control, uniformly over PCs, the crosstalk power accumulated during training. For each PC *k*, define

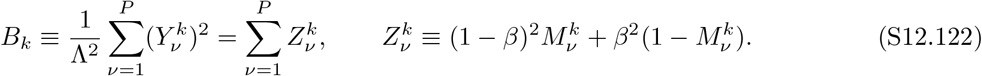

The variables 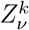 are independent over *ν*, with 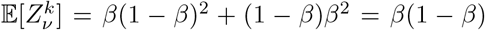. Hence *µ*_*B*_ ≡ E[*B*_*k*_] = *Pβ*(1−*β*). Since 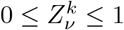, we have 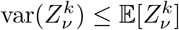 and 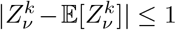. Bernstein’s inequality therefore gives, for every *D > µ*_*B*_,

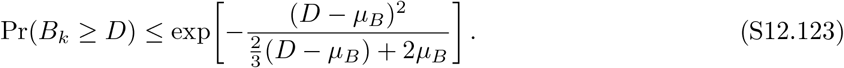

For all 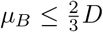, the right side is bounded above by 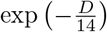. Also define

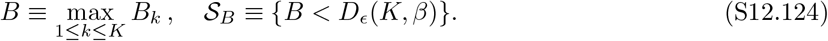

A union bound over *k* then gives that

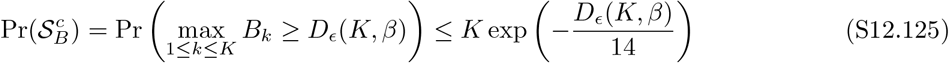

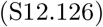

holds, subject to the condition

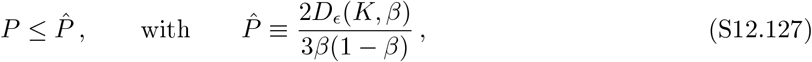

which is equivalent to the right inequality in (S12.120). From (S12.119), we also have

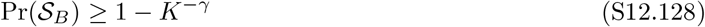

We next control the fraction of patterns for which the network received no LTD signal during training. Define the coverages *S*_*µ*_ and number *U* of uncovered patterns as follows:

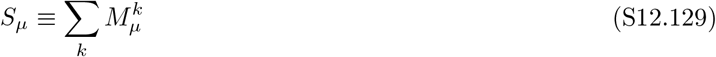

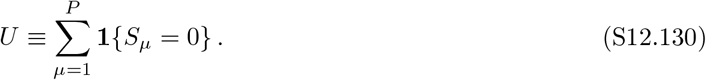

Note that *U* is a sum over *P* independent (quenched) random Bernoulli variables. The standard multiplicative Chernoff bound then gives

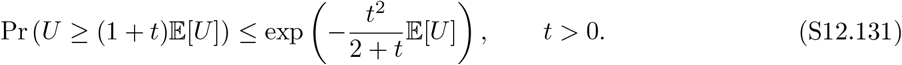

Define the event 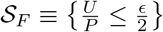. We then have

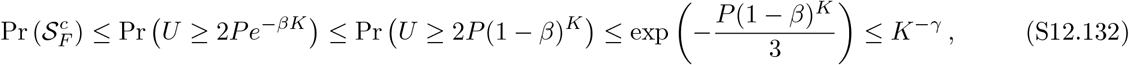

where the first inequality follows from condition (S12.118) the second from the fact that (1 − *β*)^*K*^ ≤ *e*^−*βK*^, the third from the Chernoff bound with *t* = 1 and E[*U*] = *P* (1 − *β*)^*K*^, and the last from the left inequality in (S12.120). Combining (S12.132) and (S12.128) using a union bound on the failure events 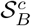 and 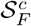 gives

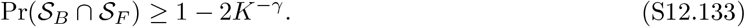

Next, we condition on a fixed realization ***M*** ∈ *S*_*B*_ ∩ *S*_*F*_. By (S12.115), every PC then satisfies

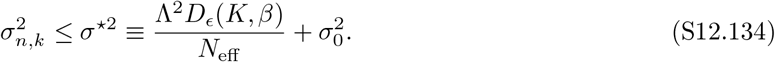

We nex t apply (S12.40) with 0 *< T <* 1, giving, because Ω(***g***) is a nonnegative integer, 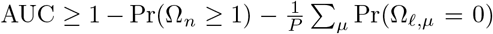. We first upper-bound the false-positive rate Pr(Ω_*n*_ ≥ 1). Since *κ <* 0, it follows from (S12.134) that

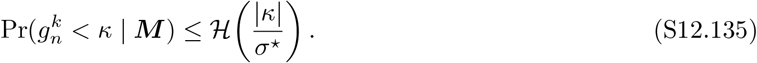

A union bound over the PCs yields

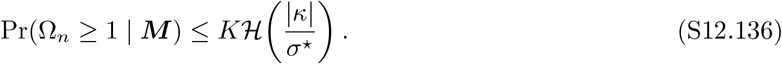

To bound the false-negative probability for the fixed realization ***M*** ∈ *S*_*B*_ ∩ *S*_*F*_, define the uncovered and covered pattern sets

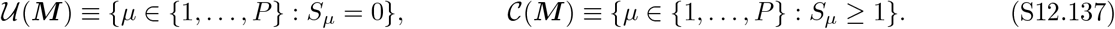

Because ***M*** ∈ *S*_*F*_, we have

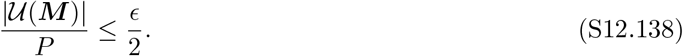

Splitting the average over uncovered and covered patterns gives

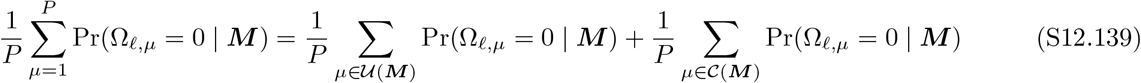

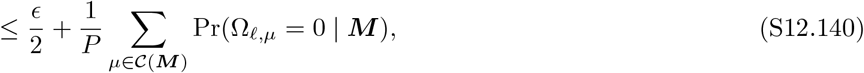

where we used the trivial bound Pr(Ω_*ℓ,µ*_ = 0 | ***M***) ≤ 1 for uncovered patterns. Now fix any *µ* ∈ *C*(***M***). Since *S* ≥ 1, there exists at least one *k* such that 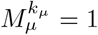. Conditional on ***M***, the input current 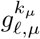 is Gaussian with mean and variance satisfying

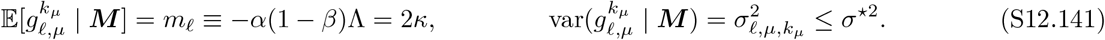

Since *κ* − *m*_*ℓ*_ = |*κ*|, it follows that

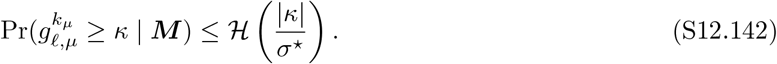

Moreover, if Ω_*ℓ,µ*_ = 0, then every PC lies above threshold, and in particular 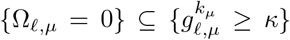. Therefore, for every 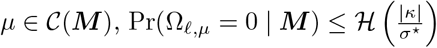. Substituting this bound into (S12.140) gives

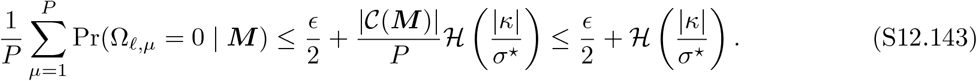

Substituting (S12.136) and (S12.143) into (S12.40) yields

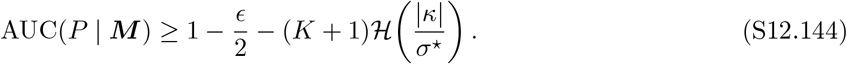

When ***M*** ∈ *S*_*B*_ ∩ *S*_*F*_, it is therefore sufficient for AUC(*P* | ***M***) ≥ 1 − *ϵ* that

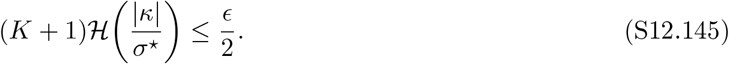

Using

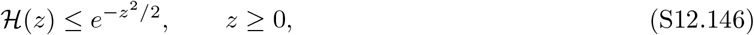

this condition is implied by

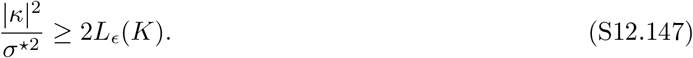

Because |*κ*|^2^ = *α*^2^(1 − *β*)^2^Λ^2^*/*4, the last inequality is equivalent to

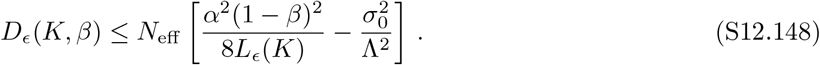

Our choice for *D*_*ϵ*_(*K, β*) (and therefore 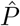) is therefore the maximum choice which satisfies this requirement. Thus for every *P* satisfying (S12.120), AUC(*P* | ***M***) ≥ 1 − *ϵ* for all ***M*** on *S*_*B*_ ∩ *S*_*F*_. Furthermore, (S12.133) shows that the event *S*_*B*_ ∩ *S*_*F*_ occurs with probability at least 1 − 2*K*^−*γ*^, completing the proof.

#### Corollary 2

(Capacity scaling under sparse plasticity). *Fix ϵ* ∈ (0, 1), *γ >* 0, *α >* 0, *and c* ≥ log(4*/ϵ*) *Suppose σ*_0_ = 0, *and define*

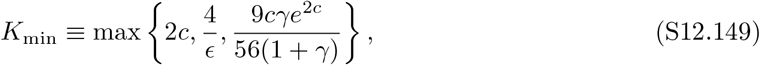

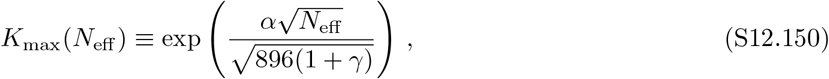

*and assume that K*_min_ ≤ *K* ≤ *K*_max_(*N*_eff_). *Suppose that we set* 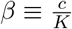 *for every K in this admissible range. Then, for every K in the admissible range and every integer P satisfying*

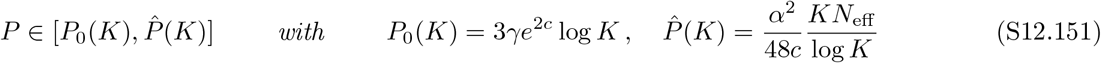

*we have*

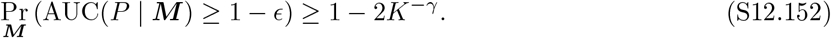

*Moreover, as K, N*_eff_ → ∞ *in the admissible range, the upper endpoint in* (S12.151) *gives the capacity lower-bound scaling*

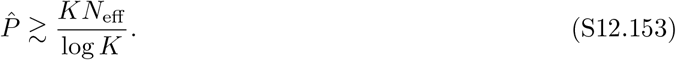

*Proof*. Because *K* ≥ *K*_min_ ≥ max{2*c*, 4*/ϵ*}, we have 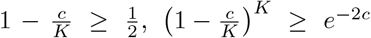, and 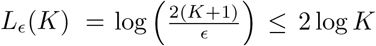. Plugging *β* = *c/K* and *σ*_0_ = 0 into (S12.117) and using the preceding bounds, we get

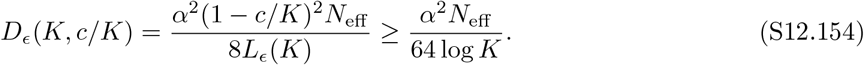

The necessary condition *D*_*ϵ*_(*K, c/K*) ≥ 14(1 + *γ*) log *K* in Theorem 3 follows easily from the upper bound *K* ≤ *K*_max_(*N*_eff_). Also, *βK* = *c* ≥ log(4*/ϵ*), verifying condition (S12.118) of Theorem 3. Substituting *β* = *c/K* into the lower load endpoint of Theorem 3 we have

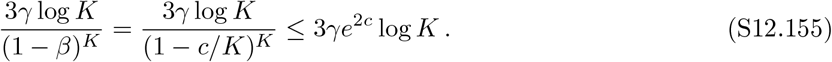

Substituting into the upper load condition gives

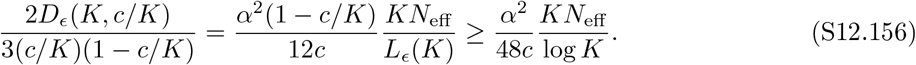

Together, (S12.155) and (S12.156) show that the hypothesis (S12.151) is a sufficient condition for the bounds on *P* in Theorem 3 (given in eq. S12.120). Theorem 3 therefore gives

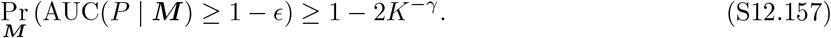

Finally, the conditions *K* ≤ *K*_max_(*N*_eff_) and *K* ≥ *K*_min_ imply

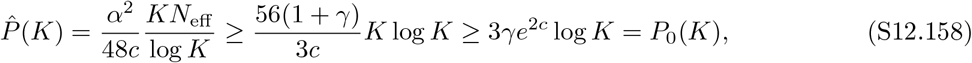

so the permissible *P*-interval is nonempty. Moreover, the ratio of the upper and lower bounds on the *P* interval is given by

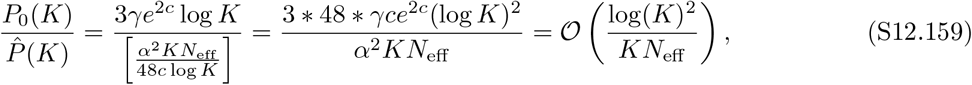

so that the size of the permissible interval is nonempty and growing in size for sufficiently large *K, N*_eff_.

#### Remark

**Capacity Scaling with width** Suppose *N*_*s*_, *N*_*m*_ and *K* and jointly scaled with a “width” parameter *w*, such that *N*_eff_ = *c*_1_*w* and *K* = *c*_2_*w*. Then for sufficiently large *w* we will have *K > K*_min_ and, because *N*_eff_ ∝ *K* ∝ *w, N*_eff_ *>* 896(1 + *γ*)(log *K*)^2^*/α*^2^ so that *K* ≤ *K*_max_(*N*_eff_) in Corollary 2. In this scaling regime we therefore have the capacity lower bound:

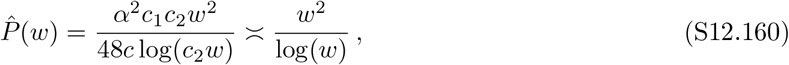

as stated in the main text.

## Footnotes

1 claude.ai

2 chatgpt.com

## References

[1] David A McCormick and Richard F Thompson. Cerebellum: essential involvement in the classically conditioned eyelid response. Science, 223(4633):296–299, January 1984.

[2] D A McCormick and R F Thompson. Neuronal responses of the rabbit cerebellum during acquisition and performance of a classically conditioned nictitating membrane-eyelid response. J. Neurosci., 4(11):2811–2822, November 1984.

[3] Michiel M ten Brinke, Henk-Jan Boele, Jochen K Spanke, Jan-Willem Potters, Katja Kornysheva, Peer Wulff, Anna C H G IJpelaar, Sebastiaan K E Koekkoek, and Chris I De Zeeuw. Evolving models of pavlovian conditioning: Cerebellar cortical dynamics in awake behaving mice. Cell Rep., 13(9):1977–1988, December 2015.

[4] Henk-Jan Boele, Saša Peter, Michiel M Ten Brinke, Lucas Verdonschot, Anna C H IJpelaar, Dimitris Rizopoulos, Zhenyu Gao, Sebastiaan K E Koekkoek, and Chris I De Zeeuw. Impact of parallel fiber to purkinje cell long-term depression is unmasked in absence of inhibitory input. Sci. Adv., 4(10):eaas9426, October 2018.

[5] Dylan J Calame, Matthew I Becker, and Abigail L Person. Cerebellar associative learning underlies skilled reach adaptation. Nat. Neurosci., 26(6):1068–1079, June 2023.

[6] David J Herzfeld, Yoshiko Kojima, Robijanto Soetedjo, and Reza Shadmehr. Encoding of action by the purkinje cells of the cerebellum. Nature, 526(7573):439–442, October 2015.

[7] David J Herzfeld, Yoshiko Kojima, Robijanto Soetedjo, and Reza Shadmehr. Encoding of error and learning to correct that error by the purkinje cells of the cerebellum. Nat. Neurosci., 21 (5):736–743, May 2018.

[8] Ehsan Sedaghat-Nejad, Jay S Pi, Paul Hage, Mohammad Amin Fakharian, and Reza Shad-mehr. Synchronous spiking of cerebellar purkinje cells during control of movements. Proc. Natl. Acad. Sci. U. S. A., 119(14):e2118954119, April 2022.

[9] Reza Shadmehr. Population coding in the cerebellum: a machine learning perspective. J. Neurophysiol., 124(6):2022–2051, December 2020.

[10] David J Herzfeld, Mati Joshua, and Stephen G Lisberger. Rate versus synchrony codes for cerebellar control of motor behavior. Neuron, 111(15):2448–2460.e6, August 2023.

[11] Mohammad Amin Fakharian, Alden M Shoup, Paul Hage, Hisham Y Elseweifi, and Reza Shadmehr. A vector calculus for neural computation in the cerebellum. bioRxivorg, page 2024.11.14.623565, November 2024.

[12] Jennifer L Raymond and Javier F Medina. Computational principles of supervised learning in the cerebellum. Annu. Rev. Neurosci., 41(Volume 41, 2018):233–253, July 2018.

[13] M Ito, M Sakurai, and P Tongroach. Climbing fibre induced depression of both mossy fibre responsiveness and glutamate sensitivity of cerebellar purkinje cells. J. Physiol., 324(1):113– 134, March 1982.

[14] D Marr. A theory of cerebellar cortex. J. Physiol., 202(2):437–470, June 1969.

[15] J Albus. A theory of cerebellar function. Bellman Prize in Mathematical Biosciences, 10(1-2): 25–61, February 1971.

[16] Ito. Cerebellum and Neural Control. Lippincott Williams and Wilkins, Philadelphia, PA, July 1984.

[17] Baktash Babadi and Haim Sompolinsky. Sparseness and expansion in sensory representations. Neuron, 83(5):1213–1226, September 2014.

[18] Ashok Litwin-Kumar, Kameron Decker Harris, Richard Axel, Haim Sompolinsky, and L F Abbott. Optimal degrees of synaptic connectivity. Neuron, 93(5):1153–1164.e7, March 2017.

[19] Samuel P Muscinelli, Mark J Wagner, and Ashok Litwin-Kumar. Optimal routing to cerebellum-like structures. Nat. Neurosci., 26(9):1630–1641, September 2023.

[20] Ilker Ozden, Megan R Sullivan, H Megan Lee, and Samuel S-H Wang. Reliable coding emerges from coactivation of climbing fibers in microbands of cerebellar purkinje neurons. J. Neurosci., 29(34):10463–10473, August 2009.

[21] Takayuki Michikawa, Takamasa Yoshida, Satoshi Kuroki, Takahiro Ishikawa, Shinji Kakei, Ryo Kimizuka, Atsushi Saito, Hideo Yokota, Akinobu Shimizu, Shigeyoshi Itohara, and Atsushi Miyawaki. Distributed sensory coding by cerebellar complex spikes in units of cortical segments. Cell Rep., 37(6):109966, November 2021.

[22] Olov Oscarsson. Functional units of the cerebellum-sagittal zones and microzones. Trends Neurosci., 2:143–145, January 1979.

[23] Martin Garwicz, Carl-Fredrik Ekerot, and Henrik Jörntell. Organizational principles of cerebellar neuronal circuitry. News Physiol. Sci., 13(1):26–32, February 1998.

[24] I Sugihara, H S Wu, and Y Shinoda. The entire trajectories of single olivocerebellar axons in the cerebellar cortex and their contribution to cerebellar compartmentalization. J. Neurosci., 21(19):7715–7723, October 2001.

[25] Izumi Sugihara and Yoshikazu Shinoda. Molecular, topographic, and functional organization of the cerebellar cortex: a study with combined aldolase C and olivocerebellar labeling. J. Neurosci., 24(40):8771–8785, October 2004.

[26] G Brochu, L Maler, and R Hawkes. Zebrin II: a polypeptide antigen expressed by a subset of purkinje cells in the cerebellar cortex. Journal of Comparative Neurology, 291(4):538–552, 1990.

[27] Richard Apps and Richard Hawkes. Cerebellar cortical organization: a one-map hypothesis. Nat. Rev. Neurosci., 10(9):670–681, September 2009.

[28] Izumi Sugihara, Hirofumi Fujita, Jie Na, Pham Nguyen Quy, Bing-Yang Li, and Daisuke Ikeda. Projection of reconstructed single purkinje cell axons in relation to the cortical and nuclear aldolase C compartments of the rat cerebellum. J. Comp. Neurol., 512(2):282–304, January 2009.

[29] Shinichiro Tsutsumi, Maya Yamazaki, Taisuke Miyazaki, Masahiko Watanabe, Kenji Sakimura, Masanobu Kano, and Kazuo Kitamura. Structure-function relationships between aldolase C/zebrin II expression and complex spike synchrony in the cerebellum. J. Neurosci., 35(2):843–852, January 2015.

[30] François G C Blot, Joshua J White, Amy van Hattem, Licia Scotti, Vaishnavi Balaji, Youri Adolfs, R Jeroen Pasterkamp, Chris I De Zeeuw, and Martijn Schonewille. Purkinje cell microzones mediate distinct kinematics of a single movement. Nat. Commun., 14(1):4358, July 2023.

[31] Richard Apps, Richard Hawkes, Sho Aoki, Fredrik Bengtsson, Amanda M Brown, Gang Chen, Timothy J Ebner, Philippe Isope, Henrik Jörntell, Elizabeth P Lackey, Charlotte Lawrenson, Bridget Lumb, Martijn Schonewille, Roy V Sillitoe, Ludovic Spaeth, Izumi Sugihara, Antoine Valera, Jan Voogd, Douglas R Wylie, and Tom J H Ruigrok. Cerebellar modules and their role as operational cerebellar processing units: A consensus paper [corrected]. Cerebellum, 17 (5):654–682, October 2018.

[32] Hirofumi Fujita, Takashi Kodama, and Sascha du Lac. Modular output circuits of the fastigial nucleus for diverse motor and nonmotor functions of the cerebellar vermis. eLife, 9, July 2020.

[33] Abigail L Person and Indira M Raman. Purkinje neuron synchrony elicits time-locked spiking in the cerebellar nuclei. Nature, 481(7382):502–505, December 2011.

[34] Tycho M Hoogland, Jornt R De Gruijl, Laurens Witter, Cathrin B Canto, and Chris I De Zeeuw. Role of synchronous activation of cerebellar purkinje cell ensembles in multi-joint movement control. Curr. Biol., 25(9):1157–1165, May 2015.

[35] I E Brown and J M Bower. Congruence of mossy fiber and climbing fiber tactile projections in the lateral hemispheres of the rat cerebellum. J. Comp. Neurol., 429(1):59–70, January 2001.

[36] J P Welsh, E J Lang, I Suglhara, and R Llinás. Dynamic organization of motor control within the olivocerebellar system. Nature, 374(6521):453–457, March 1995.

[37] S Kitazawa, T Kimura, and P B Yin. Cerebellar complex spikes encode both destinations and errors in arm movements. Nature, 392(6675):494–497, April 1998.

[38] P F Gilbert and W T Thach. Purkinje cell activity during motor learning. Brain Res., 128 (2):309–328, June 1977.

[39] L S Stone and S G Lisberger. Detection of tracking errors by visual climbing fiber inputs to monkey cerebellar flocculus during pursuit eye movements. Neurosci Lett, 72:163–168, 1986.

[40] Michiel Coesmans, John T Weber, Chris I De Zeeuw, and Christian Hansel. Bidirectional parallel fiber plasticity in the cerebellum under climbing fiber control. Neuron, 44(4):691–700, November 2004.

[41] Alexandre Mathy, Sara S N Ho, Jenny T Davie, Ian C Duguid, Beverley A Clark, and Michael Häusser. Encoding of oscillations by axonal bursts in inferior olive neurons. Neuron, 62(3): 388–399, May 2009.

[42] Heather K Titley, Mikhail Kislin, Dana H Simmons, Samuel S-H Wang, and Christian Hansel. Complex spike clusters and false-positive rejection in a cerebellar supervised learning rule. J. Physiol., 597(16):4387–4406, August 2019.

[43] Court Hull and Wade G Regehr. The cerebellar cortex. Annu. Rev. Neurosci., 45(Volume 45, 2022):151–175, July 2022.

[44] Elizabeth P Lackey, Luis Moreira, Aliya Norton, Marie E Hemelt, Tomas Osorno, Tri M Nguyen, Evan Z Macosko, Wei-Chung Allen Lee, Court A Hull, and Wade G Regehr. Specialized connectivity of molecular layer interneuron subtypes leads to disinhibition and synchronous inhibition of cerebellar purkinje cells. Neuron, 112(14):2333–2348.e6, July 2024.

[45] E P Lackey, A Norton, L Moreira, C Gaynor,A A Lee, and W G Regehr. Purkinje cell collaterals preferentially target a subtype of molecular layer interneuron. bioRxiv, page 2025.07.17.665322, July 2025.

[46] Tri M Nguyen, Logan A Thomas, Jeff L Rhoades, Ilaria Ricchi, Xintong Cindy Yuan, Arlo Sheridan, David G C Hildebrand, Jan Funke, Wade G Regehr, and Wei-Chung Allen Lee. Structured cerebellar connectivity supports resilient pattern separation. Nature, 613(7944): 543–549, January 2023.

[47] Maoz Shamir and Haim Sompolinsky. Nonlinear population codes. Neural Comput., 16(6): 1105–1136, June 2004.

[48] Chris I De Zeeuw. Bidirectional learning in upbound and downbound microzones of the cerebellum. Nat. Rev. Neurosci., 22(2):92–110, February 2021.

[49] Ehsan Sedaghat-Nejad, Mohammad Amin Fakharian, Jay Pi, Paul Hage, Yoshiko Kojima, Robijanto Soetedjo, Shogo Ohmae, Javier F Medina, and Reza Shadmehr. P-sort: an open-source software for cerebellar neurophysiology. Journal of Neurophysiology, 126(4):1055–1075, 2021.

[50] Marjorie Xie, Samuel P Muscinelli, Kameron Decker Harris, and Ashok Litwin-Kumar. Task-dependent optimal representations for cerebellar learning. Elife, 12, September 2023.

[51] P Dayan and D J Willshaw. Optimising synaptic learning rules in linear associative memories. Biol. Cybern., 65(4):253–265, August 1991.

[52] B P Graham. Pattern recognition in a compartmental model of a CA1 pyramidal neuron. Network, 12(4):473–492, November 2001.

[53] Joy T Walter and Kamran Khodakhah. The advantages of linear information processing for cerebellar computation. Proc. Natl. Acad. Sci. U. S. A., 106(11):4471–4476, March 2009.

[54] Thomas G Dietterich. Ensemble methods in machine learning. In Multiple Classifier Systems, Lecture Notes in Computer Science, pages 1–15. Springer Berlin Heidelberg, Berlin, Heidelberg, 2000.

[55] Jacob Andreas Zavatone-Veth. Chapter 7. In Statistical Mechanics of Bayesian Inference and Learning in Neural Networks, chapter 7. ProQuest Dissertations & Theses, 2024.

[56] D Jaeger and J M Bower. Synaptic control of spiking in cerebellar purkinje cells: dynamic current clamp based on model conductances. J. Neurosci., 19(14):6090–6101, July 1999.

[57] Dan-Anders Jirenhed, Fredrik Bengtsson, and Germund Hesslow. Acquisition, extinction, and reacquisition of a cerebellar cortical memory trace. J. Neurosci., 27(10):2493–2502, March 2007.

[58] Dan-Anders Jirenhed and Germund Hesslow. Are purkinje cell pauses drivers of classically conditioned blink responses? Cerebellum, 15(4):526–534, August 2016.

[59] Sungho Hong, Mario Negrello, Marc Junker, Aleksandra Smilgin, Peter Thier, and Erik De Schutter. Multiplexed coding by cerebellar purkinje neurons. Elife, 5, July 2016.

[60] Andrea Giovannucci, Aleksandra Badura, Ben Deverett, Farzaneh Najafi, Talmo D Pereira, Zhenyu Gao, Ilker Ozden, Alexander D Kloth, Eftychios Pnevmatikakis, Liam Paninski, Chris I De Zeeuw, Javier F Medina, and Samuel S-H Wang. Cerebellar granule cells acquire a widespread predictive feedback signal during motor learning. Nat. Neurosci., 20(5): 727–734, May 2017.

[61] Laura D Knogler, Daniil A Markov, Elena I Dragomir, Vilim Štih, and Ruben Portugues. Sensorimotor representations in cerebellar granule cells in larval zebrafish are dense, spatially organized, and non-temporally patterned. Curr. Biol., 27(9):1288–1302, May 2017.

[62] Aleksandra Badura and Chris I De Zeeuw. Cerebellar granule cells: Dense, rich and evolving representations. Curr. Biol., 27(11):R415–R418, June 2017.

[63] Frederic Lanore, N Alex Cayco-Gajic, Harsha Gurnani, Diccon Coyle, and R Angus Silver. Cerebellar granule cell axons support high-dimensional representations. Nat. Neurosci., 24 (8):1142–1150, August 2021.

[64] Nicolas Brunel, Vincent Hakim, and Jean-Pierre Nadal. From the perceptron to the cerebellum. arXiv [q-bio.NC], May 2025.

[65] Chulhee Yun, Suvrit Sra, and Ali Jadbabaie. Small ReLU networks are powerful memorizers: a tight analysis of memorization capacity. arXiv [cs.LG], October 2018.

[66] Roman Vershynin. Memory capacity of neural networks with threshold and rectified linear unit activations. SIAM J. Math. Data Sci., 2(4):1004–1033, January 2020.

[67] Yunliang Zang and Erik De Schutter. The cellular electrophysiological properties underlying multiplexed coding in purkinje cells. J. Neurosci., 41(9):1850–1863, March 2021.

[68] C D Aizenman and D J Linden. Regulation of the rebound depolarization and spontaneous firing patterns of deep nuclear neurons in slices of rat cerebellum. J. Neurophysiol., 82(4): 1697–1709, October 1999.

[69] J Laine and H Axelrad. Lugaro cells target basket and stellate cells in the cerebellar cortex. Neuroreport, 9(10):2399–2403, July 1998.

[70] Abigail L Person and Indira M Raman. Synchrony and neural coding in cerebellar circuits. Front. Neural Circuits, 6:97, December 2012.

[71] Ke Zhang, Zhen Yang, Michael A Gaffield, Garrett G Gross, Don B Arnold, and Jason M Christie. Molecular layer disinhibition unlocks climbing-fiber-instructed motor learning in the cerebellum. bioRxivorg, page 2023.08.04.552059, August 2023.

[72] Andrew K Wise, Nadia L Cerminara, Dilwyn E Marple-Horvat, and Richard Apps. Mechanisms of synchronous activity in cerebellar purkinje cells: Synchrony in the cerebellum. J. Physiol., 588(Pt 13):2373–2390, July 2010.

[73] Tianyu Tang, Timothy A Blenkinsop, and Eric J Lang. Complex spike synchrony dependent modulation of rat deep cerebellar nuclear activity. Elife, 8, January 2019.

[74] J F Medina and M D Mauk. Computer simulation of cerebellar information processing. Nat. Neurosci., 3 Suppl(S11):1205–1211, November 2000.

[75] Paul Dean and John Porrill. Adaptive-filter models of the cerebellum: computational analysis. Cerebellum, 7(4):567–571, 2008.

[76] Kameron Decker Harris. Additive function approximation in the brain. arXiv [cs.NE], September 2019.

[77] N Alex Cayco-Gajic, Claudia Clopath, and R Angus Silver. Sparse synaptic connectivity is required for decorrelation and pattern separation in feedforward networks. Nat. Commun., 8 (1):1116, October 2017.

[78] William Dorrell and Peter E Latham. Mapping connectomic structure to function(s) in cerebellar-like networks using kernel regression. arXiv [q-bio.NC], January 2026.

[79] Nicolas Brunel, Vincent Hakim, Philippe Isope, Jean-Pierre Nadal, and Boris Barbour. Optimal information storage and the distribution of synaptic weights: perceptron versus purkinje cell. Neuron, 43(5):745–757, September 2004.

[80] Claudia Clopath and Nicolas Brunel. Optimal properties of analog perceptrons with excitatory weights. PLoS Comput. Biol., 9(2):e1002919, February 2013.

[81] Blake Bordelon and Cengiz Pehlevan. Population codes enable learning from few examples by shaping inductive bias. Elife, 11, December 2022.

[82] Naoki Hiratani. Local gated-hebbian learning of deep cerebellar networks with quadratic classification capacity. bioRxiv, page 2026.04.17.718957, April 2026.

[83] Adrian Holtrup, Ramin Khajeh, and Wei-Chung Allen Lee. Random innervation of cerebellar purkinje cells as a substrate for diverse representational learning. bioRxiv, page 2026.06.26.734775, June 2026.

[84] Lyudmila Kushnir and Stefano Fusi. Neural classifiers with limited connectivity and recurrent readouts. J. Neurosci., 38(46):9900–9924, November 2018.

[85] Javier F Medina and Stephen G Lisberger. Variation, signal, and noise in cerebellar sensory-motor processing for smooth-pursuit eye movements. J. Neurosci., 27(25):6832–6842, June 2007.

[86] Spencer T Brown, Mauricio Medina-Pizarro, Meghana Holla, Christopher E Vaaga, and Indira M Raman. Simple spike patterns and synaptic mechanisms encoding sensory and motor signals in purkinje cells and the cerebellar nuclei. Neuron, 112(11):1848–1861.e4, June 2024.

[87] Michael A Long, Michael R Deans, David L Paul, and Barry W Connors. Rhythmicity without synchrony in the electrically uncoupled inferior olive. J. Neurosci., 22(24):10898– 10905, December 2002.

[88] Simon R Schultz, Kazuo Kitamura, Arthur Post-Uiterweer, Julija Krupic, and Michael Häusser. Spatial pattern coding of sensory information by climbing fiber-evoked calcium signals in networks of neighboring cerebellar purkinje cells. J. Neurosci., 29(25):8005–8015, June 2009.

[89] Joseph Chaumont, Nicolas Guyon, Antoine M Valera, Guillaume P Dugué, Daniela Popa, Paikan Marcaggi, Vanessa Gautheron, Sophie Reibel-Foisset, Stéphane Dieudonné, Aline Stephan, Michel Barrot, Jean-Christophe Cassel, Jean-Luc Dupont, Frédéric Doussau, Bernard Poulain, Fekrije Selimi, Clément Léna, and Philippe Isope. Clusters of cerebellar purkinje cells control their afferent climbing fiber discharge. Proc. Natl. Acad. Sci. U. S. A., 110(40):16223–16228, October 2013.

[90] Kim M Gruver, Jenny W Y Jiao, Eviatar Fields, Sen Song, Per Jesper Sjöström, and Alanna J Watt. Structured connectivity in the output of the cerebellar cortex. Nat. Commun., 15(1):5563, July 2024.

[91] Andrew M Davidson, Shivam Kaushik, and Toshihide Hige. Dopamine-dependent plasticity is heterogeneously expressed by presynaptic calcium activity across individual boutons of the drosophila mushroom body. eNeuro, 10(10), October 2023.

[92] Yujie Wu and Wolfgang Maass. A simple model for behavioral time scale synaptic plasticity (BTSP) provides content addressable memory with binary synapses and one-shot learning. Nat. Commun., 16(1):342, January 2025.

[93] Leo Breiman. Bagging predictors. Mach. Learn., 24(2):123–140, August 1996.

[94] Leo Breiman. Random forests. Mach. Learn., 45(1):5–32, October 2001.

[95] J Neyman and E S Pearson. IX. on the problem of the most efficient tests of statistical hypotheses. Philos. Trans. R. Soc. Lond., 231(694-706):289–337, February 1933.

[96] J A Hanley and B J McNeil. The meaning and use of the area under a receiver operating characteristic (ROC) curve. Radiology, 143(1):29–36, April 1982.

[97] J M Wozencraft and I M Jacobs. Principles of Communications Engineering. John Wiley & Sons, Nashville, TN, January 1966.

[98] Adrian Holtrup, Ramin Khajeh, Wei-Chung Allen Lee. Parallel fiber subsampling in the cerebellar cortex as a substrate for diverse learning. COSYNE, March 2026.

[99] Shu Ho, Rebecca Lajaunie, Marion Lerat, Mickaël Le, Valérie Crépel, Karine Loulier, Jean Livet, Jean-Pierre Kessler, and Païkan Marcaggi. A stable proportion of purkinje cell inputs from parallel fibers are silent during cerebellar maturation. Proc. Natl. Acad. Sci. U. S. A., 118(45):e2024890118, November 2021.

[100] B Laurent and P Massart. Adaptive estimation of a quadratic functional by model selection. Ann. Stat., 28(5):1302–1338, October 2000.

